# Proteome-wide probing of the dual NMT-dependent myristoylation tradeoff unveils potent, mechanism-based suicide inhibitors

**DOI:** 10.1101/2021.06.22.449375

**Authors:** Frédéric Rivière, Cyril Dian, Rémi F. Dutheil, Carmela Giglione, Thierry Meinnel

## Abstract

N-myristoyltransferases (NMTs) catalyze protein myristoylation, a major and ubiquitous lipid modification. Originally thought to modify only N-terminal glycine α-amino groups (G-myristoylation), NMTs are now known to modify lysine ε-amino groups (K-myristoylation), the significance of which is uncertain. Here we exploited systematic structural proteomics analyses and a novel pipeline involving the *Shigella* IpaJ protease to discriminate K- and G-myristoylation with unprecedented accuracy and identify the specific features driving each modification. NMT-dependent K-myristoylation occurs post-translationally and only on lysines 1, 2, or 3 following G-myristoylation or caspase cleavage. Direct interactions between the substrate’s reactive amino group and the NMT catalytic base slow K-myristoylation catalysis. IpaJ unmasked novel K-myristoylation sites in a dozen human proteins. The unique properties of NMT-driven K-myristoylation allowed us to design potent, mechanism-based suicide NMT inhibitors. These analyses unravel the respective paths towards K-myristoylation, G-myristoylation, or NMT inhibition, which rely on a very subtle tradeoff embracing the chemical landscape around the reactive group.

**SIGNIFICANCE STATEMENT:** We report the specific and unique elements guiding N-myristoyltransferase to either alpha or epsilon myristoylation, allowing us to establish the post-translational nature of N-myristoyltransferase-dependent lysine myristoylation and design novel, potent N-myristoyltransferase inhibitors.

## INTRODUCTION

Myristoylation is an essential lipidation that adds a C:14:0 fatty acid to proteins in all eukaryotes including pathogens^1–4^. The lipid moiety anchors soluble proteins to membranes, where they interact with partners to initiate signal transduction^5–7^. N-myristoyltransferases (NMTs; glycylpeptide N-tetradecanoyltransferases) are the only enzyme class known to catalyze myristoylation and, given their central role in pathobiology, are promising therapeutic targets^8–12^. NMTs were long thought to exclusively target proteins with an N-terminal glycine (G-myristoylation) usually arising from co-translational methionine excision^1,13^ or, less frequently, post-translation cleavage exposing new N-terminal glycines^14,15^. Proteome-wide approaches have delivered an exhaustive list of substrates undergoing co-translational G-myristoylation - the so-called G-myristoylome - covering approximately 2% of the human proteome (~600 proteoforms^16^). Additionally, high-resolution co-crystallography of human NMT1 with a number of reactive substrates displaying an N-terminal Gly^17^ has established a specific water channel in NMT that provides a water-mediated bond (Wat2) between the N-terminal group of the G-starting substrate and the C-terminal carboxy group (Q496) of NMT, which acts as the catalytic base for deprotonation and further reactivity^1,17^. These data have been instrumental in revealing that NMTs further support myristoylation of lysine side chains (K-myristoylation), significantly expanding the range of known substrates^17,18^. The mammalian small G-protein ARF6, which features a K3 following an N-terminal G2, is the only protein currently known to undergo both N-terminal G- and K-myristoylation^18^, allowing it to attach to plasma membranes when GTP is bound and permitting K-myristoylation by sirtuin deacylases upon GTP hydrolysis. In this cycle, NMT ensures the novel ARF6 K-myristoylation after GTP refueling.

NMT-dependent G-myristoylation occurs on the α-amino group of the target protein via an amide bond, creating an extremely stable attachment *in vivo* and resistance to chemical cleavage *in vitro*^19–21^. Accordingly, G-myristoylation was considered to be irreversible. However, the discovery of IpaJ, a pathogenic cysteine-dependent protease produced by several disease-causing bacteria including *Shigella flexneri*, challenged this four decade-old dogma^22^. IpaJ cleaves the N-terminal myristoylated G from several proteins. IpaJ relies on the presence of a G-myristoylated group embedded within a small dipeptide as a minimal chassis^23^, with strong specificity for C:14 chains over longer (C:16) or shorter (C:10) fatty acyl chains^23^. Since this study predated the discovery of NMT-driven K-myristoylation, known IpaJ substrates are limited *in vivo* to the small G-proteins of the ADP-ribosylation factor family (ARF and ARL) with the exception of ARF6^23,24^. Critically, it is unknown whether IpaJ contributes to the NMT-dependent K-myristoylome. One major challenge to establishing this lies in the difficulty in identifying and proving the NMT-dependent K-myristoylome and the type of myristoylation taking place, the latter usually relying on mass spectrometry (MS). Unequivocal MS/MS (MS2) spectra of K-myristoylation profiles are not available and, because several acyl hydrolases targeting K-myristoylation have been described including SIRT2, SIRT6, HDAC8, and HDAC11, their action might prevent the identification of K-myristoylated substrates *in vivo*. New approaches for characterizing the K-myristoylome are urgently required.

To this end, here we combine mass spectrometry, kinetic studies, *in silico* analysis, and crystallography of human NMT1 with synthetic peptides to assess the molecular basis of NMT-dependent K-myristoylation. In doing so, we provide an overview of its coverage in the human proteome. We reveal that K-myristoylation results from post-translational events as illustrated by IpaJ-induced cleavage triggering K-myristoylation of a number of targets. Additionally, we show that G-myristoylation outcompetes K-myristoylation, since G- and K-myristoylation share a similar recognition chassis at the NMT active site. Nevertheless, K-myristoylation features a number of unique recognition elements, based on which we design novel, selective, and potent mechanism-based suicide NMT inhibitors.

## RESULTS

### Co-translational G-myristoylation predominates over K-myristoylation on N-terminal GK-starting peptides

We previously reported unforeseen NMT-binding clefts accommodating peptide substrate side-chains (pockets 2-8 in **Fig. 1a**) and the N-terminal myristoylome based on the hypothesis that NMT-dependent myristoylation only occurs on glycines^16^. We and others later reported NMT-dependent K-myristoylation on position 3 lysines^17,18^. An updated NMT reaction scheme with the two myristoylation types is displayed in **Supplementary Fig. 1**.

**Fig. 1.**
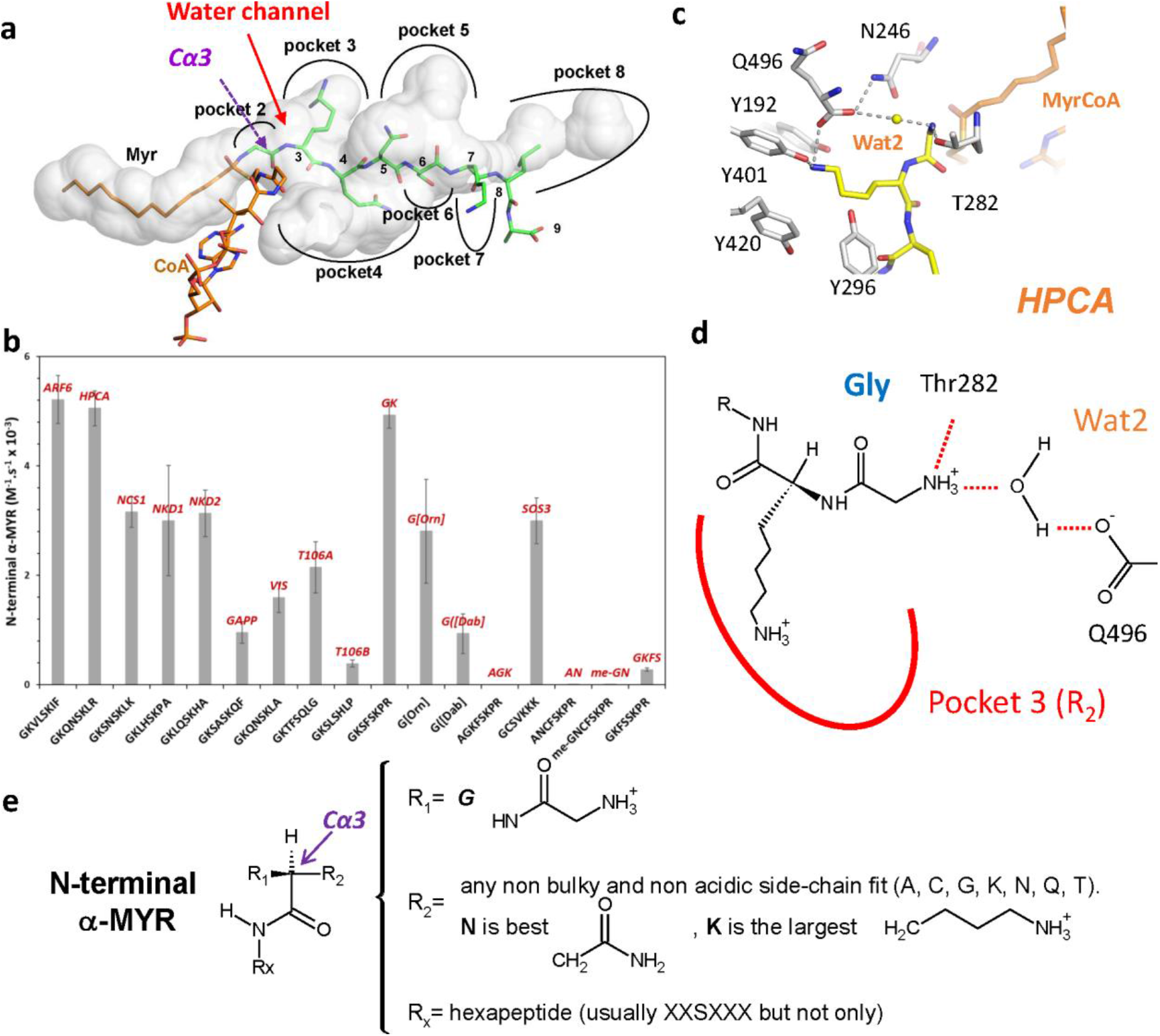
G-myristoylation rules in the context of GK-starting substrates. All peptides are derived from the GK peptide unless otherwise stated (see short name on top of panel a). (**a**) Crystal structure of the HPCA peptide with HsNMT; the various pockets and their extents are displayed and Cα3 is indicated. (**b**) Catalytic efficiencies of GK-starting peptides (see all parameters in **Supplementary Table 1a**). (**c**) 3D details of the active site organization of the HPCA complex. (**d**) Planar representation of panel c. (**e**) Types of chemical groups allowed around Cα3 leading to classic N-terminal myristoylation with G acceptor residue at R_1_ (G-myristoylation).

As a model peptide with a K3 (G_2_KSFSKPR; reference, GK) was an efficient NMT substrate (**Fig. 1b** and **Supplementary Table 1a**), we assessed which type of myristoylation (G- and/or K) *Homo sapiens* (Hs)NMT1 catalyzed on this peptide by MALDI-Tof-Tof including MS-MS (MS2). Together with kinetic analysis, K3 favored G-myristoylation in this model context (**Supplementary Table 1a, Supplementary Dataset 1** and **Fig. 1b**). We next examined the myristoylation status of all human proteins starting with N-terminal GKs, including the six closely related members of the HPCA/NCS1 family and ARF6. In all cases, only G-myristoylation was unambiguously retrieved (**Fig. 1b**, **Supplementary Dataset 1** and **Supplementary Table 1a**), with efficiencies in the same range as reported for the G-myristoylome, as exemplified by the SOS3 variant (**Fig. 1b**). Remarkably, most N-terminal GK-starting proteins harbored the S_6_K_7_ motif, which is known to facilitate myristoylation^25^. Indeed, the salt bridges of K7 with the three contiguous Ds of the NMT Ab-loop were recently shown to promote the movement of the Ab-loop from an open to a closed conformation (SuppFig.7 in ^17^), favoring catalysis through distortion of the MyrCoA thioester bond in the closed conformation and facilitating transition to the tetrahedral state.

A new crystal structure of HsNMT1 with a peptide corresponding to the N-terminus of HPCA revealed that the K3 side chain buries tightly in pocket 3 of NMT and that its positive charge bonds the catalytic base (**Fig. 1c**). Full occupation of pocket 3 with K3 slightly displaces the reactive alpha amino group of G2 from the MyrCoA thioester group whilst retaining the Wat2 interaction of the reactive N-terminal α-amino group required for G-myristoylation^17^ (**Fig. 1c,d**). This K3-induced displacement of G2 confirms that, at position 3, a K residue is less favorable to myristoylation than any amino acid with a smaller, uncharged side chain.

To examine whether GK-starting proteins have the same recognition rules as those in the G[^K] context (where ^X means any amino acid other than X)^16^, we analyzed HsNMT1’s capacity to acylate a series of changes in the N-terminal peptide derived from NCS1. Only G-myristoylation occurred when K7 was changed into G or S6 into A, although HsNMT1’s catalytic efficiency significantly decreased (**Supplementary Table 1b** and **Supplementary Dataset 1**). Therefore, the S6K7 motif is equally important for GK- and G[^K]-starting proteins, i.e., the role of S6 in substrate binding and K7 in the MyrCoA binding-induced conformational switch of the conserved flexible acidic Ab-loop of NMT is equal to that already reported and starting with G[^K]^17^. We next assessed whether modifying the side-chain length but retaining the basic character at position 7 impacted the myristoylation type. Shortening the basic side chain to ornithine (Orn, reduced by one CH_2_) or di-amino butyrate (Dab, reduced by two CH_2_s) had little impact on G-myristoylation (**Supplementary Table 1b**), suggesting maintenance of the salt bridge with the Ab-loop. Further shortening to di-amino propionate (Dap, short basic chain CH_2_NH_2_) or removal of the basic character with homocysteine (Hcy) significantly reduced catalytic efficiency but still promoted only G-myristoylation (**Supplementary Table 1b** and **Supplementary Dataset 1**).

To further investigate the rules governing myristoylation by NMT, we substituted the G2 with an A in the GK-context variant devoid of K3 (A_2_NCFSKPR; AN). There was unambiguous α-myristoylation by both MS and crystallographic analyses (**Extended Data 1a**, **Supplementary Dataset 1** and **Supplementary Table 1a**). Catalytic data indicated that the substrate was very inefficiently modified by NMT (**Fig. 1b**), and the crystal structure of AN confirmed that the active site was filled with a methyl addition, improperly orientating the reactive amino group. To decipher whether an A resulted in significant myristoylation in another sequence context, we substituted G2 with A in G_2_SSVSKKK, which was not myristoylated (**Supplementary Dataset 1**), clearly indicating that the S_6_K_7_ motif is insufficient for reactivity in the A2 context. The same was also true for an AGK variant (**Fig. 1b, Supplementary Table 1a**). In all cases, the G2A substitution dramatically reduced myristoylation but eventually led to G-myristoylation.

We therefore wondered whether the strong hindrance to the active site promoted by the additional methyl chain of A was similar if this group was grafted directly on the N-terminus. The N-methyl-GN peptide (meGN, meG is an isomer of A) led to very poor efficiency G-myristoylation (**Fig. 1b**, **Supplementary Table 1a**). Furthermore, the crystal structure of HsNMT1 in complex with meGN revealed that the methyl rotated the side-chain of N3 around Cα3, which was orientated towards the catalytic base, whereas the methylated N-terminus was positioned in pocket 3 (**Extended Data 1b,c**), explaining the low performance of this substrate for G-myristoylation. Therefore, any subtle modification around G2 protrudes aa2 into pocket 3, resulting in aa3 oriented in pocket 2 at the catalytic center^17^.

Together, these data indicate that GK-starting peptides lead to G-myristoylation, which relies on G2 and the S6K7 motif. In the less favorable K3 context, S_6_K_7_ becomes even more crucial for G-myristoylation, consistent with previous reports suggesting strong dependence of G-myristoylation on K3 and K7^18,26^. However, considering that the N-termini of T106A/B (GKSLSHLP, GKT<u>F</u>S<u>Q</u>LG) are unambiguously NMT substrates (**Fig. 1b** and **Supplementary Table 1a**) despite the absence of K7, the deduced motif for G-myristoylation with K3 is GKXXSX (**Fig. 1e**). We noticed that both T106A/B harbor a highly hydrophobic residue at position 5 (L and F, respectively), another feature favoring G-myristoylation^16,25^, which probably compensates for the absence of K7 and leads to efficient G-myristoylation. In addition, both ARF6 and GK possess a hydrophobic residue at position 5 (**Fig. 1b**) so, consistent with its positive role, permutation of positions 4 and 5 in the GK background (GKFS) significantly reduced myristoylation efficiency (**Fig. 1b**).

### K3 myristoylation of N-acetylated peptides with the S6K motif

In apparent contradiction to the above results, the GK-starting protein ARF6 was recently shown to accept K-myristoylation *in vivo*^18^, but only if the G2 α-amino group was unreactive due to previous N-acetylation (*ac*) or G-myristoylation. We therefore examined any modification around the reactive G2 amino group allowing K-myristoylation of either the HPCA, ARF6, and/or GK-starting peptides. As expected, *ac* of the three G-starting peptides led to side-chain myristoylation, provided that K occupied position 3 (**Fig. 2a**, **Supplementary Table 1c** and **Supplementary Dataset 1**). When the K side chain was shortened to Orn, in the context of N-acetylated peptides, side chain myristoylation was still observed at position 3 (**Fig. 2a**). The catalytic efficiencies of the *ac* series were one order of magnitude lower than those of G-myristoylation (**Supplementary Table 1a**, **c**), with the catalytic constant (*k_cat_*) accounting for most of the *k_cat_/K_m_* decrease in both cases. This suggested that the affinity provided by the peptide core was similar but that catalysis had a limiting step.

**Fig. 2.**
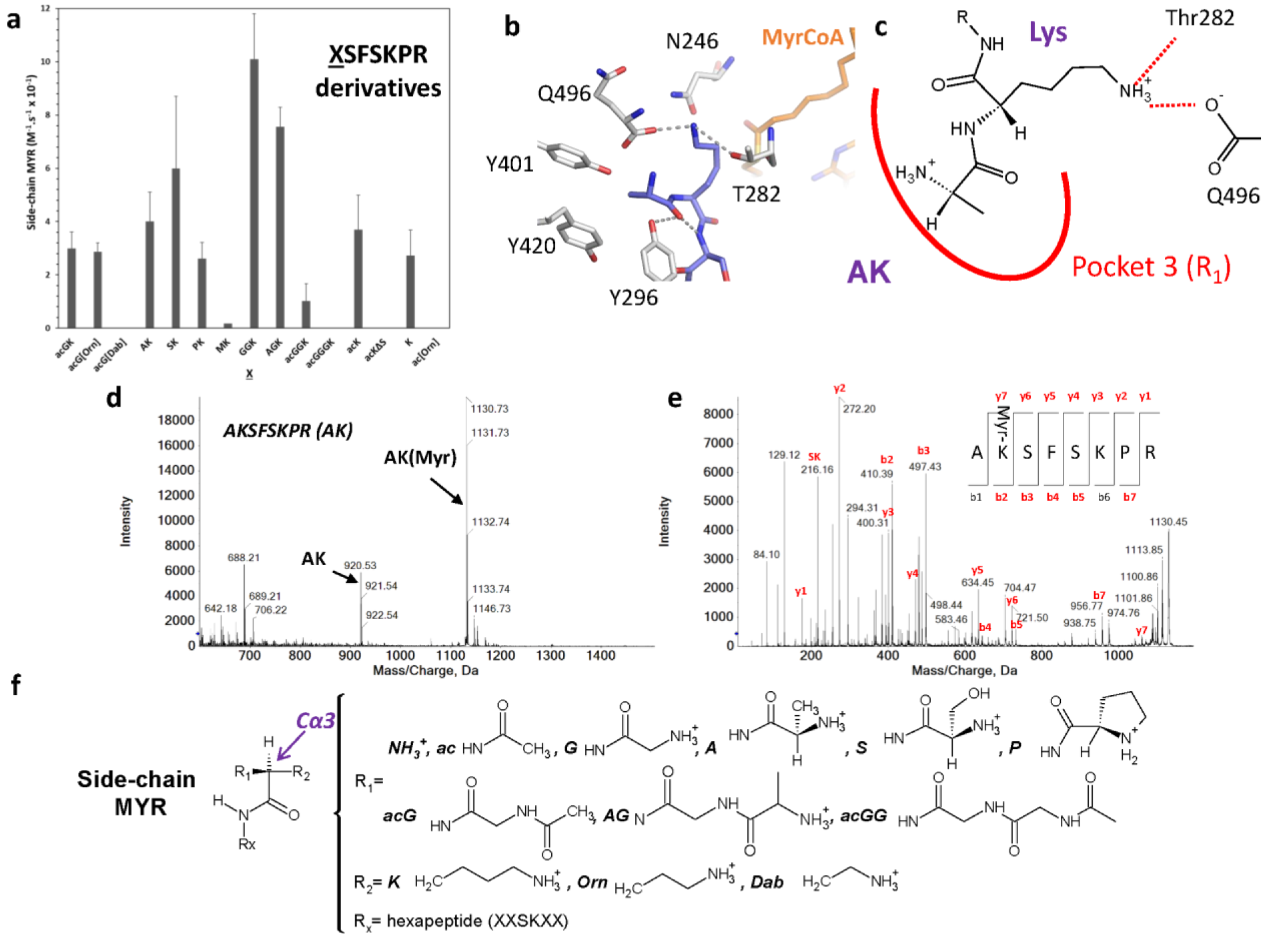
K-myristoylation and G-myristoylation rules. The peptides are derived from the GK peptide. (**a**) Main K-myristoylation catalytic efficiencies measured (see details in **Supplementary Table 1a**). (**b**) Crystal structures of HsNMT1 in complex with MyrCoA and the AK variant. The panel displays a close-up of the structural environment around the catalytic base, the C-terminus of the C-terminal residue Q496. (**c**) corresponds to plane representations of panel b. (**d**) MS1 analysis of the AK derivative after incubation with HsNMT1. (**e**) MS2 analysis of the 1,131 Da myristoylated peptide in panel d showing K-myristoylation. (**f**) New knowledge on the type of acceptor amino group leading to K-myristoylation. The panel shows that if a lysine side chain or a non-natural basic side-chain with reduced length occurs at R_2_, and depending on the substitutions made around the N-terminus, i.e., while blocking it (*ac* derivatives) or slightly modifying its plane orientation (e.g., if G is substituted for A, S, or P), then K-myristoylation (ε-myristoylation) is favored over N-terminal myristoylation (α-myristoylation)^17,18^.

The crystal structures of HsNMT1 in complex with acG[Orn] or an acGN non-reactive derivative were obtained and compared with the acGK and acGN variants. In both cases, there was rotation around the Cα of the aa3 side chain (Cα3), as expected from the *ac* addition. The acG group moved to pocket 3, and the aa3 side chain contacted the catalytic base (**Extended Data 2a-d**). The K side chain of acGK bonded directly with the C-terminal catalytic base of NMT, unlike the G-myristoylation mechanism, which involves a water-mediated bond (**Fig. 1c,d**). acG[Orn] led to a weaker salt bridge with the catalytic base than acGK, as the distance was increased from 3.1 to 3.6 Å (**Extended Data 2e**). Nevertheless, the associated K-myristoylation mechanism still involved direct interaction between the catalytic base and the ammonium group of Orn, like with acGK (**Extended Data 2c-f**). Further reduction to Dab dramatically decreased K-myristoylation (**Fig. 2a**, **Supplementary Dataset 1** and **Supplementary Table 1c**). Therefore, tight direct binding of the K side chain reactive amino group to the catalytic base most likely explains effective but slower K-myristoylation compared to G-myristoylation.

### Side chain myristoylation at K3 is preferred over N-terminal α-myristoylation when the N-terminal residue is not G

Therefore, Cα3 tends to rotate and embed the N-terminal group into pocket 3 provided any change in G2 (**Fig. 1e**) and, if aa3 displays an amino group (e.g., K3), it becomes reactive and promotes K-myristoylation. This suggests that the strong negative effect of the G2A substitution might favor K-myristoylation in the presence of K3. To test this hypothesis, we examined G2A HPCA (A-HPCA), ARF6 (A-ARF6), and GK variants (AK), all featuring a K3. In all cases, crystallographic and/or MS analysis unambiguously showed K-myristoylation (**Fig. 2b-e**, **Extended Data 3** and **Supplementary Table 1c**). The catalytic efficiencies with AK and A-ARF6 were (i) significantly greater than with A-HPCA and (ii) of the same order of magnitude as the acetylated versions of the corresponding variants. Therefore, the K side chain was preferred over the α-amino provided that a side chain (e.g., methyl of A) was grafted onto the reactive residue instead of G2. The crystal structure of AK complexed with HsNMT1 (**Fig. 2b,c**) showed that the K3 side chain bonded to the C-terminal catalytic base (Q496) and to T282 and the A2 side chain buried in pocket 3. Again, direct bonding to the catalytic base likely explains the reduced catalysis compared to the G-myristoylation water-mediated bonding of the GK or HPCA peptide. The lower *K_m_* of the acG derivatives may be related to additional interactions of the entire A2 chain in pocket 3 (**Fig. 2b**).

We next interrogated the human proteome to discover possible K-myristoylation targets featuring AKXXSK immediately following an M start residue (M1). Only tuberin (TCS2; P49815; **Supplementary Table 2a**) was identified, but no myristoylation was observed with the corresponding peptide derivative (AKPTSKDS; **Supplementary Table 1c**), perhaps due to the P4 inducing a local conformational kink disfavoring both G- and K-myristoylation by misaligning S6K7 with the reactive amino upstream group. The absence of a hydrophobic side chain at residue 5 (similar to A-HPCA *vs* A-ARF6 or AK) might also explain this phenomenon, as noted above for G-myristoylation.

We next examined whether non-G N-termini other than A might also lead to K-myristoylation by replacing the N-terminal G2 of the GK peptide with M, S, and P. K-myristoylation occurred in all cases with catalytic efficiencies inversely proportional to side chain length (**Fig. 2a**, **Supplementary Table 1c** and **Supplementary Dataset 1**). We concluded that the nature of the N-terminal residue was a major determinant guiding N-terminal K-myristoylation, provided that aa3 harbored a proximal acceptor group (**Fig. 2f**). Due to improper positioning of the N-terminal amino group, any non-G-starting peptides with the S6K7 motif may be modified by NMT but with extremely poor catalytic efficiencies several orders of magnitude lower than the corresponding G derivative. A K3 downstream of any amino acid at position 2 but G permits K-myristoylation with dramatically increases associated catalytic efficiencies (**Supplementary Table 1a**). Higher efficiencies are possible when the peptide context is favorable, as with AK and A-ARF6 peptides (this work and ^18^). Remarkably, both peptides feature a bulky, hydrophobic, favorable residue at position 5 (F or L, respectively), already shown to be important for G-myristoylation (see ^16^). Finally, interrogation of data libraries indicated that no protein with A2 (i.e., originating from M1 excision) undergoes K-myristoylation in the human proteome.

Taken together, our data reveal that K-myristoylation does not require a given side chain of the first amino acid but relies on optimal residues at position 5-7. The [^G]KX[FL]SK motif emerges as the determinant of K-myristoylation (where ^X means any residue but X). Any non-G2 side chains appear to privilege pocket 3 binding, preventing their reactivity and ensuring high specificity for G2 undergoing G-myristoylation.

In the human proteome, 7548 hits harbored a KXXSK motif (**Supplementary Table 2a**) corresponding to 2831 unique entries with 3615 sites in 6764 proteoforms (14-15% of the proteome). 572 sequences displayed F or L at position 3, like in ARF6 or the GK peptide. Among the 29 sequences with translation start sites at positions 1 or 2 (i.e. arising from cotranslational N-Met removal rules), 10 of the 15 known G-myristoylated N-termini starting with GK were found ^16^ plus a few additional sequences, most not comprising F or L3, like TSC2. The other 3595 sequences corresponded to internal sequences starting beyond codon 3, and 285 were preceded by a G suggesting large scope for post-translational G-myristoylation in this subset. Post-translational myristoylation, however, requires prior unmasking by proteolytic cleavage provided before the first XK. Indeed, a number of K-myristoylation sites among the 543 [^G]KX[FL]SK internal sequences were preceded by a classic caspase motif (**Supplementary Table 2b-e**). Therefore, unlike G-myristoylation, K-myristoylation should solely arise from post-translational myristoylation.

### Myristoylation may occur on Lys4

We next wondered whether extending the amino acid sequence upstream of the reactive K might lead to K-myristoylation, provided that a two-amino acid spacer between the accepting K and the S6K7 motif was retained. We therefore grafted an extra N-terminal A or G to the GK peptide (A- or G-GKSFSKP; AGK or GGK) and checked whether HsNMT1 could acylate the K at position 4. By MS and kinetic analysis, K-myristoylation was as efficient with these A/G-grafted peptides as with the acG peptides (**Fig. 2a**; **Supplementary Table 1c** and **Extended Data 3**). We concluded that the pocket usually hosting residue 3 could host a dipeptide moiety as large as AG provided that a close and available K-reactive side chain was properly spaced from the crucial S6K7 motif. We noted that, with five bonds and a positive charge, a K side chain was the most bulky moiety tolerated in NMT pocket 3 for G-myristoylation. With seven bonds, the AG backbone is much bulkier than K but still promoted K-myristoylation (**Fig. 2f**). To establish whether larger chains would make K-myristoylation possible, we examined large, poorly-branched N-terminal grafts larger than GG or AG (AcGGK, AcGGGK, AcGGGGK, and AcGGGGGK with 9, 12, 15, and 18 bonds, respectively) upstream of K3 in the GK series. With AcGGK and AcGGGK, there was unequivocal K-myristoylation by MALDI analysis (**Supplementary Dataset 1**). K-myristoylation catalytic efficiency progressively decreased with increasing chain length to very low values, and K-myristoylation was not detected for AcGGGGK or AcGGGGGK (**Supplementary Table 1c**). The crystal structure obtained with the GGK derivative showed that the GG chain is fully buried in pocket 3 and propels the main backbone peptide chain into a more remote location at Cα3 (**Supplementary Fig. 2a,b**); there is also a greater distance between the reactive K amino group and both the catalytic base and T282. The 7-bond GG or AG chain corresponds to the bulkiest chains accepted by pocket 3 for efficient K-myristoylation (**Fig. 2f**). When longer than 4 residues, as in the case of K, the backbone chain tends to not be perfectly aligned, but this is not responsible for the lower catalytic rates observed.

We concluded that to allow K-myristoylation, the [^G]KX[FL]SK motif may be associated with a one amino acid extension or *ac* at the N-terminus, provided that an *ad hoc* proteolytic cellular machinery can produce such extremities. In addition, about two dozen internal XXKXXSK K-myristoylation motifs were noted in the human proteome displaying an upstream classic caspase cleavage site (**Supplementary Table 2a,b,c**).

With the knowledge that K4 may be reactive in the context of a G_2_XK_4_ derivative, we next assessed whether G2- and K4-myristoylation could compete within the same protein. For this, we first altered the G-myristoylation S_6_K_7_ motif of the GK peptide by introducing an additional S between S6 and K7, pushing the K back to position 8 (G_2_G_3_K_4_F_5_S_6_S_7_K_8_PR). The resulting peptide therefore featured two overlapping myristoylation motifs separated by a two amino acid spacer, one for modification of the α-group of G2, relying on motif S_6_S_7_, and a second on the ε-amino of K4, relying on motif S_7_K_8_. This modification reduced the catalytic efficiency compared to the GK peptide (**Supplementary Table 1a**). Furthermore, the MS2 spectra of the 1,173 Da myristoylated product favored G-myristoylation over K-myristoylation (**Extended Data 4a**), with evidence of the major G-myristoylation product as unambiguous characteristic b_2_, y_7_, and y_8_ ions (**Extended Data 4b**). Nevertheless, we could not exclude the possibility of K-myristoylation, as the associated prototypic K-myristoylation ions could only be identified at low intensity. Therefore, other experimental approaches were required to assess for simultaneous K-myristoylation.

### The IpaJ protease as a tool to unravel ε-myristoylation

Whatever the context, G-myristoylation is favored over K-myristoylation when in competition. Since only an N-terminal G appears to promote myristoylation, K-myristoylation is unlikely to originate from a co-translational event. However, because K-myristoylation might be further added through post-translational insertion, identifying the conditions favoring this event would be useful. To date, two physiological proteolytic states have been shown to impact myristoylation: caspase and IpaJ proteolytic cleavage, and indeed several internal K-myristoylation sites may arise from caspase cleavage.

The virulence factor IpaJ from *Shigella flexneri* is a cysteine protease that specifically recognizes the myristoyl (Myr)-G moiety of several human proteins *in vivo*^24^. As a result of its activity on Myr-G-peptides (**Supplementary Fig. 3a**), IpaJ is predicted to induce a 267 Da shift (Myr-Gly product) in MALDI MS spectra. Since it cleaves off G-myristoylation sites, we anticipated that IpaJ - provided it specifically processes any G-myristoylated peptide – could be used to unravel unambiguous and undiscovered K-myristoylation modification sites.

The *S. flexneri IpaJ* gene was cloned and the protein overexpressed and purified to homogeneity (**Supplementary Fig. 3b,c**). An inactive variant (IpaJ-C64S) was also produced as a control. Under specific conditions (**Supplementary Fig. 3d**), IpaJ was sufficient to cleave off the G-myristoylated moiety from any tested peptide *in vitro*, unlike IpaJ-C64S (**Supplementary Fig. 4a,b,c**). Additionally, IpaJ was inactive on an N-acetylated G-peptide, an A2-myristoylationed peptide, and a K3-myristoylationed peptide derived from ARF6 (**Supplementary Fig. 4d** and **Supplementary Table 1d,e**). Due to its unique and high specificity for G-myristoylation and the observed complete cleavage of any G-myristoylated peptide, we concluded that IpaJ could be used as powerful tool to distinguish G-myristoylation from K-myristoylation and, additionally, investigate IpaJ-dependent post-translational K-myristoylation.

We therefore established an IpaJ pipeline that determined the myristoylation type of any peptide subjected to NMT (**Fig. 3a**). The workflow faithfully reproduced the sequence of events occurring in the context of cellular IpaJ activity. As proof-of-concept, a dozen peptides were carefully chosen for their sequence diversity and ability to undergo G-myristoylation (**Supplementary Fig. 5a,b**; see ARF6 in **Supplementary Fig. 6**). Combined with previous data^23^, IpaJ efficiently cleaved any G-myristoylated peptide regardless of the sequence downstream of the G residue (**Supplementary Fig. 5c**).

**Fig. 3.**
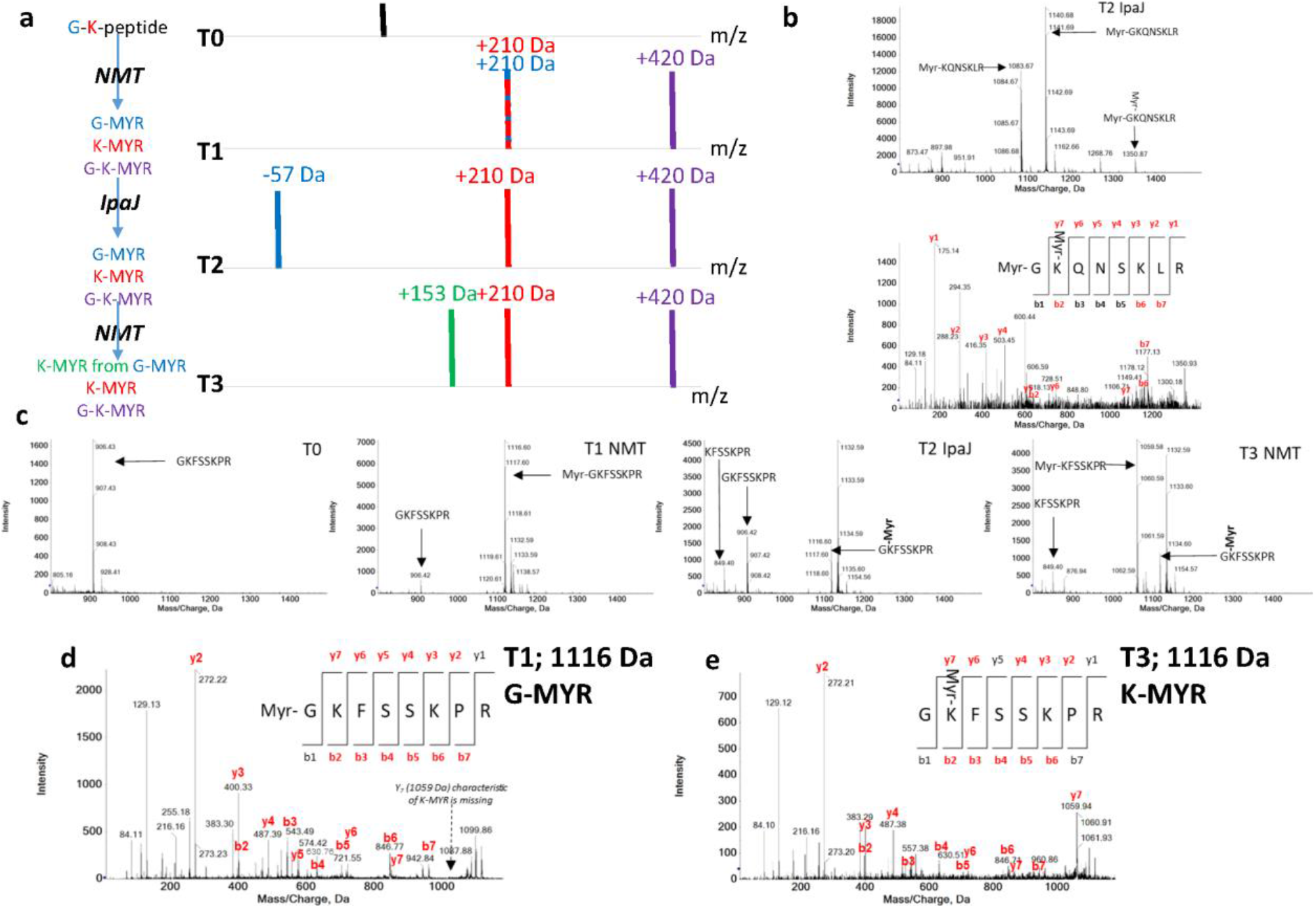
A pipeline to investigate NMT/IpaJ-coupled impact on myristoylation. (**a**) The schematized NMT/IpaJ pipeline, expected m/z at each step, and corresponding isoforms. In brief, peptides are first incubated with NMT to allow myristoylation and visualization of the +210 Da addition (T1). Next, the reactor is mixed with IpaJ to allow complete removal at low temperature of the G-myristoylated moiety, which results in a −57 Da shortening compared to the starting peptide and −267 Da signature of the product compared to the myristoylated peptide (T2). Finally, the reactor is mixed with high NMT concentrations to check whether substrate myristoylation results from the action of the first two steps (T3). (**b**) HPCA peptide at T2 (top) and MS2 spectrum of the double myristoylation product (bottom). (**c**) GKFS peptide (GKFSSKPR) in the course of the different steps of the pipeline. (**d**) MS2 spectrum of the 1,116 Da peptide at T1; the missing y_7_ (1,059 Da) characteristic ion of K-myristoylation is indicated with an arrow. (**e**) MS2 spectrum of the 1,116 Da peptide at T3 with the 1,059 Da ion now visible.

Taking the double myristoylation of ARF6 at both the N-terminus of G2 and the side chain of K3 into account^18^, we used the IpaJ pipeline to identify double myristoylation of other GK-starting NMT substrates. We identified dual myristoylation of a peptide derived from HPCA/HPCL1/NCALD in the presence of IpaJ (**Fig. 3b; Supplementary Fig. 7**). Finally, double myristoylation was not observed with other closely related substrates of HPCA in the same NCS1 family of calcium sensors sharing the GK motif (**Supplementary Dataset 1**). We cannot exclude that the extremely high hydrophobicity of such acylated products prevented their crystallization in the matrix and consequently their identification.

### Competition between G- and K-myristoylation

With the IpaJ/NMT pipeline in hand, we next examined the ambiguous myristoylation state of the GGKFSSKPR peptide. At T1, MS2 analysis unambiguously revealed G-myristoylation as a product (**Extended Data 4b**). At T2 (i.e., after incubation with NMT and IpaJ), both an uncleaved myristoylated peptide and a shorter, 267 Da peptide resulting from G-myristoylation cleavage were identified (T2, **Extended Data 4c**). There was also unambiguous and unique K-myristoylation at K4 of the GGKFSSKPR peptide at T3 (**Extended Data 4c,d**).

Interestingly, the IpaJ product of G-myristoylation of this peptide started with GK to form GKFSSKPR (GKFS), which displayed reduced myristoylation efficiency (**Fig. 1b**). This peptide could be further myristoylated at T3 to produce a 1,116 Da form. MS2 analysis revealed K-myristoylation at the N-terminal K3 of this peptide (**Extended Data 4d**). To obtain a simpler set of products for the pipeline, we further characterized the myristoylation profile of GKFS in the IpaJ workflow (**Fig. 3c,d,e**). This peptide underwent both G- and K-myristoylation. GKFS features a bulky residue at position 4 and a small residue at position 5. We hypothesized that while a spacer is mandatory for both myristoylation types, either myristoylation might be favored depending on as yet unknown amino acid requirements occurring in that spacer (i.e., aa4-5, the two residues switched in the GK series). This is likely, as both pockets 4 and 5 are large enough to accommodate all side chains^16^, so some combinations could favor K-myristoylation in a GK-starting context.

We found no straightforward natural chassis in human sequences starting with a G2 that leads to competition between K- and G-myristoylation and a systematic preference for G-myristoylation in sequences originating from co-translational myristoylation. When we applied the IpaJ pipeline to GK-starting peptides derived from the human myristoylome and exploited its enhanced sensitivity to detect K-myristoylation, we observed that most displayed the myristoylated peptide at T2 after IpaJ cleavage (**Supplementary Table 1e**). Therefore, competition exists between G- and K-myristoylation on GK-starting peptides. Furthermore, none of these peptides displayed a hydrophobic residue at position 4, explaining why the K-myristoylation/G-myristoylation ratio was smaller than with the GKFS peptide and why prototypic K-myristoylation ions were not detected in MS2 spectra. Nevertheless, as myristoylation can also arise from post-translational addition provided that a cleavage generates a new K–accepting site, competition favoring K-myristoylation is likely. Indeed, as noted above, hundreds of internal proteins display an F/L4 profile (**Supplementary Table 2a**; see columns aa3, F3, and L3 corresponding to this position, as all sequences start at the cleavage site and not the unprocessed M1).

In conclusion, a G at the N-terminus of peptides with GKFXSK motifs sustains competition between K- and G-myristoylation. These data also indicate that the KXXSK motif – not only [^G]KX[FL]SK – is indicative of K-myristoylation independent of the residue preceding the K.

### IpaJ reveals that K-myristoylation may also occur on free or acetylated K1

When the IpaJ/NMT pipeline was applied to either the myristoylated GGKFSSKPR or the GKFSSKPR peptides (see above and **Fig. 4a**), IpaJ-induced cleavage of this reaction product also led to myristoylation of an unexpected K-starting peptide resulting from reiterated G-myristoylation IpaJ cleavage of each new G-myristoylation product. Although we could not distinguish alpha and epsilon myristoylation, side-chain modification was expected (K-myristoylation). Taking into account the S_6_K_7_ clamping of the peptide at the peptide-binding site, 3D molecular modeling indicated that the N-terminal free amino group would be too distant from the catalytic base for myristoylation. To test the hypothesis of K-myristoylation, we first prepared an acetylated (i.e., N-α-blocked derivative) of the GK peptide starting with K3 (acK). In the presence of MyrCoA and NMT, this peptide was myristoylated with catalytic efficiencies similar to the previously tested peptides (**Fig. 2a**, **Supplementary Table 1c**). When the N-terminal acK was installed at the N-terminus of a peptide with the aa4-5 spacer shortened to only one residue, myristoylation was no longer observed (acKΔS, **Supplementary Table 1c**). Finally, we checked that such K-starting peptides derived from the GK (K) and HPCA chassis (K-HPCA) underwent K-myristoylation (**Fig. 2a**, **Supplementary Table 1c**). The crystal structure of the complex between NMT and an acK derivative revealed two alternative structures, both of which supported a K-myristoylation mechanism involving direct interaction between the K side chain and the catalytic base (**Fig. 4b**). The group around Cα3 could be retrieved in two opposite directions: the first showed the *ac* group entering either pocket 3 mimicking an N side chain (**Fig. 4c**) while the overall active site positioning mimicked that observed with GK; and the second revealed the *ac* moiety occupying part of pocket 4, which is large enough to host two moieties (i.e., *ac* and S4 in acK) from the same peptide (**Fig. 4b,c**, right). We concluded that side chain myristoylation can also occur on an N-terminal K (K1) regardless of acetylation status and that might result from post-translational proteolytic cleavages such as those induced by IpaJ during the course of *Shigella* infection.

**Fig. 4.**
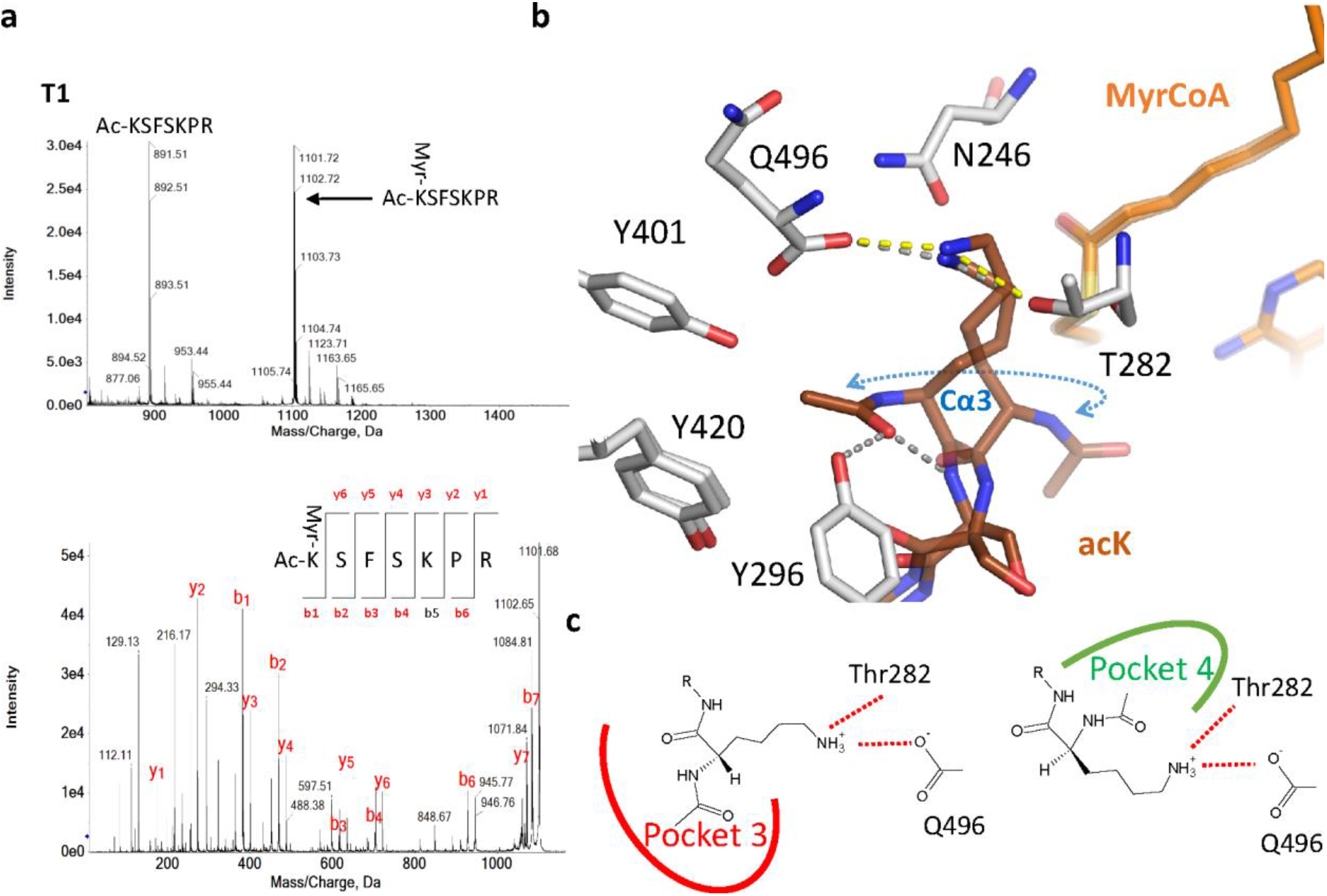
K-myristoylation mechanism on an N-terminal side chain. The crystal structure of the complex made between acK and HsNMT1 is displayed. (**a**) MS analysis showing myristoylation of the acK; top MS1, bottom MS2. (**b**) The two conformations are displayed, and the rotation point around Cα3 is indicated. (**c**) Planar representation showing how each conformation may position the *ac* chain within pocket 3 (conformation A, left hand side) or pocket 4 (conformation B, right hand side).

We next interrogated the 15 GK starting peptides derived from different human proteins using the IpaJ pipeline (**Supplementary Table 1e**). As noted above, because they mostly resulted in G-myristoylation, they are likely to generate K1 N-termini after IpaJ cleavage. **Supplementary Fig. 7** summarizes such behavior with HPCA. As expected, there was evidence of K1-myristoylation at T3, further indicating that all such peptides can undergo K-myristoylation after IpaJ cleavage (**Supplementary Dataset 1**). We conclude that all GK-starting proteins likely undergo post-translational K-myristoylation after IpaJ cleavage.

### K-myristoylation-inspired design of potent NMT inhibitors

The data above use synthetic K mimics to assess the effects of the side-chain length, and in doing so we noticed that ac[Orn], ac[Dap], A[Orn], and acG[Dab] do not undergo myristoylation even at the highest NMT concentrations (**Fig. 2a** and **Supplementary Dataset 1**), unlike their close homologs, acK, AK, and acGK. We hypothesized that such compounds might act as NMT inhibitors. While the ac[Dap] derivative was a poor inhibitor with an inhibition constant of 11 ± 1 μM, the ac[Orn] derivative showed an order of magnitude higher affinity (**Fig. 5a**), suggesting that the N-terminal reactive amino group – if properly aligned with the α-carbon through 2-3 carbon bonds like in Orn or Dab – might contribute to binding while not promoting catalysis. This encouraged us to further design and study compounds with variations around N-terminal Orn and Dab derivatives based on the GK sequence.

**Fig. 5.**
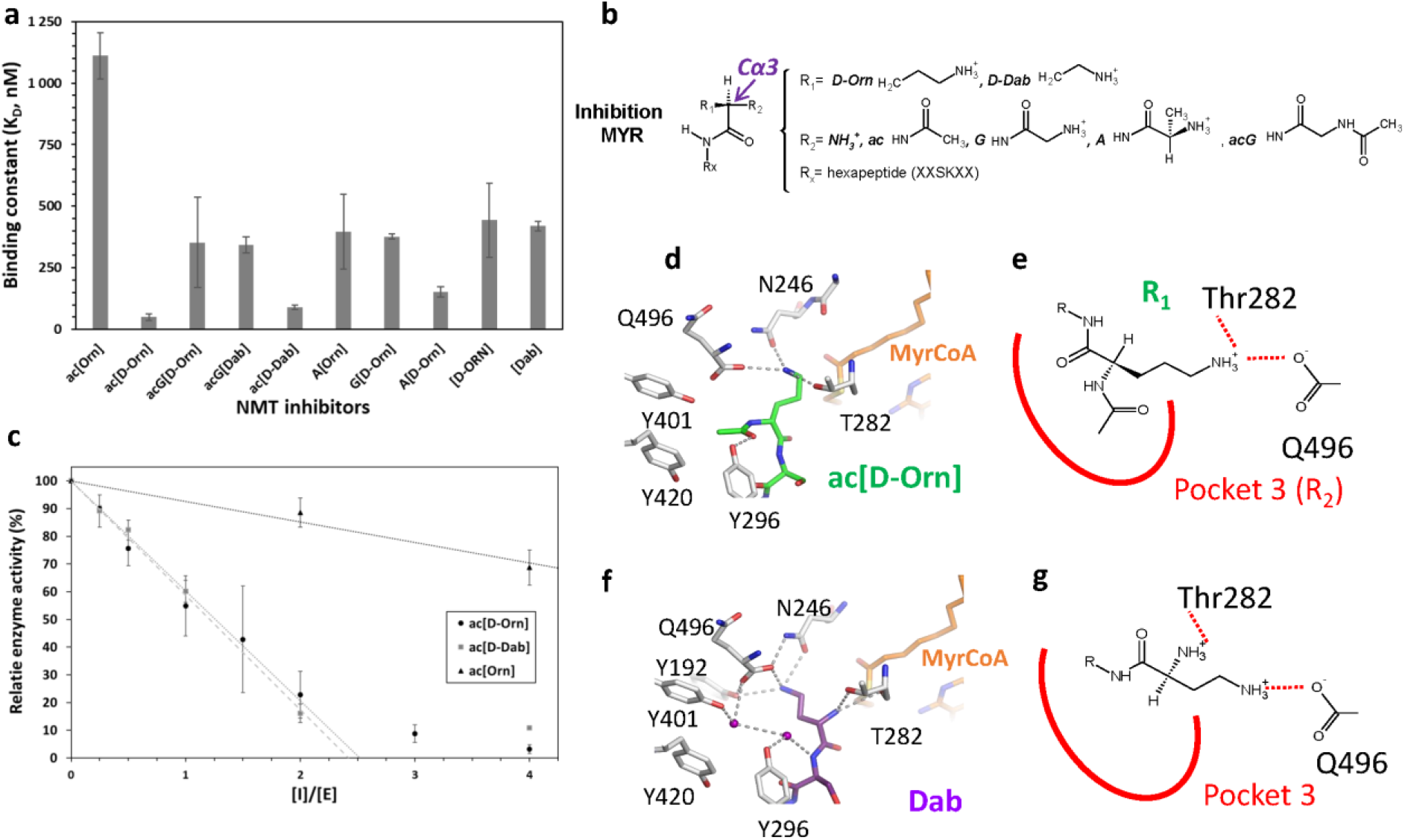
Characterization of K-myristoylation-inspired suicide inhibitors of NMT. The peptides are derived from the GK peptide. (**a**) Apparent binding constants were derived from IC_50_ value measurements according to ^46^. All reactants but the reactive peptide SOS3 were pre-incubated in the presence of the two peptides for 30 minutes. The reaction was triggered by the addition of SOS3 and followed over time for 60 minutes. Because of the bi:bi mechanism, MyrCoA is required to form the complex^5,50^. (**b**) 3D and (**c**) plane representations of the interaction networks around the ac[D-Orn] derivative bound to HsNMT1. (**d**) Plot showing that the ac[D-Orn] (black circles) and ac[D-Dab] (grey squares) derivatives display low partition ratios. The ac[L-Orn] inhibition pattern is indicated (black triangles). The reaction was triggered by the addition of the productive substrate after pre-incubation of the inhibitor for 60 minutes in the presence of MyrCoA and HsNMT1. Time-course analysis was next achieved to determine the remaining enzyme activity. A value of 100 was given to the value obtained in the absence of inhibitor. 3D (**e**) and plane (**f**) representations of the interaction network around Dab bound to HsNMT1. (**e**) Representation in a model similar to that of **Fig. 1b and 2a** for G- or K-myristoylation, respectively substrates of the chemical requirement leading to NMT inhibition.

We first assessed acG[Dab], [Dab], and A[Orn], all of which acted as potent competitive inhibitors with IC_50_ values of the order of the NMT concentration used in the assay, indicating a tight binding constant in the low micromolar range (i.e., <0.4 μM; **Fig. 5a**). Dab and Orn but not Lys clearly led to strong inhibition.

To modify the amino group orientation towards the catalytic base and to mimic the G-myristoylation productive mechanism, we built two derivatives with D-Orn - the enantiomeric stereoisomer of L-Orn (**Fig. 5b**; symmetric branching of R_1_ and R_2_ with respect to Cα3) - either with *ac* or acG to optimize the filling of pocket 3. Interestingly, the ac[D-Orn] derivative was a remarkably close mimic of a GN3 variant (**Extended Data 5a**), while our previous work revealed that N is favored in pocket 3^1,16^. Both D-Orn derivatives were potent inhibitors, with the *ac* version the most potent (*K_D_*=50 nM). A similar result was observed with ac[D-Dab] or A[D-Orn] derivatives (**Fig. 5a**), and acG[D-Orn], G[D-Orn], or [D-Orn] derivatives were also strong inhibitors but to a lesser extent than ac[D-Orn]. When ac[D-Orn] was installed on the NCS1 chassis – i.e., without a hydrophobic position at position 5 – significant, though weak, myristoylation was observed (**Supplementary Table 1c**). This indicated that potent inhibition was only possible with an optimal fit for K-myristoylation including a hydrophobic residue filling pocket 5.

Prolonged incubation with NMT followed by MS analysis of the reaction products of the above inhibitors revealed product formation, though to a much lower level than the substrate. Furthermore, these peptides acted as suicide inhibitors (i.e., the unreleased product was the effective inhibitor). Indeed, pre-incubation of the inhibitors with the enzyme and MyrCoA rapidly inactivated the enzyme, indicating conversion of the inhibitors bound to the enzyme into a one- to two-order of magnitude more potent inhibitor. We concluded that these inhibitors behave as single-turnover substrates^27^. There was also a partition ratio of 1.5 of either D-Dab or D-Orn derivatives of the *ac* series (**Fig. 5c**), indicating very high inactivation efficiency with an inhibition on-rate similar to that of the off-rate.

Finally, we obtained the crystal structure of ac[D-Orn] and [Dab] in complex with NMT and retrieved the complex before conversion of the substrate into a product (**Fig. 5d-g**). The complex featured direct binding of the D-Orn and Dab amino groups to the carboxy terminal base. The reactive amino group of the Dab side chain bonded directly with the carboxy-terminal base of NMT through two alternative orientations flipped around Cα3, one of which revealed a bond between the Nα ammonium and Thr282 (**Fig. 5f-g**, **Extended Data 5b,c**). Such direct bonding of the reactive amino group of both inhibitors is unlike the efficient GK peptide, which requires water-mediated binding, instead being more similar to the AK substrate derivative binding (**Fig. 3**). According to our data, this direct salt bridge created in the context of the Orn or Dab side chains, which leads to optimal sizing, is expected to be stronger than the one made with K and at least 100-fold less prone to reaction as a result. This likely explains the lesser reactivity and the inhibitory behavior. Both compounds bound to Thr282 (**Fig. 5d-g**). The non-acetylated Dab compound featured a water network within pocket 3 indicative of less bonding compared to both acetyl derivatives, perhaps explaining its lower binding constant.

In conclusion, (i) the chain length bringing the reactive amino group, (ii) the chiral conformation of the substituents R_1_ and R_2_ of the Cα group facing pocket 3 (Cα3; compare **Figs. 1e**, **2f** and **5b**), and (iii) optimal occupation of pocket 3 by a moderately bulky N-terminal moiety mimicking N were mandatory for effective K-myristoylation mechanism-based design of NMT inhibitors.

## DISCUSSION

N-myristoyltransferases (NMT), also known as glycylpeptide N-tetradecanoyltransferases (EC 2.3.1.97; see BRENDA resource ^28^), are the only enzymes known to catalyze N-myristoylation in eukaryotes. It was long believed that NMT only co-translationally transferred a lipid (myristate) to the α-amino group of the N-terminal amino group of G-starting proteins mostly exposed by the prior action of methionine aminopeptidases (MetAP). Recent crystal structures of NMTs co-crystallized with G-starting substrates have revealed how the side chains of these substrates are recognized and lie within dedicated cavities (pockets 3-8; ^1,16,17^), characteristics that contribute to the G-starting recognition pattern. Although this G-starting recognition pattern appeared to clearly define a global specificity, it did not explain the entire NMT-dependent myristoylation landscape. This was even more evident when we and others reported the possibility that NMTs catalyze K-myristoylation^17,18^. Although this K-myristoylation function should expand the range of NMT substrates, the extent and clues ruling NMT-dependent K-myristoylation are unknown.

Here we disclose the unique and common specificities underlying NMT-dependent G- and K-myristoylation. We reveal that the main specificity rules for NMT-dependent G- and K-myristoylation partially overlap, leading to competition between the two types of myristoylation in GK-starting contexts. The myristoylation rules are summarized in **Figs. 1e and 2f**. The very narrow shape of pocket 2 ensures reactivity of a primary amino borne only on unbranched moieties such as N-terminal glycine or side chains like lysine, which facilitates the 70° rotation of the amino-reactive group from the catalytic base towards the thio-reacting group of MyrCoA. This motion allows Wat3 to enter from the water channel and play a crucial role at the transition state^17^. The addition of a methyl group such as in A2 or meG variants makes this motion unlikely and dramatically decreases catalysis. The rules directing G- or side chain-myristoylation are predominantly determined by the R_1_ and R_2_ chemical groups branched at Cα3. NMT usually favors G- over K-myristoylation on proteins starting with G, while K-myristoylation dominates on any other N-terminal chain.

Overall, NMT-dependent K-myristoylation is less efficient than G-myristoylation whatever the context. Although the two myristoylation types may directly act on an N-terminal residue, K-myristoylation appears less strict than G-myristoylation in this regard; indeed, K-myristoylation does not require a given side chain of the first amino acid but relies on optimized residues at positions 5-7. A major difference between the two myristoylation types is that K-myristoylation needs the KXXSK-restricted pattern. This is unlike G-myristoylation, which allows the modification of many proteins not exhibiting the S_6_K_7_ motif by NMT on their N-terminal G (see ^16^ and **Supplementary Table 1a**). The KXXSK motif of K-myristoylation can be associated with a one amino acid extension or acetylation at the N-terminus provided that an *ad hoc* proteolytic machinery can produce these extremities. The specificity of MetAPs - which remove the first M from proteins - favors new N-terminal residues with short side chains like A, G, P, or S. A survey of the human proteome reveals, however, no K-myristoylation on either K2, K3, or K4 and 15 G-myristoylation substrates in the subset of proteins arising from MetAP-guided co-translational cleavage rules. Among MetAP unprocessed substrates, five MK-starting proteins were identified, but we ruled out that they might undergo K-myristoylation. A number of internal KXXSK K-myristoylation motifs are preceded by caspase cleavage sites in the human proteome suggesting that K-myristoylation - in contrast to mostly co-translational G-myristoylation - results from post-translational addition, for instance initiated by cleavage by specific caspases during apoptosis^15,29^.

We show that post-translational myristoylation may occur either after initial G-myristoylation, causing double myristoylation, or upon protease cleavage. This includes the action of bacterial protease IpaJ, which we used as a case study and as a unique tool to discriminate between G- and K-myristoylation. Our data confirm that ARF6 is myristoylated in both states^17,18^, in contrast to the other ARFs (ARF1/3/4/5) or ARL1, which do not feature a K3 like ARF6. This insensitivity of ARF6 to IpaJ was initially proposed to be due to ARF6 not being located at the ER/Golgi, unlike the other members of the ARF/ARL family, but rather at the plasma membrane. ARF6 translocates to endosomes, where it acts as a major regulator of endocytosis^30,31^. Our data indicate that ARF6 resistance to IpaJ might also result from the unmasking of a myristoylatable K3 upon G-myristoylation cleavage. As a result, ARF6 would still be myristoylated and retained on myristoylation affinity azido-biotin prior to and after IpaJ cleavage.

We noticed that K-myristoylation or any other side chain myristoylation systematically reduced *k_cat_* values, making the reaction poorly efficient. The associated *k_cat_* values were significantly lower than G-myristoylation and in the low range of reported values^32^. These data, together with the observation of direct, non-water mediated bonding of the reactive amino group of the side chain to the catalytic base, suggested the occurrence of long-lasting enzyme-ligand or enzyme–product intermediary complexes prior to product release. Taking advantage of this observation, we designed a series of NMT inhibitors based on our new knowledge of the determinants of side chain myristoylation. Specifically, a K-myristoylation substrate could be turned into a very potent suicide inhibitor through simple manipulation around the acceptor side chain, i.e., the stereo-isomeric D version of a K-mimicking, shorter side chain, including Orn or Dab. This reflects the tight fit of the active site and mechanism of both K- and G-myristoylation and highlights that even very small modifications (such as methylene variations, e.g., G2 to A or me-Gly for G-myristoylation, acK to ac[Orn] for K-myristoylation) produce distinct and significant effects on the reaction outcome.

To conclude, this study extents our knowledge of the subproteome, which is sensitive to NMT-guided myristoylation, and provides new understanding about the subtle requirements for specific NMT inhibition.

## ACKNOWLEDGMENTS

This work was supported by French National Research Agency (ANR) DynaMYT (ANR-20-CE44-0013) and Fondation ARC (ARCPJA32020060002137) grants to TM. This work has benefited from the support of a French State grant (ANR-17-EUR-0007, EUR SPS-GSR) managed by the ANR under an Investments for the Future program (ANR-11-IDEX-0003-02), from the I2BC crystallization platform supported by FRISBI (ANR-10-INSB-05-01), and from the facilities and expertise of the I2BC proteomic platform SICaPS, supported by IBiSA, Ile de France Region, Plan Cancer, CNRS and Paris-Sud University. FR is supported by grants from Région Ile-de-France (17012695) and Fondation pour la Recherche Médicale (FDT202001010779). We warmly acknowledge Odile Schiltz (IPBS, Toulouse), Virginie Redeker, Jean-Pierre Le Caer, Laila Sago and David Cornu (all at SICaPS, Gif/Yvette) for their extensive support with mass spectrometry analyses. We thank the French National Synchrotron Facility (SOLEIL) for providing synchrotron radiation facilities and the staff of the Proxima 1 & 2 beamlines.

## AUTHOR CONTRIBUTIONS

FR conducted all MS experiments and kinetic analyses, characterized IpaJ variants, and set up the IpaJ/NMT pipeline. CD undertook cloning, mutagenesis, and purification of IpaJ and performed all NMT structural analyses. RFD completed and consolidated the kinetic analysis. TM and CG conceived the project, supervised the experiments and analyzed the data. FR, CD, CG and TM, wrote the manuscript.

## DATA AVAILABILITY

The nine crystal structures of NMT in complex with MyrCoA and the peptides reported here have been deposited at the PDB under codes 7OWM, 7OWN, 7OWO, 7OWP, 7OWQ, 7OWR, 7OWS, 7OWT and 7OWU.

## ADDITIONAL INFORMATION

Supplementary information is available for this paper.

## METHODS

### Chemicals

All peptides (see all sequences in **Supplementary Tables 1**) were purchased at 95% purity (Genscript, Leiden, Netherlands). NAD was purchased from Roche (Basel, Switzerland). All other chemicals were purchased from Sigma-Aldrich (St. Quentin, France). Stock solutions of myristoyl-CoA (0.2 mM) were prepared in sodium acetate, pH 5.6, and 1% Triton X-100, except for MALDI analysis, where cholate was used to reduce background.

Because the N-terminal myristoylated G residue is usually the second in the original open reading frame (ORF) after the initial M, we call it aa2 as the extreme N-terminal residue throughout this work for consistency with previous work from ourselves and others.

### *IpaJ* cloning

Wild-type full-length IpaJ numbering is used throughout the text. The nucleotide sequence encoding residues including the final TGA stop codon of *S. flexneri IpaJ* ORF (UniProt code Q54150) was optimized for *E. coli* expression (**Supplementary Table 3**) and synthesized *in vitro* (Genscript). For protein expression, the 30-259 fragment was cloned between the *Nco*I and *Hind*III restriction sites of expression vector pETM30 as a C-terminal fusion with glutathione S-transferase (GST, **Extended Data 2a**). A His-tag was placed at the N-terminus of the fusion to facilitate protein purification (see below). A TEV cleavage sequence inserted between the two ORFs allowed release of IpaJ from GST with only an additional N-terminal G-Ala-Met-Ala tetrapeptide sequence upstream of R30, the first residue of IpaJ. Two variants were created by site-directed mutagenesis using a QuikChange site-directed mutagenesis kit (Stratagene, San Diego, CA) with the primer pairs displayed in **Supplementary Table 3**. The first variant was inactive (C64S), while the other (3M) was fully active with improved solubility and stability over time (L97N/L99N/C103S).

### Enzyme production and purification

#### HsNMT1

The HsNMT1*l* isoform containing residues 81-496 corresponds to the major isoform^3^. HsNMT1*l* was cloned into pET16b as an N-terminal His-tag fusion^3^, and the recombinant protein was expressed and purified as described^16^. Isoform HsNMT1*s* containing residues 99-496 was cloned into pET28 and purified as described^17^.

The three soluble 30-259 variants of the protease IpaJ were produced as follows. Rosetta2pLysS cells expressing the pETM30 derivative were grown in 1 L of 2xTY medium supplemented with kanamycine (50 μg/mL) and chloramphenicol (34 μg/mL) at 37°C under vigorous shaking. Protein overexpression was induced with 0.5 mM IPTG at OD_600_=0.4. Cells were transferred at 20°C and grown for 20 hours. Cells were centrifuged at 10,000 × g, and the pellet was resuspended (10 mL/g) in lysis buffer (20 mM Tris pH=8.0, 0.2 M NaCl, 5 mM 2-mercaptoethanol, 5 mM imidazole, 5% glycerol). Cells were lysed with a sonicator Q700 (amplitude 50%, 10 s on and 20 s off) for 3 min at 4°C. The sample was then centrifuged at 40,000 × g (Rotor JA20) for 30 min and the pellet discarded. The supernatant was loaded on a Ni-IMAC-HisTrap™ FF 5 mL column (GE Healthcare, Chicago, IL) at 3.5 mL/min; elution was achieved with a liner 0-0.5M imidazole gradient run over 100 ml. Purified proteins were dialyzed in Spectra-Por7 semi-permeable dialysis membranes (8 kDa cut-off; Thermo Fisher Scientific, Waltham, MA) for 48 hours against conservation buffer (20 mM Tris pH=8.0, 0.2 M NaCl, 5 mM dithiothreitol, 55% glycerol) at 4°C. The sample was stored at −20°C.

Protein concentrations were measured with the Bio-Rad Protein Assay Kit using bovine serum albumin (Sigma-Aldrich) as reference.

### Protein crystallization and structure determination

#### HsNMT1:MyrCoA:X structure

HsNMT1*s* was used to solve the structures of the complexes with nine different peptides in the course of this study (HPCA, AK, AcK, AcG[Orn], AN, meGN, GGK, Dab and ac[D-Orn] with PDB entries 7OWM, 7OWN, 7OWO, 7OWP, 7OWU, 7OWQ, 7OWR, 7OWS and 7OWT, respectively. Suitable crystals of HsNMT1:MyrCoA:peptide substrates or inhibitors were obtained by co-crystallization using the hanging-drop vapor diffusion method at 20°C in the crystallization conditions previously described^16,17^. Briefly, crystallization droplets were formed by mixing 2 μL of the of HsNMT1:MyrCoA:peptide complex (ratio 1:1.5:1.5) at 6-9 mg/mL with 2 μL of the precipitant solution containing either 0.1 M MgCl_2_, 0.2 M NaCl, 0.1 M sodium citrate pH 5.6, and 18-24% (w/v) PEG 6K or 0.1 M sodium acetate pH 4.6, and 18-24% (w/v) PEG 6K. Crystals were cryoprotected in the reservoir solution supplemented with 15% (v/v) glycerol and flash cooled in liquid nitrogen. Complete X-ray datasets of complex were collected at 100*K* a single wavelength from a single crystal at the French National Synchrotron Facility (SOLEIL) PROXIMA1 (for HPCA and AcG[Orn] at λ=0.98400 Å and AcK at λ=0.97856 Å) or PROXIMA2 (for AK and GGK at λ= 0.98012 Å and Dab and Ac[D-ORN] at λ = 0.97563 Å) beamlines and at European Radiation Synchrotron Facility (ESRF) ID30A1 (for AN at λ=0.96600 Å) and ID30A3 (for me-GN λ=0.96775 Å) beamlines. Datasets were integrated with XDS^33^ and scaled and reduced using AIMLESS from the CCP4 package^34^. For crystals that suffered from anisotropic diffraction (HPCA, AcK and GGK), data were processed with STARANISO on unmerged data^35^ before AIMLESS data reduction. Crystals belonged essentially to the space group P2_1_2_1_2 with similar unit cell parameters with the exception of GGK and meGN belonging both to C2 with similar unit cell parameters (summarized in **Supplementary Table 4**). In both space groups identified, unit cells contained two NMT molecules per asymmetric unit. Structure resolution was accomplished in all cases using the molecular replacement method and solved using PHASER^36^ and the HsNMT1:MyrCoA:peptide ternary complex (PDB entry 5O9T or 6SK2) as a search model. The structure of AN was solved using MOLREP^37^ and protein coordinates of HsNMT1:MyrCoA (PDB entry 5O9T) as a search model. Structures were subjected to alternating refinement cycles using PHENIX and manual model building using COOT^38–40^. The good quality of the electron density maps also enabled the refinement of substrate peptide, reaction intermediate, and reaction product molecules bound to HsNMT1 in each complex. Chemical compound libraries were generated using PRODRG server^41^ in combination with eLBOW from the PHENIX suite. The geometry of the final models was validated using MOLPROBITY^42^. Figures were generated using PYMOL (DeLano Scientific LLC, http://pymol.sourceforge.net/). X-ray data collection and refinement statistics are summarized in **Supplementary Table 4**.

### Measurements of activity and inhibition parameters

HsNMT1 activity was assayed at 30°C in a coupled assay using an updated version of the previously described protocol^43^. The reaction mixture contained 50 mM Tris-HCl (pH 8.0), 1 mM MgCl_2_, 0.193 mM EGTA, 0.32 mM DTT, 0.2 mM thiamine pyrophosphate, 2 mM pyruvate, 0.1 mg/mL of BSA, 0.1% Triton X-100, 2.5 mM NAD^+^, 0.125 units/mL of porcine heart PDH (0.33 units/mg), 40 μM myristoyl-CoA, and 1-2000 μM peptides. Unless otherwise stated for tight binding studies, the reaction mixture was pre-incubated for 3 min at 30°C before starting the reaction by adding MyrCoA. A final volume of 200 μL was used in 96-well black plates (Grenier Bio One and Dominique Dutscher, Brumath, France; the optical path for 0.2 mL is 0.55 cm). A value of 6300 M^−1^.s^−1^ was used as the molecular extinction coefficient of NADH at 340 nm. An Infinite M Nano^+^ plate reader equipped with micro-injectors (Tecan, Lyon, France) was set at 340 nm to monitor the absorbance over time at 30°C. Briefly, a reaction mixture containing HsNMT1*l* at different concentrations of peptide acceptors was pre-incubated for 3 min at 30°C before starting the reaction with MyrCoA.

Myristoylation kinetics were monitored continuously for 30 min, and the data were fitted as to obtain the initial velocities associated to each peptide concentration. Curve fits to obtain kinetic parameters were achieved by non-linear regression with GraphPad Prism 9.1 (GraphPad Software, La Jolla, CA). Parameters with standard errors were computed for all parameters using the complete dataset including replicates. Both *k*_cat_ and *K*_m_ kinetic parameters were obtained by fitting to the Michaelis–Menten equation. *k_cat_/K_m_* values and the associated standard deviations were obtained by taking advantage of the *k_SP_* approach^44^ with *k_SP_*=*k_cat_/K_m_* and v_0_/[E]= *k_SP_*[S]/(1+[S]/*K_m_*), where v_0_ is the reaction rate measured at NMT concentration [E_0_].

For inhibition assays, we used the same reaction mixture for 30 min at 30°C incubation in the presence of the inhibitor peptide and MyrCoA. The reaction was triggered by the addition of SOS3 peptide and monitored for 30 min. Inhibition curves and associated IC_50_ values were fitted with the absolute IC_50_ module with baseline set at 0. Inhibition constants were calculated as described in ^45–47^ with *K_I_* = (IC_50_-[E]/2)/(1+[S]/*K_m_*). The HsNMT1 concentration was 0.25 μM, and the substrate (G_2_CSVKKK; SOS3; *K_m_*=18±3 μM^43,48^) was used at a concentration of 20 μM.

The NMT/IpaJ pipeline described in **Fig. 5b** first involved incubation of the peptide (100 μM) for 1 hour in the presence of 0.5 μM HsNMT1 or HsNMT2 (T1). The buffer was the same as for NMT activity measurement but contained cholate (5 μM) instead of Triton X-100. Full IpaJ cleavage conditions involved further incubation at 20°C for 1 hour in the presence of 10 μM IpaJ-3M (T2). 5 μM HsNMT1 was finally added for 60 min at 30°C (T3). T0 corresponded to the T3 time (3 hours) with the peptide diluted in the incubation buffer but in the absence of any enzyme.

### Mass spectrometry

300 μL of a mixture containing 50 mM Tris (pH 8), 0.193 mM EGTA, 1 mM MgCl_2_, 1 mM DTT, 5 μM sodium cholate, 40 μM Myr-CoA solution (stock solution 0.2 mM Myr-CoA, 10 mM sodium acetate, 2.5 μM sodium cholate), 0.5 μM NMT, and 100 μM of synthetic peptide were pre-incubated at 30°C. The myristoylation reaction was followed over time by collection of 10 μL samples further diluted in 90 μL of water/acetonitrile (90/10) solution. The different samples were then diluted five times in the matrix solution made of 5 mg/mL of α-cyano-4-hydroxycinnamic acid solubilized in water/formic acid/acetonitrile (50/50/0.1%). 1 μL of each dilution was spotted on a metal target and dried. MS and MS/MS spectra of each sample were acquired with an AB SCIEX 5800 MALDI-Tof-Tof instrument in positive ion mode as reported in SI. Survey scans were performed using delayed extraction (390 ns) in reflector mode for a total of 15,000 shots. MS/MS scans were operated with a collision energy of 1 kV. Peptide and fragment mass tolerances were set at 10 ppm and 0.8 Da, respectively. Mass spectra were analyzed with Peakview software (AB Sciex, Macclesfield, UK), and MS2 analysis was performed with ProteinProspector v6.2.2 (https://prospector.ucsf.edu/prospector/mshome.htm)^49^. All spectra are available in **Supplementary Dataset 1**. Theoretical mass values were calculated at https://web.expasy.org/peptide_mass/.

## EXTENDED DATA

**Extended Data 1:**
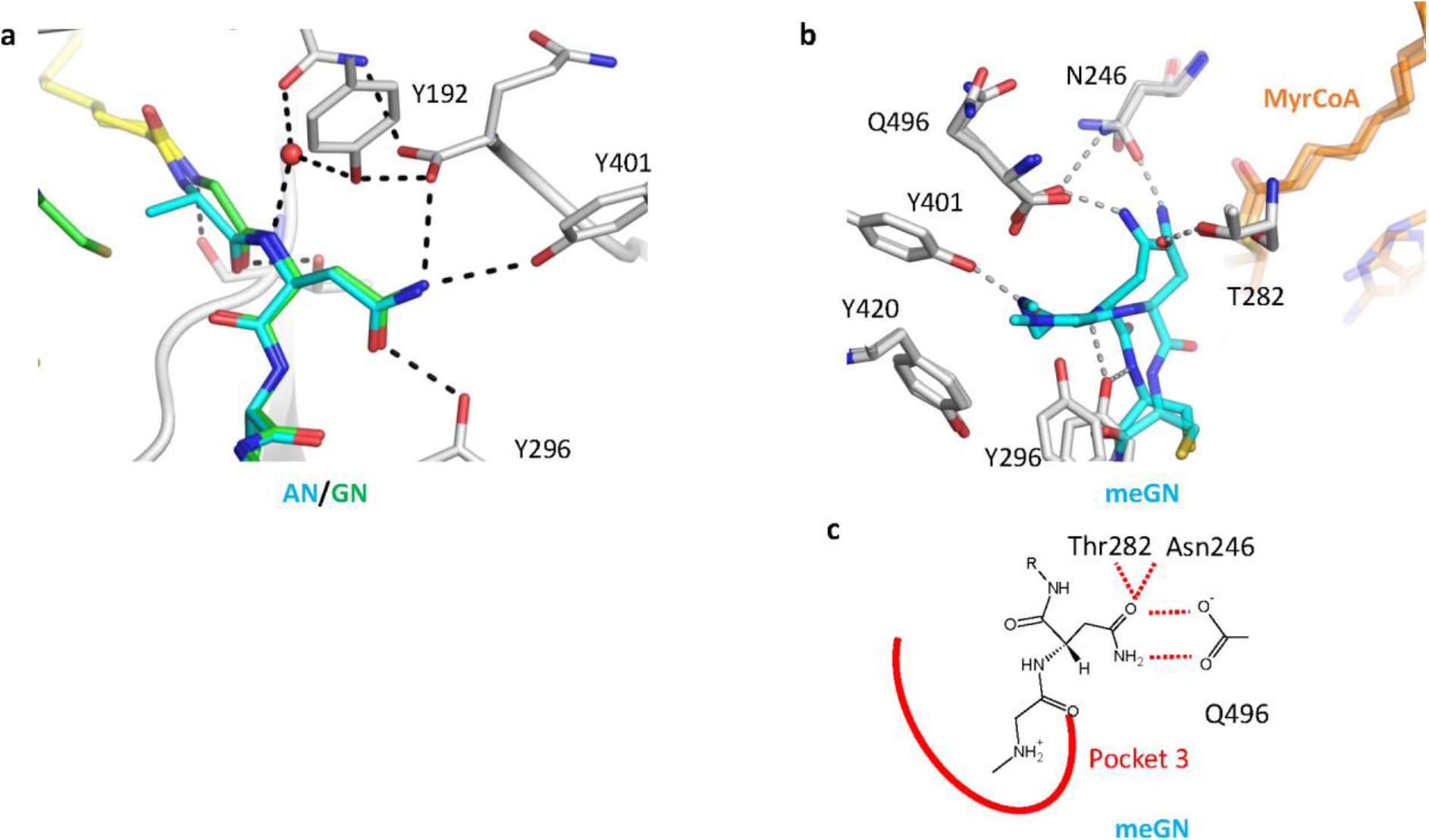
Structural impact of the addition a methyl group on aa1. Derivatives of the reference peptide (G_2_KSFSKPR) at positions 2 and/or 3 (GN and AN) were studied, and the crystal structures of their complexes with HsNMT1 were determined. (**a**) Superimposition of the GN (green) and AN derivatives (light blue). (**b**) Overlap of the two alternative conformations of the N-methyl GN derivative (meGN). (**c**) Plane structure of the environment at the active site of the meGN derivative.

**Extended Data 2:**
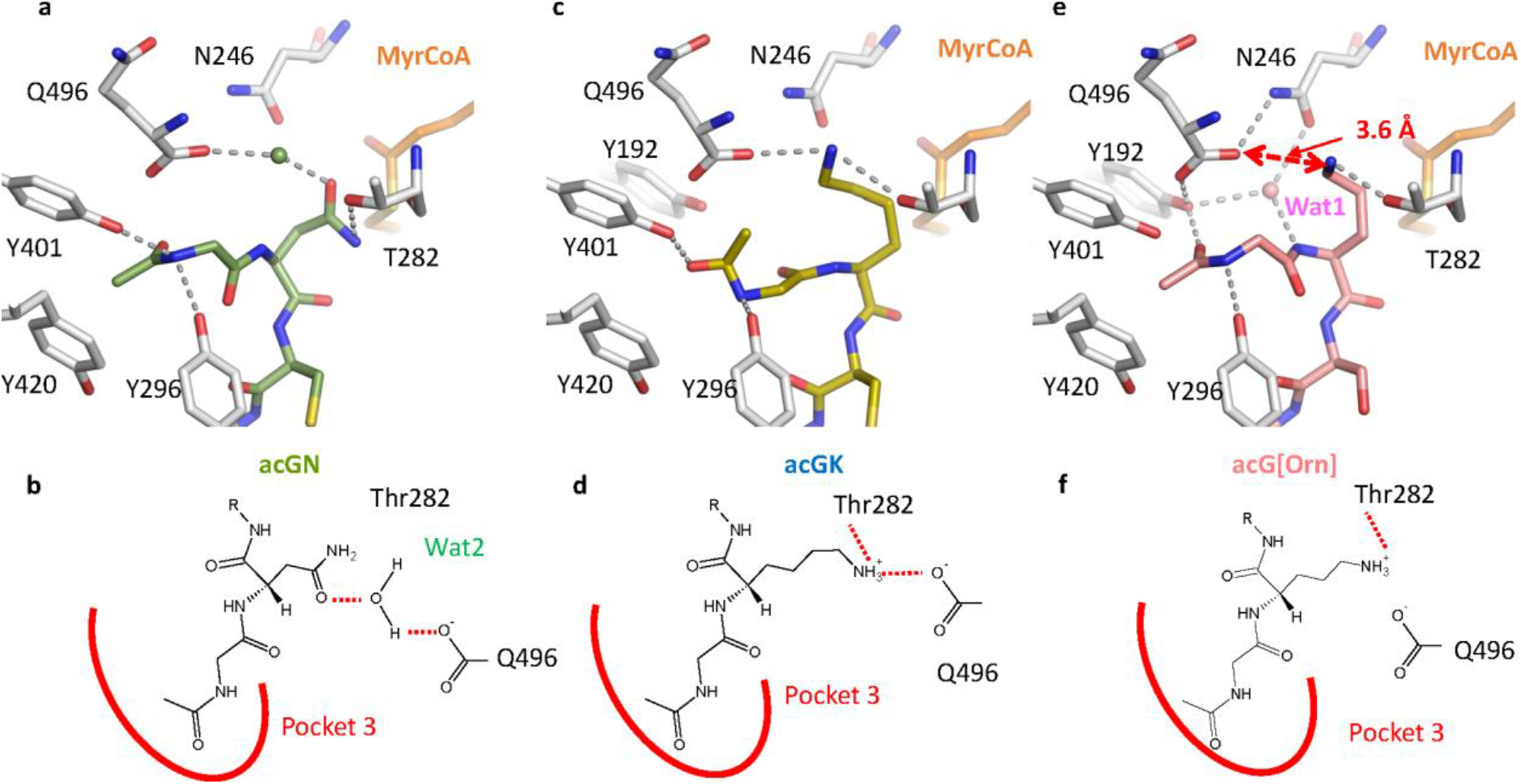
Structural impact of the nature of the side chain at position 3 on myristoylation. All three peptides are acetylated (ac) derivatives of the GK peptide (**Fig. 2a**). The 3D close up of the active site around the N-terminal extremity of the peptide ligand is displayed in the top panels together with a plane version of the corresponding peptides (bottom panels). (**a**) 3D and (**b**) plane close-ups of the active site environment with acGN; (**c**) 3D and (**d**) plane close-ups with acGK (data from ^17^, PDB entry 6SK2, subunit A). (**e**) 3D and (**f**) plane close-ups with acG[Orn].

**Extended Data 3:**
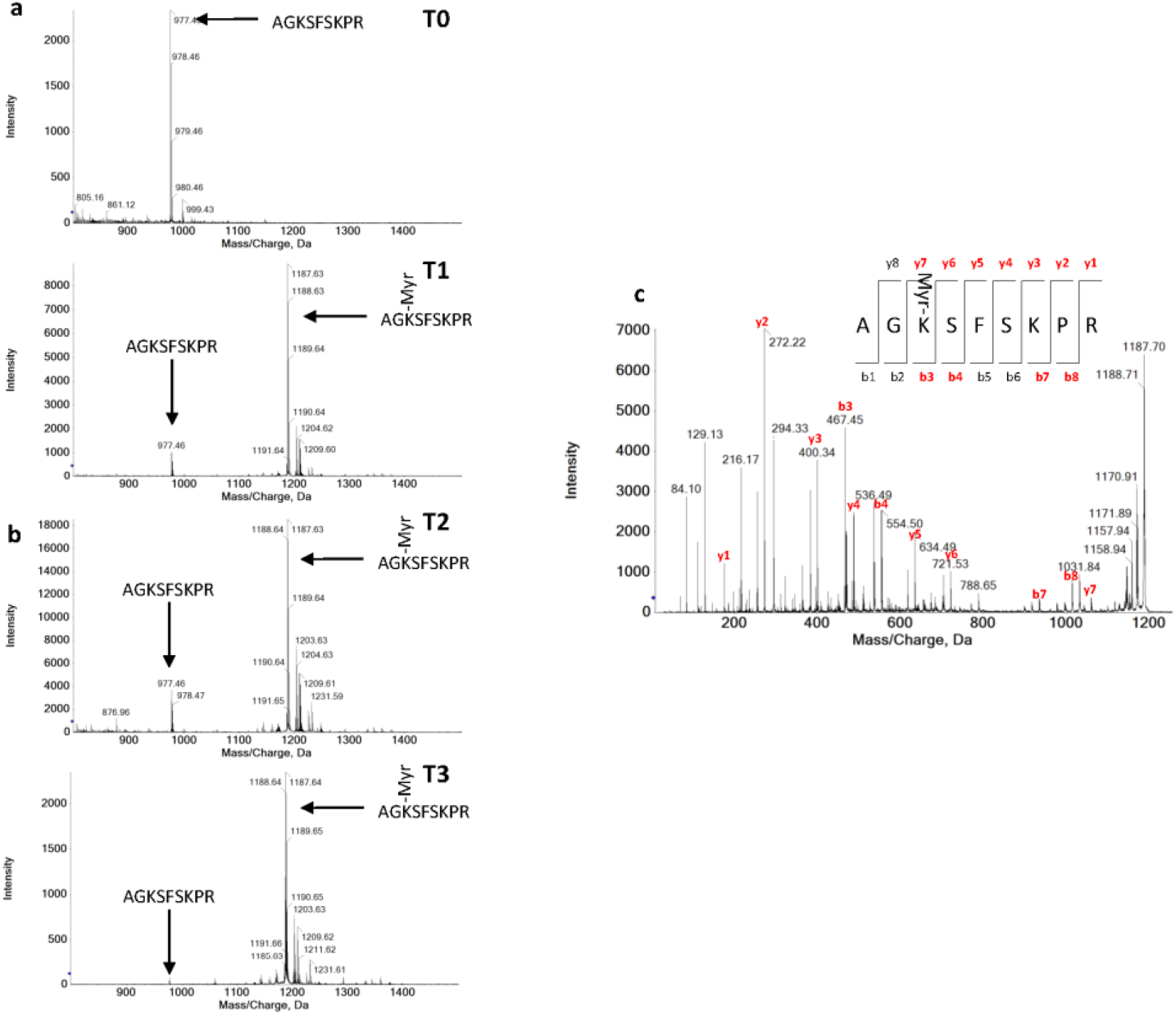
Mass spectrometry analysis of peptide <u>AG</u>KSFSKPR showing K4 myristoylation. (**a**) and (**b**) Analysis at various time points by MS1. (**c**) MS2 analysis of the 1,187 Da peptide.

**Extended Data 4.**
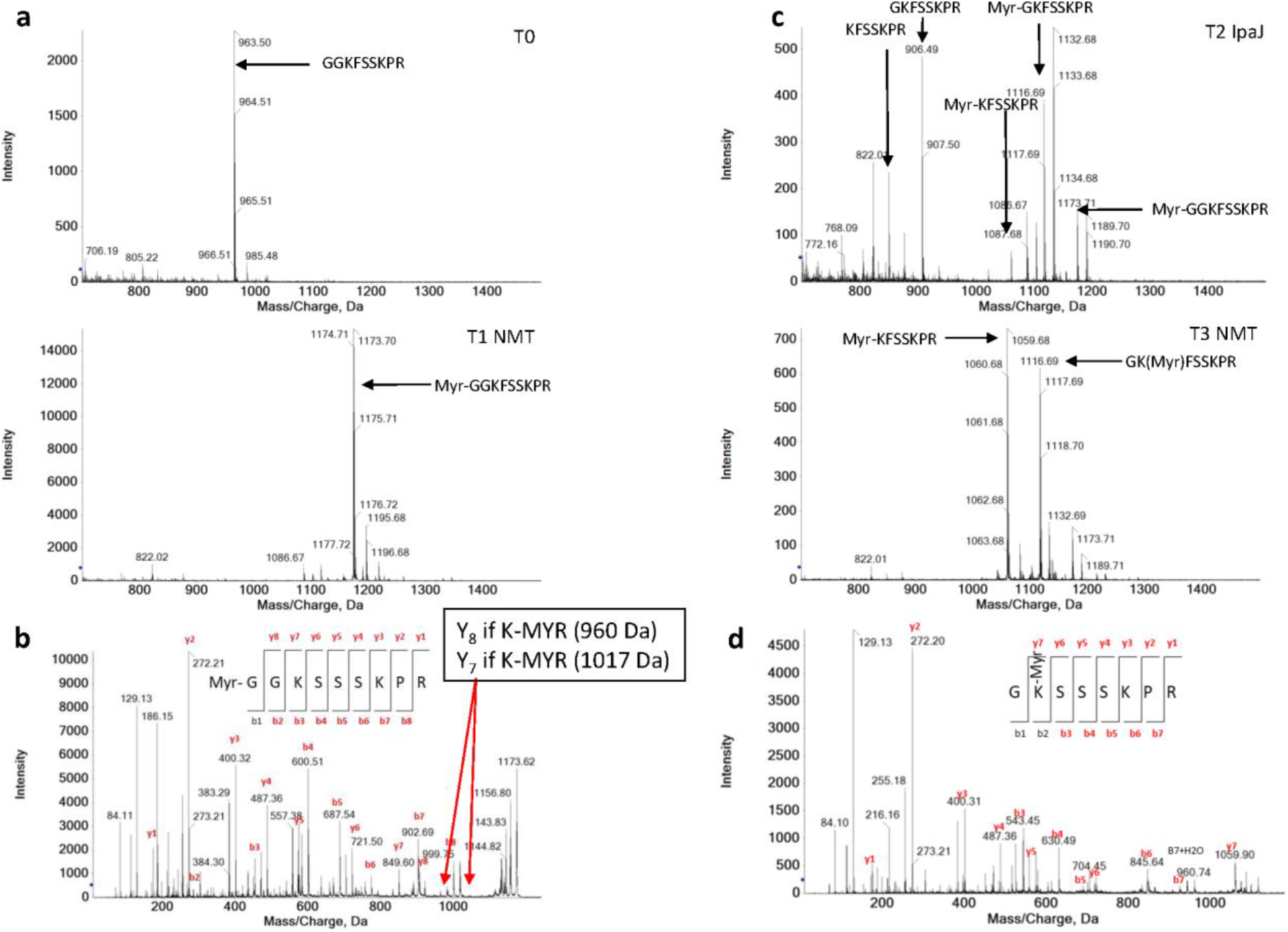
IpaJ action reveals that a substrate starting with G may feature competition between G- and K-myristoylation. Peptide GGKFSSKPR was used as the substrate. (**a**) MS1 of various spectra recorded at T0 (peptide only), T1 (incubation with 0.5 μM HsNMT1 with MyrCoA for 10 min), IpaJ treatment for 120 min (T2), and further HsNMT1 incubation (5 μM for 120 min, T3). (**b**) MS2 spectra of the 1116 Da ion after T3. The 1059 y_7_ ion is characteristic of K-myristoylation. No b_1_ ion is characteristic of G-myristoylation at 267 Da. **Fig. 3c,d** show the analysis of the intermediate derivative GKFSSKPR to confirm K-myristoylation. No double myristoylation was observed in either case.

**Extended Data 5:**
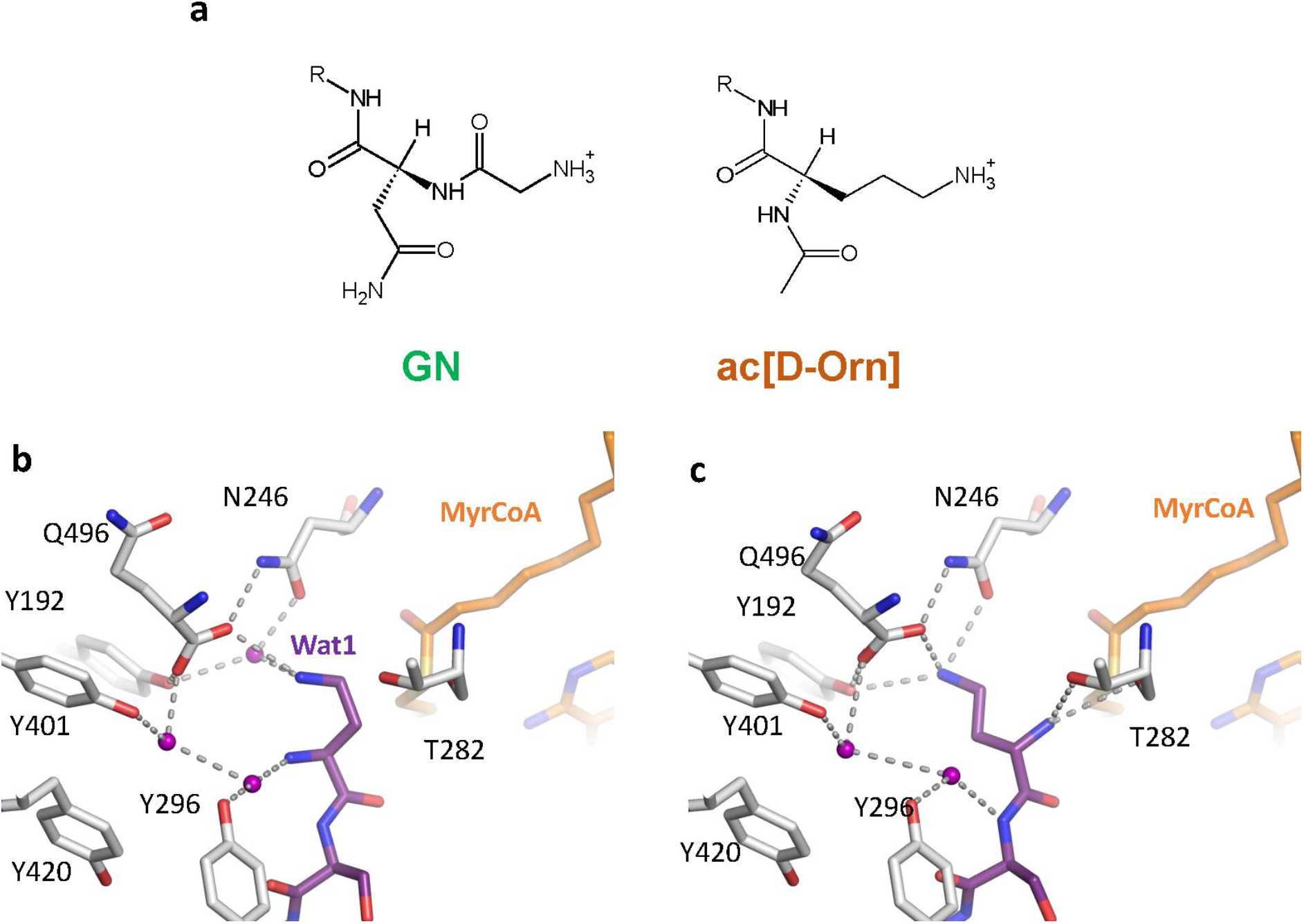
Two conformations of Dab make direct interaction with the catalytic base. (**a**) Comparison of the plane structure of GN and ac[D-orn] showing their very similar structure. (**b**) and (**c**) Close up of the active site of HsNMT1 crystallized with the Dab derivative showing the two alternative conformations of the reactive side chain of Dab entering the active site next to the catalytic base, the C-terminus of Q496. (**b**) Subunit A, (**c**) Subunit B. NB Wat1 does not make any contact with the Dab peptide and is a recurrent interactor in most peptide NMT complexes with Y192 and Y180 (see ^17^).

