## Supplementary Figures for "Proteome-wide probing of the dual NMT-dependent myristoylation tradeoff unveils potent, mechanism-based suicide inhibitors"

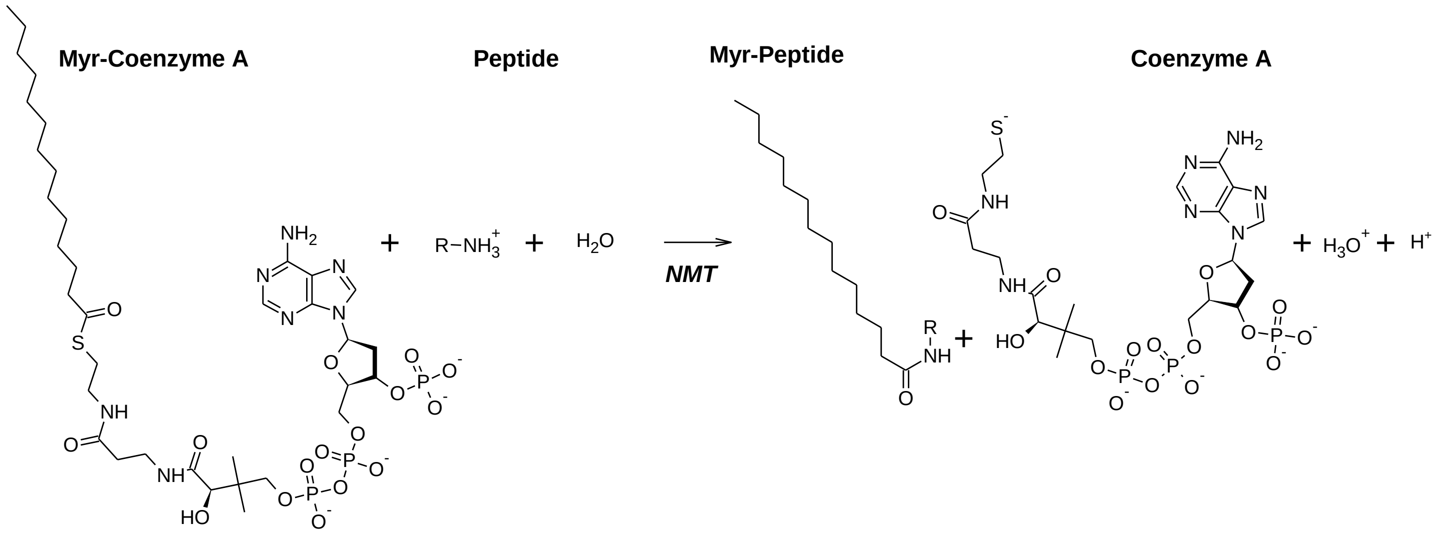


**Supplementary Figure 1. The reaction catalyzed by NMT**
NMT-catalyzed G-myristoylation. The use of a water molecule for the reaction is indicated.

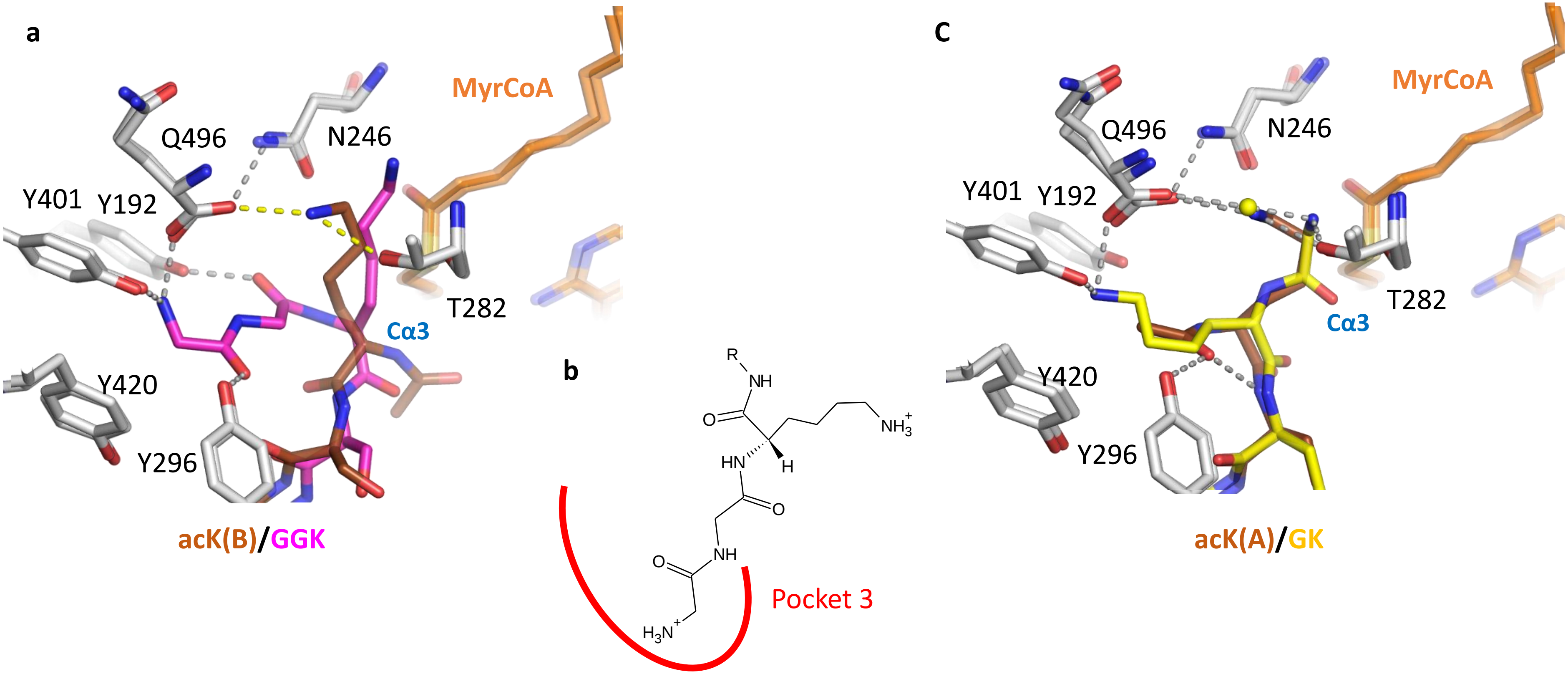


**Supplementary Figure 2: Two alternative paths involving rotation around Cα3 of the main chain may lead to K-myristoylation**

Close up of the 3D structure of the active site of variants of the reference peptide. (**a**) Overlap of acK (B subunit) with GGK. (**b**) Overlap of GK (HPCA) with acK (A subunit).

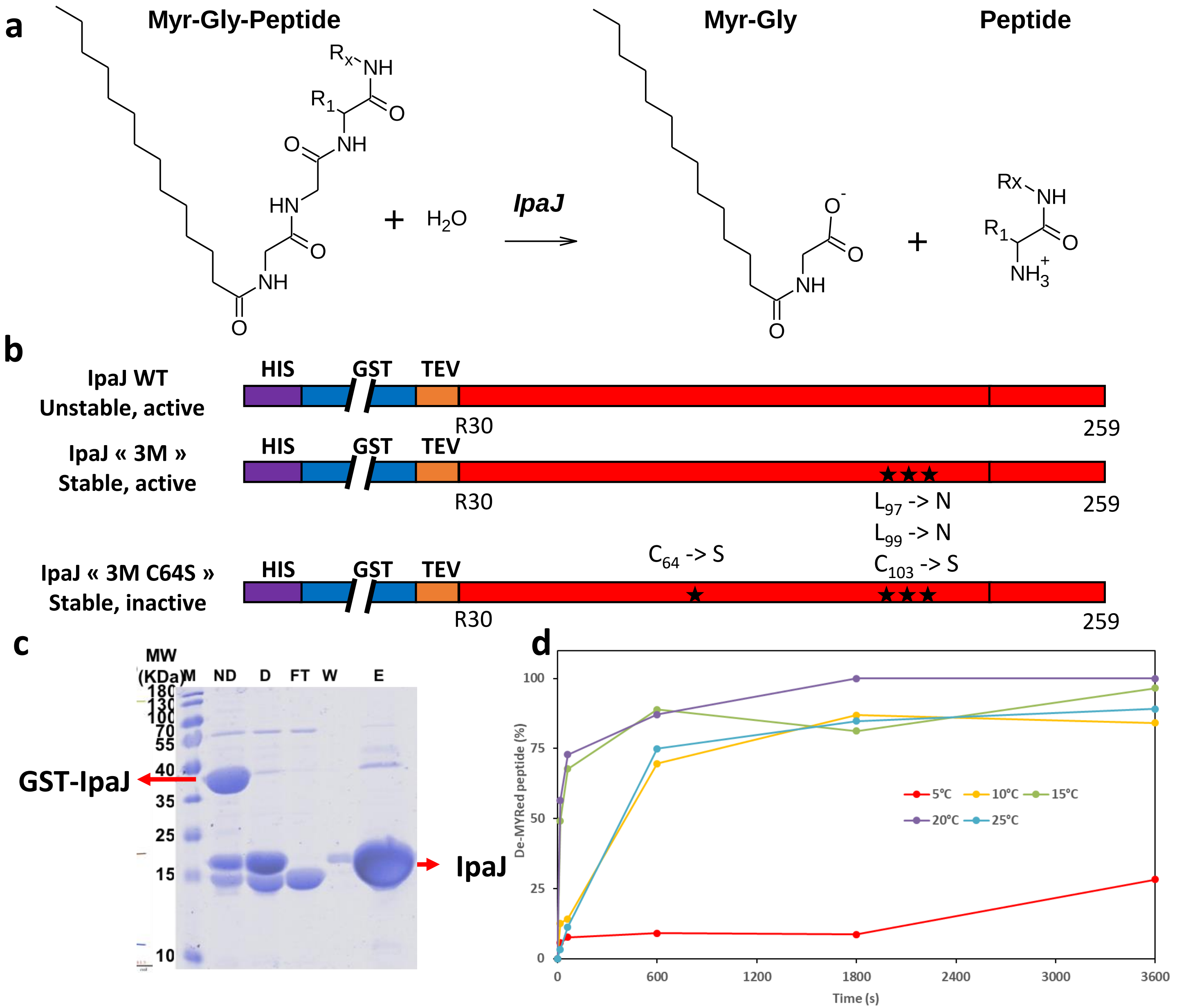


**Supplementary Figure 3: A stable variant of IpaJ allows efficient peptide cleavage at low temperatures**

(**a**) Scheme summarizing the IpaJ-catalyzed chemical reaction. (**b**) Cartoon summarizing the three constructs used in this study. (**c**) Purification of IpaJ as a GST fusion and GST cleavage with protease. (**d**) Impact of temperature incubation on the activity of IpaJ-3M; peptide Myr-GQGPSGGLNR was used as the substrate. IpaJ was used at 1 µM with 100 µM peptide and incubated for 60 min to allow visualization of the various species by MS. The cleaved/uncleaved peak intensity yield is plotted as a function of time.

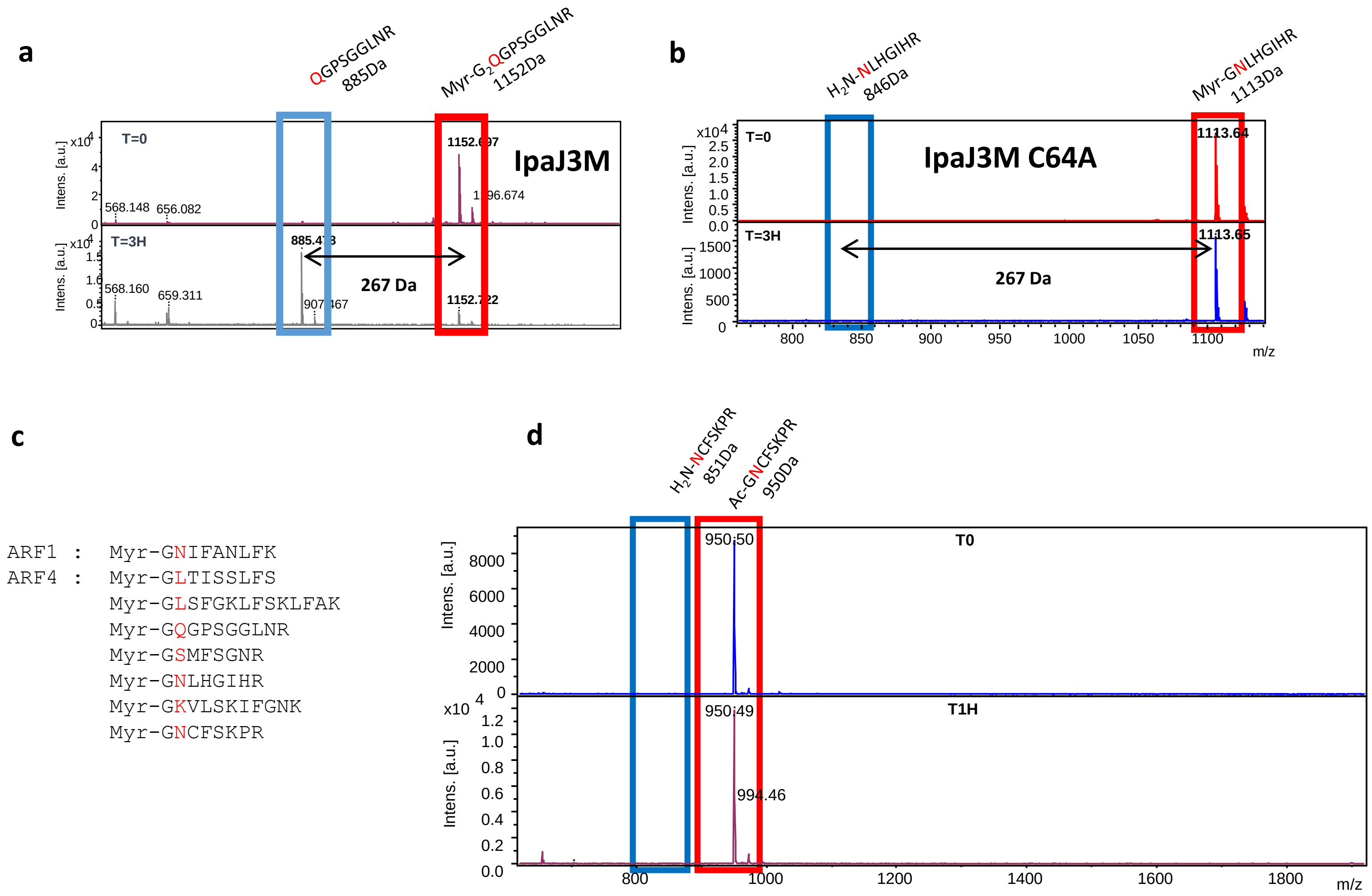


**Supplementary Figure 4: The tight substrate selectivity of IpaJ**

(**a**) Cleavage of a peptide by IpaJ; IpaJ was used at a low (1 µM) concentration and incubated for 10 min to allow visualization of the various species. (**b**) Demonstration that variant IpaJ C64A is inactive. Enzyme concentration was 10 µM and incubation time was 180 min. (**c**) Various tryptic peptides used to challenge IpaJ *in vitro*. (**c**) IpaJ-3M does not cleave either Myr-Ala or acetyl-Gly derivatives. IpaJ concentration was 10 µM and incubation time was 180 min.

a

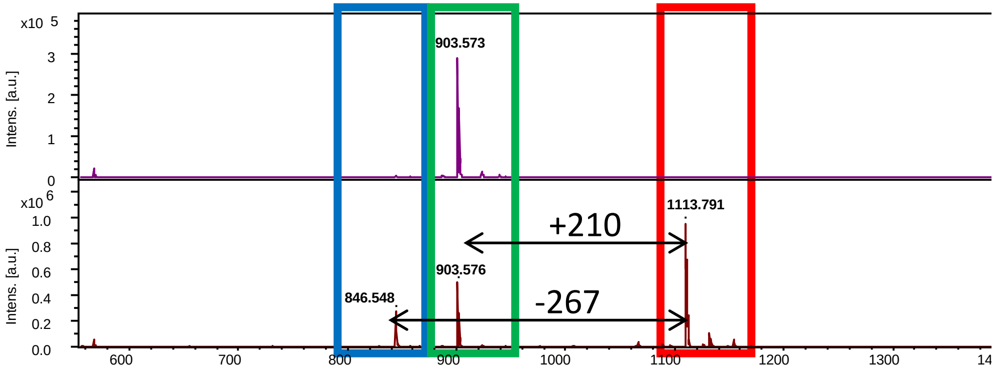


b

| | |
|---|---|
| $H_2N$-$G_2$IGWSLHN | $H_2N$-$G_2$FCFSKFG |
| $H_2N$-$G_2$NLHGIHR | $H_2N$-$G_2$RKWSGPT |
| $H_2N$-$G_2$CFHSKAA | $H_2N$-$G_2$TSVSKPV |
| $H_2N$-$G_2$HRHSKSK | $H_2N$-$G_2$MVFGKIA |
| $H_2N$-$G_2$KVLSKIF | $H_2N$-$G_2$GTTSTRR |

c

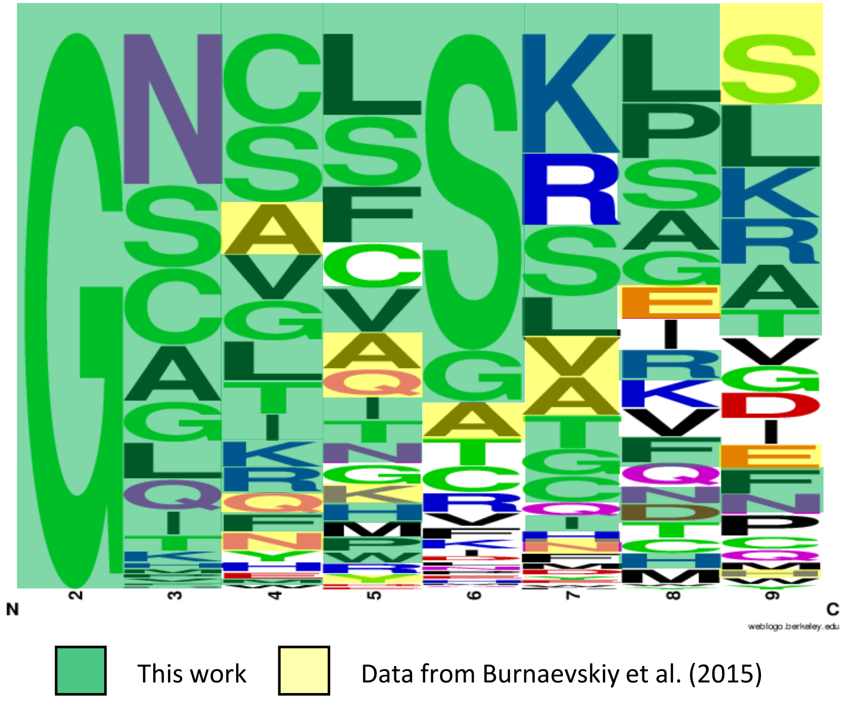


**Supplementary Figure 5: MALDI-TOF analyses with the NMT-IpaJ workflow**

(**a**) Example of the behavior of an NMT substrate at T0 and T2. (**b**) Various peptide sequences verified in the current analysis. (**c**) Logo recapitulating the sequence space covered by our (colored in green) or other (yellow; see [23] experiments).

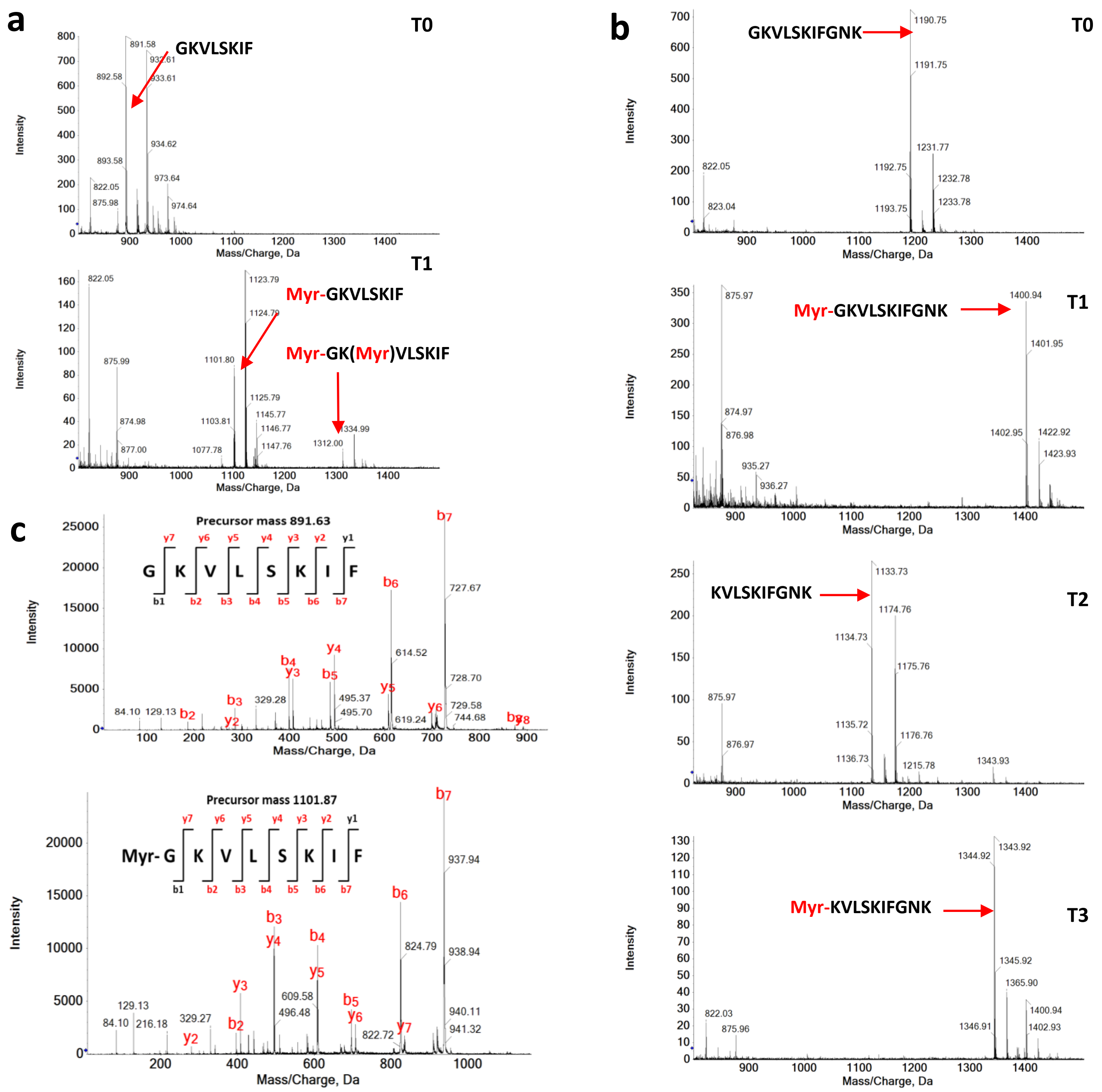


**Supplementary Fig. 6. The many ARF6 myristoylation products as revealed by the IpaJ pipeline**

(**a**) The indicated octapeptide derived from ARF6 was incubated in the presence of HsNMT2. (**b**) An 11-mer version of the ARF6 peptide was subjected to the IpaJ pipeline; MS1 MALDI spectra are displayed at T0, T1, T2, and T3 to reveal four isoforms. (**c**) MS2 spectra of the unmodified and mono modified short ARF6 peptide, from data in panel a.

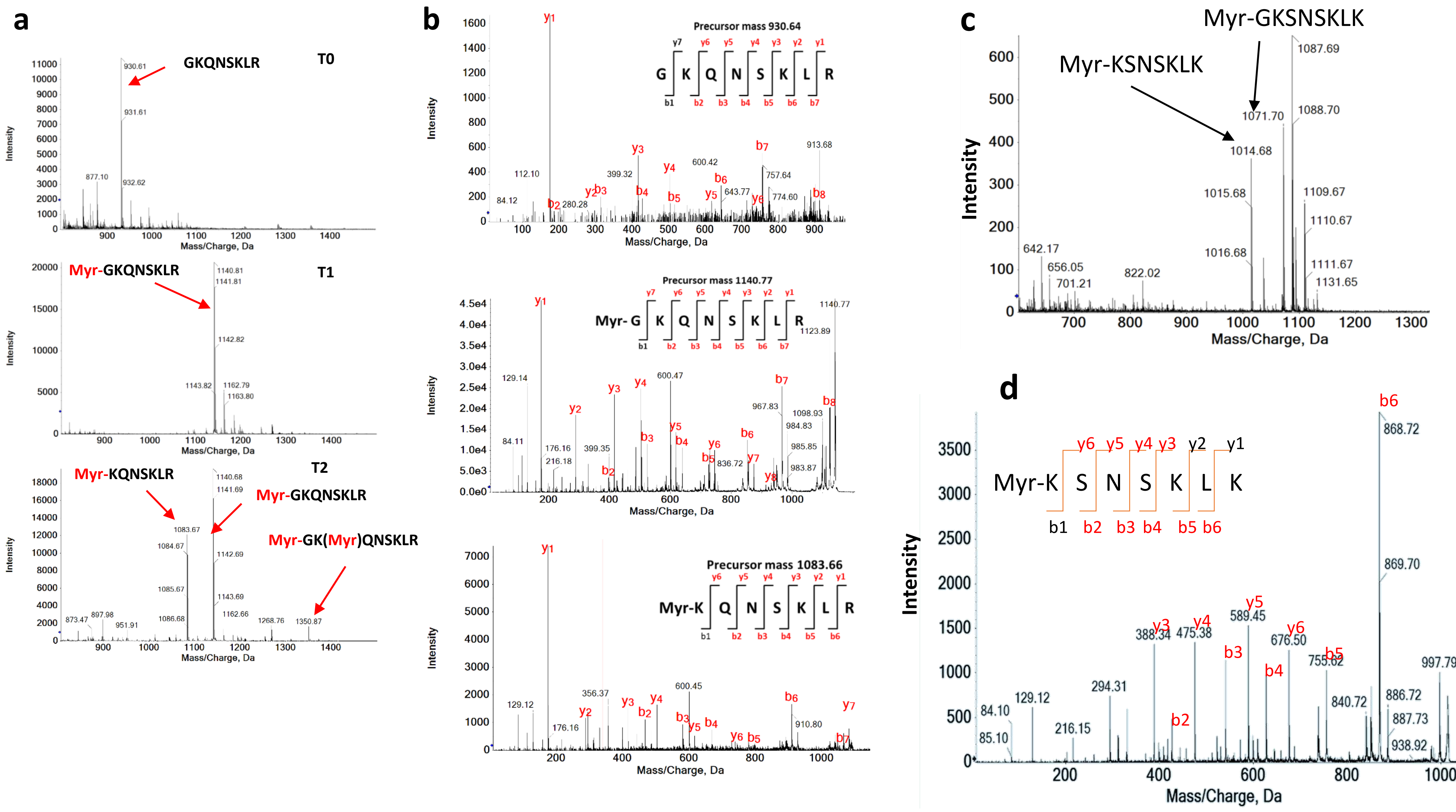


**Supplementary Figure 7. Neuron-specific calcium-binding proteins may undergo all the many different interconversions before and upon IpaJ cleavage**

Two octapeptides were derived from neuron-specific calcium-binding protein hippocalcin isoforms (HPCA/HPCL1/NCALD, ID P84074/P37235/P61601) and neuronal calcium sensor 1 (NCS1; P62166), respectively. The NMT/IpaJ pipeline was applied in the presence of HsNMT1, and the various products were identified. (**a**) HPCA peptide at T0, T1, and T2. (**b**) MS2 spectrum of the double myristoylation product from T2 (panel A). (**c**) MS2 spectrum of the mono myristoylation products from T0, T1, and T2 (panel a). (**d**) NCS1 MS1 spectrum at T2 (**e**) NCS1 MS2 spectrum of the 1014 Da species at T2.
