## Supplementary Table 1 for "Proteome-wide probing of the dual NMT-dependent myristoylation tradeoff unveils potent, mechanism-based suicide inhibitors"

**Supplementary Table 1: List of all peptides and diagnoses**

MALDI and kinetic data of peptides used in this study. MS spectra are displayed in **Supplementary Dataset 1**. Reference peptide sequence is GKSFSKPR. The page number is indicated in one column of each table. (**a**) Peptides displaying G-myristoylation, including those from the human proteome. (**b**) All peptides with K-myristoylation. (**c**) Series with variations on the side chains at positions 6 and 7. (**d**) IpaJ cleavage on various peptides. (**e**) myristoylation after IpaJ cleavage (T2 and T3).

#### Supplementary Table 1a: GK-starting peptides derived from the human proteome reveal N-terminal alpha-MYR as the major modification made by NMT

| Series[a] | Peptide sequence | Short name (text & Figures) | MS data page in Supplementary Dataset 1a | Crystal structure | $k_{cat}$ ($s^{-1}$) | $K_m$ (µM) | $k_{cat}/K_m$ ($s^{-1}.M^{-1}$) | Relative $k_{cat}/K_m$ (%)[c] |
|---|---|---|---|---|---|---|---|---|
| Reference GK-peptide variants | GKSFSKPR | GK | 2 | - | 0.050 ± 0.0002 | 10 ± 2 | 4 931 ± 626 | 165 |
| | G{Orn}SFSKPR | | 3 | - | 0.018 ± 0.001 | 6 ± 2 | 2808 ± 951 | 94 |
| | G{Dab}SFSKPR | | 4 | - | 0.015 ± 0.002 | 16 ± 7 | 935 ± 362 | 31 |
| | ANCFSKPR | AN | 5 | 2.16, ED1a | 0.0004 ± 0.0001 | >300 | 1.2 ± 1.1 | 0.05 |
| | AGKFSKPR | | 6 | - | NM | NM | <0.02 | <0.01 |
| | me-GNCFSKPR | meGN | 7 | 3, ED1b | 0.0018 ± 0.0001 | 1 068 ± 51 | 1.65 ± 0.05 | 0.06 |
| | GKSWSKGR | | 8 | - | 0.090 ± 0.002 | 9 ± 1 | 10 000 | 334 |
| | GKFSSKPR[b] | GKFS | 9/10 | - | 0.11 ± 0.02 | 346 ± 75 | 330 ± 33 | 11 |
| Human GKX | GKVLSKIF | ARF6 | 11 | - | 1.01 ± 0.04 | 194 ± 14 | 5 215 ± 436 | 174 |
| | GKQNSKLR | HPCA | 12 | 1.5, Fig.1a,c | 1.03 ± 0.03 | 204 ± 11 | 5 055 ± 320 | 169 |
| | GKTNSKLA | HPCL4 | 13 | - | 0.58 ± 0.06 | 48 ± 18 | 12 026 ± 3 560 | 401 |
| | GKQNSKLA | VIS | 14 | - | 0.092 ± 0.004 | 58 ± 11 | 1 589 ± 265 | 53 |
| | GKSNSKLK | NCS1 | 15 | 5O9S, Ref.Castrec | 0.047 ± 0,001 | 15 ± 1 | 3 157 ± 277 | 105 |
| | GKSLSHLP | T106B | 16 | - | 0.31 ± 0.04 | 811 ± 224 | 386 ± 66 | 13 |
| | GKTFSQLG | T106A | 17 | - | 0.43 ± 0.03 | 200 ± 53 | 2 147 ± 467 | 72 |
| | GKSASKQF | GAPP | 18 | | 0.065 ± 0.04 | 69 ± 17 | 952 ± 198 | 32 |
| | GKLHSKPA | NKD1 | 19 | - | 0.10 ± 0.01 | 40 ± 2 | 2 500 ± 200 | 83 |
| | GKLQSKHA | NKD2 | 20 | - | 0.10 ± 0.01 | 41 ± 2 | 2 501 ± 200 | 83 |
| GXK | GAKQSGPA | ZNRF2 | 21 | - | | | 103 | 3 |
| | GGKQSTAA | ZNRF1 | 22 | | 1.45 ± 0.23 | 2 320 ± 573 | 623 | 21 |
| | GGKFSSKPR | | 23/24 | - | 0.14±0.01 | 85 ± 32 | 1 647 | 21 |
| | GGKLSKKK | BASP | 25 | - | | | 3 900 | 130 |
| SOS3 | GCSVSKKK | SOS3 | 26 | - | 0.055 ±0.002 | 18 ± 3 | 2 997 ± 415 | **100** |
| | ASSVSKKK | | 27 | - | NM | NM | <1 | <0.01 |

[a] X means any aminoacid or N-acetyl (ac), which corresponds to a Gly devoid of $NH_2$, the reactive moiety for G-MYR

[b] both G- and K-MYR

**Supplementary Table 1b: Impact of the S6K7 motif on G-MYR**

| Peptide sequence | MS data page in SupDataset 1b | WT | | |
|---|---|---|---|---|
| | | $k_{cat}$ ($s^{-1}$) | $K_m$ (µM) | $k_{cat}/K_m$ ($s^{-1}.M^{-1}$) |
| GKSNSKLK | 2 | 0.047±0.001 | 15 ± 2 | 3133 |
| GKSNSGLKP | 3 | NM | NM | 2.2±0.2 |
| GKSNS{DAP]LKP | 4 | 0.280±0.002 | 1042±113 | 269 |
| GKSNS{DAB]LKP | 5 | 0.096±0.003 | 23±4 | 4174 |
| GKSNS{ORN]LKP | 6 | 0.16±0.005 | 29±4 | 5517 |
| GKSNS{HCY}LKP | 7 | NM | NM | 40 |
| GKSNS{bhLYS}LKP | 8 | 0.10±0.01 | 92±26 | 1087 |
| GKVWSQGV | 9 | 0.123 | 329 | 373 |
| GKSNAKLKP | 10 | NM | NM | 69 |

### Supplementary Table 1c: Substitutions around the N-terminus of peptides promoting side chain MYR or NMT inhibition

| Series [a] | Peptide sequence | Shortname (text & Figures) | Series sequence[d] | MS data page in Supplementary Dataset 1c | Crystal structure[e] | Kinetic parameters with HsNMT1[f] | | |
|---|---|---|---|---|---|---|---|---|
| | | | | | | $k_{cat}$ ($s^{-1}$) | $K_m$ (µM) | $k_{cat}/K_m$ ($s^{-1}.M^{-1}$) |
| acGK | ac-GKSFSKPR | acGK | | 2 | 6SK2, ED2e,f, Ref.[illegible] | 0.0017 ± 0.0002 | 58 ± 15 | 30 ± 6 |
| | ac-G{Orn}SFSKPR | acG[Orn] | | 3 | 1.81, ED2,cd | 0.0006 ± 0.0001 | 22 ± 3 | 29 ± 3 |
| | ac-G{Dab}SFSKPR | acG[Dab][c] | | 4 | - | NM | NM | <<0.01 |
| | ac-G{D-Orn}SFSKPR | acG[D-Orn][c] | | 5 | - | NM | NM | <<0.01 |
| | ac-GNCFSKPR | acGN | | 6 | ED2a,b | NM | NM | <<0.01 |
| | ac-GKVLSKIF | acGK | ARF6 | 7 | - | 0.0019 ± 0.0001 | 37 ± 4 | 51 ± 6 |
| XK | AKQNSKLR | A-HPCA | HPCA | 8 | - | >0.0004 | >1000 | 0.4 |
| | AKVLSKIF | A-ARF6 | ARF6 | 9 | - | 0.0005 ± 0.0001* | 8 ± 3* | 64 ± 18 |
| | AKSFSKPR | AK | | 10 | Fig.2b,c | 0.0006 ± 0.0002* | 14 ± 5* | 40 ± 11 |
| | AKPTSKDSGLK | TCS2 | TCS2 | 11 | - | NM | NM | <<0.01 |
| | A{Orn}SFSKPR | A{Orn}[c] | | 12 | - | NM | NM | <<0.01 |
| | A{D-Orn}SFSKPR | A{D-Orn}[c] | | 13 | - | NM | NM | <<0.01 |
| | SKSFSKPR | SK | | 14 | - | 0.0025 ± 0.0003 | 43 ± 23 | 60 ± 27 |
| | MKSFSKPR | MK | | 15 | - | >0.0001 | >100 | 1.7 |
| | PKSFSKPR | PK | | 16 | - | 0.00068 ± 0.00005 | 26 ± 8 | 26 ± 6 |
| | G{D-ORN}SFSKPR | G[D-Orn][c] | | 17 | - | NM | NM | <<0.01 |
| | ac-KSFSKPR | acK | | 18 | 1.7, Fig.4a,b | 0.0009 ± 0.0001 | 24 ± 10 | 37 ± 13 |
| | ac-{Orn}SFSKPR | ac[Orn][c] | | 19 | - | NM | NM | <<0.01 |
| | ac-{D-Orn}SFSKPR | ac[D-Orn][c] | | 20 | Fig.5d,e | NM | NM | <<0.01 |
| | ac-{D-Dab}SFSKPR | ac[D-Dab][c] | | 21 | - | NM | NM | <<0.01 |
| | ac-{Dap}SFSKPR | ac[Dap] | | 22 | - | NM | NM | <<0.01 |
| K | K | K | | 23 | - | 0.00043 ± 0.00004 | 16 ± 6 | 27 ± 6 |
| | {Orn}SFSKPR | [Orn] | | 24 | - | >0.0002 | | 2 |
| | {D-Orn}SFSKPR | [D-Orn][c] | | 25 | - | NM | NM | <<0.01 |
| | {Dab}SFSKPR | [Dab][c] | | 26 | ED5 | NM | NM | <<0.01 |
| | KQNSKLRP | K-HPCA | HPCA | 27 | ? | 0.0013 ± 0.0002 | 133 ± 49 | 9 |
| | ac-KFSKPR | acKΔS | | 28 | - | NM | NM | <<0.01 |
| | KSFS{Orn}PR | | | 29 | - | 0.0029 ± 0,0002 | 59 ± 15 | 49 |
| | KSFS{Dab}PR | | | 30 | - | 0.0024 ± 0,0009 | 145 ± 109 | 17 |
| | KSFS{Dap}PR | | | 31 | - | 0.0012 ± 0.00003 | 39 ± 4 | 31 |
| | ac-[D-Orn}SNSKLK | | NCS1 | 32 | - | | | 1.5 |
| | KSNAKLKP | | NCS1 | 33 | - | NM | NM | <<0.01 |
| ZXGK | GGKSFSKP | GGK | | 34/35 | SupFig.2 | 0.089 ± 0.012 | 882 ± 263 | 101 ± 17 |
| | AGKSFSKPR | AGK | | 36 | - | 0.0024 ± 0.0004 | 31 ± 3 | 76 ± 7 |
| | ac-GGKSFSKPR | acGGK | | 37 | - | 0.00062 ± 0.00005 | 82 ± 24 | 8 ± 2 |
| | ac-GGGKSFSKPR | acGGGK | | 38 | - | NM | NM | <<0.01 |
| | ac-GGGGKSFSKPR | | | 39 | - | NM | NM | <<0.01 |
| | ac-GGGGGKSFSKPR | | | 40 | - | NM | NM | <<0.01 |

* only concentrations lower than 50 µM were taken into account as susbstrate concentration inhibition was observed at higher concentrations

**Supplementary Table 1d: IpaJ acts as a cysteine protease with strict specificity for alpha MYR-Gly/peptide cleavage**

| Sequence | Cleavage by IpaJ | MS data page in Supplementary Dataset 1e |
|---|---|---|
| Myr-GLSFGKLFSKLFAK | + | 2 |
| Myr-GSMFSGNR | + | 3 |
| Myr-GQGPSGGLNR | + | 4 |
| Myr-GNLHGIHR | + | 5 |
| Myr-GKVLSKIFGNK | + | 6 |
| Myr-GNCFSKPR | + | 7 |
| ac-GNCFSKPR | - | 8 |
| Myr-ATNGSKVA | - | 9 |
| | **IpaJ cleavage after MYR** | |
| GIGWSLHN | + | 10 |
| GNLHGIHR | + | 11 |
| GCFHSKAA | + | 12 |
| GHRHSKSK | + | 13 |
| GMVFGKIA | + | 14 |
| GGTTSTRR | + | 15 |
| GKVLSKIF | + | 16 |
| GFCFSKFG | + | 17 |
| GRKWSGPT | + | 18 |
| GTSVSKPV | + | 19 |

#### Supplementary Table 1e: Epsilon myristoylation is induced after IpaJ cleavage

| Peptide sequence | Delta mass (Da) after | Myr after cleavage | Myr-peptide after IpaJ cleavage | MS data page in Supplementary Dataset 1e |
|---|---|---|---|---|
| met-GNCFSKPR | Nd | No | - | 2 |
| GKSFSKPR | 267,2 | Yes | + | 3 |
| G{ORN}SFSKPR | 267,26 | Yes | + | 4 |
| GKVLSKIFGNK | 267,21 | Yes | + | 5/6/7/8 |
| GKVLSKIFGN | 267,22 | Yes | + | 5/6/7/8 |
| GKVLSKIF | 267,41 | Yes | + | 5/6/7/8 |
| Myr-GKVLSKIFGNK | 267,21 | Yes | + | 9 |
| G-K(ε-MYR)VLSKIFGNK | Nd | No | + | 10 |
| GKFSSKPR | 267,2 | Yes | + | 11 |
| GKQNSKLR | 267,35 | Yes | + | 12 |
| KQNSKLRP | Nd | No | - | 13 |
| GKQNSKLA | Nd | Yes | + | 14 |
| GKSNSKLK | Nd | Yes | + | 15 |
| GKTNSKLA | Nd | Yes | + | 16 |
| GKSASKQF | 267,21 | Yes | + | 17 |
| GAKQSGPA | Nd | No | - | 18 |
| GGKQSTAA | Nd | No | - | 19 |
| GGKLSKKK | Nd | Nd | - | 20 |
| GGKWSKLS | 267,21 | Yes | - | 21 |
| AKSFSKPR | No | No | - | 22 |
| GKSWSKGR | 267,2 | Yes | + | 23 |
| AKQNSKLR | Nd | No | - | 24 |
| GGKQSKAA | Nd | No | - | 25 |
| GGKSFSKP | No | No | - | 26 |
| GGKFSSKPR | 267,21/324,23 | Yes/Yes | +/+ | 27/28 |
| AKPTSKDSGLK | Nd | No | - | 29 |
| AGKFSKPR | No | No | - | 30 |
| AGKSFSKPR | No | No | - | 31 |
