## Supplementary Table 3 for "Proteome-wide probing of the dual NMT-dependent myristoylation tradeoff unveils potent, mechanism-based suicide inhibitors"

### Supplementary Table 3: Primers used in this study

Cloning and site-directed mutagenesis primers for IpaJ.

| Sequence | Use |
|---|---|
| TGGCCGCACCGCTAGAATTATCATACATTTGTTTGC | C64S-for |
| GCAAACAAATGTATGATAATTCTAGCGGTGCGGCCA | C64S-rev |
| AGATACAGATCGCGTTCACTACGGTTGTCATTATCATTAGAGCTTTTACGGGTCATTTCCGACATTG | L97N-L99N-C102S-for |
| CAATGTCGGAAATGACCCGTAAAAGCTCTAATGATAATGACAACCGTAGTGAACGCGATCTGTATCT | L97N-L99N-C102S-rev |
| AATCTTTATTTTCAGGGCGCCATGGCGCGTGAATATATGATTAGCCATCAG | IpaJ-Nt30-for |
| TCGAGTGCGGCCGCAAGCTTTCACAGTGCTTCGTTGGTAATAA | IpaJ-Ct259-rev |
| ATGTCCGAACAACGCAAACCGTGTAAACGTGGCTGTATTCATACGGGTGTGATGCTGTATGGTGTCCTGCTGCAAGGTGC<br>TATTCCGCGTGAATATATGATTAGCCATCAGACCGATGTCCGTGTGAACGAAAATCGCGTCAACGAACAGGGCTGCTTTC<br>TGGCGCGCAAACAAATGTATGATAATTCTTGCGGTGCGGCCAGTCTGCTGTGTGCAGCTAAAGAACTGGGCGTTGACAAA<br>ATTCCGCAATACAAAGGTTCAATGTCGGAAATGACCCGTAAAAGCTCTCTGGATCTGGACAACCGTTGTGAACGCGATCT<br>GTATCTGATCACGTCCGGCAACTACAATCCGCGTATTCACAAAGATAATATCGCGGACGCCGGTTACTCTATGCCGGACA<br>AAATTGTTATGGCCACCCGTCTGCTGGGTCTGAACGCCTATGTGGTTGAAGAATCAAACATCTTTTCGCAGGTTATCAGC<br>TTCATCTACCCGGATGCCCGTGACCTGCTGATTGGCATGGGTTGCAACATCGTCCATCAGCGTGATGTGCTGAGTTCCAA<br>TCAACGCGTGCTGGAAGCAGTTGCTGTCAGCTTTATCGGCGTGCCGGTTGGTCTGCACTGGGTTCTGTGCCGCCCGGATG<br>GCAGTTATATGGACCCGGCAGTCGGTGAAAACTACAGTTGTTTCTCCACGATGGAACTGGGTGCTCGTCGCAGCAATTCT<br>AACTTTATCGGCTACACCAAAATCGGCATCAGCATTGTTATTACCAACGAAGCACTGTGA | Full synthetic DNA sequence for *S. flexneri* IpaJ (1-259) |
