## Supplementary Table 4 for "Proteome-wide probing of the dual NMT-dependent myristoylation tradeoff unveils potent, mechanism-based suicide inhibitors"

**Supplementary Table 4: Data collection and refinement statistics of HsNMT1 in complex with peptide substrates and inhibitors**

| | HPCA | AK | AcK | AcG[ORN] | AN | me-GN | GGK |
|---|---|---|---|---|---|---|---|
| **PDB accession code** | 7OWM | 7OWN | 7OWO | 7OWP | 7OWU | 7OWQ | 7OWR |
| **Data collection** | | | | | | | |
| Space group | $P2_12_12$ | $P2_12_12$ | $P2_12_12$ | $P2_12_12$ | $P2_12_12$ | *C121* | *C121* |
| Cell dimensions | | | | | | | |
| *a*, *b*, *c* (Å) | 79.88, 178.4, 58.3 | 79.1, 178.1, 58.2 | 79.32, 178.3, 58.2 | 79.32, 178.2, 58.2 | 78.74 178.0 58.3 | 93.65, 58.88, 154.21 | 90.4, 61.4, 158.9 |
| α, β, γ (°) | 90.0, 90.0, 90.0 | 90.0, 90.0, 90.0 | 90.0, 90.0, 90.0 | 90.0, 90.0, 90.0 | 90.0, 90.0, 90.0 | 90.0, 90.0, 90.0 | 90.0, 90.0, 90.0 |
| Resolution (Å) | 48.8–1.5 (1.54-1.50)* | 48.90–2.01 (2.06-2.01) | 48.7–1.70 (1.74-1.70) | 48.7–1.81 (1.85-1.81) | 46.90–2.04 (2.08-2.04) | 47.69–3.0 (3.18-3.00) | 48.4–2.39 (2.44-2.39) |
| $R_{sym}$ or $R_{merge}$ | 0.115 (1.532) | 0.171 (1.486) | 0.11 (1.401) | 0.100 (0.562) | 0.060 (0.317) | 0.134 (0.601) | 0.090 (0.481) |
| $I / \sigma I$ | 12.2 (1.4) | 15.9 (2.0) | 14.4 (1.9) | 16.9 (4.8) | 13.0 (4.3) | 11.2 (3.0) | 12.7 (3.12) |
| Completeness S(%) | 96.1 (77.5) | 99.4 (92.1) | 86.6 (48.7) | 99.1 (98.2) | 99.6 (99.9) | 98.6 (98.7) | 99.4 (91.3) |
| Completeness E(%) | 98.1 (97.8) | | 97.2 (96.1) | | | | 99.4 (91.3) |
| Redundancy | 13.9 (8.9) | 14.9 (15.7) | 13.2 (12.3) | 15.0 (14.7) | 3.4 (3.3) | 6.3 (5.9) | 5.71 (4.9) |
| **Refinement** | | | | | | | |
| Resolution (Å) | 48.8–1.50 | 48.8–2.10 | 48.7–1.70 | 47.6–1.81 | 46.86–2.08 | 47.7–3.00 | 48.4–2.39 |
| No. reflections | 34542 | 49330 | 79532 | 75344 | 49769 | 32321 | 34536 |
| $R_{work}$ / $R_{free}$ | 15.44/19.58 | 16.63/21.50 | 18.29/22.32 | 15.52/19.10 | 17.86/22.36 | 21.05/25.51 | 17.34/21.87 |
| No. atoms | | | | | | | |
| Protein | 6433 | 6411 | 6489 | 6538 | 6517 | 6330 | 6387 |
| Ligand/ion | 219 | 150 | 207 | 144 | 156 | 126 | 144 |
| Water | 1006 | 537 | 676 | 790 | 525 | | 366 |
| *B*-factors | | | | | | | |
| Protein | 23.98 | 31.88 | 25.65 | 31.01 | 29.75 | 45/.17 | 30.48 |
| Ligand/ion | 25.97 | 36.67 | 29.69 | 40.45 | 36.23 | 53.26 | 26.83 |
| Water | 32.68 | 35.25 | 31.85 | 37.61 | 35.30 | | 30.37 |
| R.m.s. deviations | | | | | | | |
| Bond lengths (Å) | 0.006 | 0.007 | 0.007 | 0.006 | 0.004 | 0.003 | 0.003 |
| Bond angles (°) | 0.874 | 0.913 | 0.899 | 0.862 | 0.793 | 0.700 | 0.823 |

The datasets were collected from a single crystal for each hsNMT1 complex.
*Highest-resolution shell is shown in parentheses.

**Supplementary Table 4 (following): Data collection and refinement statistics of HsNMT1 in complex with peptide substrates and inhibitors**

| | Ac[D-ORN] | DAB |
|---|---|---|
| **PDB accession code** | 7OWS | 7OWT |
| **Data collection** | | |
| Space group | $P2_12_12$ | $P2_12_12$ |
| Cell dimensions | | |
| *a*, *b*, *c* (Å) | 79.8, 179.1, 58.4 | 78.74 178.0 58.3 |
| α, β, γ (°) | 90.0, 90.0, 90.0 | 90.0, 90.0, 90.0 |
| Resolution (Å) | 48.9–1.90 (1.94-1.90)* | 48.8–1.90 (1.94-1.90) |
| $R_{sym}$ or $R_{merge}$ | 0.080 (0.678) | 0.057 (0.584) |
| $I / \sigma I$ | 21.6 (5.0) | 25 (4.5) |
| Completeness S(%) | 99.9 (99.7) | 100 (99.9) |
| Completeness E(%) | | |
| Redundancy | 14.9 (15.7) | 13.0 (13.1) |
| **Refinement** | | |
| Resolution (Å) | 48.9–1.90 | 48.8–1.90 |
| No. reflections | 66831 | 65500 |
| $R_{work}$ / $R_{free}$ | 15.78/19.48 | 17.19/21.50 |
| No. atoms | | |
| Protein | 6553 | 6484 |
| Ligand/ion | 179/1 | 158/2 |
| Water | 616 | 487 |
| *B*-factors | | |
| Protein | 30.52 | 37.70 |
| Ligand/ion | 29.64/43.62 | 41.73/49.53 |
| Water | 37.57 | 40.78 |
| R.m.s. deviations | | |
| Bond lengths (Å) | 0.007 | 0.007 |
| Bond angles (°) | 0.940 | 0.943 |

The datasets were collected from a single crystal for each hsNMT1 complex.
*Highest-resolution shell is shown in parentheses.
