## Supplementary Dataset 1a for "Proteome-wide probing of the dual NMT-dependent myristoylation tradeoff unveils potent, mechanism-based suicide inhibitors"

### Sup Dataset 1a

| Series[a] | Peptide sequence | Short name (text & Figures) | MS data in Supplementary Dataset 1a |
|---|---|---|---|
| Reference GK-peptide variants | GKSFSKPR | GK | 2 |
| | G{Orn}SFSKPR | | 3 |
| | G{Dab}SFSKPR | | 4 |
| | ANCFSKPR | AN | 5 |
| | AGKFSKPR | | 6 |
| | me-GNCFSKPR | meG | 7 |
| | GKSWSKGR | | 8 |
| | GKFSSKPR[c] | GKFS | 9/10 |
| Human GKX | GKVLSKIF | ARF6 | 11 |
| | GKQNSKLR | HPCA | 12 |
| | GKTNSKLA | HPCL4 | 13 |
| | GKQNSKLA | VIS | 14 |
| | GKSNSKLK | NCS1 | 15 |
| | GKSLSHLP | T106B | 16 |
| | GKTFSQLG | T106A | 17 |
| | GKSASKQF | GAPP | 18 |
| | GKLHSKPA | NKD1 | 19 |
| | GKLQSKHA | NKD2 | 20 |
| GXK | GAKQSGPA | ZNRF2 | 21 |
| | GGKQSTAA | ZNRF1 | 22 |
| | GGKFSSKPR | | 23/24 |
| | GGKLSKKK | BASP | 25 |
| SOS3 | GCSVSKKK | SOS3 | 26 |
| | ASSVSKKK | | 27 |

### GKSFSKPR

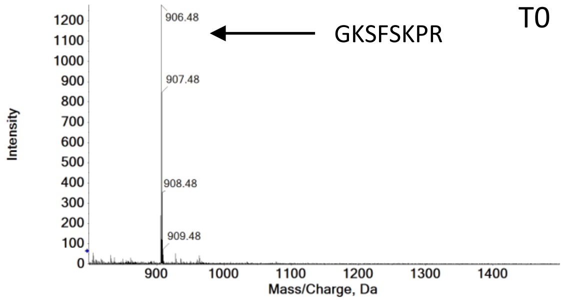
T0
GKSFSKPR
906.48
907.48
908.48
909.48
Intensity
Mass/Charge, Da

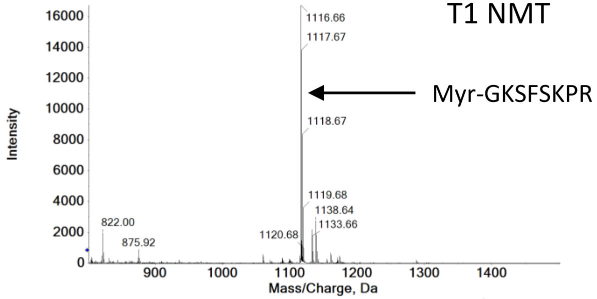
T1 NMT
Myr-GKSFSKPR
1116.66
1117.67
1118.67
1119.68
1138.64
1133.66
1120.68
822.00
875.92
Intensity
Mass/Charge, Da

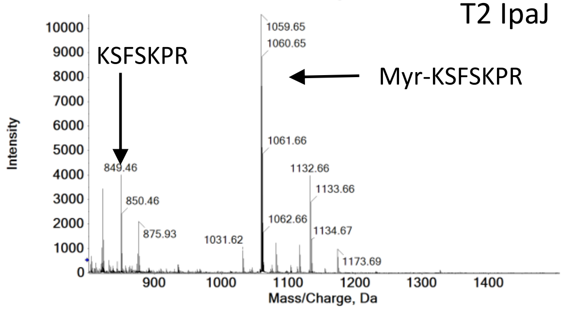
T2 IpaJ
KSFSKPR
Myr-KSFSKPR
1059.65
1060.65
1061.66
1062.66
1132.66
1133.66
1134.67
1173.69
1031.62
849.46
850.46
875.93
Intensity
Mass/Charge, Da

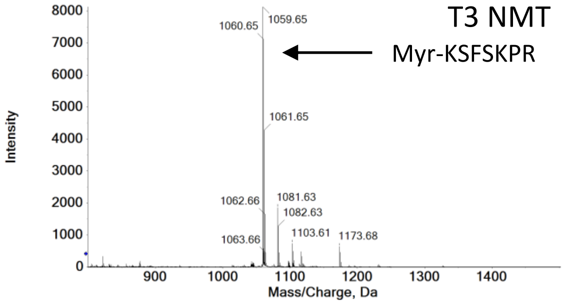
T3 NMT
Myr-KSFSKPR
1060.65
1059.65
1061.65
1062.66
1063.66
1081.63
1082.63
1103.61
1173.68
Intensity
Mass/Charge, Da

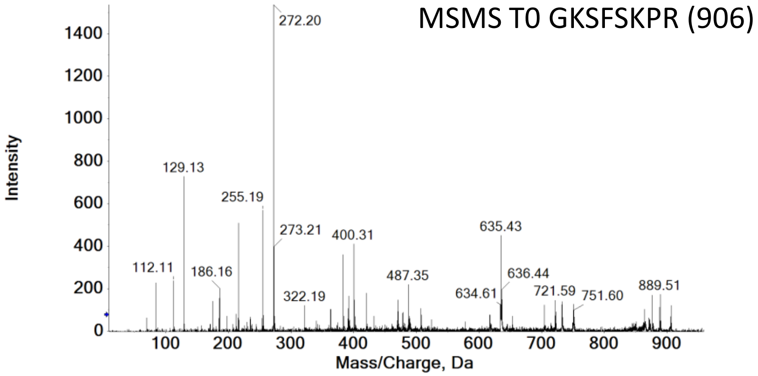
MSMS T0 GKSFSKPR (906)
272.20
129.13
255.19
273.21
400.31
635.43
112.11
186.16
322.19
487.35
636.44
634.61
721.59
751.60
889.51
Intensity
Mass/Charge, Da

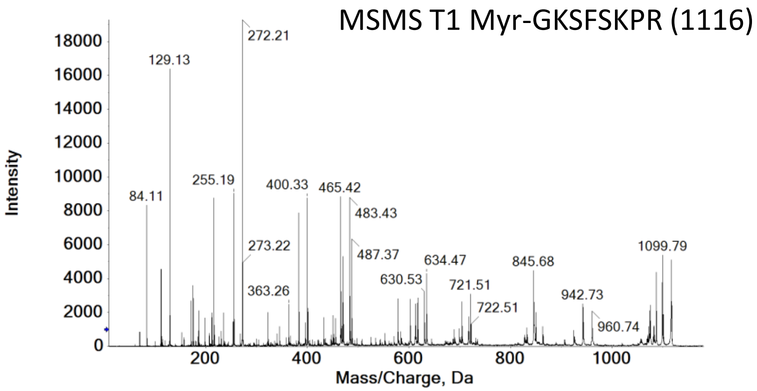
MSMS T1 Myr-GKSFSKPR (1116)
272.21
129.13
255.19
400.33
465.42
483.43
84.11
273.22
487.37
634.47
845.68
1099.79
630.53
721.51
722.51
363.26
942.73
960.74
Intensity
Mass/Charge, Da

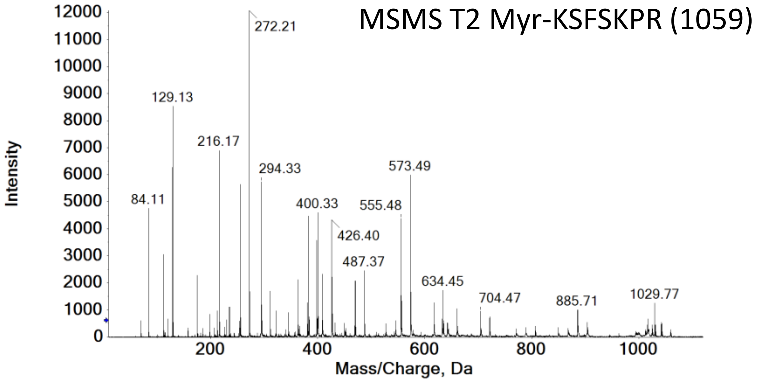
MSMS T2 Myr-KSFSKPR (1059)
272.21
129.13
216.17
294.33
573.49
84.11
400.33
555.48
426.40
487.37
634.45
704.47
885.71
1029.77
Intensity
Mass/Charge, Da

### G{ORN}SFSKPR

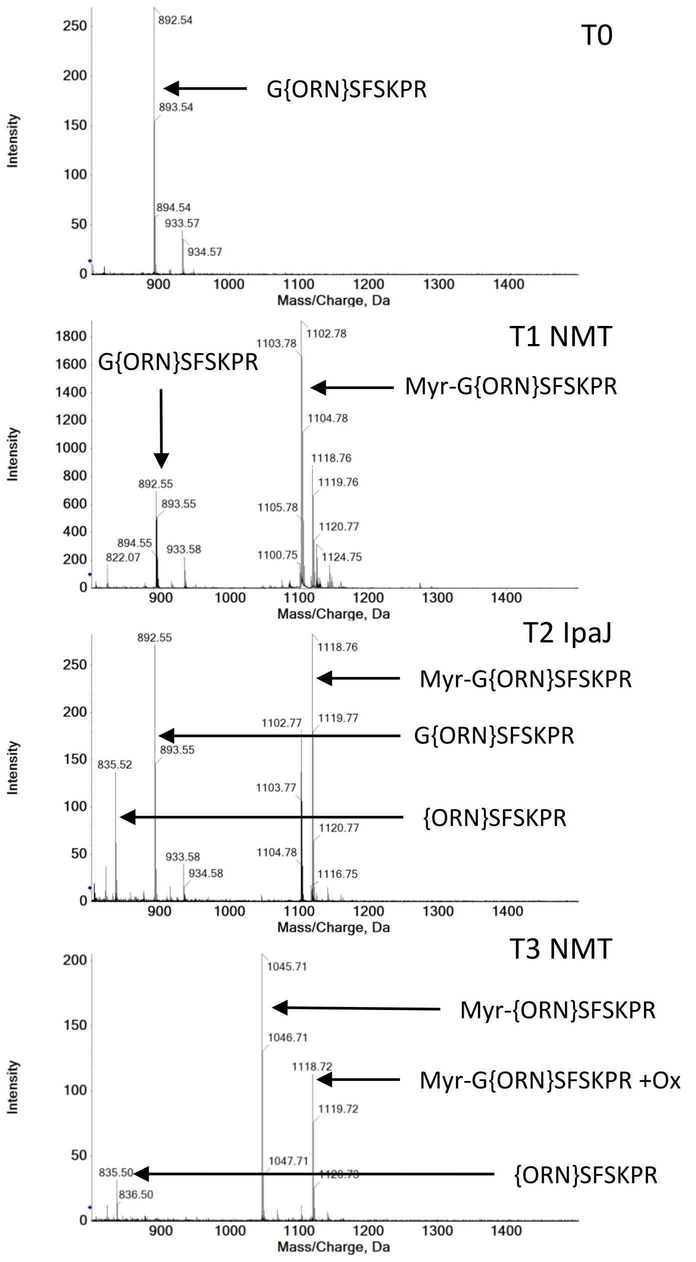


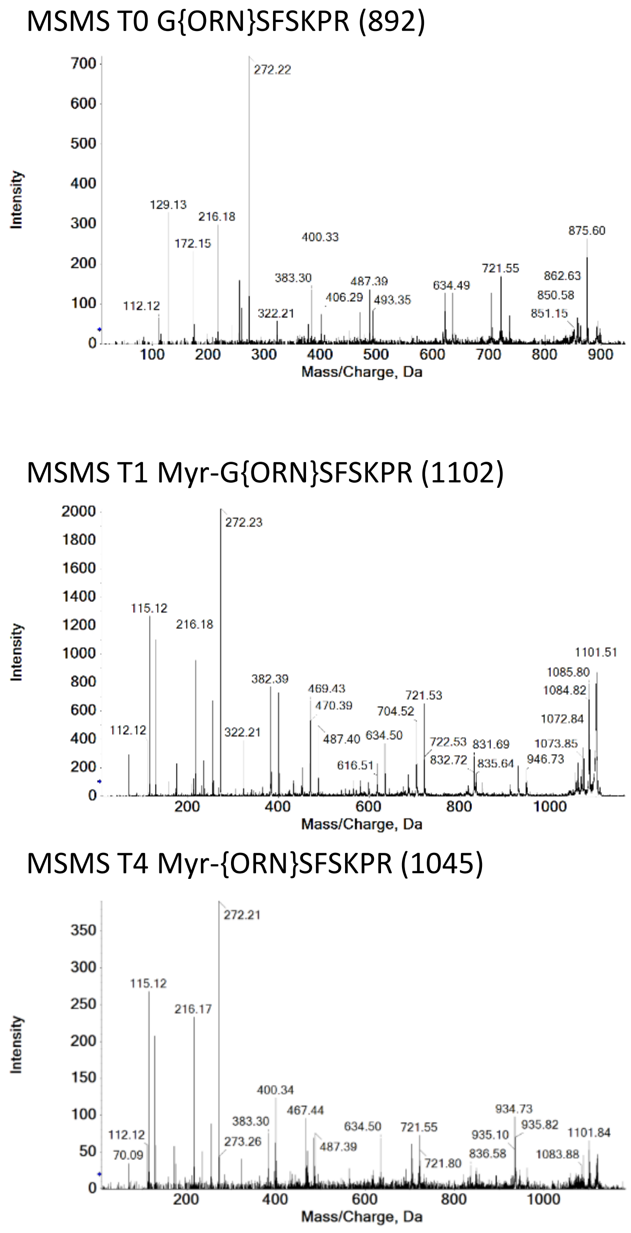


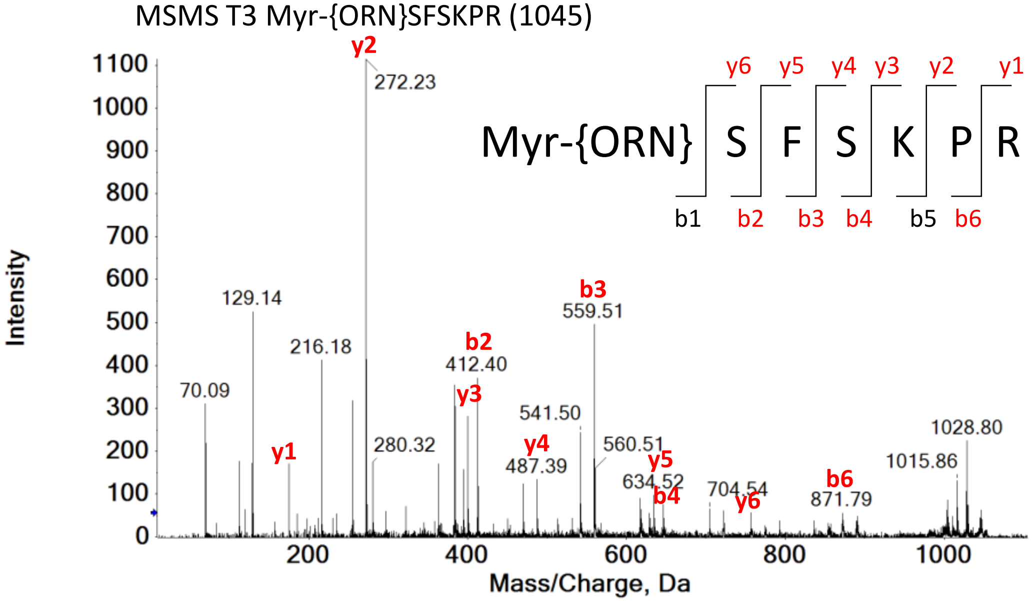


| | Mass/Charge (Da) | Intensity | ion |
|---|---|---|---|
| 875 | 175,15 | 170,50 | y1 |
| 1983 | 272,22 | 1113,88 | y2 |
| 4152 | 400,33 | 282,50 | y3 |
| 4407 | 412,37 | 371,60 | b2 |
| 5781 | 487,39 | 134,58 | y4 |
| 6965 | 559,47 | 496 | b3 |
| 8430 | 634,48 | 97,41 | y5 |
| 8686 | 646,51 | 76,39 | b4 |
| 9878 | 721,51 | 61,96 | y6 |
| 11989 | 871,78 | 56,62 | b6 |

### G{Dab}SFSKPR

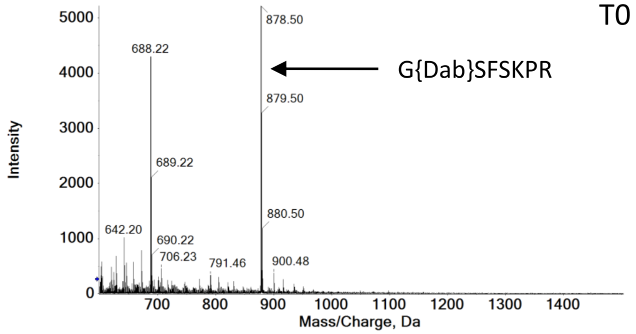
T0
G{Dab}SFSKPR
Intensity
Mass/Charge, Da
878.50
688.22
879.50
689.22
880.50
642.20
690.22
706.23
791.46
900.48

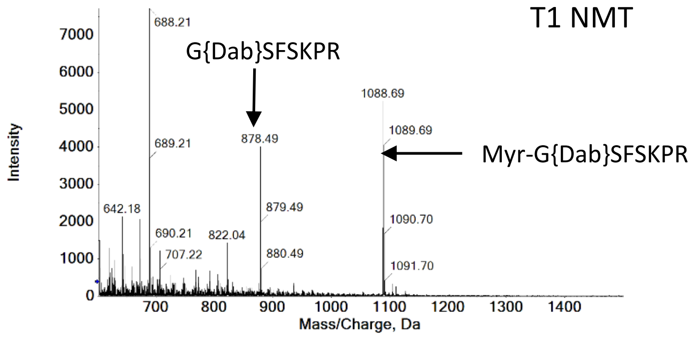
T1 NMT
G{Dab}SFSKPR
Myr-G{Dab}SFSKPR
Intensity
Mass/Charge, Da
688.21
689.21
878.49
1088.69
1089.69
879.49
642.18
690.21
822.04
1090.70
707.22
880.49
1091.70

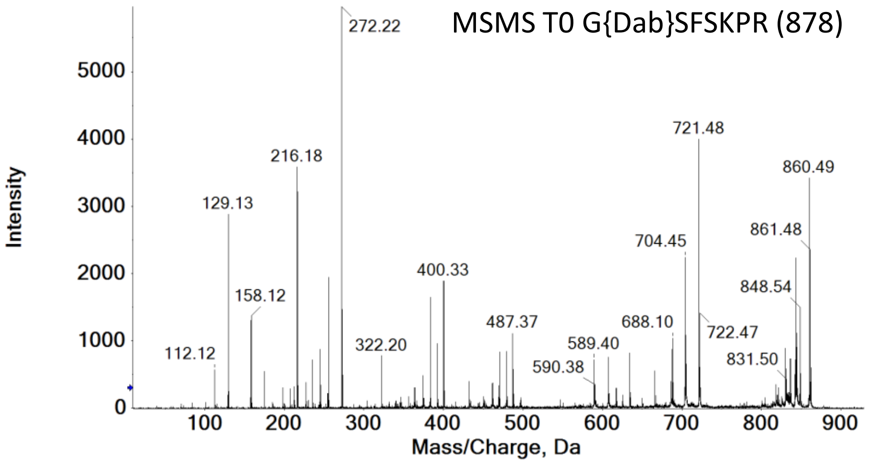
MSMS T0 G{Dab}SFSKPR (878)
Intensity
Mass/Charge, Da
272.22
721.48
216.18
860.49
129.13
861.48
704.45
400.33
848.54
158.12
487.37
688.10
722.47
112.12
322.20
589.40
831.50
590.38

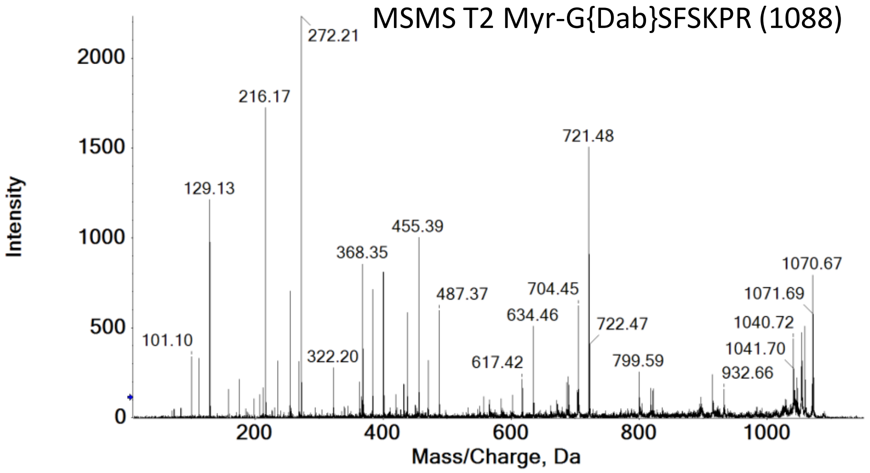
MSMS T2 Myr-G{Dab}SFSKPR (1088)
Intensity
Mass/Charge, Da
272.21
216.17
721.48
129.13
455.39
368.35
1070.67
487.37
704.45
1071.69
634.46
722.47
1040.72
101.10
1041.70
322.20
617.42
799.59
932.66

### ANCFSKPR

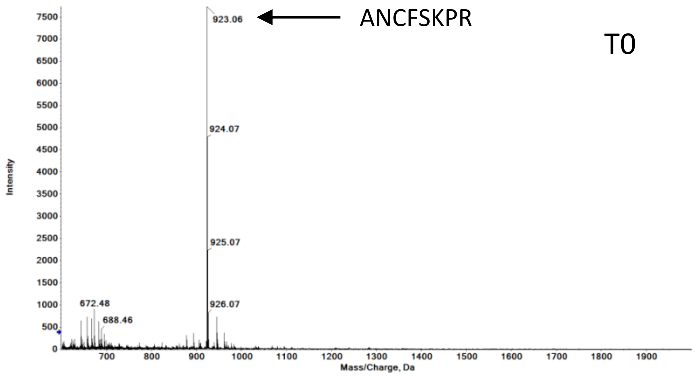
923.06
924.07
925.07
926.07
672.48
688.46
ANCFSKPR
T0
Intensity
Mass/Charge, Da

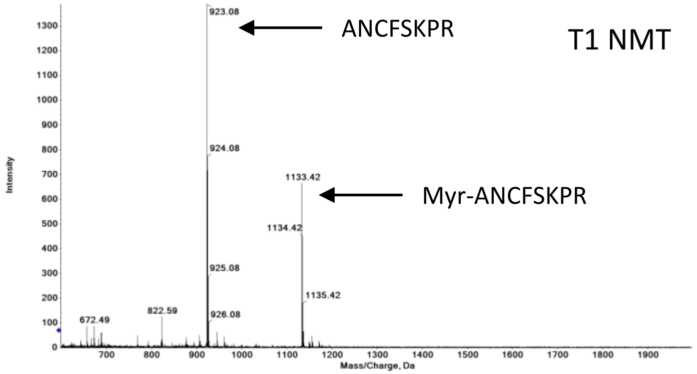
923.08
924.08
925.08
926.08
1133.42
1134.42
1135.42
672.49
822.59
ANCFSKPR
Myr-ANCFSKPR
T1 NMT
Intensity
Mass/Charge, Da

### AGKFSKPR

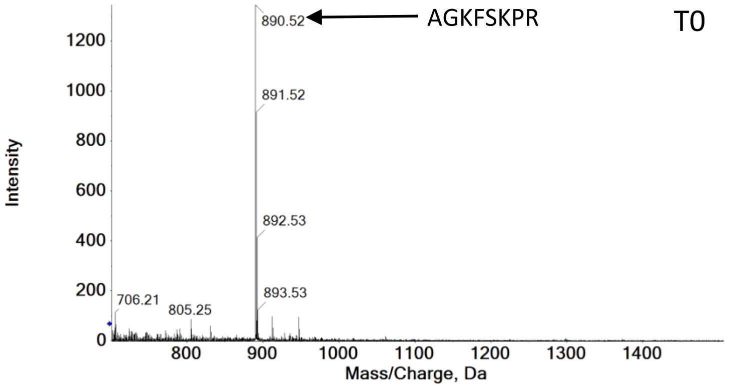


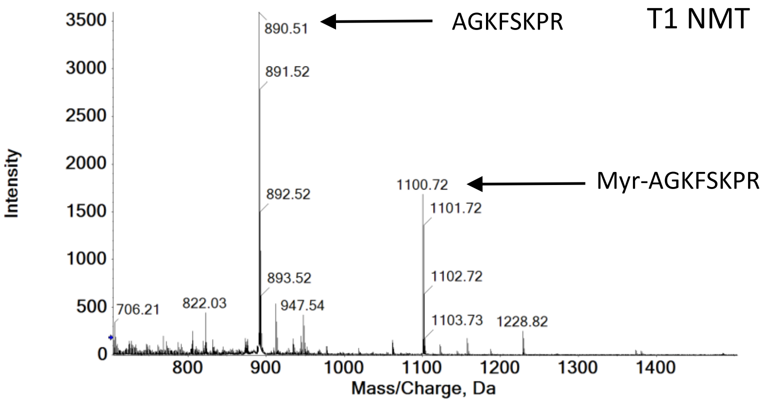


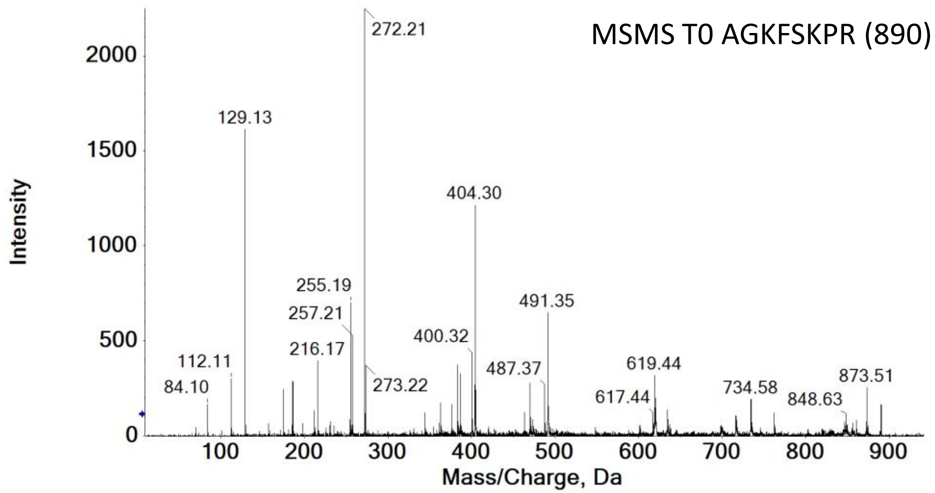


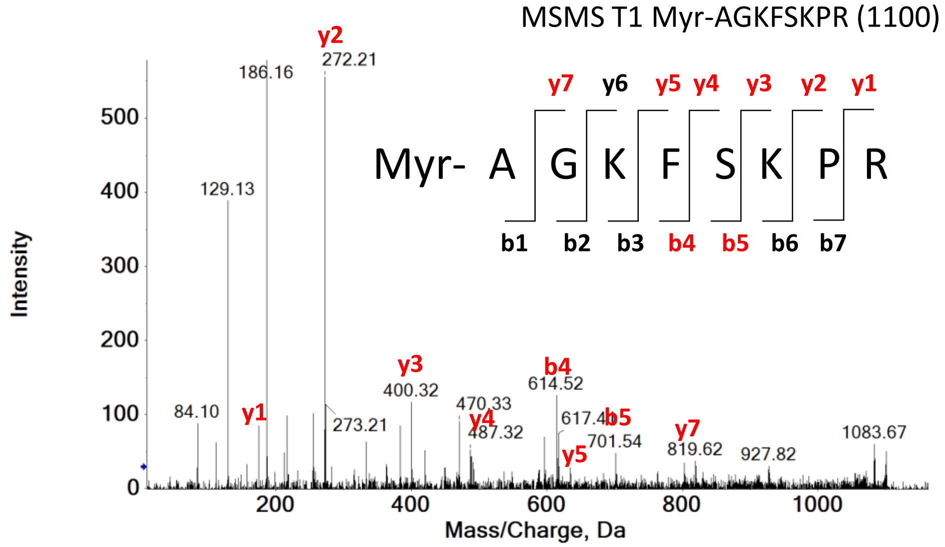


| L4 1100 | Mass/Charge (Da) | Intensity | |
|---|---|---|---|
| 542 | 129,138 | 388,8628 | K |
| 801 | 175,1467 | 84,86275 | y1 |
| 894 | 186,1513 | 578,353 | GK |
| 1149 | 216,181 | 98,66667 | SK |
| 1590 | 272,2057 | 555,9216 | y2 |
| 2024 | 333,2545 | 63,21569 | GKF |
| 2715 | 400,3234 | 116,8627 | y3 |
| 2896 | 420,3111 | 51,45098 | GKFS |
| 3333 | 470,3304 | 90,82353 | y4-NH3 |
| 3503 | 487,3036 | 52,07843 | y4 |
| 4591 | 614,5147 | 126,1177 | b4 |
| 4836 | 634,4205 | 28,07843 | y5 |
| 5421 | 701,5098 | 47,68628 | b5 |
| 6483 | 819,5577 | 37,17647 | y7 |

### me-GNCFSKPR

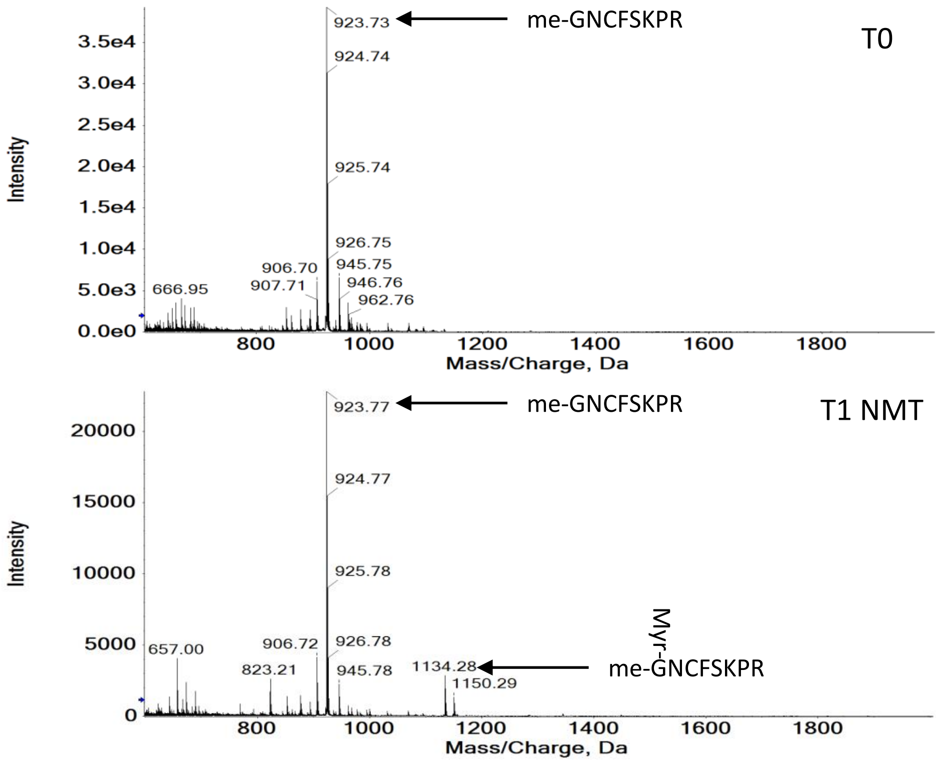


MSMS T1 me-(Myr)GNCFSKPR (1134)

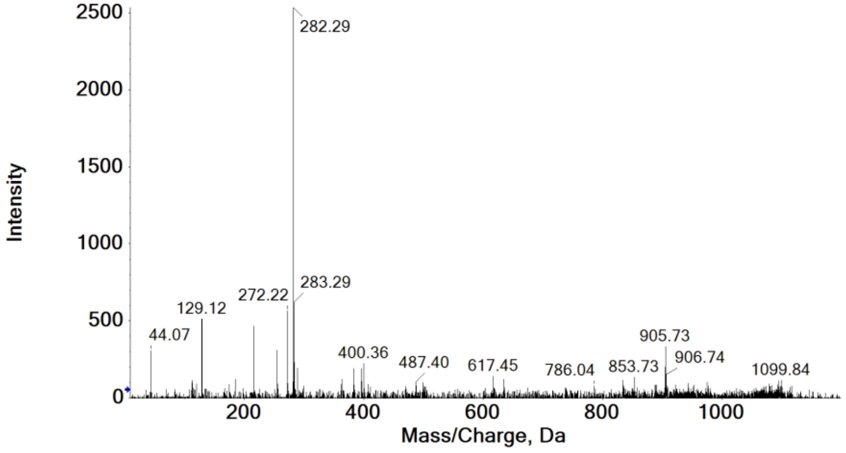

### GKSWSKGR

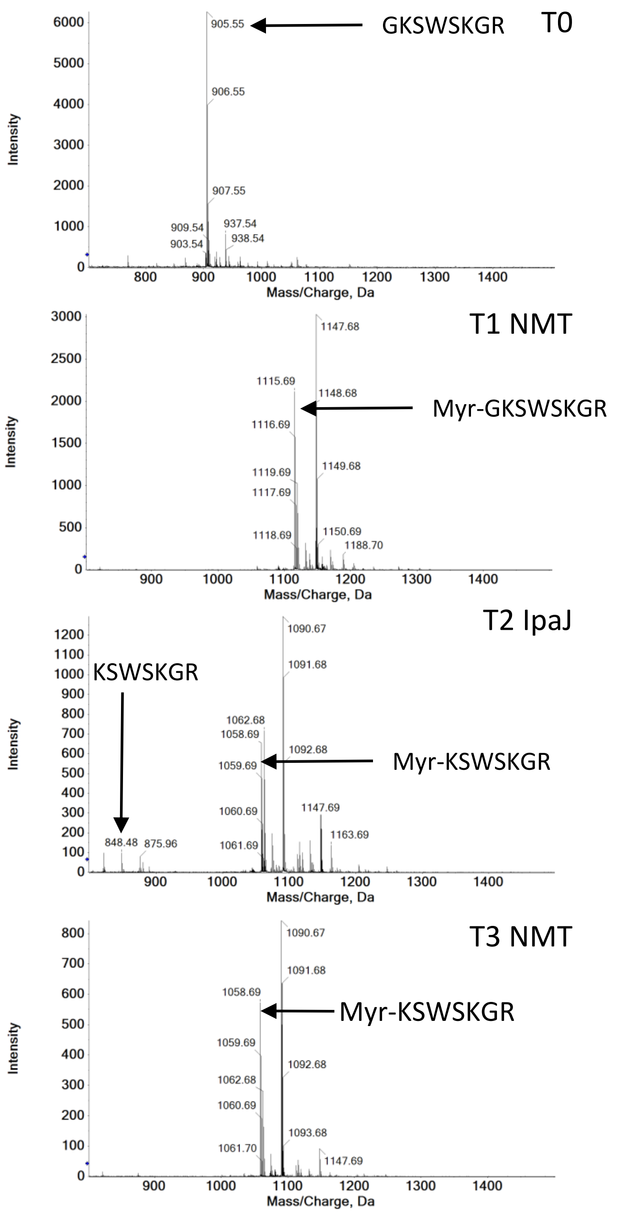


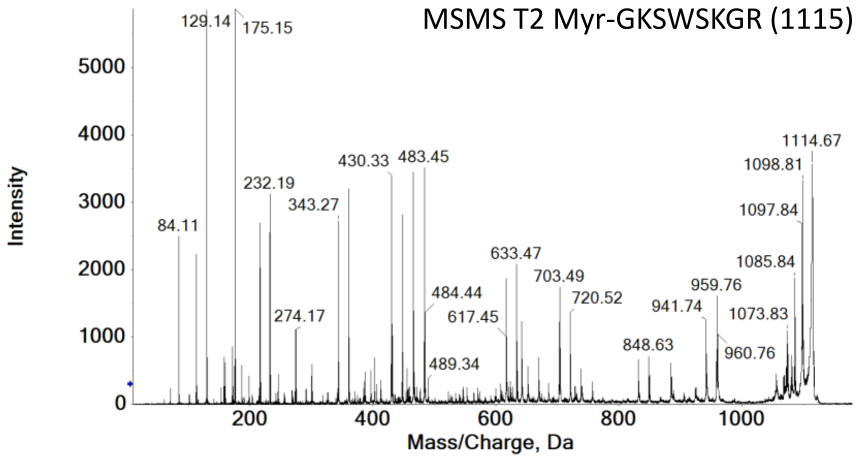


| | Mass/Charge (Da) | Intensity | |
|---|---|---|---|
| 1520 | 175,1429 | 1428,392 | y1 |
| 2460 | 232,1888 | 496,1569 | y2 |
| 5185 | 360,3024 | 479,3726 | y3 |
| 7074 | 426,405 | 552,1569 | b2 |
| 7655 | 447,3432 | 485,4902 | y4 |
| 11575 | 612,537 | 578,0392 | b3 |
| 12102 | 633,4509 | 383,6863 | y5 |
| 13648 | 699,5988 | 61,80392 | b4 |
| 14137 | 720,4839 | 172,0784 | y6 |
| 17256 | 884,7047 | 301,9608 | b6 |

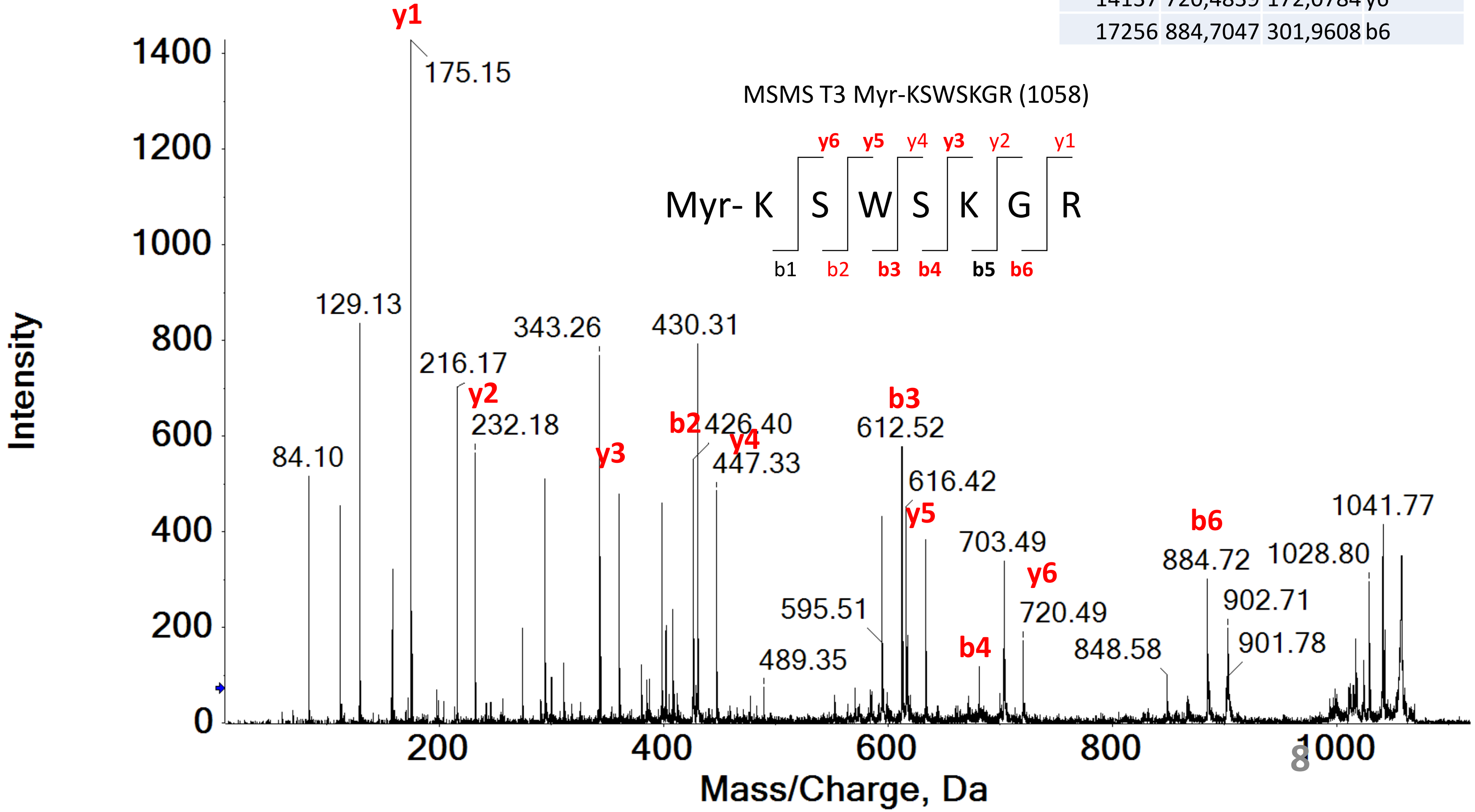

### GKFSSKPR

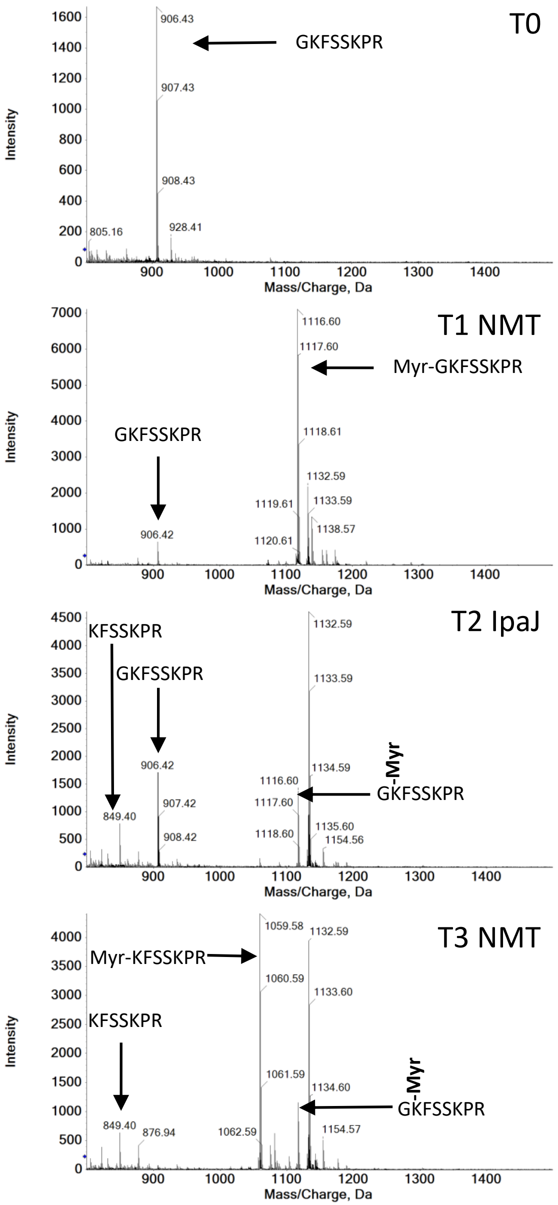
T0
GKFSSKPR
906.43
907.43
908.43
928.41
805.16
T1 NMT
Myr-GKFSSKPR
GKFSSKPR
1116.60
1117.60
1118.61
1119.61
1120.61
1132.59
1133.59
1138.57
906.42
T2 IpaJ
KFSSKPR
GKFSSKPR
Myr-
GKFSSKPR
1132.59
1133.59
1134.59
1116.60
1117.60
1118.60
1135.60
1154.56
906.42
907.42
908.42
849.40
T3 NMT
Myr-KFSSKPR
KFSSKPR
Myr-
GKFSSKPR
1059.58
1060.59
1061.59
1062.59
1132.59
1133.60
1134.60
1154.57
849.40
876.94
Intensity
Mass/Charge, Da

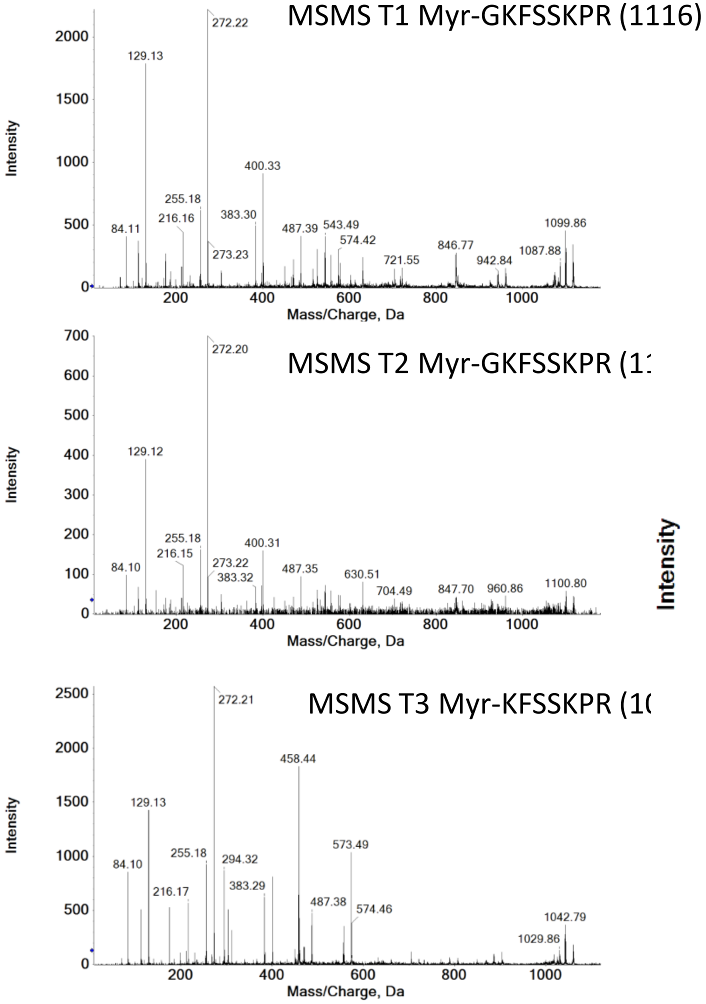
MSMS T1 Myr-GKFSSKPR (1116)
272.22
129.13
400.33
255.18
216.16
383.30
84.11
487.39
543.49
574.42
273.23
846.77
721.55
942.84
1087.88
1099.86
MSMS T2 Myr-GKFSSKPR (1:
272.20
129.12
255.18
216.15
400.31
273.22
383.32
84.10
487.35
630.51
704.49
847.70
960.86
1100.80
MSMS T3 Myr-KFSSKPR (1(
272.21
458.44
129.13
573.49
255.18
294.32
84.10
383.29
216.17
487.38
574.46
1042.79
1029.86
Intensity
Mass/Charge, Da

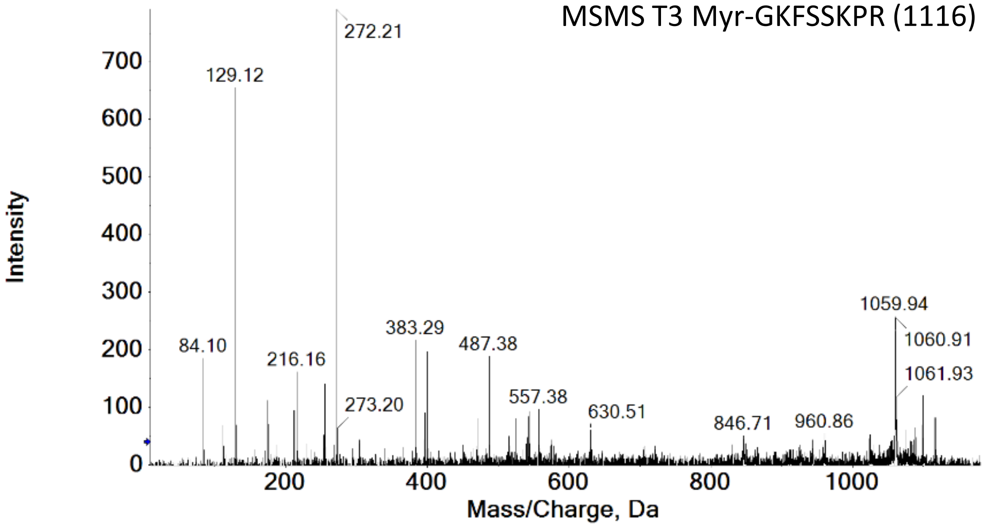
MSMS T3 Myr-GKFSSKPR (1116)
272.21
129.12
84.10
216.16
273.20
383.29
487.38
557.38
630.51
846.71
960.86
1059.94
1060.91
1061.93
Intensity
Mass/Charge, Da

MSMS T1 Myr-GKFSSKPR (1116)

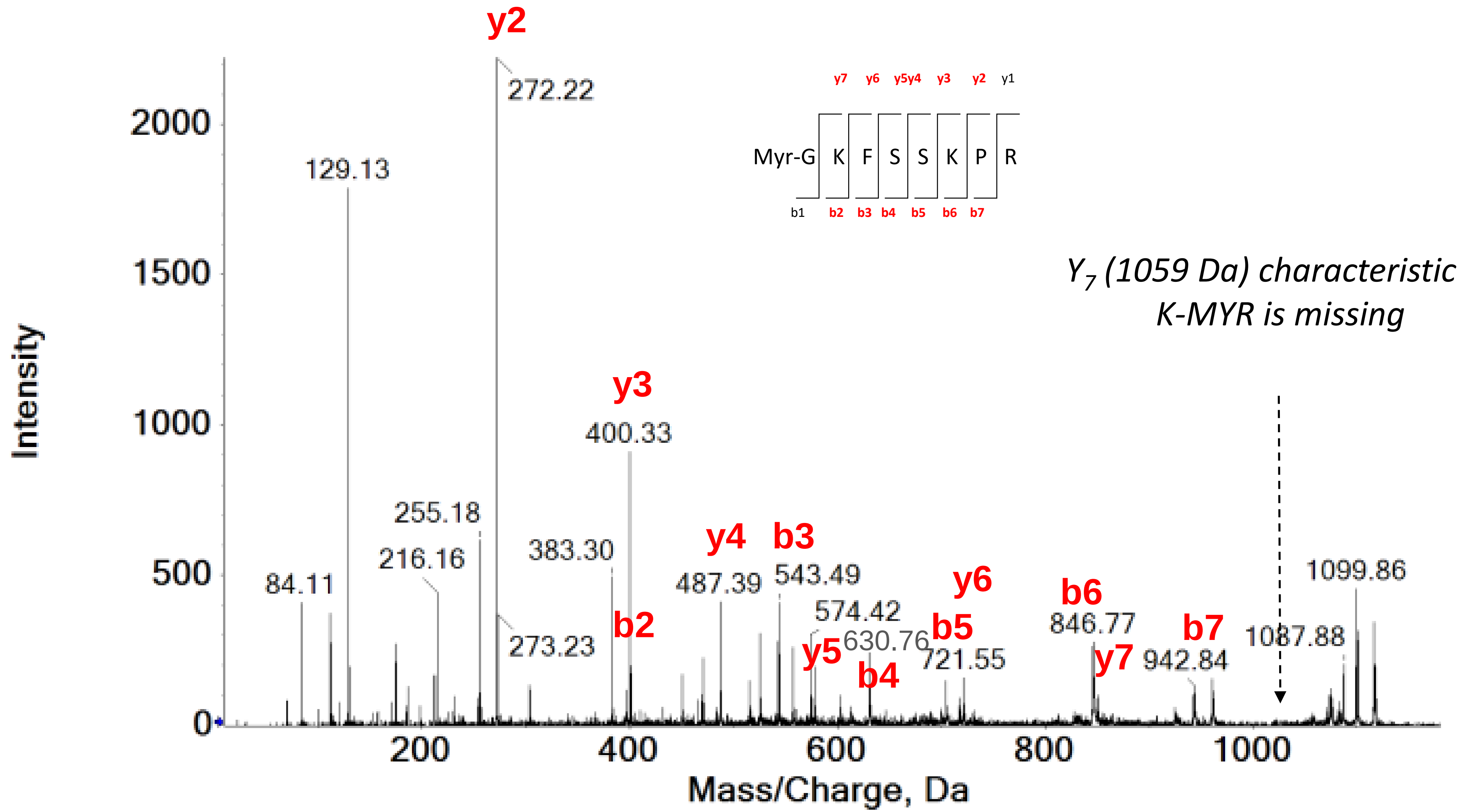


*$Y_7$ (1059 Da) characteristic of K-MYR is missing*

| L9 1116.6 | Mass/Charge (Da) | Intensity | |
|---|---|---|---|
| 1118 | 175,1541 | 270,5883 | y1 |
| 2531 | 272,2059 | 2221,647 | y2 |
| 4646 | 396,3899 | 116,549 | b2 |
| 4763 | 400,3088 | 912,1569 | y3 |
| 6619 | 487,3806 | 411,9216 | y4 |
| 7923 | 543,4726 | 409,5686 | b3 |
| 8670 | 574,7245 | 101,1765 | y5 |
| 9901 | 630,7576 | 151,5294 | b4 |
| 11807 | 717,5554 | 91,76471 | b5 |
| 11897 | 721,5382 | 156,7059 | y6 |
| 14106 | 845,9461 | 136 | b6 |
| 14189 | 849,7706 | 87,52941 | y7 |
| 15804 | 942,8001 | 135,0588 | b7 |

#### **GKVLSKIF** NMT1

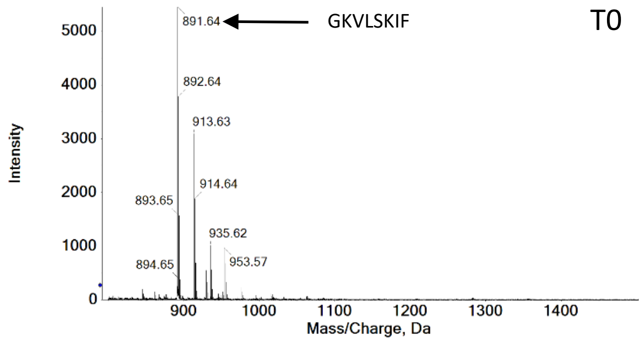


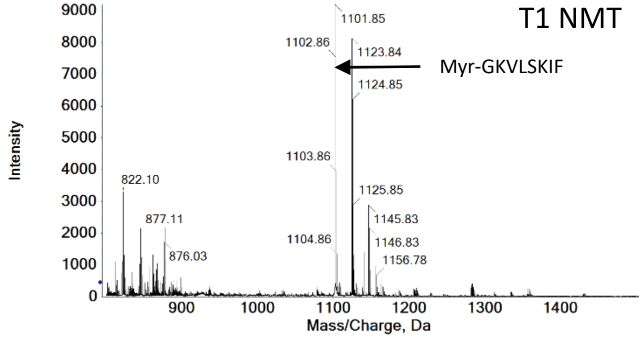


MSMS T0 GKFVLSKIF (891)

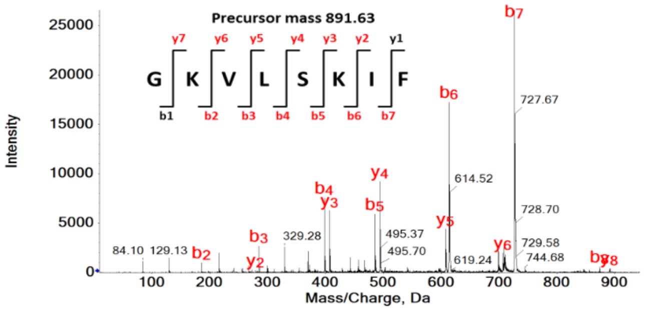


MSMS T1 Myr-GKFVLSKIF (1101)

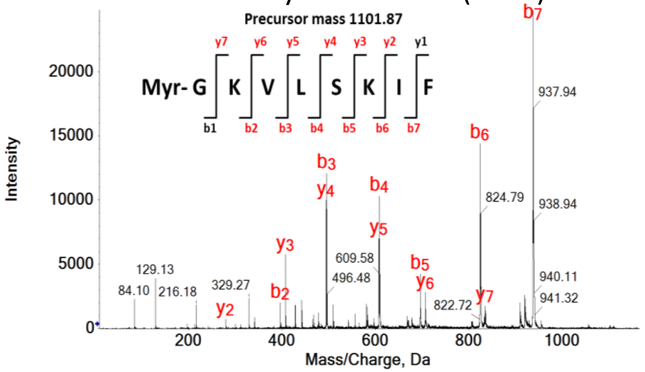


#### GKVLSKIF NMT2

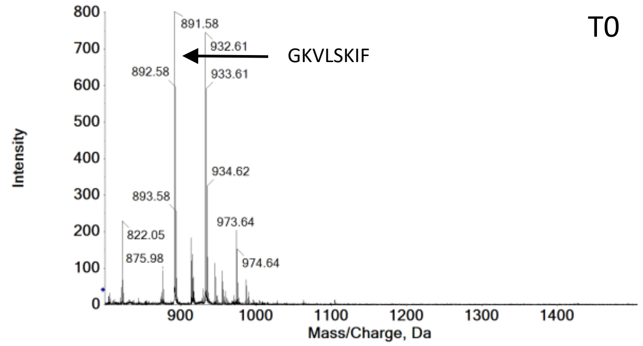


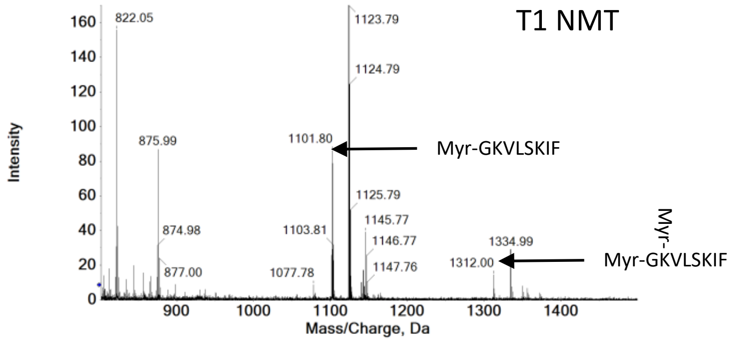

### GKQNSKLR

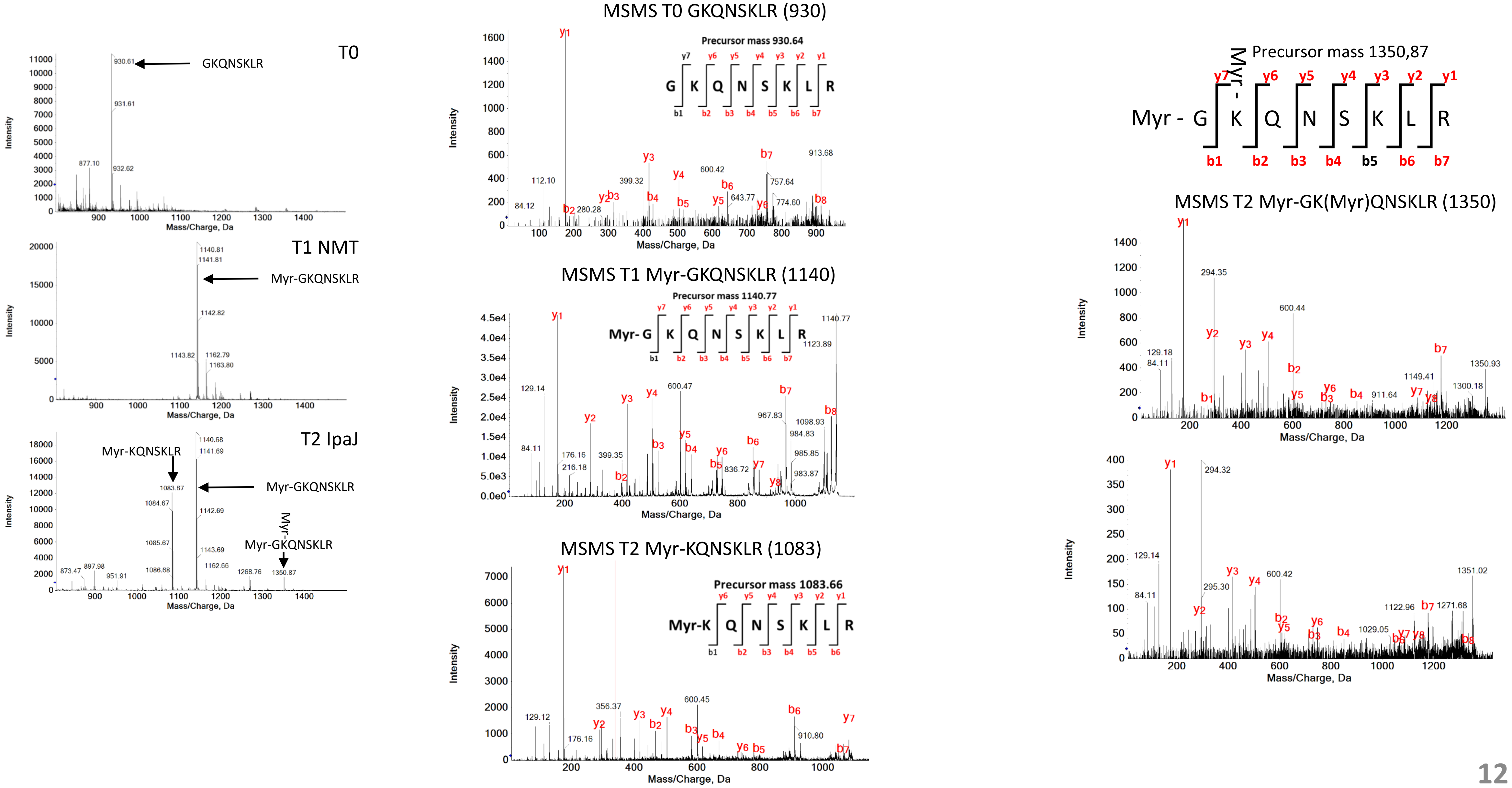
T0
GKQNSKLR
T1 NMT
Myr-GKQNSKLR
T2 IpaJ
Myr-KQNSKLR
Myr-GKQNSKLR
Myr-GK(Myr)QNSKLR
MSMS T0 GKQNSKLR (930)
Precursor mass 930.64
MSMS T1 Myr-GKQNSKLR (1140)
Precursor mass 1140.77
MSMS T2 Myr-KQNSKLR (1083)
Precursor mass 1083.66
Precursor mass 1350,87
MSMS T2 Myr-GK(Myr)QNSKLR (1350)
Intensity
Mass/Charge, Da

### GKTNSKLA

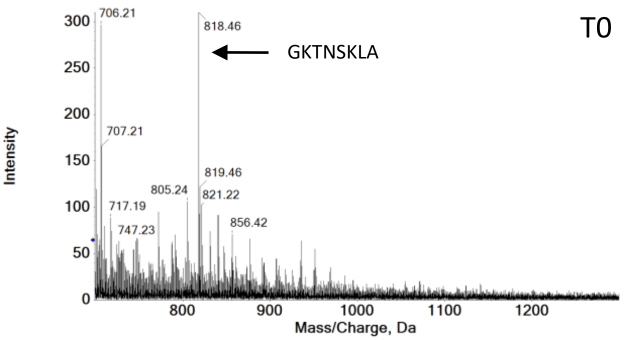
T0
GKTNSKLA
706.21
818.46
707.21
819.46
805.24
821.22
717.19
747.23
856.42
Intensity
Mass/Charge, Da

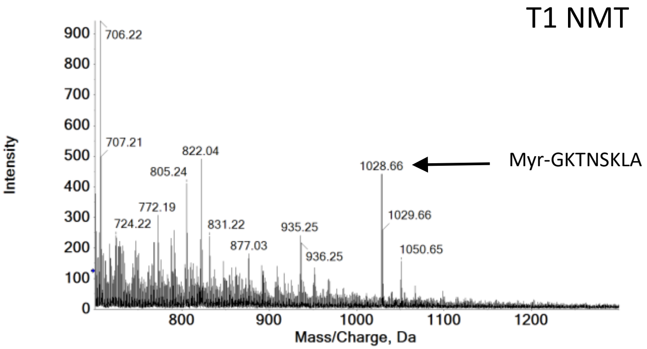
T1 NMT
Myr-GKTNSKLA
706.22
707.21
822.04
805.24
1028.66
772.19
724.22
831.22
935.25
1029.66
877.03
936.25
1050.65
Intensity
Mass/Charge, Da

### GKQNSKLA

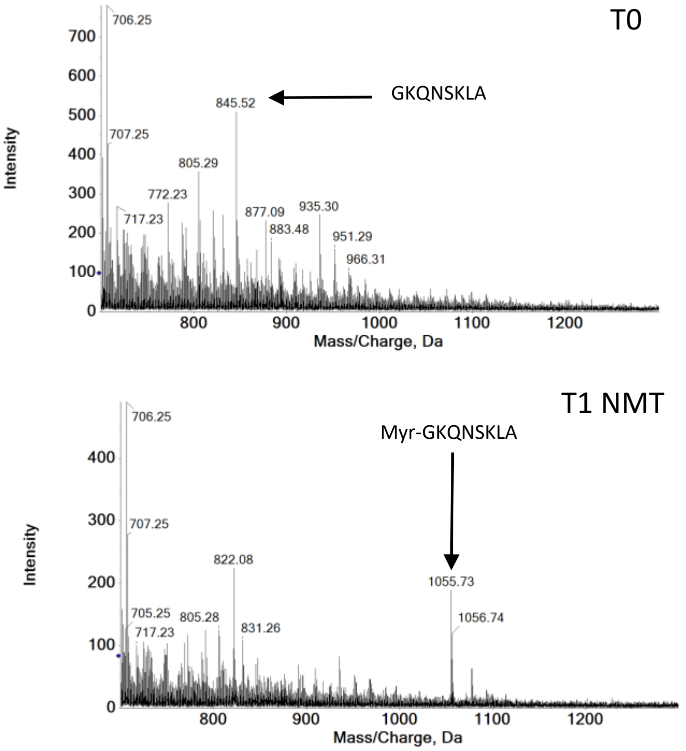


MSMS T1 Myr-GKQNSKLA (1055)

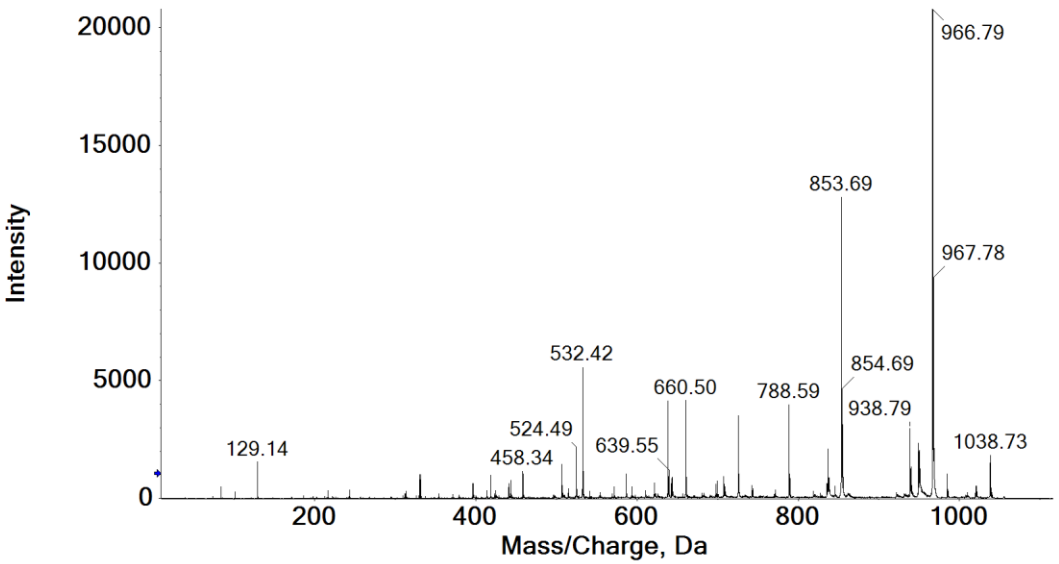

### GKSNSKLK

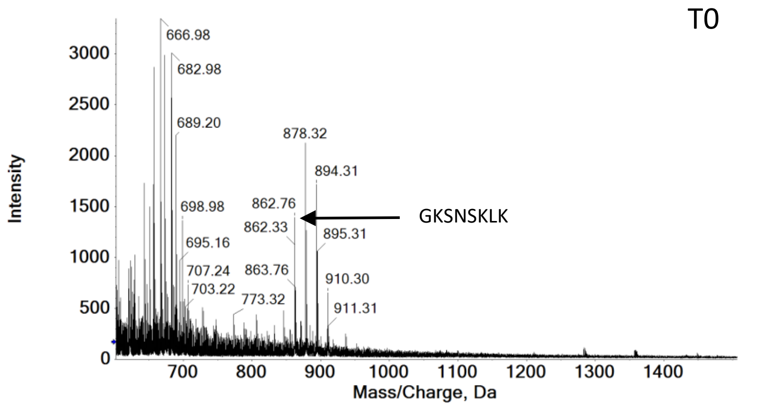


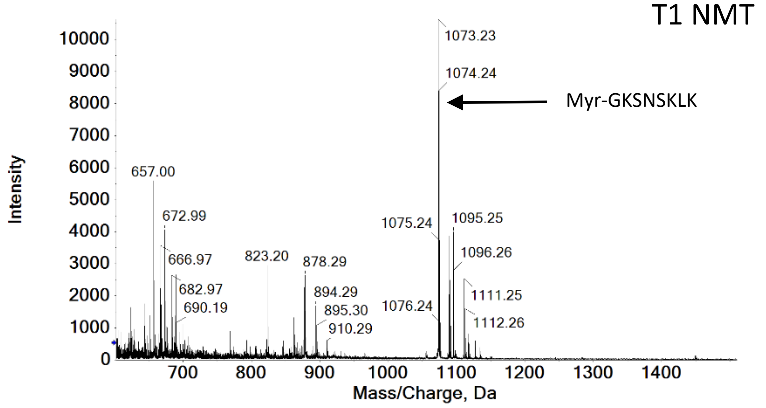


MSMS T1 Myr-GKSNSKLK (1073) (1)

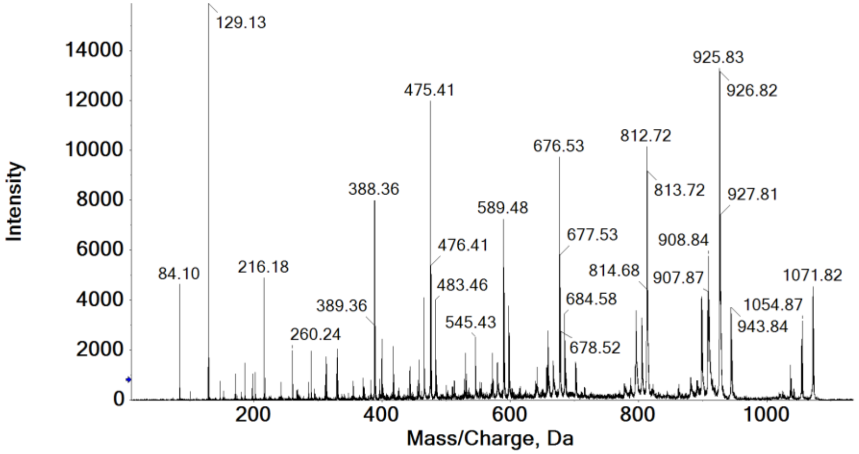


MSMS T1 Myr-GKSNSKLK (1073) (2)

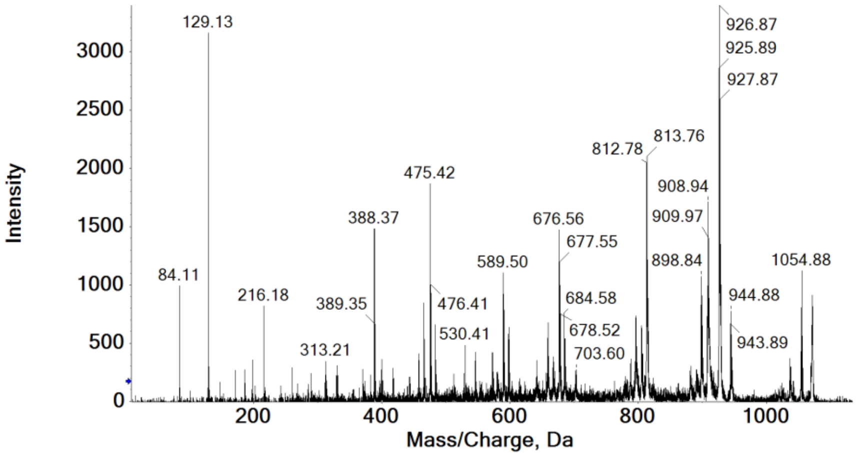

### GKSLSHLP

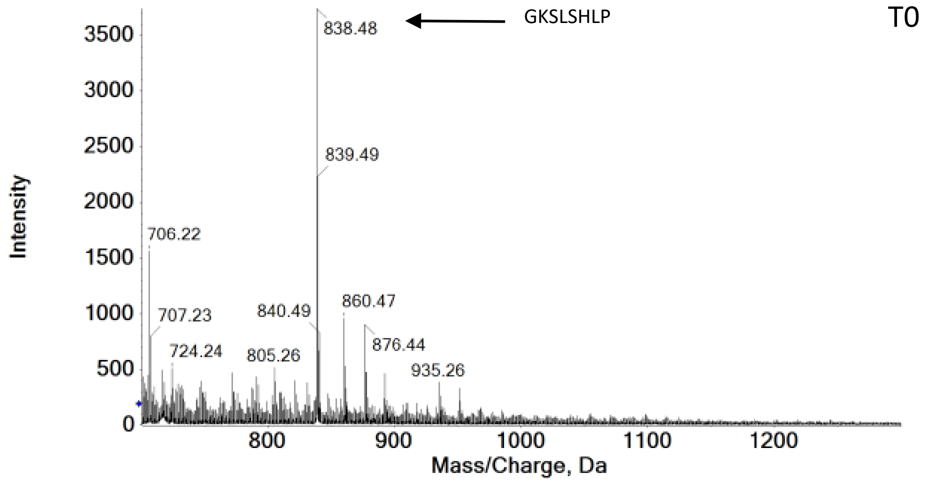
T0
GKSLSHLP
838.48
839.49
706.22
707.23
724.24
805.26
840.49
860.47
876.44
935.26
Intensity
Mass/Charge, Da

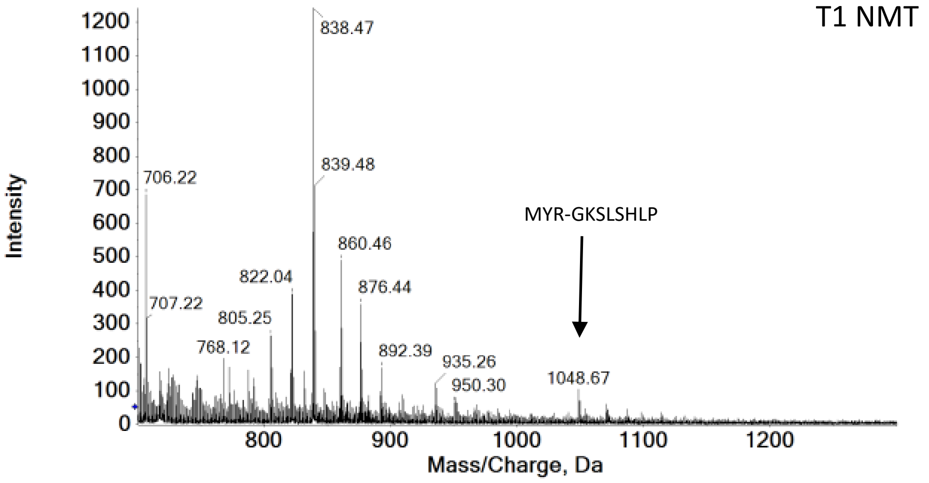
T1 NMT
838.47
839.48
706.22
MYR-GKSLSHLP
860.46
822.04
876.44
707.22
805.25
768.12
892.39
935.26
950.30
1048.67
Intensity
Mass/Charge, Da

### GKTFSQLG

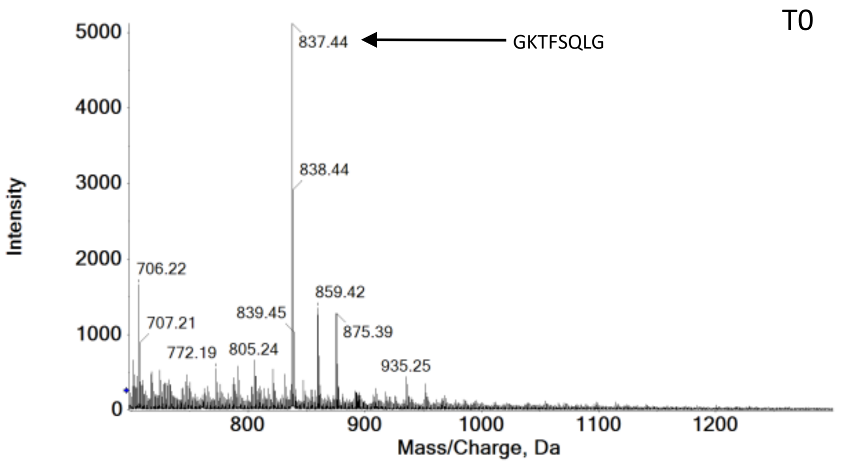
T0
GKTFSQLG
837.44
838.44
706.22
707.21
772.19
805.24
839.45
859.42
875.39
935.25
Intensity
Mass/Charge, Da

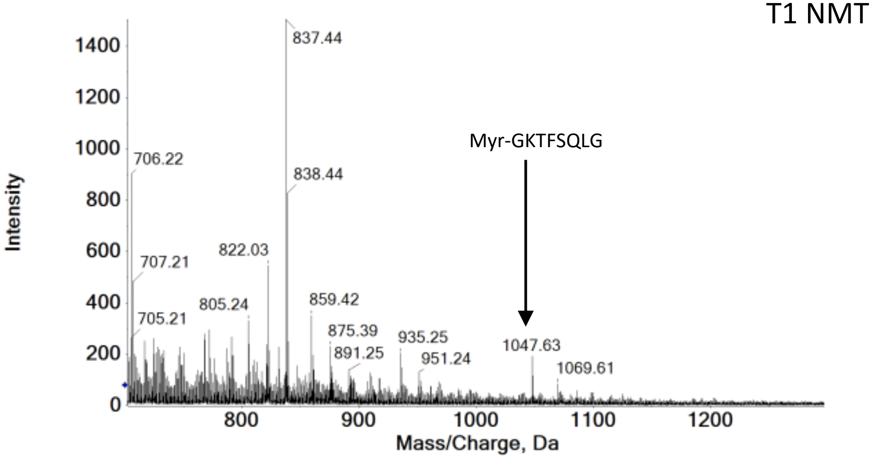
T1 NMT
Myr-GKTFSQLG
837.44
838.44
706.22
707.21
705.21
805.24
822.03
859.42
875.39
891.25
935.25
951.24
1047.63
1069.61
Intensity
Mass/Charge, Da

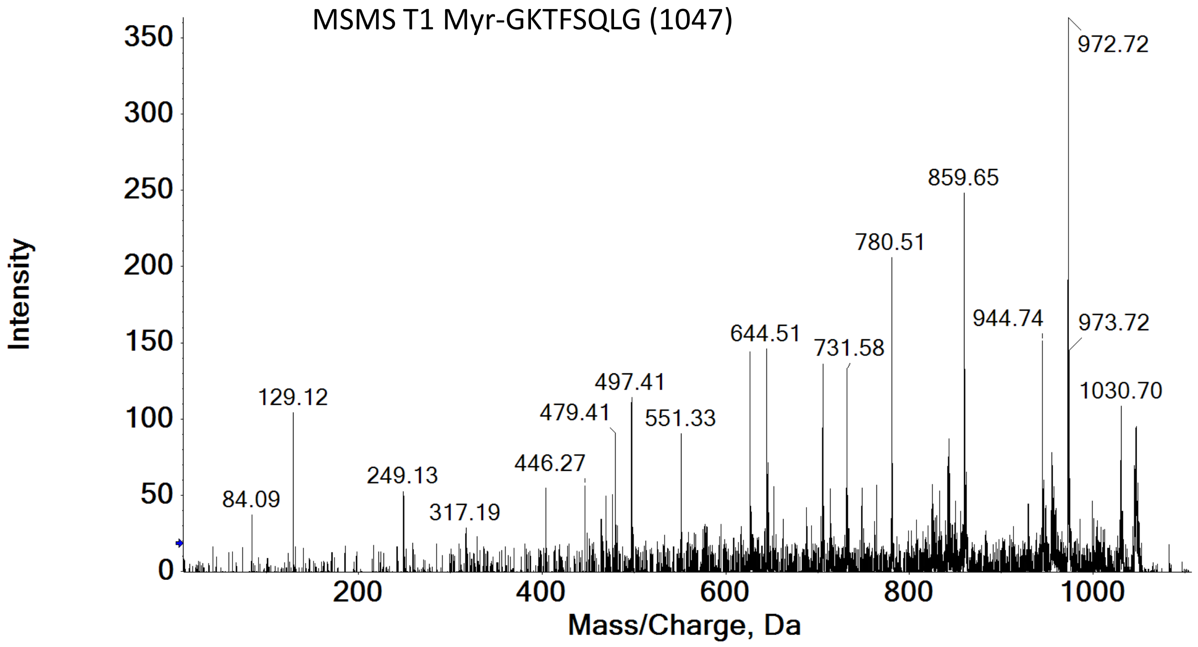
MSMS T1 Myr-GKTFSQLG (1047)
972.72
859.65
780.51
944.74
973.72
644.51
731.58
497.41
479.41
551.33
129.12
1030.70
446.27
249.13
84.09
317.19
Intensity
Mass/Charge, Da

### GKSASKQF

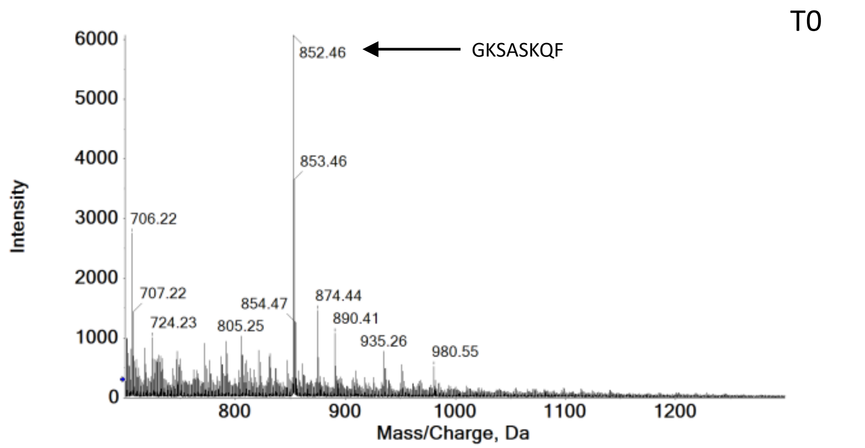
T0
GKSASKQF
852.46
853.46
706.22
707.22
724.23
805.25
854.47
874.44
890.41
935.26
980.55
Intensity
Mass/Charge, Da

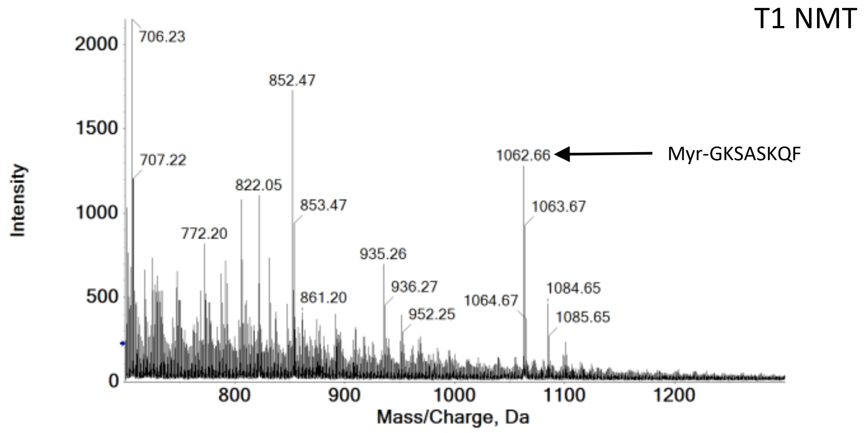
T1 NMT
706.23
852.47
707.22
822.05
772.20
853.47
1062.66
Myr-GKSASKQF
1063.67
935.26
936.27
952.25
861.20
1064.67
1084.65
1085.65
Intensity
Mass/Charge, Da

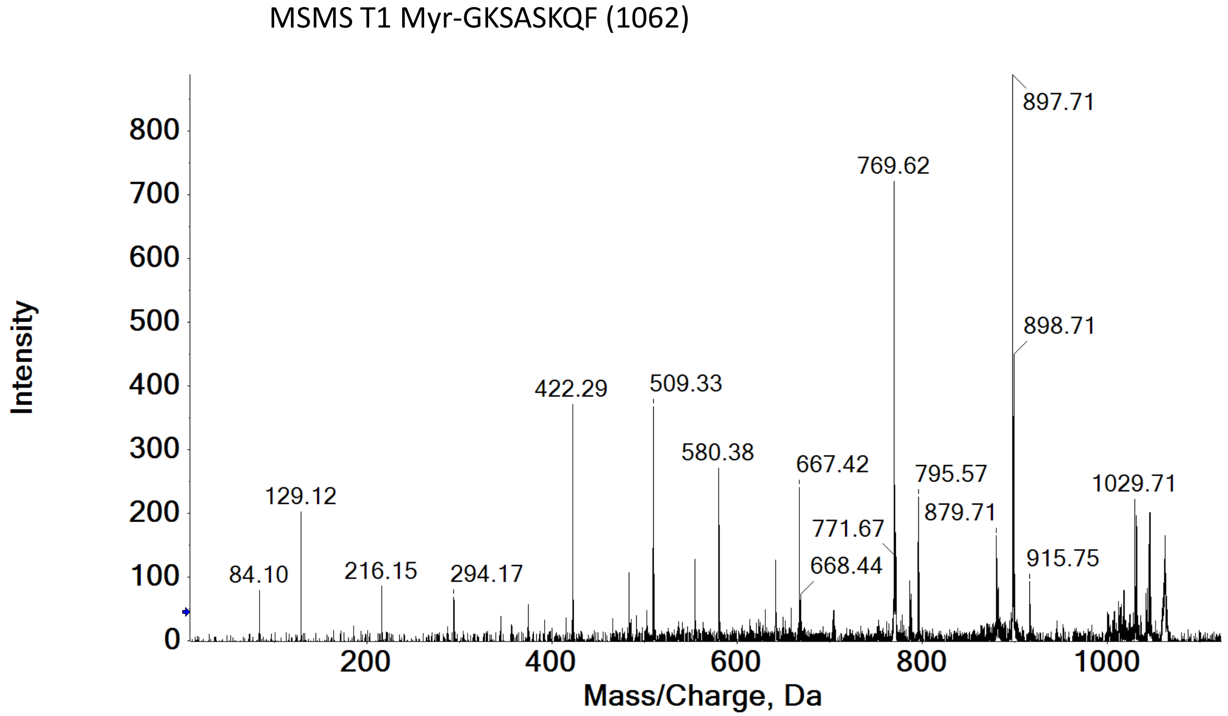
MSMS T1 Myr-GKSASKQF (1062)
897.71
769.62
898.71
422.29
509.33
580.38
667.42
795.57
1029.71
129.12
879.71
771.67
668.44
915.75
84.10
216.15
294.17
Intensity
Mass/Charge, Da

#### GKLQSKHA

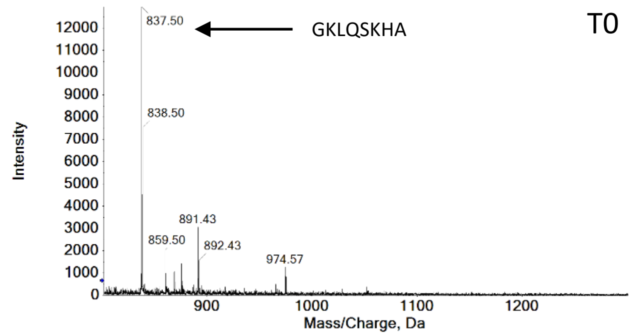
T0
GKLQSKHA
837.50
838.50
891.43
859.50
892.43
974.57
Intensity
Mass/Charge, Da

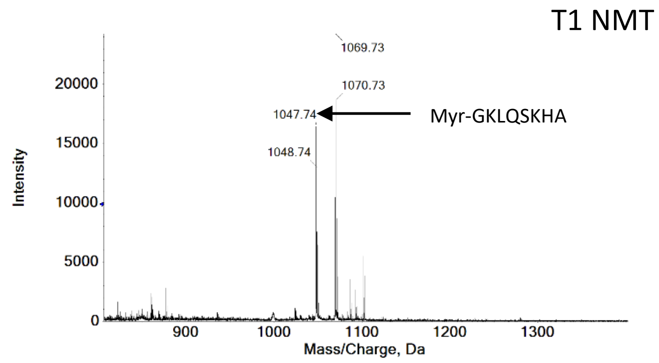
T1 NMT
1069.73
1070.73
1047.74
Myr-GKLQSKHA
1048.74
Intensity
Mass/Charge, Da

### GKLQSKHA

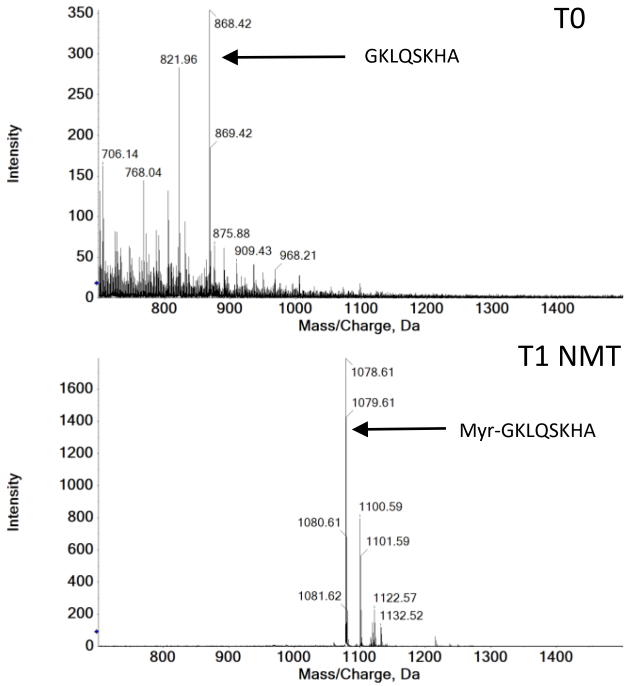

T0
GKLQSKHA
Intensity
Mass/Charge, Da
868.42
821.96
869.42
706.14
768.04
875.88
909.43
968.21
T1 NMT
Myr-GKLQSKHA
1078.61
1079.61
1080.61
1100.59
1101.59
1081.62
1122.57
1132.52


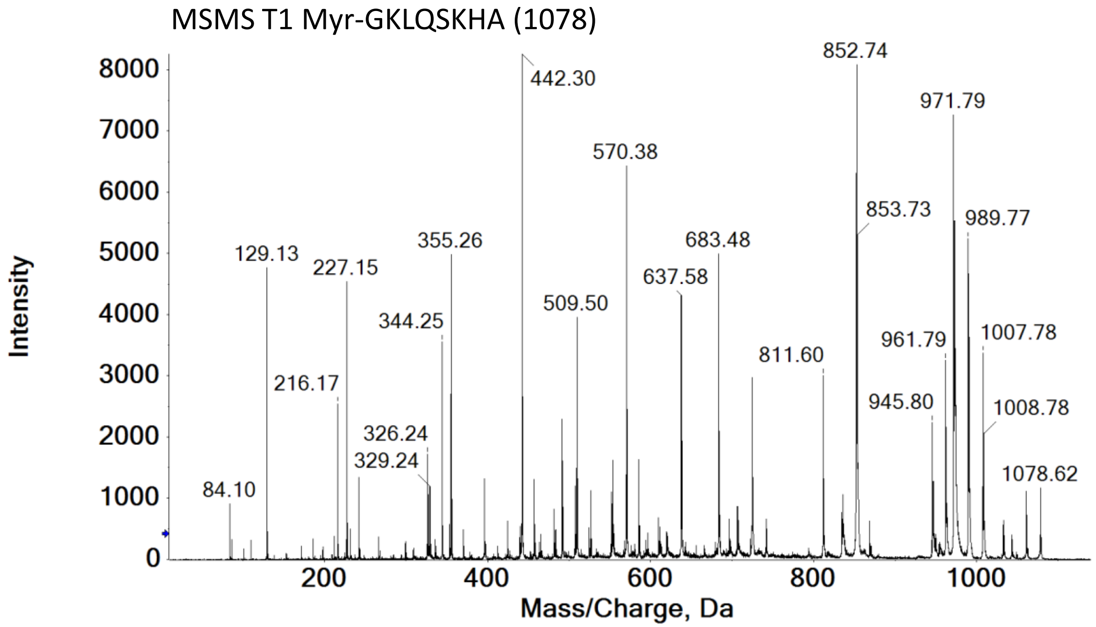

MSMS T1 Myr-GKLQSKHA (1078)
Intensity
Mass/Charge, Da
852.74
442.30
971.79
570.38
853.73
989.77
129.13
227.15
355.26
683.48
637.58
509.50
344.25
961.79
1007.78
811.60
216.17
945.80
1008.78
326.24
329.24
1078.62
84.10

### GAKQSGPA

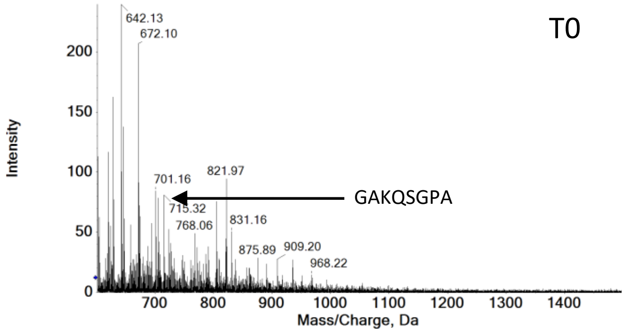
T0
642.13
672.10
701.16
821.97
715.32
768.06
831.16
875.89
909.20
968.22
GAKQSGPA
Intensity
Mass/Charge, Da

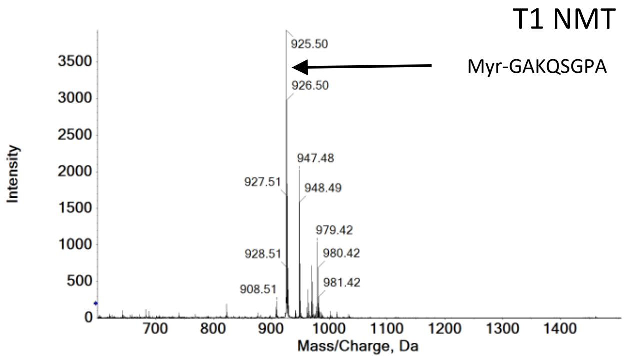
T1 NMT
925.50
926.50
Myr-GAKQSGPA
947.48
927.51
948.49
979.42
928.51
980.42
981.42
908.51
Intensity
Mass/Charge, Da

### GGKQSTAA

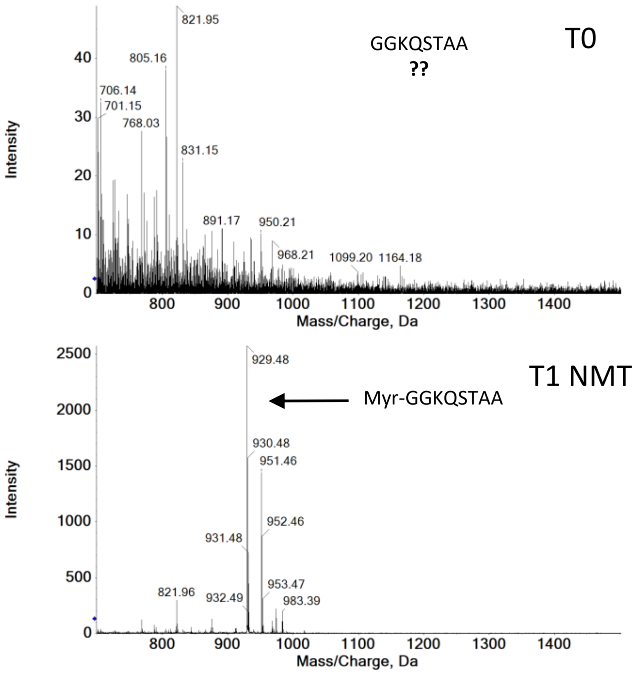


MSMS T1 Myr-GGKQSTAA (929)

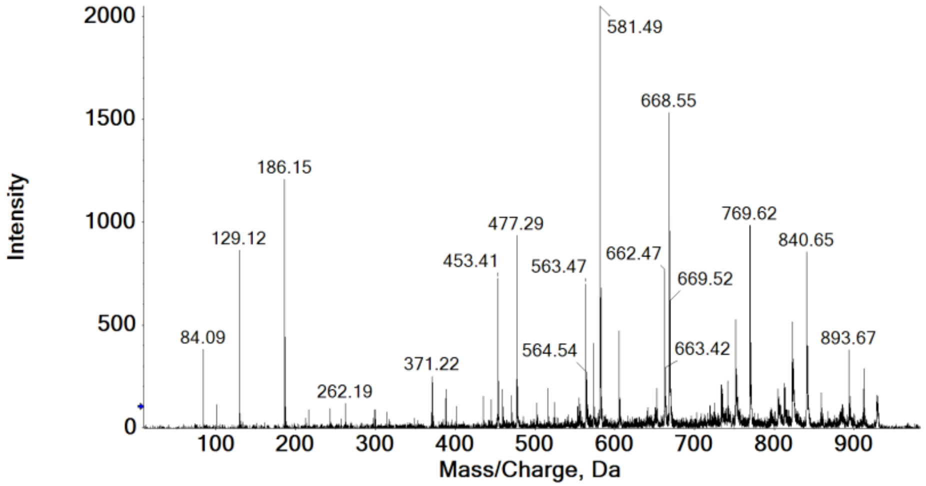

### GGKFSSKPR

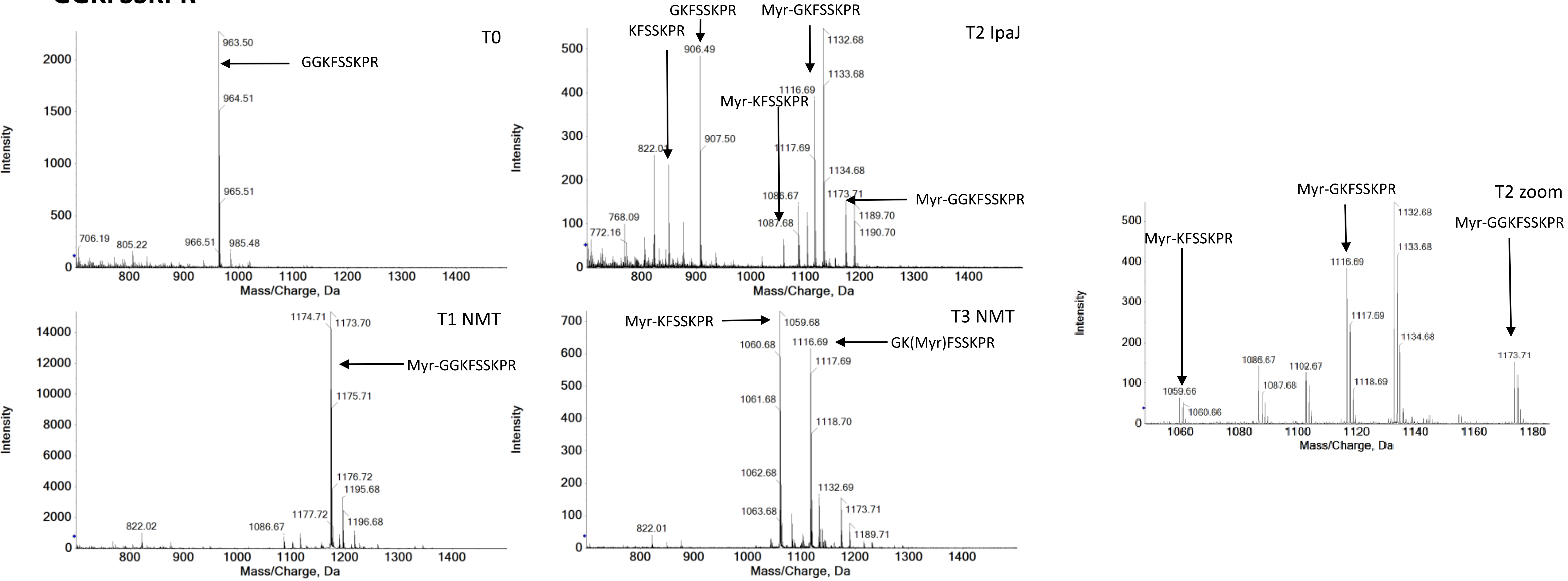
T0
GGKFSSKPR
963.50
964.51
965.51
966.51
985.48
706.19
805.22
Intensity
Mass/Charge, Da
T2 IpaJ
GKFSSKPR
Myr-GKFSSKPR
KFSSKPR
Myr-KFSSKPR
Myr-GGKFSSKPR
906.49
907.50
822.01
768.09
772.16
1132.68
1133.68
1134.68
1116.69
1117.69
1086.67
1087.68
1173.71
1189.70
1190.70
T1 NMT
Myr-GGKFSSKPR
1174.71
1173.70
1175.71
1176.72
1195.68
1196.68
1177.72
1086.67
822.02
T3 NMT
Myr-KFSSKPR
GK(Myr)FSSKPR
1059.68
1060.68
1061.68
1062.68
1063.68
1116.69
1117.69
1118.70
1132.69
1173.71
1189.71
822.01
T2 zoom
Myr-KFSSKPR
Myr-GKFSSKPR
Myr-GGKFSSKPR
1059.66
1060.66
1086.67
1087.68
1102.67
1116.69
1117.69
1118.69
1132.68
1133.68
1134.68
1173.71

### GGKFSSKPR

MSMS T0 GGKFSSKPR (963)

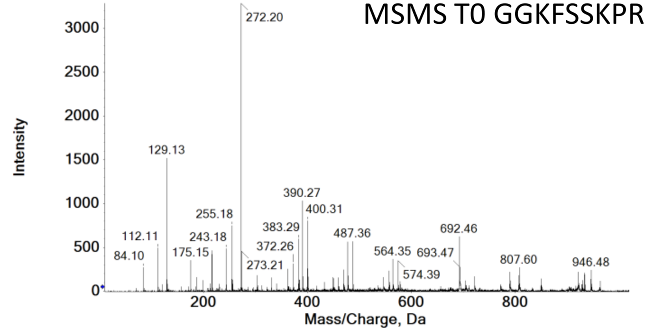


MSMS T3 GK(Myr)FSSKPR (1116)

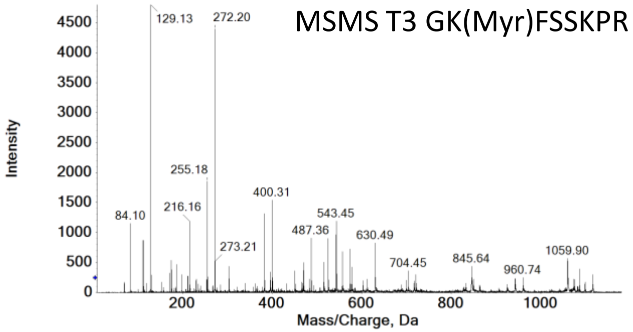


MSMS T3 Myr-KFSSKPR (1059)

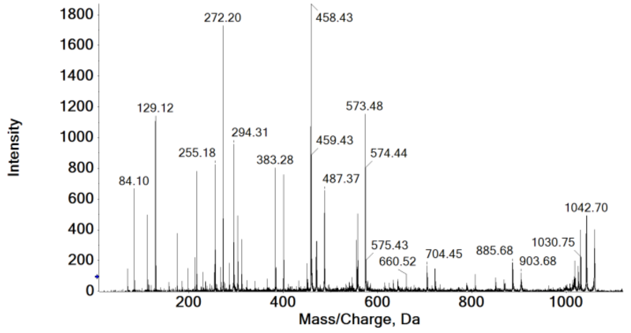


MSMS T0 Myr-GGKFSSKPR (1173)

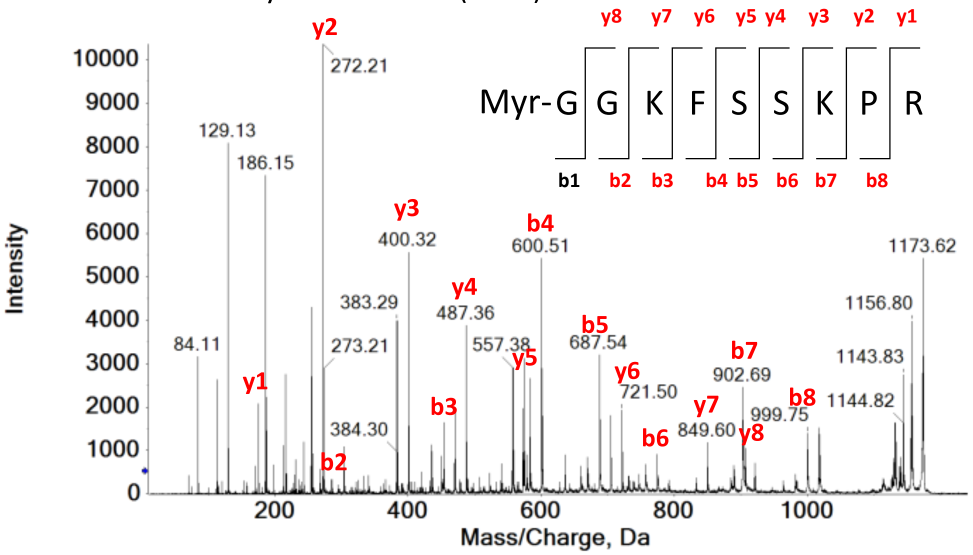


| K20 1173 | Mass/Charge (Da) | Intensity | |
|---|---|---|---|
| 4372 | 175,1503 | 2072 | y1 |
| 8355 | 272,1954 | 10360,16 | y2 |
| 10293 | 325,2772 | 320 | b2 |
| 12791 | 400,3083 | 5570,51 | y3 |
| 14427 | 453,4173 | 1645,961 | b3 |
| 15423 | 487,3767 | 3883,765 | y4 |
| 17825 | 574,4145 | 3119,059 | y5 |
| 18508 | 600,4936 | 5425,726 | b4 |
| 20684 | 687,5204 | 3206,745 | b5 |
| 21495 | 721,4933 | 1923,922 | y6 |
| 22724 | 774,5721 | 912,7844 | b6 |
| 24390 | 849,6013 | 1189,02 | y7 |
| 25523 | 902,6548 | 2460,706 | b7 |
| 25606 | 906,606 | 1046,588 | y8 |
| 27511 | 999,7244 | 1402,039 | b8 |

MSMS T3 GK(Myr)FSSKPR (1116)

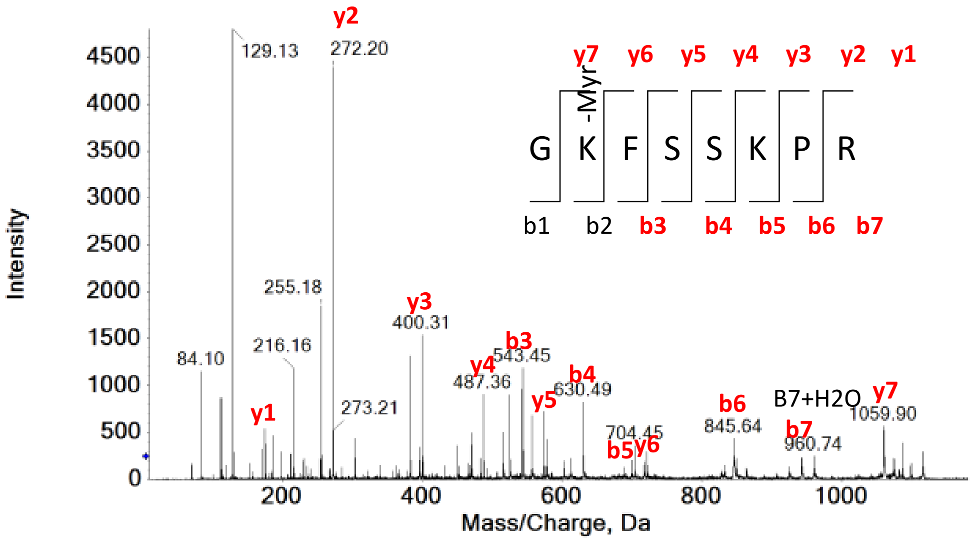


| K22 1116 | Mass/Charge (Da) | Intensity | |
|---|---|---|---|
| 2008 | 175,1541 | 538,35 | y1 |
| 2891 | 216,1627 | 1182,90 | SK |
| 3999 | 255,1691 | 1852,24 | y2-NH3 |
| 4553 | 272,2057 | 4394,67 | y2 |
| 5462 | 303,1855 | 437,33 | SSK |
| 7699 | 383,2735 | 1310,43 | SSKP-NH3 |
| 8218 | 400,3085 | 1541,02 | y3 |
| 9657 | 450,2645 | 359,53 | FSSK |
| 10244 | 470,3054 | 500,24 | y4-NH3 |
| 10729 | 487,3455 | 905,41 | y4 |
| 11494 | 515,4305 | 506,04 | a3 |
| 12266 | 543,4355 | 1191,69 | b3 |
| 13097 | 574,384 | 724,55 | y5 |
| 14532 | 630,4798 | 822,75 | b4 |
| 16634 | 717,5125 | 190,59 | b5 |
| 16727 | 721,4527 | 298,67 | y6 |
| 19445 | 845,6232 | 432,31 | b6 |
| 21453 | 942,7505 | 231,69 | b7 |
| 21812 | 960,7414 | 246,90 | b7+H2O |
| 23625 | 1059,864 | 569,10 | y7 |

### GGKLSKKK

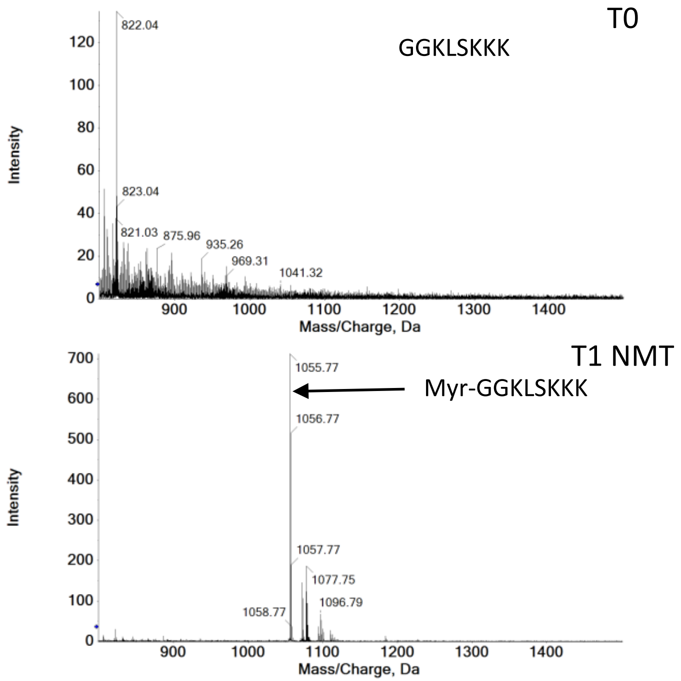


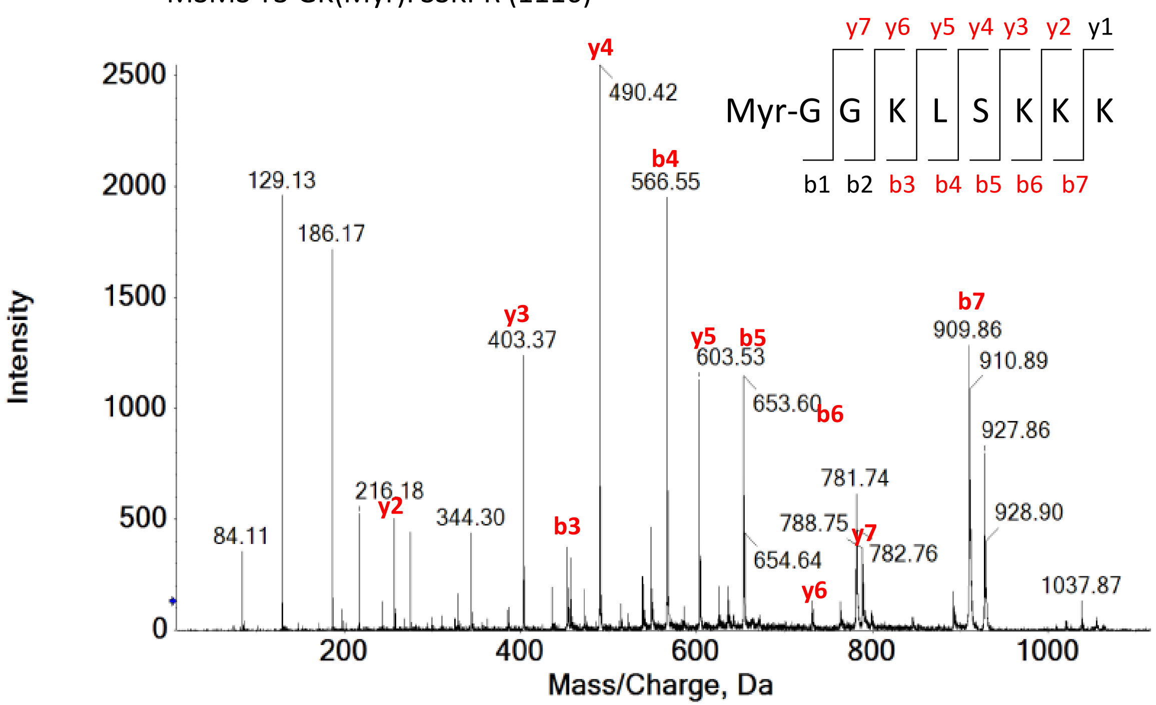


| | | | |
|---|---|---|---|
| y2 | 2642 | 275,2397 | 400,9412 |
| y3 | 5363 | 403,3621 | 1237,961 |
| b3 | 6526 | 453,4303 | 375,8431 |
| y4 | 7591 | 490,8696 | 179,6078 |
| b4 | 9598 | 566,5328 | 1952,314 |
| y5 | 10561 | 603,5288 | 1129,098 |
| b5 | 11824 | 653,5715 | 1144,941 |
| y6 | 13686 | 731,6654 | 131,2941 |
| b6 | 14812 | 781,6852 | 615,8431 |
| y7 | 14969 | 788,6453 | 359,6863 |
| b7 | 17555 | 909,8929 | 1265,882 |

#### GCSVKKK

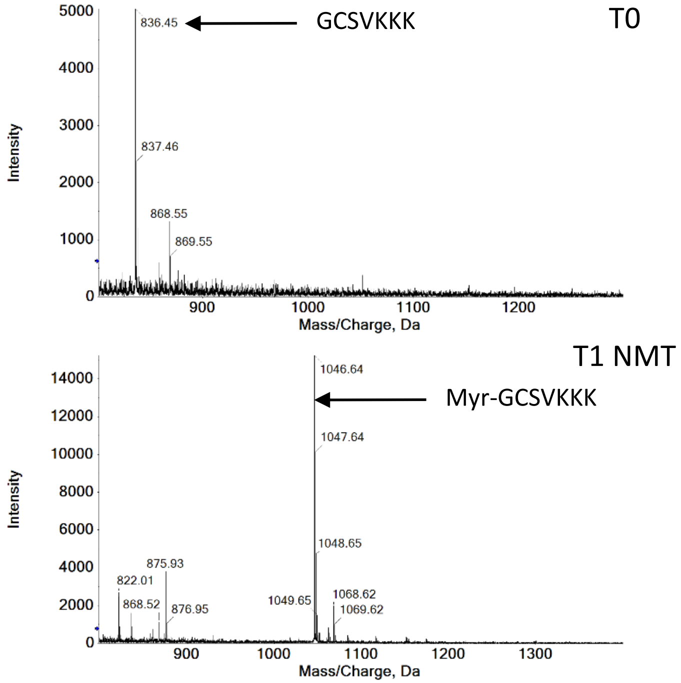
T0
GCSVKKK
836.45
837.46
868.55
869.55
Intensity
Mass/Charge, Da
T1 NMT
Myr-GCSVKKK
1046.64
1047.64
1048.65
1049.65
1068.62
1069.62
822.01
875.93
868.52
876.95

### ASSVKKK

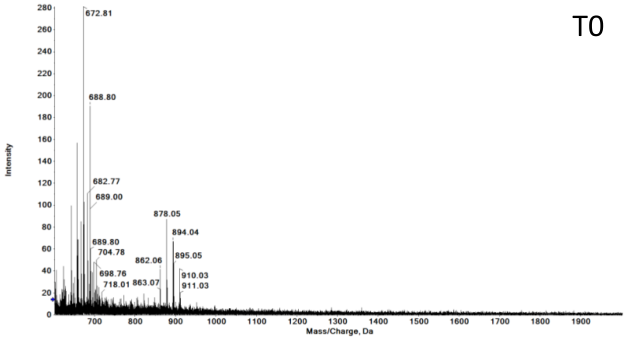
T0
Intensity
Mass/Charge, Da
672.81
688.80
682.77
689.00
878.05
894.04
689.80
704.78
862.06
895.05
698.76
718.01
863.07
910.03
911.03

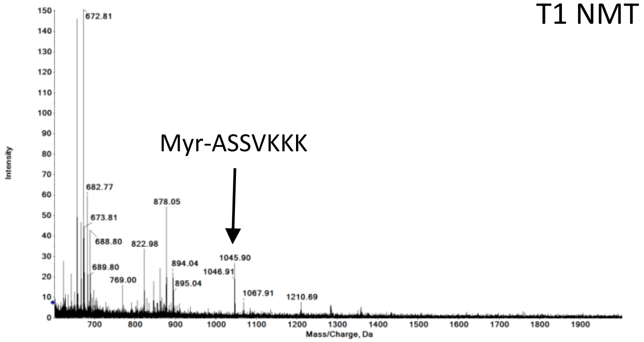
T1 NMT
Myr-ASSVKKK
Intensity
Mass/Charge, Da
672.81
682.77
878.05
673.81
688.80
822.98
1045.90
894.04
1046.91
689.80
769.00
895.04
1067.91
1210.69
