## Supplementary Dataset 1b for "Proteome-wide probing of the dual NMT-dependent myristoylation tradeoff unveils potent, mechanism-based suicide inhibitors"

### Sup Dataset 1b

| Peptide sequence | MS data page in SupDataset 1b |
|---|---|
| GKSNSKLK | 2 |
| GKSNSGLKP | 3 |
| GKSNS{DAP]LKP | 4 |
| GKSNS{DAB]LKP | 5 |
| GKSNS{ORN]LKP | 6 |
| GKSNS{HCY}LKP | 7 |
| GKSNS{bhLYS}LKP | 8 |
| GKVWSQGV | 9 |
| GKSNAKLKP | 10 |

### GKSNSKLK

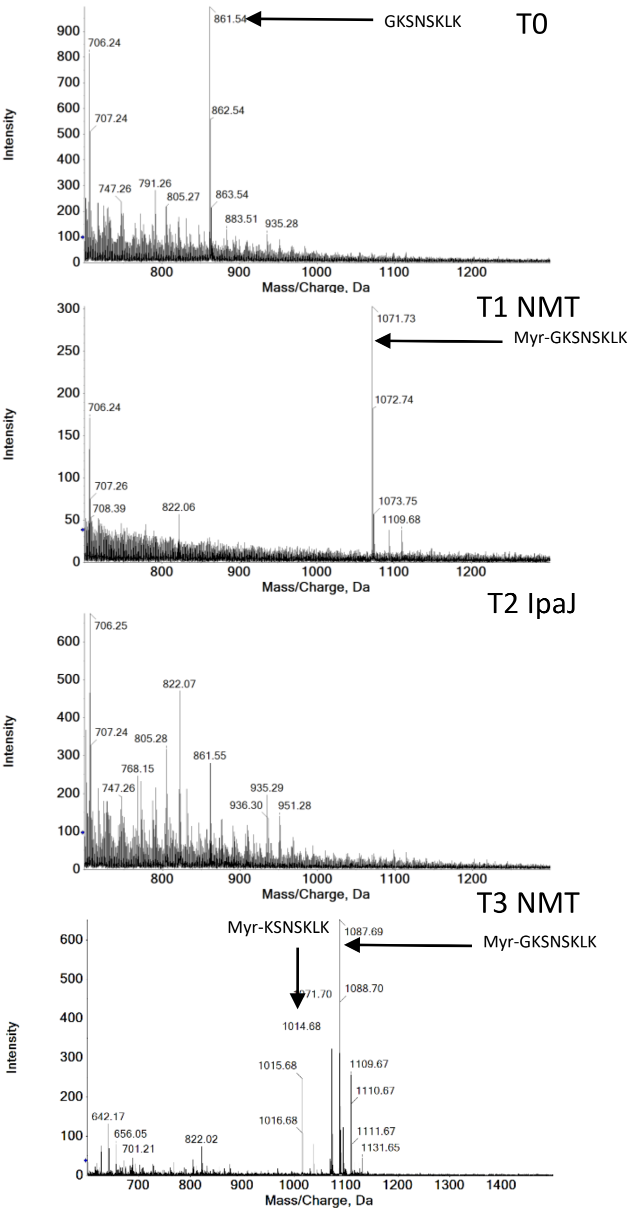


MSMS T3 Myr-GKSNSKLK (1014)

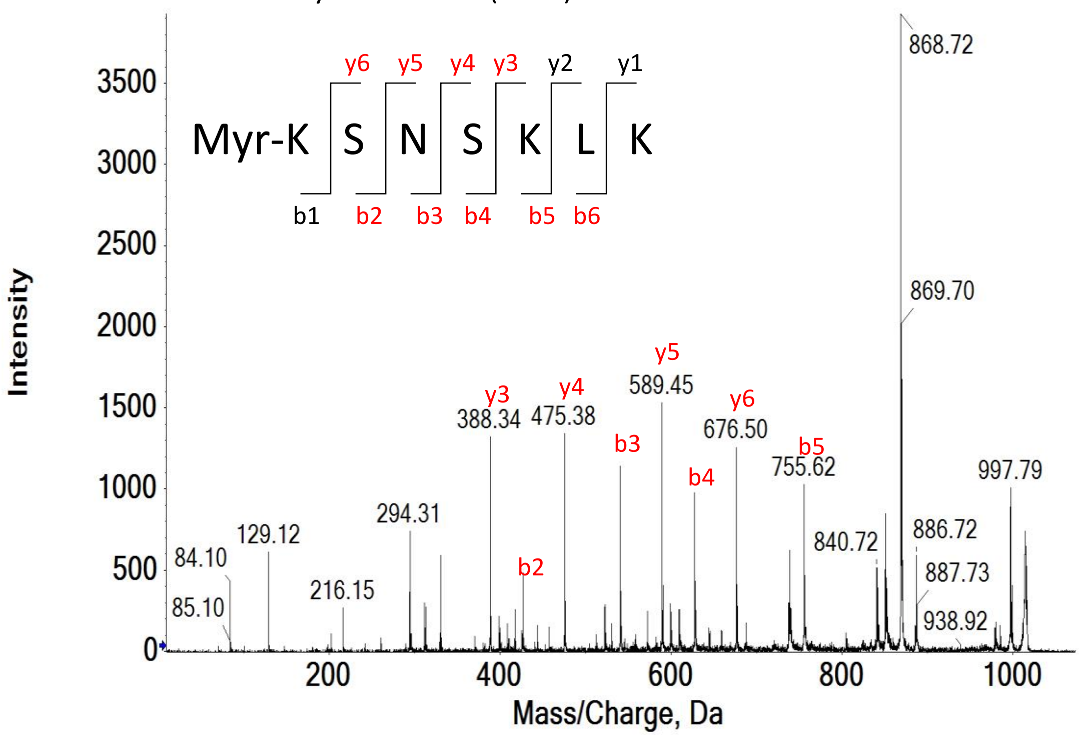


| F9_MSMS_1014.6000_2 | | | ion |
|---|---|---|---|
| pic | Mass/Charge (Da) | Intensity | ion |
| 4036 | 388,3268 | 1320,784 | y3 |
| 5032 | 426,3984 | 484,2353 | b2 |
| 6101 | 475,3693 | 1344,157 | y4 |
| 7622 | 540,4544 | 1142,118 | b3 |
| 8831 | 589,4373 | 1535,059 | y5 |
| 9779 | 627,5093 | 978,5098 | b4 |
| 10957 | 676,5035 | 1258,196 | y6 |
| 12771 | 755,6361 | 1029,804 | b5 |
| 15221 | 868,7002 | 3926,588 | b6 |

#### GKSNSGLKP

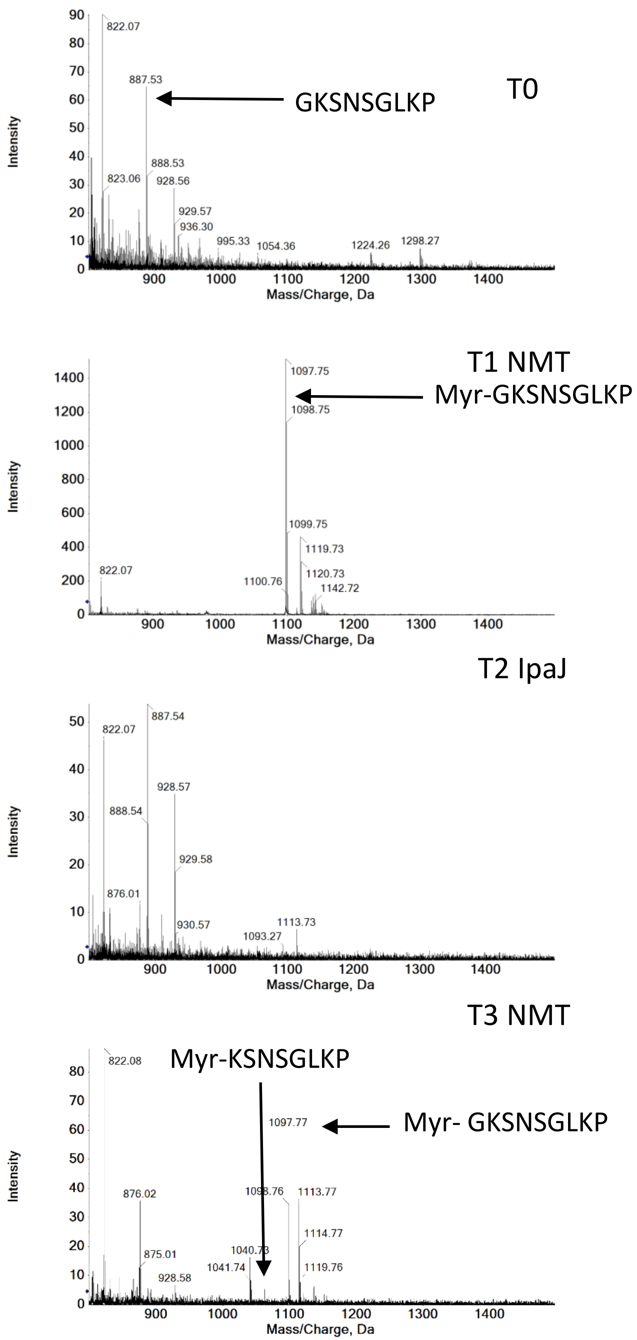

T0
GKSNSGLKP
822.07
887.53
888.53
823.06
928.56
929.57
936.30
995.33
1054.36
1224.26
1298.27
T1 NMT
Myr-GKSNSGLKP
1097.75
1098.75
1099.75
1119.73
1120.73
1142.72
1100.76
822.07
T2 IpaJ
822.07
887.54
928.57
888.54
929.58
876.01
930.57
1113.73
1093.27
T3 NMT
Myr-KSNSGLKP
Myr- GKSNSGLKP
822.08
1097.77
876.02
1113.77
1114.77
1119.76
875.01
928.58
1041.74
Intensity
Mass/Charge, Da


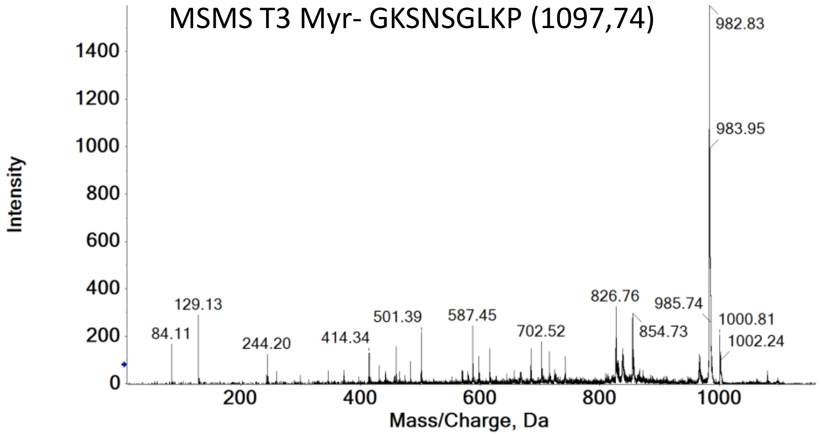

MSMS T3 Myr- GKSNSGLKP (1097,74)
982.83
983.95
129.13
84.11
244.20
414.34
501.39
587.45
702.52
826.76
854.73
985.74
1000.81
1002.24
Intensity
Mass/Charge, Da


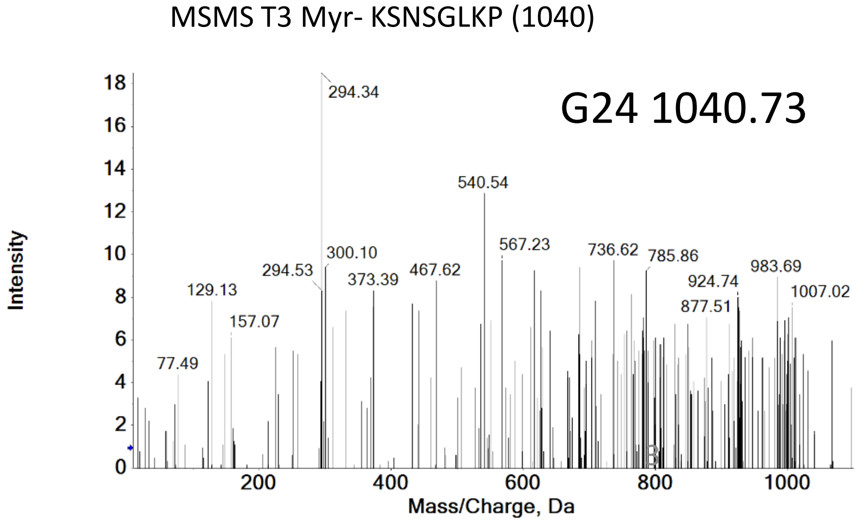

MSMS T3 Myr- KSNSGLKP (1040)
G24 1040.73
294.34
540.54
567.23
736.62
785.86
300.10
294.53
373.39
467.62
129.13
157.07
77.49
924.74
877.51
983.69
1007.02
Intensity
Mass/Charge, Da

### GKSNS{DAP}LKP

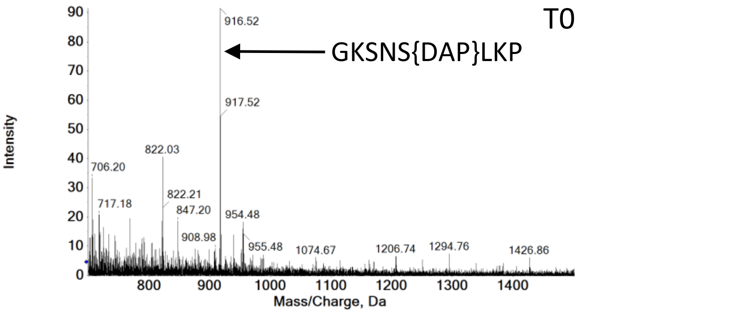


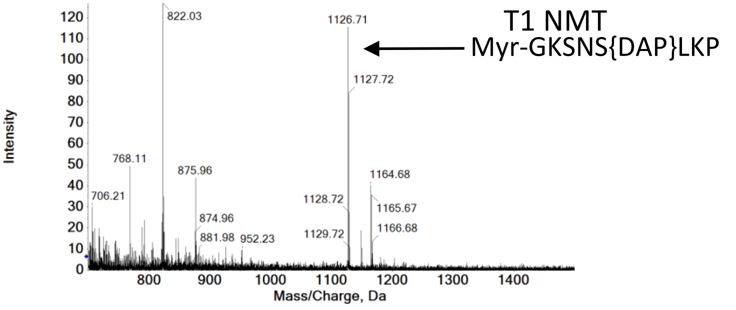


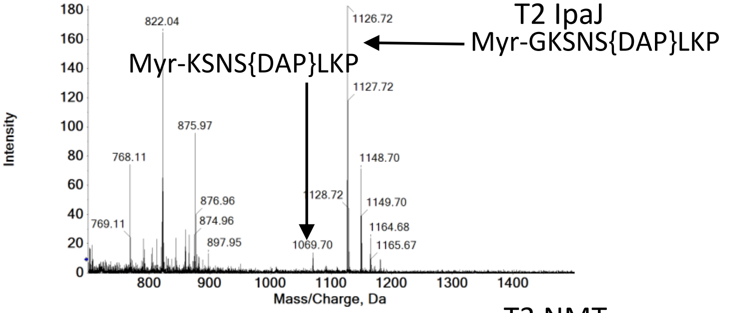


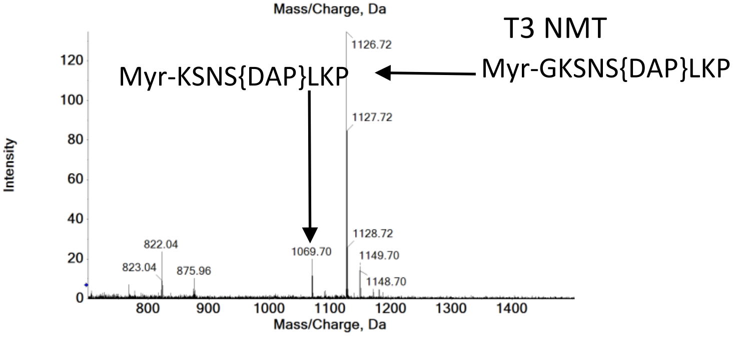


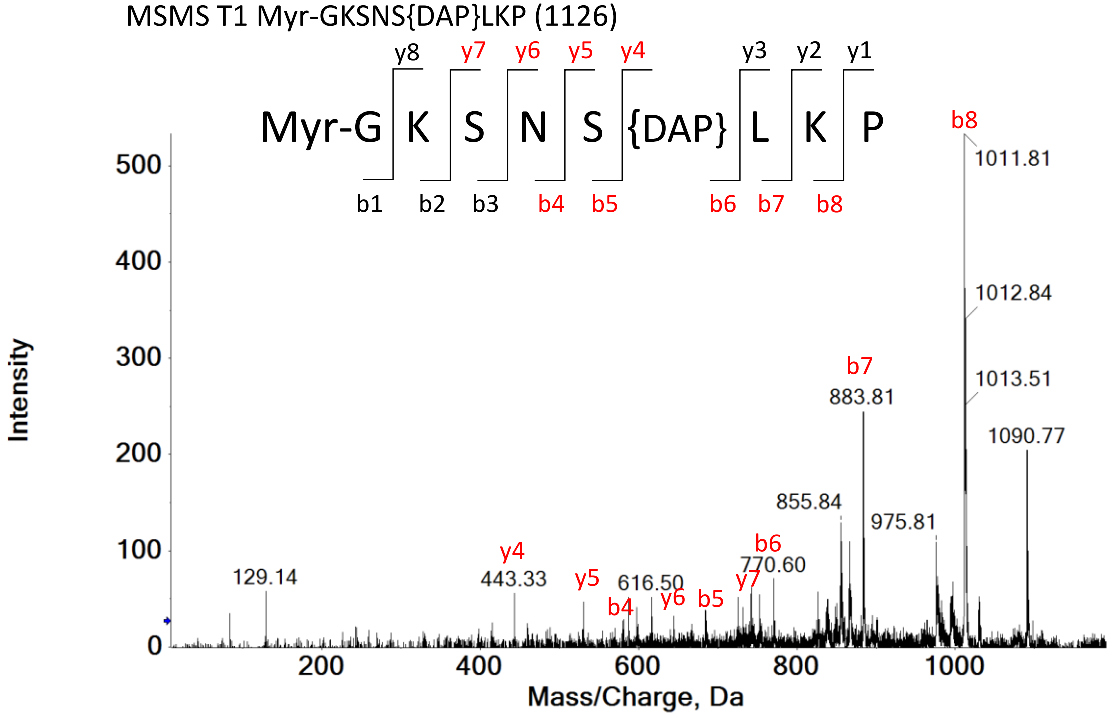


| H24_MS MS_1126 .7100_2 | Mass/Charge (Da) | Intensity | |
|---|---|---|---|
| 1664 | 443,3184 | 55,84314 | y4 |
| 2540 | 530,3643 | 47,21569 | y5 |
| 3164 | 587,4767 | 51,60785 | b4 |
| 3605 | 616,4448 | 51,92157 | SNSK(-42)LK |
| 3998 | 644,3611 | 32,31373 | y6 |
| 4636 | 684,5457 | 38,43137 | b5 |
| 5443 | 731,7719 | 41,41177 | y7 |
| 6224 | 770,6158 | 70,90196 | b6 |
| 7758 | 855,6877 | 122,0392 | a7 |
| 8332 | 883,8343 | 245,0196 | b7 |
| 10628 | 1011,857 | 533,3334 | b8 |

### GKSNS{DAB]LKP

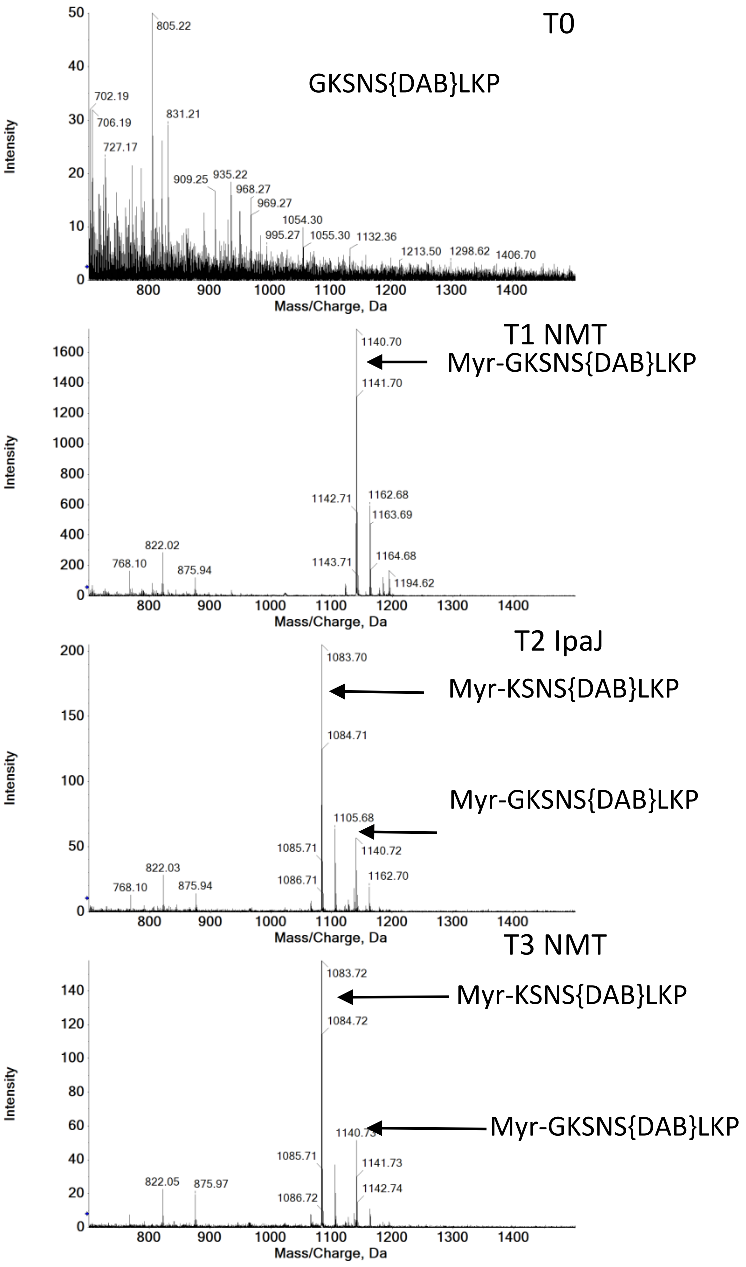


MSMS T1 Myr-GKSNS{DAB}LKP (1140)

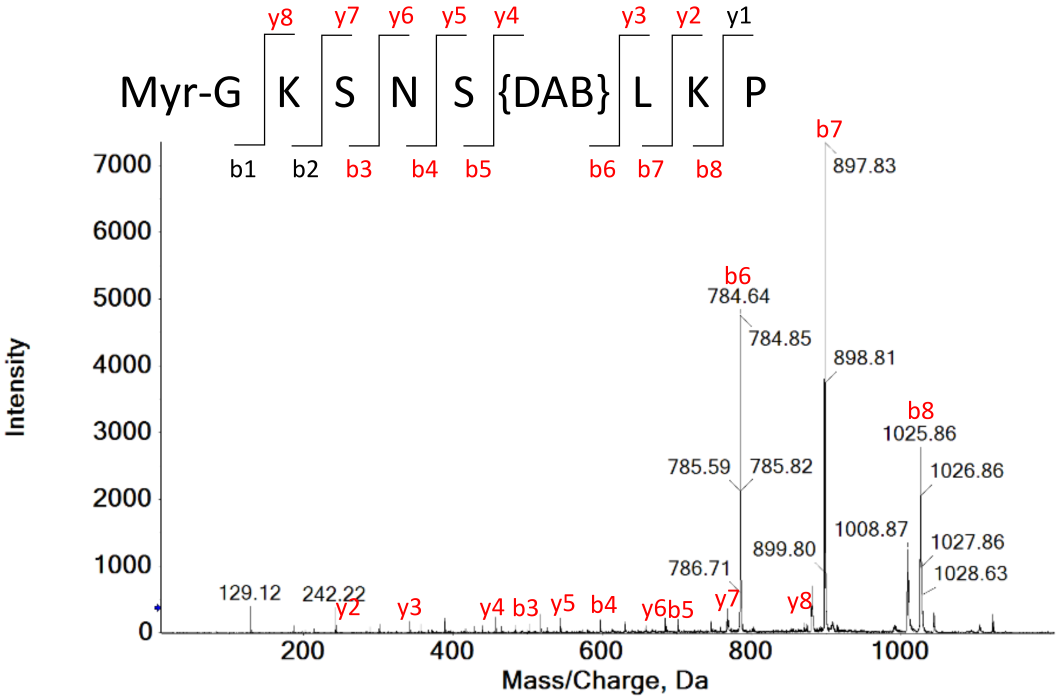


| H19_MSMS _1140.6900 _2 | Mass/Charge (Da) | Intensity | |
|---|---|---|---|
| 2456 | 242,2261 | 376,9412 | LK |
| 2532 | 244,2031 | 115,1373 | y2 |
| 4318 | 357,2703 | 131,6078 | y3 |
| 6462 | 457,3843 | 234,8235 | y4 |
| 7032 | 483,4075 | 116,8627 | b3 |
| 7891 | 517,3342 | 283,6078 | KSNSK(-28) |
| 8561 | 544,3931 | 224,3137 | y5 |
| 9868 | 597,4956 | 180,2353 | b4 |
| 11358 | 658,4525 | 103,8431 | y6 |
| 11985 | 684,5401 | 220,3922 | b5 |
| 12393 | 701,5362 | 204,3922 | c5 |
| 13419 | 745,4974 | 157,6471 | y7 |
| 14318 | 784,8348 | 4759,687 | b6 |
| 16184 | 869,7394 | 146,8235 | a7 |
| 16271 | 873,8047 | 141,1765 | y8 |
| 16781 | 897,8309 | 7348,549 | b7 |
| 19047 | 1008,825 | 1253,177 | b8-NH3 |
| 19384 | 1025,869 | 2782,431 | b8 |

MSMS T3 Myr-KSNS{DAB}LKP (1083)

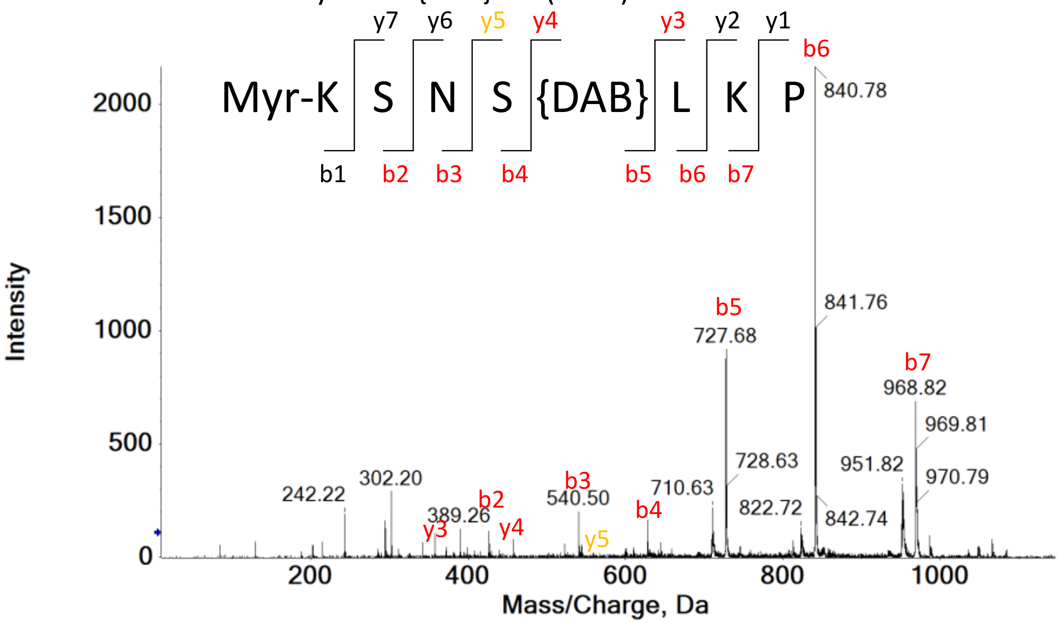


| H21_MSMS _1083.7100 _2 | Mass/Charge (Da) | Intensity | |
|---|---|---|---|
| 1610 | 357,2901 | 105,4118 | y3 |
| 2513 | 426,3815 | 116,2353 | b2 |
| 2908 | 457,3836 | 81,09805 | y4 |
| 3906 | 540,4713 | 202,1961 | b3 |
| 4011 | 544,3975 | 57,72549 | y5 |
| 5351 | 627,5062 | 164,7059 | b4 |
| 6940 | 710,5687 | 219,9216 | b5-NH3 |
| 7330 | 727,6011 | 920,1569 | b5 |
| 9182 | 822,6997 | 130,353 | b6-H2O |
| 9577 | 840,8204 | 2164,549 | b6 |
| 11624 | 951,7894 | 324,0784 | b7-NH3 |
| 11970 | 968,7888 | 688,3137 | b7 |

### GKSNS{ORN]LKP

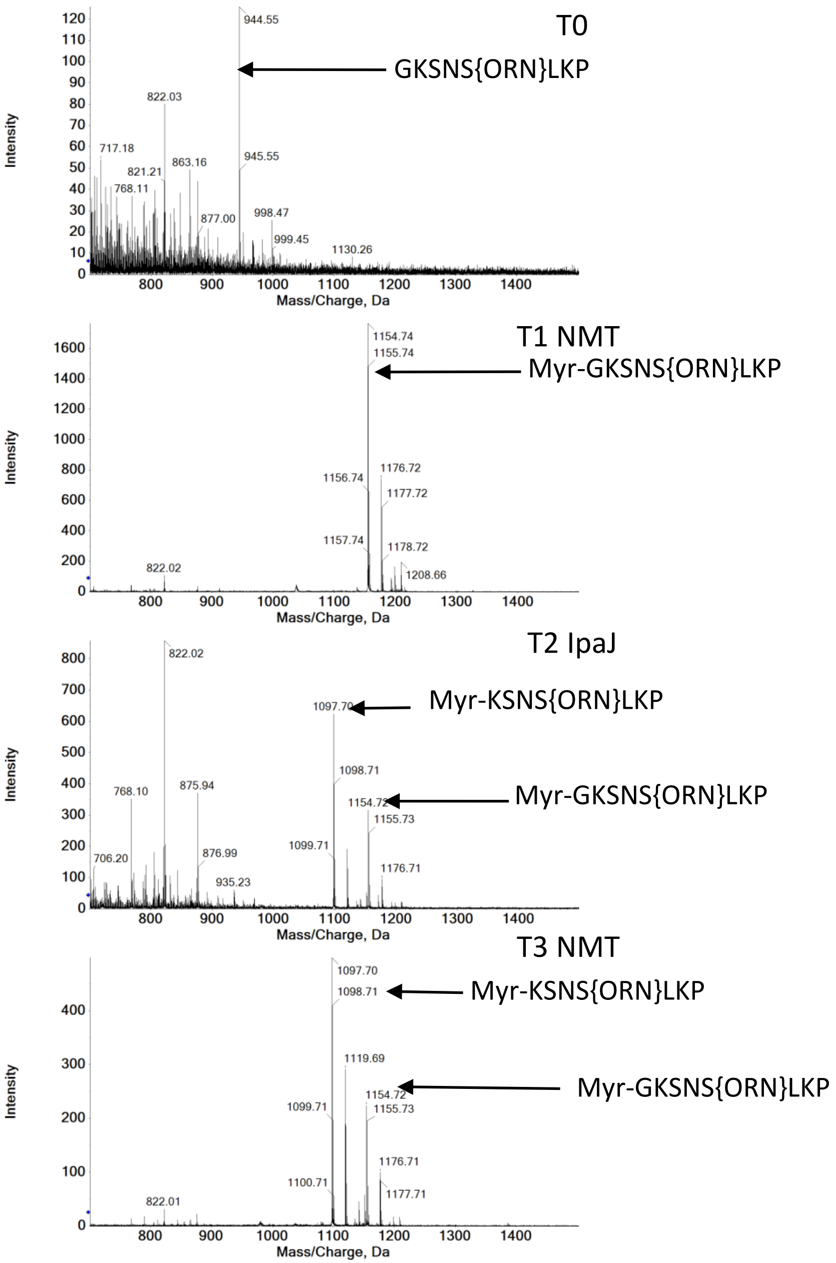


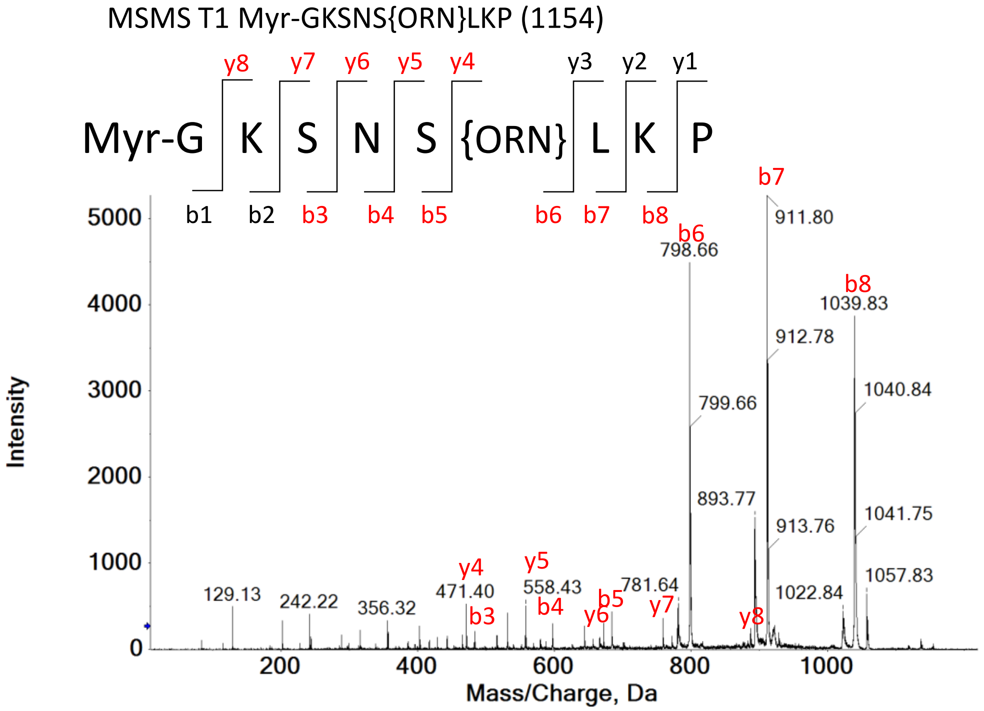


| I3_MSMS _1154.7300_2 | Mass/Charge (Da) | Intensity | |
|---|---|---|---|
| 5743 | 471,3982 | 528,3137 | y4 |
| 6033 | 483,4253 | 213,0196 | b3 |
| 7833 | 558,4199 | 504,7843 | y5 |
| 8842 | 597,5019 | 298,8235 | b4 |
| 10683 | 672,4924 | 301,6471 | y6 |
| 10976 | 684,5358 | 438,7451 | b5 |
| 12735 | 759,5251 | 359,6863 | y7 |
| 13621 | 798,6034 | 4492,079 | b6 |
| 15564 | 887,8157 | 240,3137 | y8 |
| 15689 | 893,7206 | 1531,765 | b7-H2O |
| 16070 | 911,8423 | 5269,333 | b7 |
| 18657 | 1039,824 | 3872,628 | b8 |

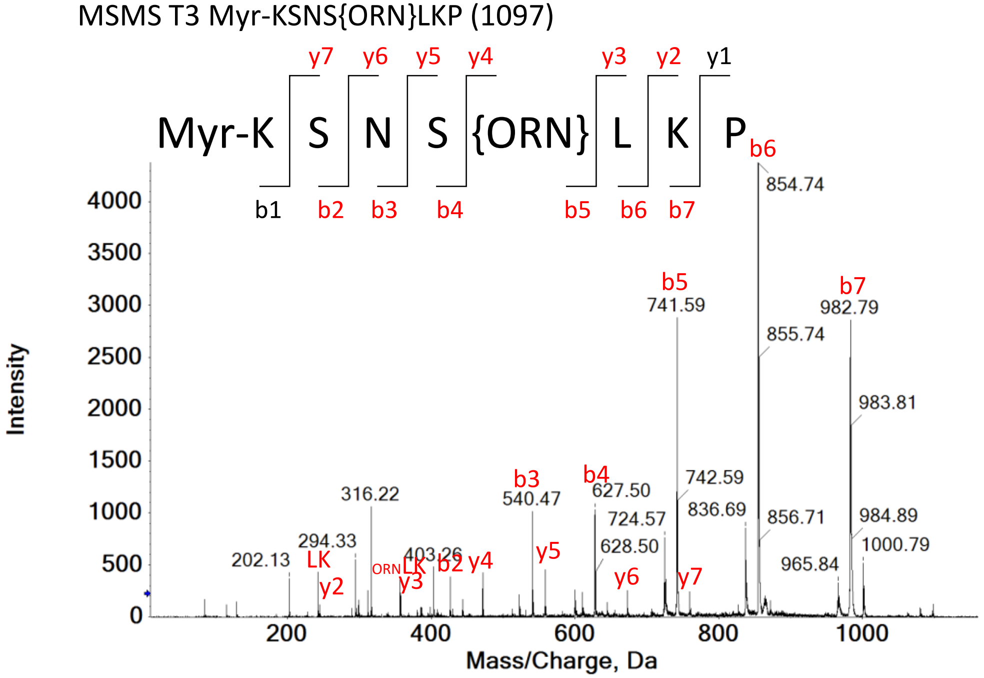


| I5_MSMS _1097.7000_2 | Mass/Charge (Da) | Intensity | |
|---|---|---|---|
| 921 | 242,2105 | 432,4706 | LK |
| 975 | 244,1879 | 116,2353 | y2 |
| 2121 | 316,2183 | 1062,902 | NSK(-14) |
| 2755 | 356,3063 | 358,902 | K(-14)LK |
| 2788 | 357,2847 | 229,6471 | y3 |
| 3759 | 403,2611 | 479,8431 | SNSK(-14) |
| 4264 | 426,4007 | 383,8431 | b2 |
| 4651 | 443,383 | 167,5294 | SK(-14)LK |
| 5194 | 471,4 | 426,8235 | y4 |
| 6713 | 540,4425 | 1013,804 | b3 |
| 7165 | 558,4264 | 452,2353 | y5 |
| 8818 | 627,5032 | 1036,235 | b4 |
| 9919 | 672,5188 | 246,7451 | y6 |
| 11532 | 741,5885 | 2886,588 | b5 |
| 11946 | 759,515 | 245,1765 | y7 |
| 14059 | 854,6556 | 4378,196 | b6 |
| 16703 | 982,7454 | 2862,275 | b7 |

### GKSNS{HCY}LKP

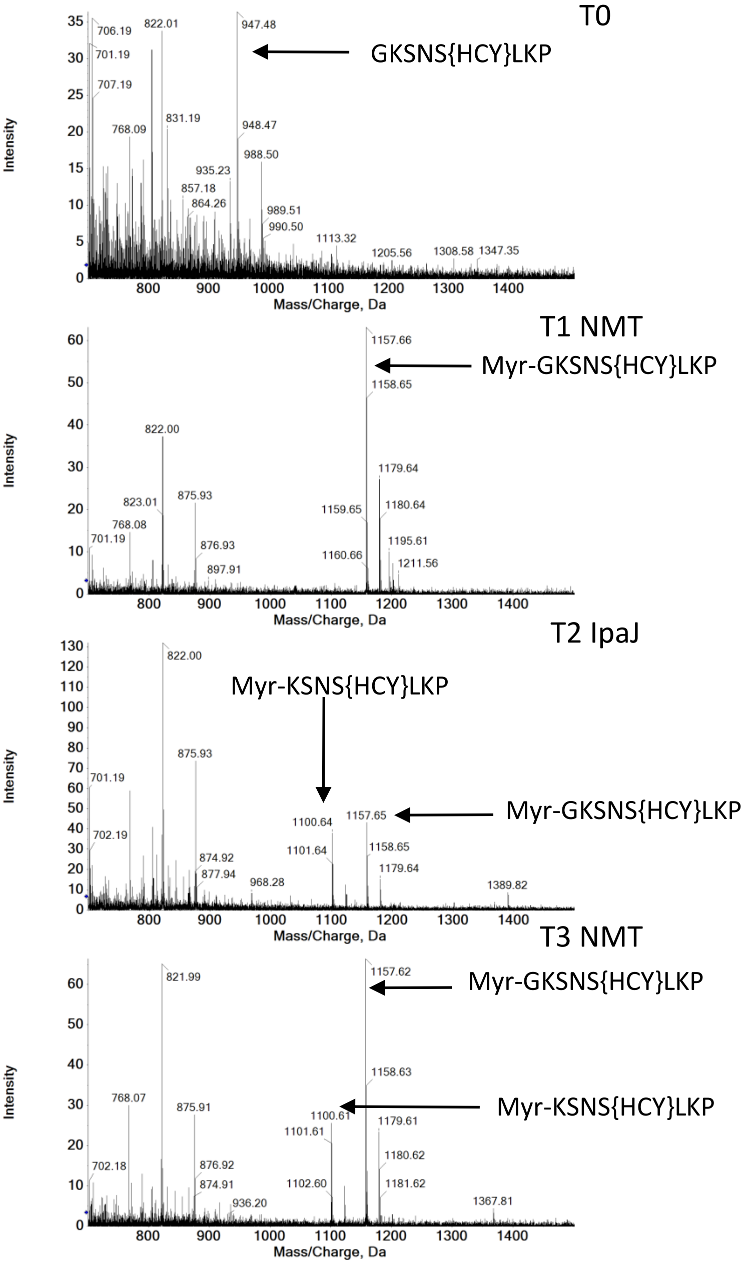


MSMS T1 Myr-GKSNS{HCY}LKP (1157)

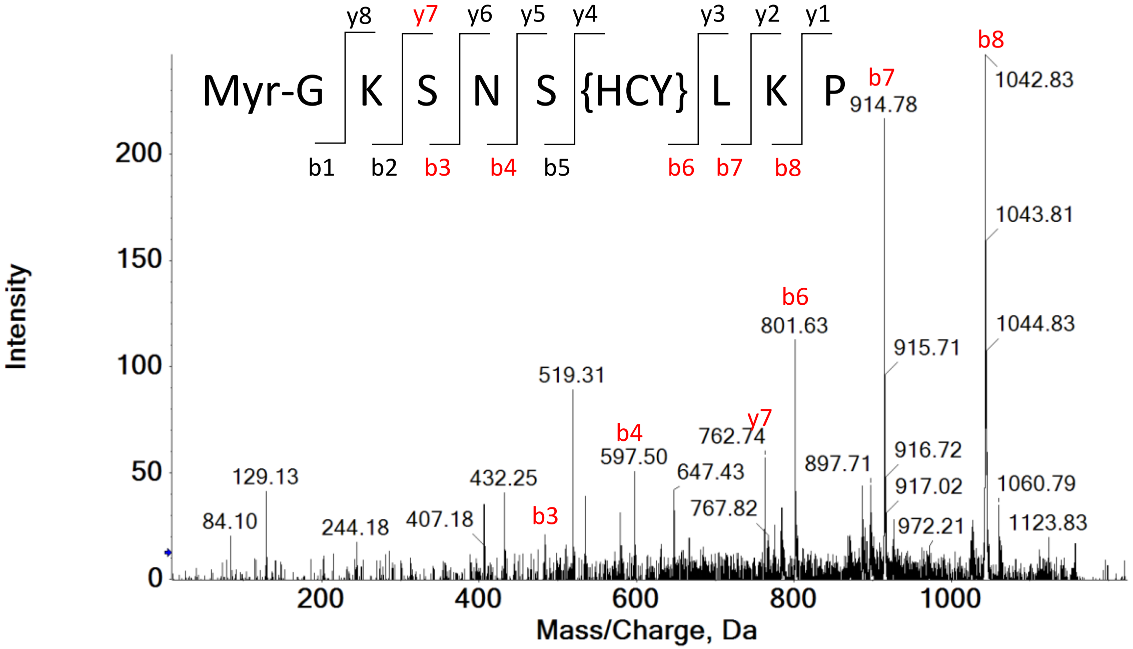


| H6_MSMS_1157.6500_2 | Mass/Charge (Da) | Intensity | |
|---|---|---|---|
| 820,00 | 406,15 | 35,29 | SNSK(-11) |
| 947,00 | 432,24 | 40,94 | NSK(-11)L |
| 1128,00 | 483,31 | 20,86 | $b_3$ |
| 1393,00 | 519,31 | 89,10 | SNSK(-11)L |
| 1490,00 | 534,45 | 39,06 | KSNSK(-11) |
| 1882,00 | 597,49 | 50,67 | $b_4$ |
| 2269,00 | 647,39 | 42,20 | KSNSK(-11)L |
| 2270,00 | 647,43 | 34,82 | SNSK(-11)LK |
| 3429,00 | 762,73 | 57,25 | $y_7$ |
| 3602,00 | 775,53 | 25,57 | KSNSK(-11)LK |
| 3744,00 | 784,48 | 33,88 | $b_6$-$NH_3$ |
| 4004,00 | 801,62 | 112,63 | $b_6$ |
| 4985,00 | 886,75 | 43,92 | $a_7$ |
| 5180,00 | 897,72 | 44,24 | $b_7$-$NH_3$ |
| 5489,00 | 914,78 | 216,63 | $b_7$ |
| 7034,00 | 1042,81 | 246,75 | $b_8$ |

MSMS T3 Myr-KSNS{HCY}LKP (1100)

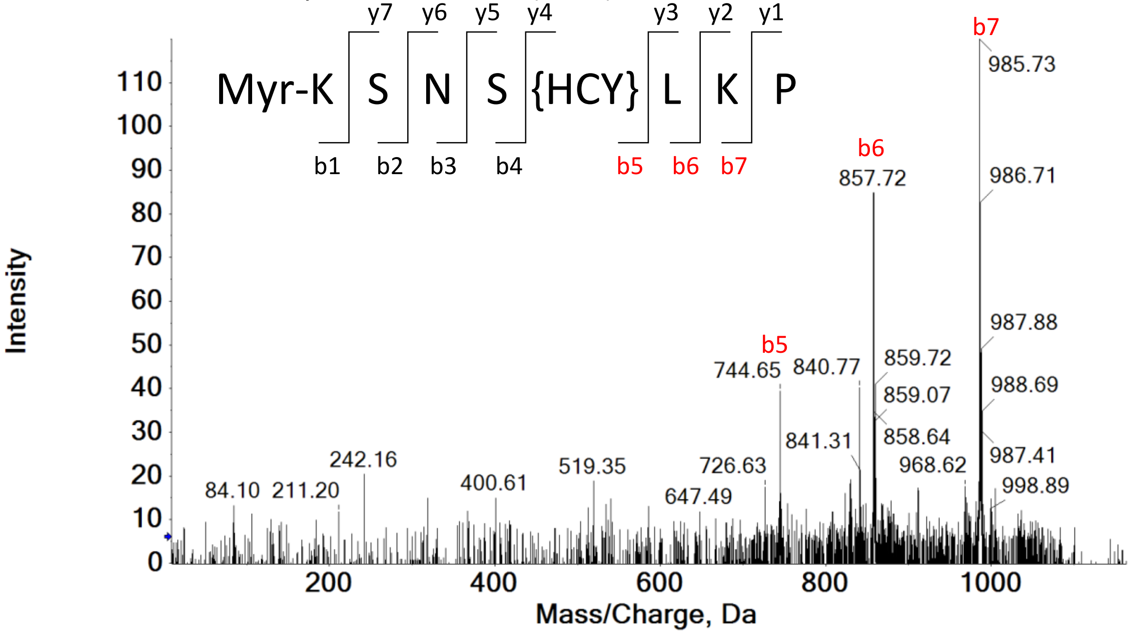


| H7_MSMS_1100.6000_2 | Mass/Charge (Da) | Intensity | |
|---|---|---|---|
| 1983 | 726,6255 | 17,41177 | b5-H2O |
| 2148 | 744,6686 | 39,37255 | $b_5$ |
| 2803 | 840,7583 | 40,15686 | b6-NH3 |
| 2999 | 857,7784 | 84,86275 | b6 |
| 4161 | 985,7468 | 120 | b7 |

### GKSNS{bhLYS}LKP

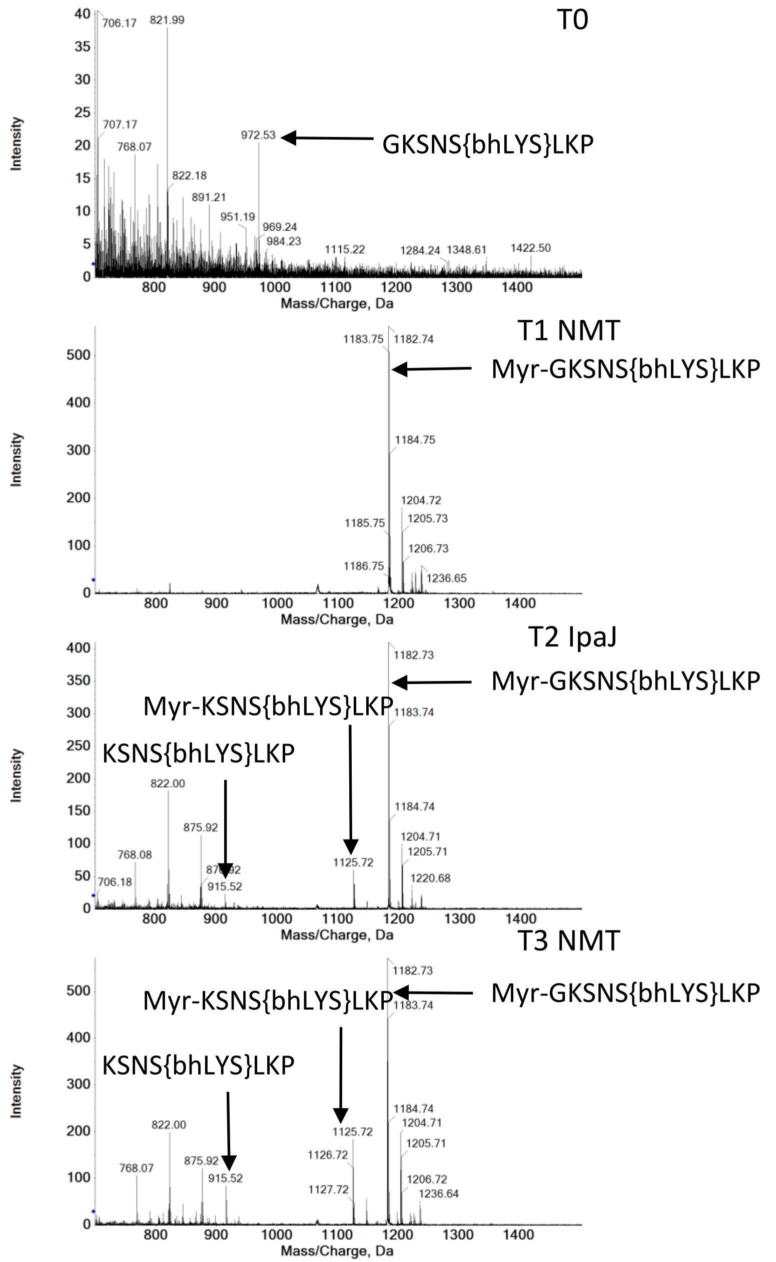


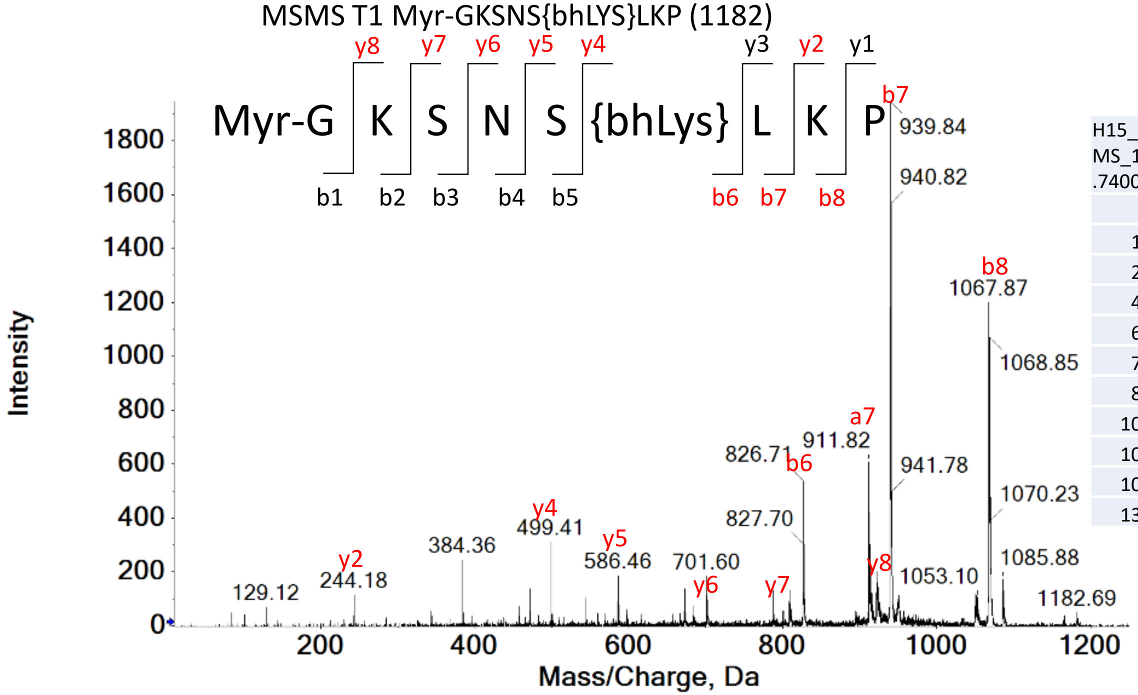


| H15_MS MS_1182 .7400_2 | Mass/Charge (Da) | Intensity | |
|---|---|---|---|
| 403 | 244,1983 | 114,8235 | y2 |
| 1275 | 384,3645 | 245,3333 | K(+14)LK |
| 2792 | 499,3974 | 311,2157 | y4 |
| 4033 | 586,4408 | 186,6667 | y5 |
| 6142 | 700,519 | 112 | y6 |
| 7786 | 787,5328 | 133,0196 | y7 |
| 8616 | 826,6849 | 538,5098 | b6 |
| 10328 | 911,7731 | 609,4118 | a7 |
| 10398 | 915,1217 | 114,5098 | y8 |
| 10912 | 939,9027 | 1945,255 | b7 |
| 13351 | 1067,865 | 1199,216 | b8 |

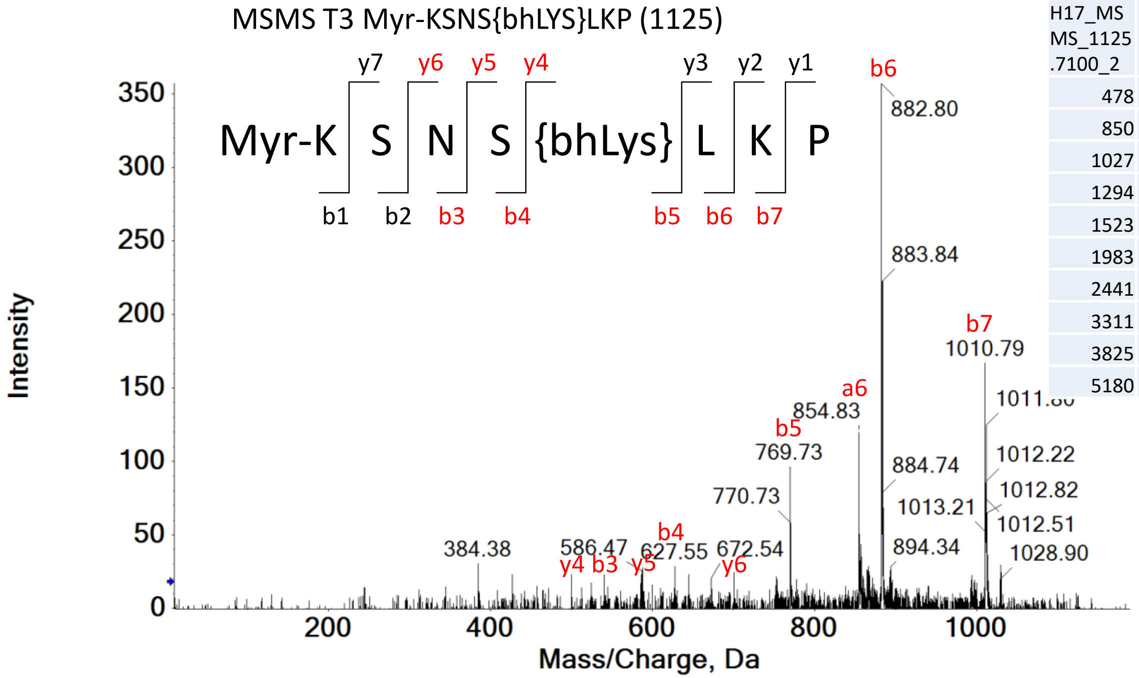


| H17_MS MS_1125 .7100_2 | Mass/Charge (Da) | Intensity | |
|---|---|---|---|
| 478 | 384,3568 | 30,58824 | K(+14)LK |
| 850 | 499,3685 | 22,90196 | y4 |
| 1027 | 540,4617 | 23,21569 | b3 |
| 1294 | 586,4889 | 25,72549 | y5 |
| 1523 | 627,5412 | 28,54902 | b4 |
| 1983 | 700,5056 | 24,47059 | y6 |
| 2441 | 769,5708 | 70,58824 | b5 |
| 3311 | 854,8636 | 119,8431 | a6 |
| 3825 | 882,8086 | 357,0196 | b6 |
| 5180 | 1010,709 | 166,902 | b7 |

### GKVWSQGV

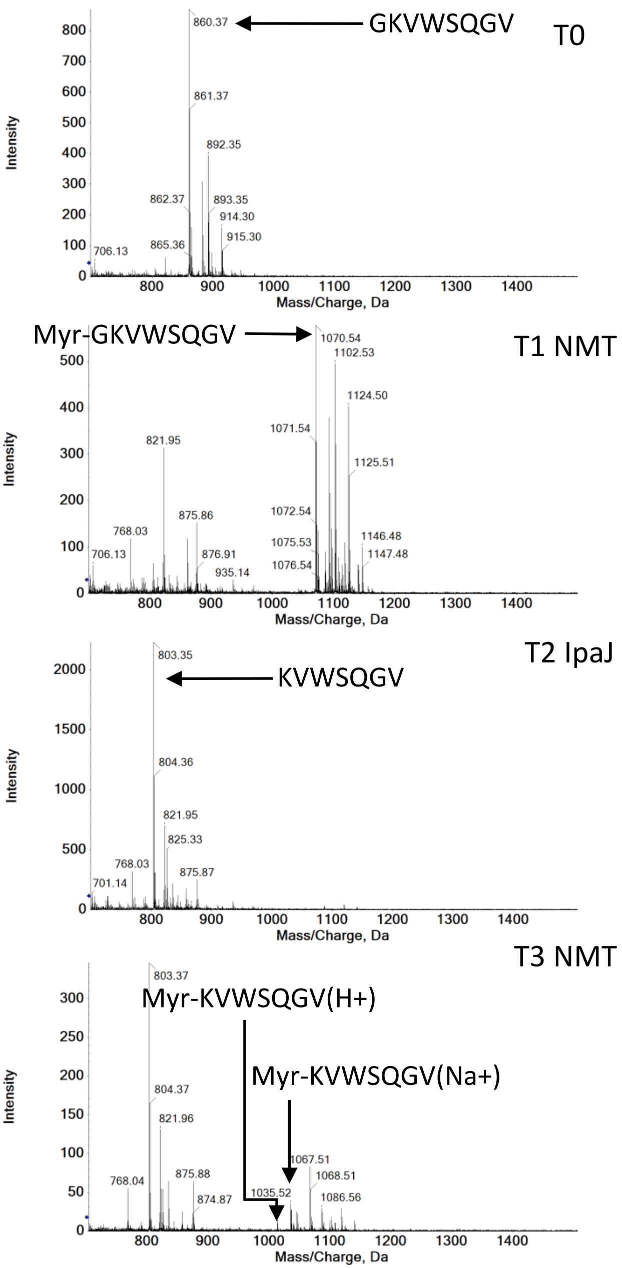


MSMS T1 Myr-GKVWSQGV (1070)

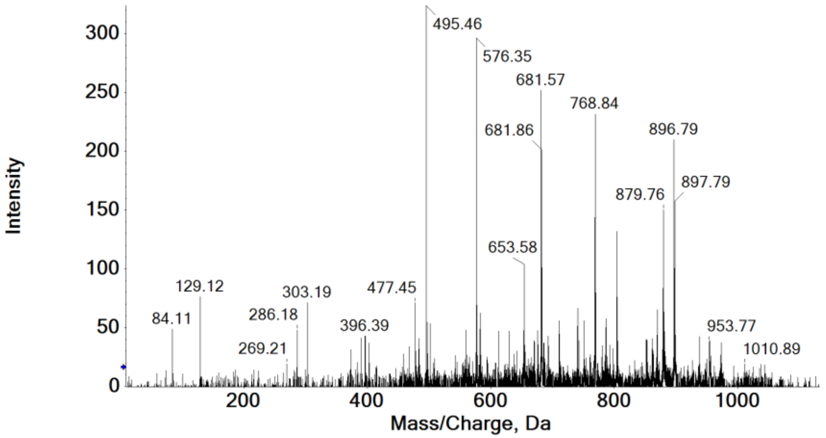


MSMS T2 KVWSQGV (803)

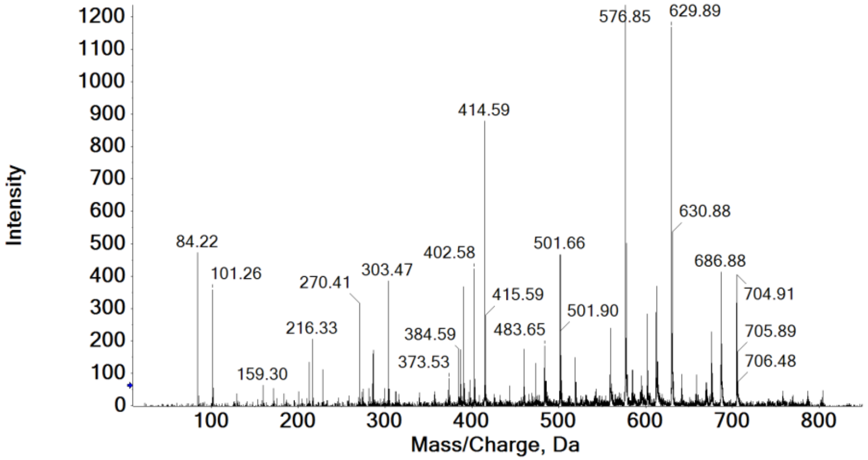


MSMS T2 Myr-KVWSQGV(H+) (1013)

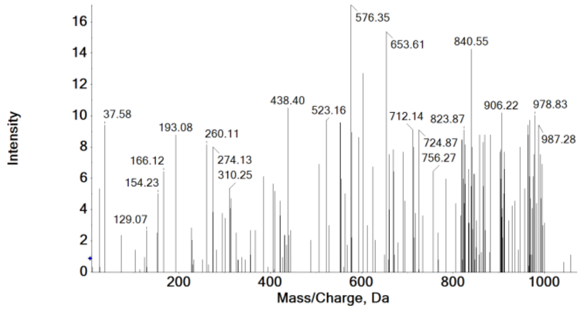


MSMS T2 Myr-KVWSQGV(Na+) (1035)

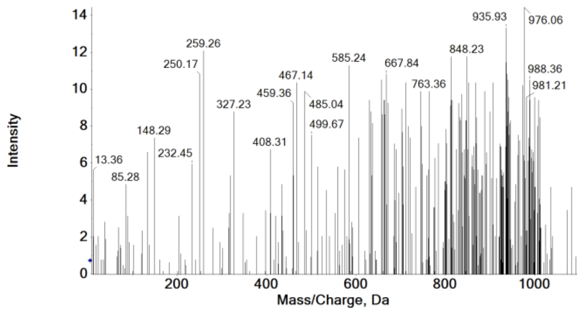

### GKSNAKLKP

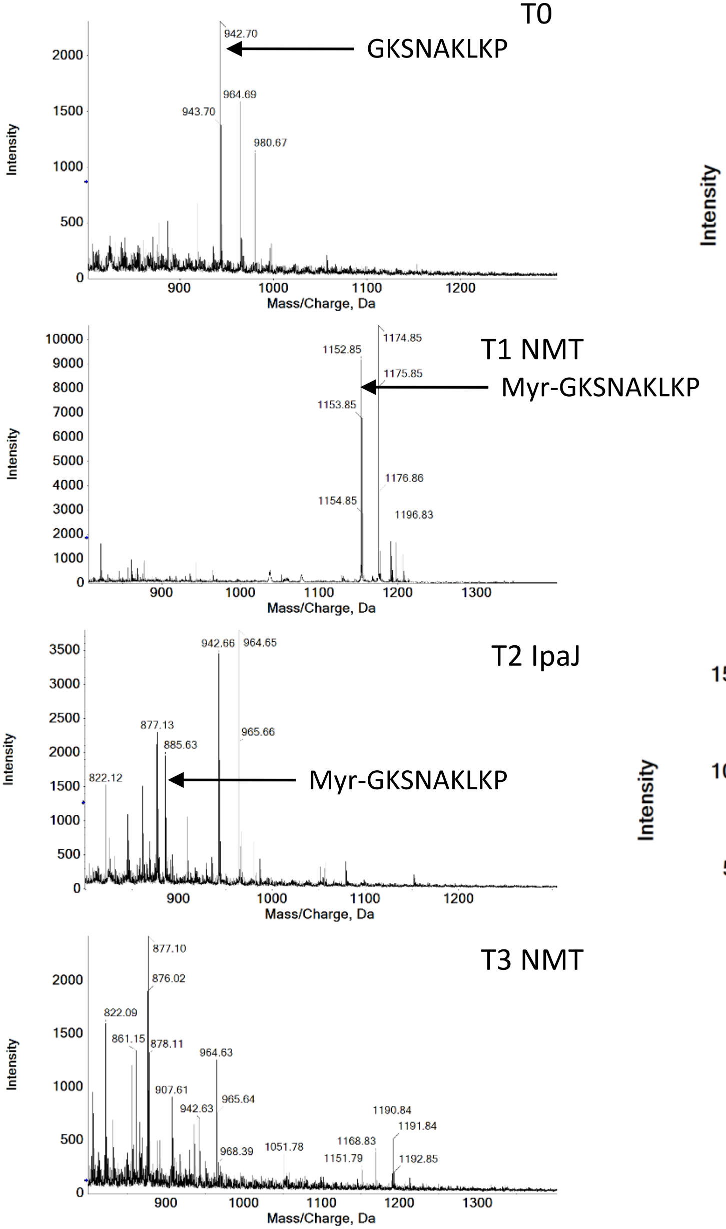


MSMS T1 Myr-GKSNAKLKP (1152)

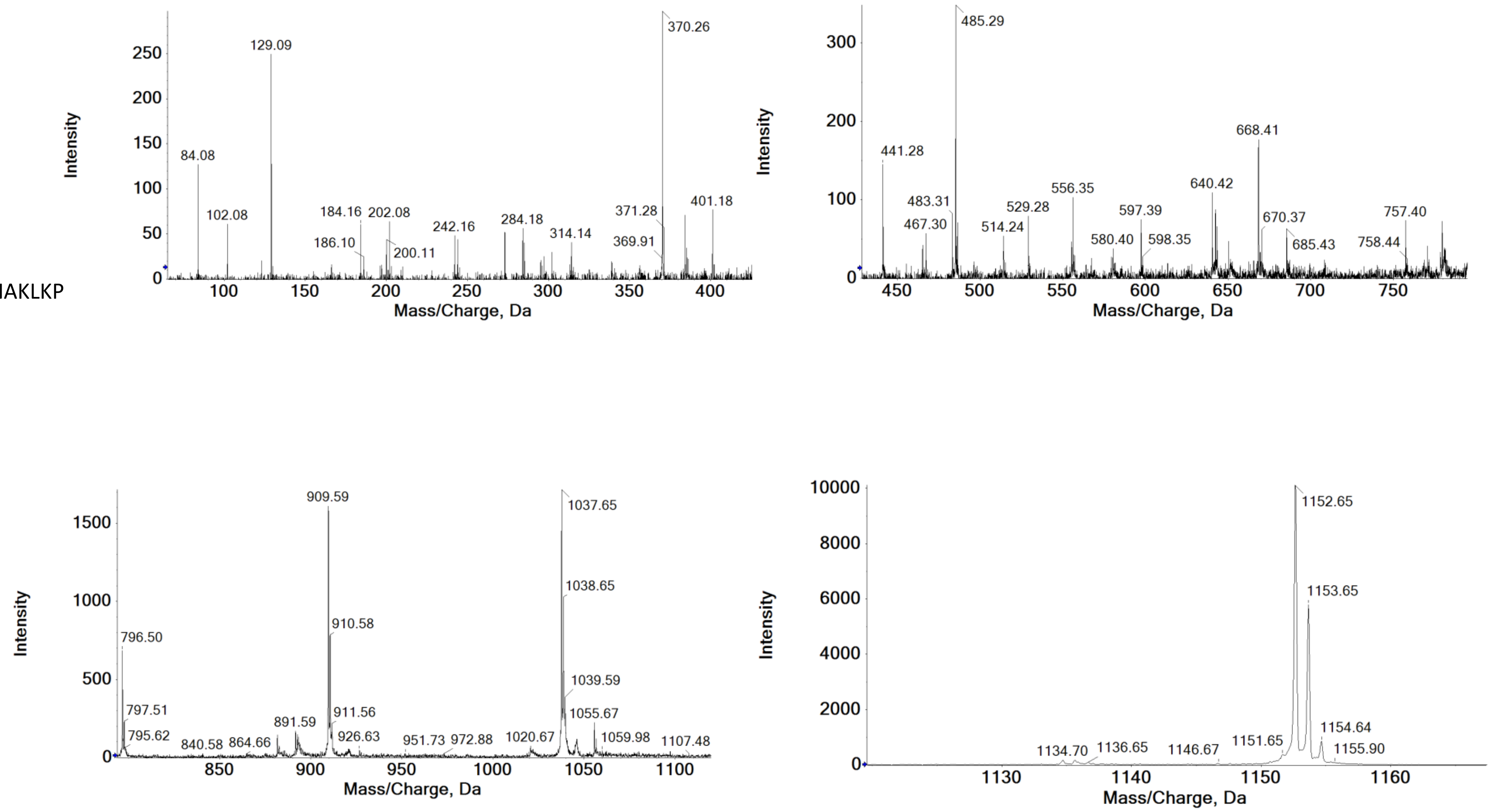
