## Supplementary Dataset 1c for "Proteome-wide probing of the dual NMT-dependent myristoylation tradeoff unveils potent, mechanism-based suicide inhibitors"

### Sup Dataset 1c

| Series [a] | Peptide sequence | Shortname (text & Figures) | Series sequence[d] | MS data page in Supplementary Dataset 1c |
|---|---|---|---|---|
| acGK | ac-GKSFSKPR | acGK | | 2 |
| | ac-G{Orn}SFSKPR | acG[Orn] | | 3 |
| | ac-G{Dab}SFSKPR | acG[Dab] [c] | | 4 |
| | ac-G{D-Orn}SFSKPR | acG[D-Orn] [c] | | 5 |
| | ac-GNCFSKPR | acGN | | 6 |
| | ac-GKVLSKIF | acGK | ARF6 | 7 |
| XK | AKQNSKLR | A-HPCA | HPCA | 8 |
| | AKVLSKIF | A-ARF6 | ARF6 | 9 |
| | AKSFSKPR | AK | | 10 |
| | AKPTSKDSGLK | TCS2 | TCS2 | 11 |
| | A{Orn}SFSKPR | A{Orn} [c] | | 12 |
| | A{D-Orn}SFSKPR | A{D-Orn} [c] | | 13 |
| | SKSFSKPR | SK | | 14 |
| | MKSFSKPR | MK | | 15 |
| | PKSFSKPR | PK | | 16 |
| | G{D-ORN}SFSKPR | G[D-Orn] [c] | | 17 |
| | ac-KSFSKPR | acK | | 18 |
| | ac-{Orn}SFSKPR | ac[Orn] [c] | | 19 |
| | ac-{D-Orn}SFSKPR | ac[D-Orn] [c] | | 20 |
| | ac-{D-Dab}SFSKPR | ac[D-Dab] [c] | | 21 |
| | ac-{Dap}SFSKPR | ac[Dap] | | 22 |
| K | K | K | | 23 |
| | {Orn}SFSKPR | [Orn] | | 24 |
| | {D-Orn}SFSKPR | [D-Orn] [c] | | 25 |
| | {Dab}SFSKPR | [Dab] [c] | | 26 |
| | KQNSKLRP | K-HPCA | HPCA | 27 |
| | ac-KFSKPR | acKΔS | | 28 |
| | KSFS{Orn}PR | | | 29 |
| | KSFS{Dab}PR | | | 30 |
| | KSFS{Dap}PR | | | 31 |
| | ac-[D-Orn}SNSKLK | | NCS1 | 32 |
| | KSNAKLKP | | NCS1 | 33 |
| ZXGK | GGKSFSKP | GGK | | 34/35 |
| | AGKSFSKPR | AGK | | 36 |
| | ac-GGKSFSKPR | acGGK | | 37 |
| | ac-GGGKSFSKPR | acGGGK | | 38 |
| | ac-GGGGKSFSKPR | | | 39 |
| | ac-GGGGGKSFSKPR | | | 40 |

### Ac-GKSFSKPR

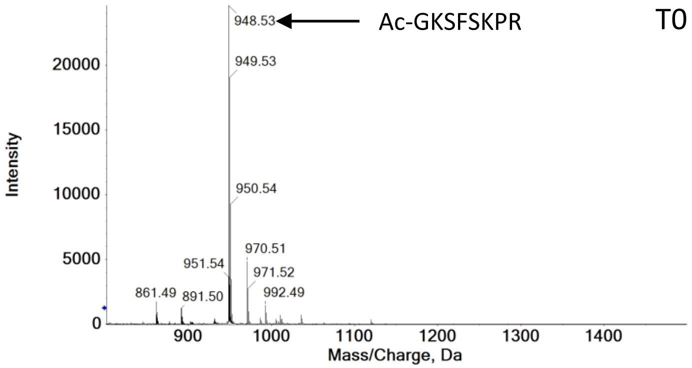


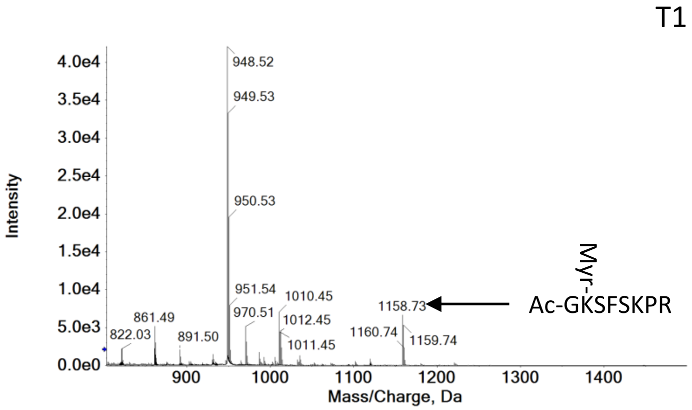


MSMS T0 Ac-GKSFSKPR (948)

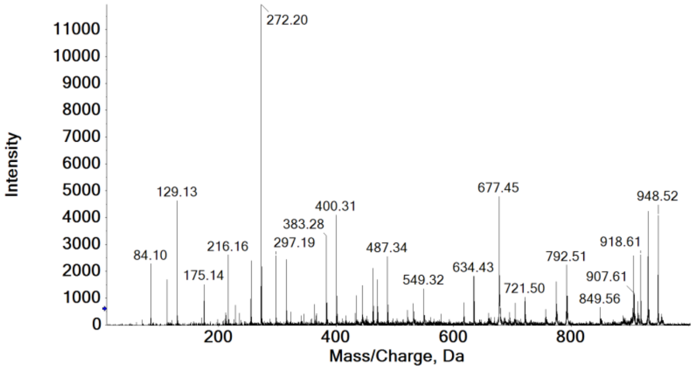


MSMS T1 Ac-GK(Myr)SFSKPR (1158)

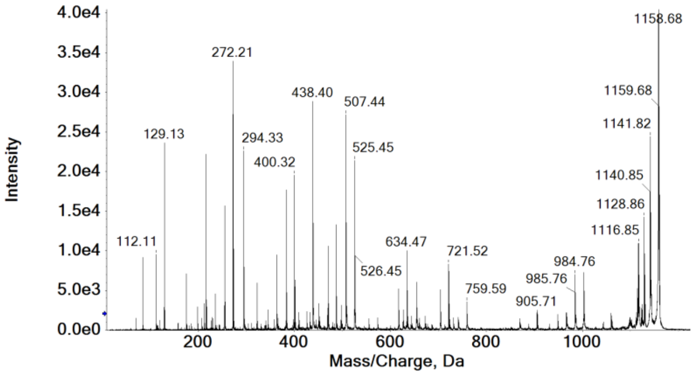


MSMS T1 Ac-GK(Myr)SFSKPR (1158) (2)

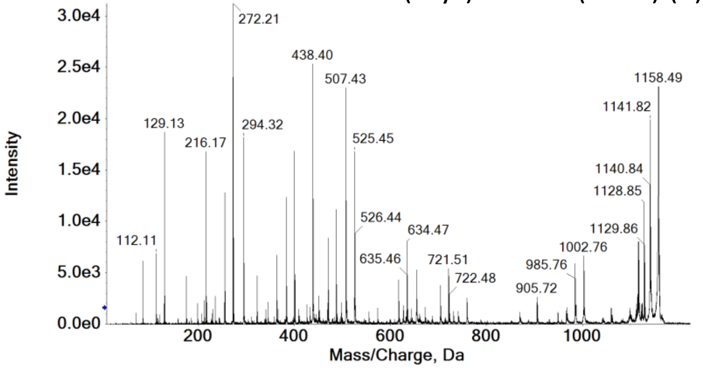


MSMS T0 Ac-GKSFSKPR (948)

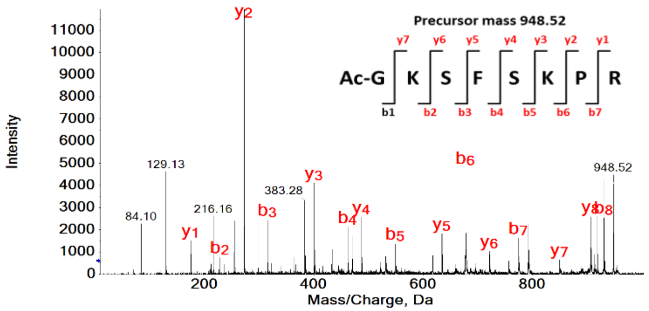


MSMS T1 Ac-GK(Myr)SFSKPR (1158)

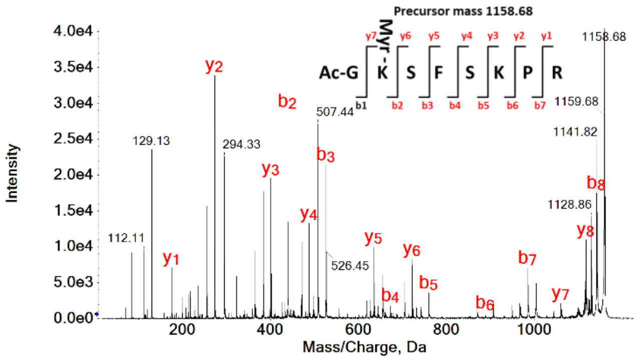

### Ac-G{ORN}SFSKPR

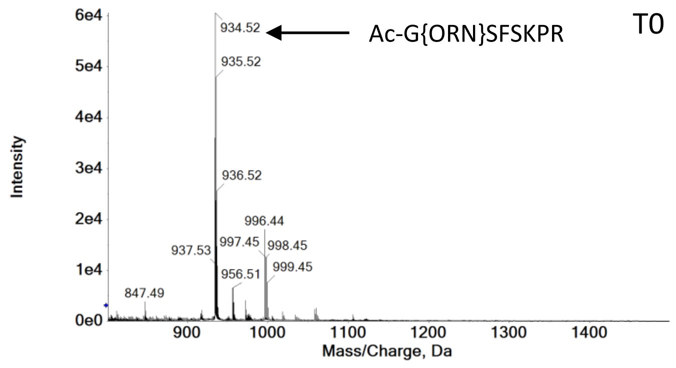


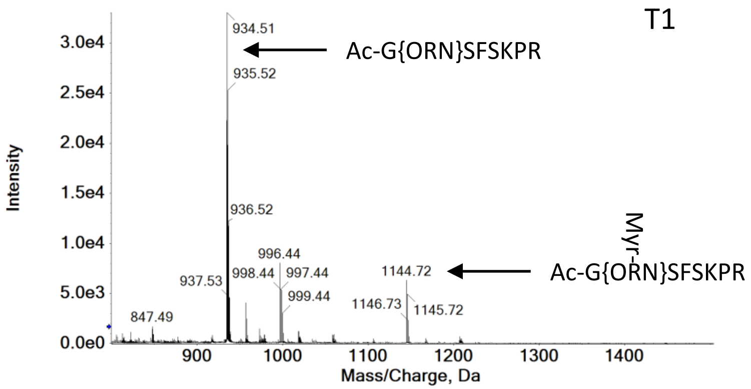

### Ac-G{DAB}SFSKPR

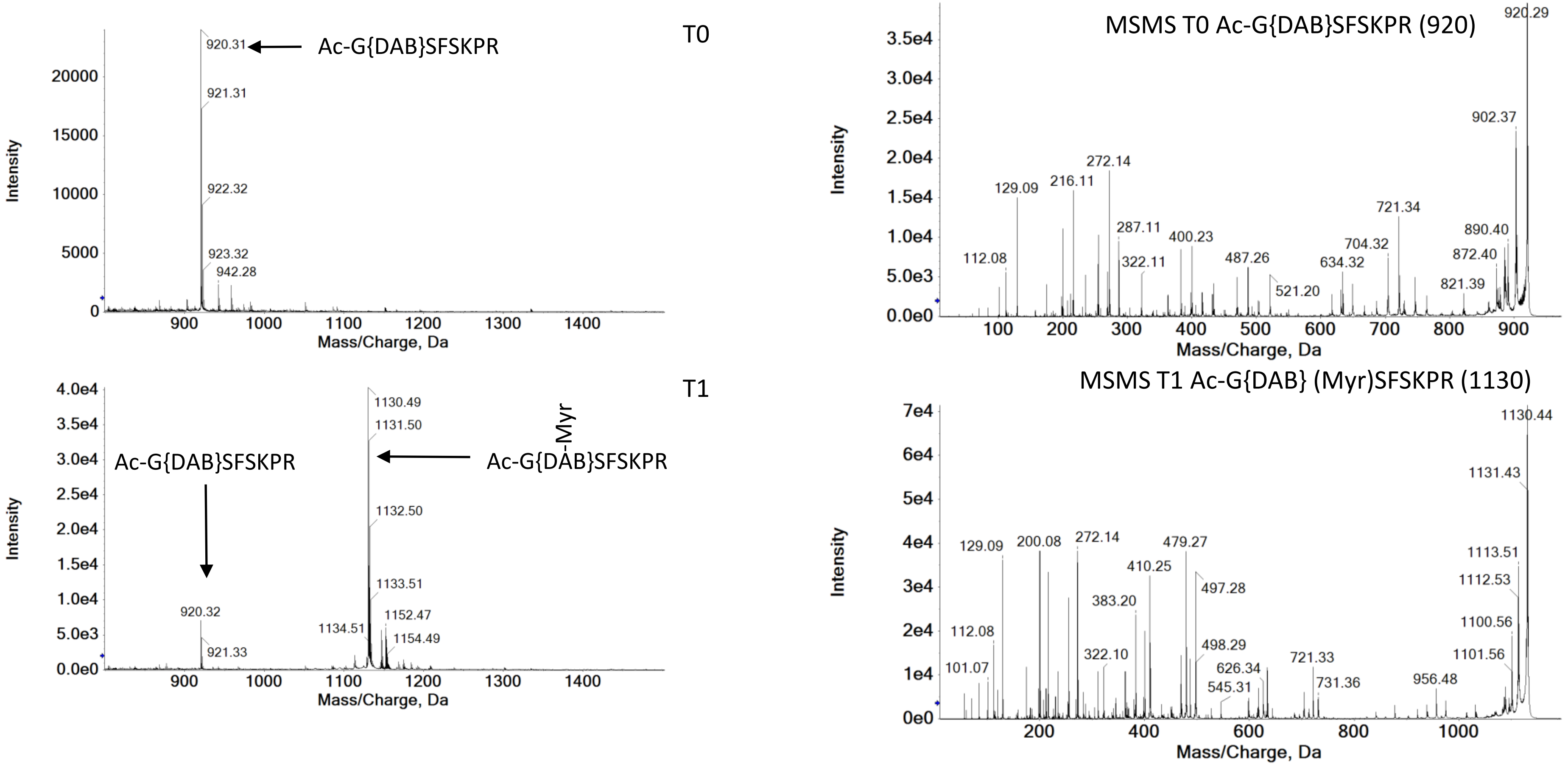

### Ac-G{D-ORN}SFSKPR

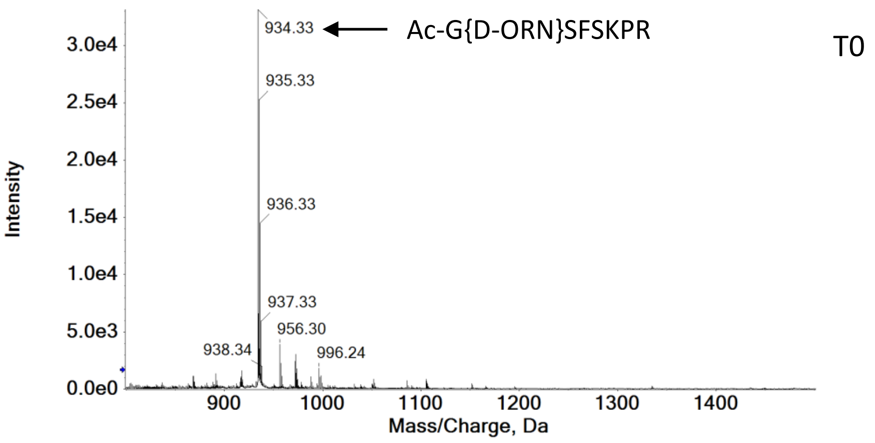


MSMS T0 Ac-G{D-ORN}SFSKPR (934)

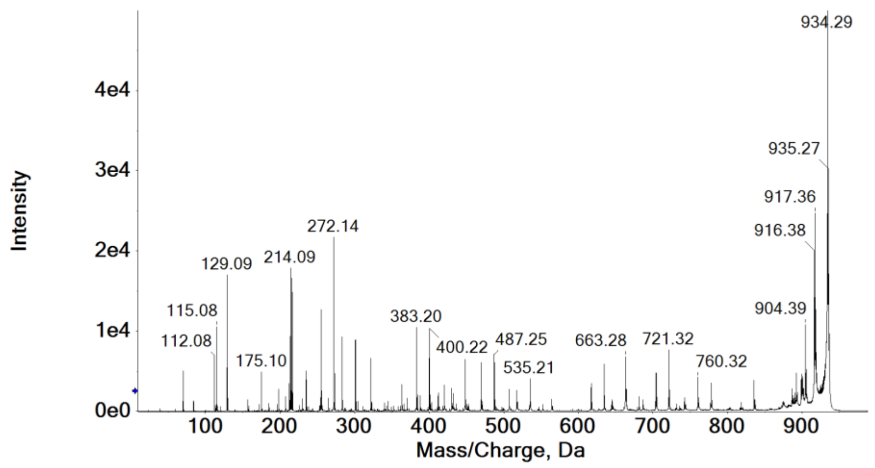


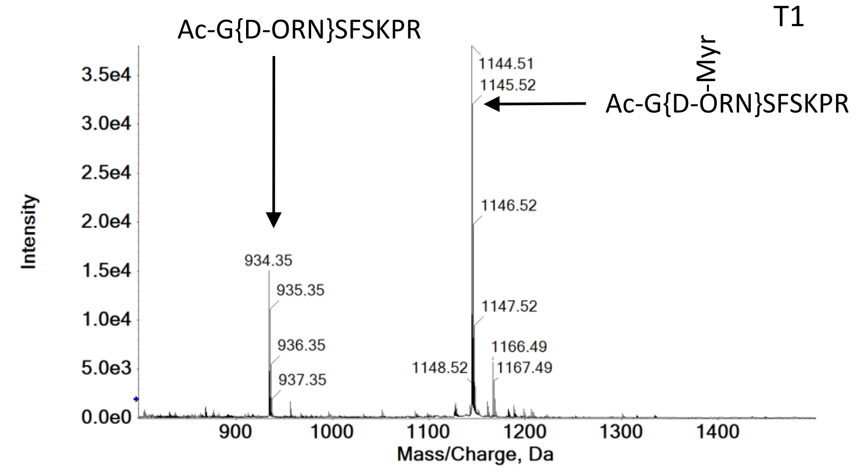


MSMS T1 Ac-G{D-ORN}(Myr)SFSKPR (1144)

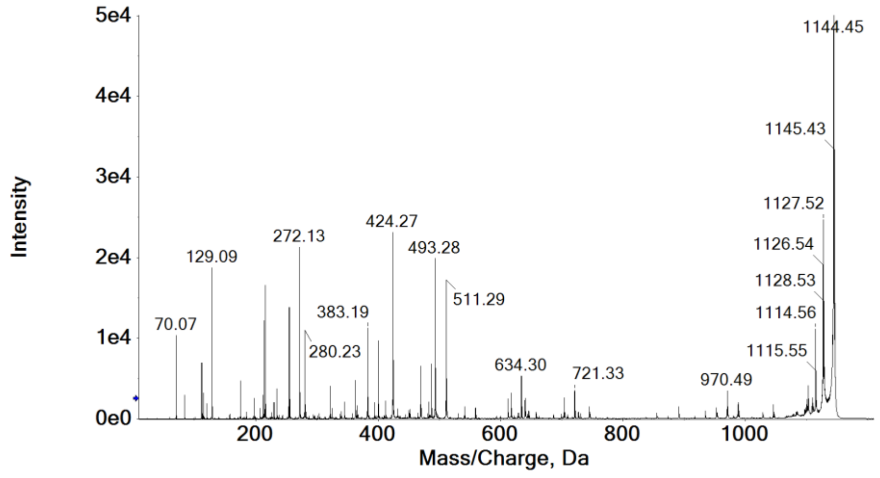

### Ac-GNCFSKPR

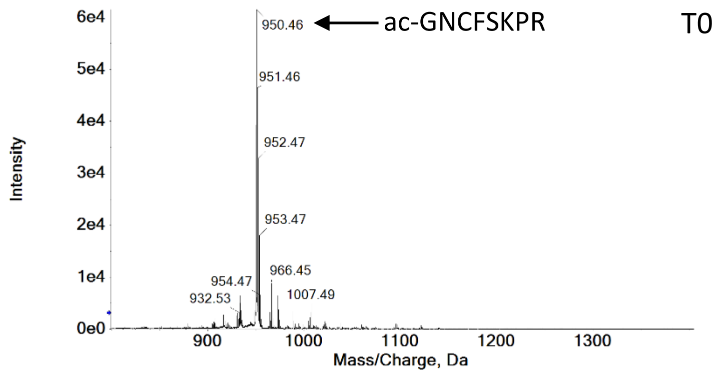

T0
ac-GNCFSKPR
950.46
951.46
952.47
953.47
966.45
954.47
932.53
1007.49
6e4
5e4
4e4
3e4
2e4
1e4
0e0
Intensity
900
1000
1100
1200
1300
Mass/Charge, Da


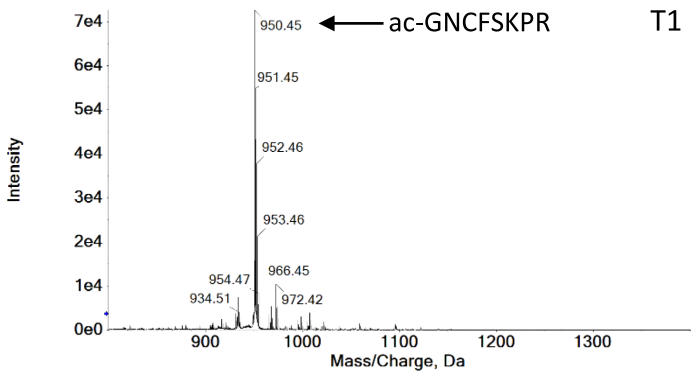

T1
ac-GNCFSKPR
950.45
951.45
952.46
953.46
966.45
954.47
934.51
972.42
7e4
6e4
5e4
4e4
3e4
2e4
1e4
0e0
Intensity
900
1000
1100
1200
1300
Mass/Charge, Da

### Ac-GKVLSKIF

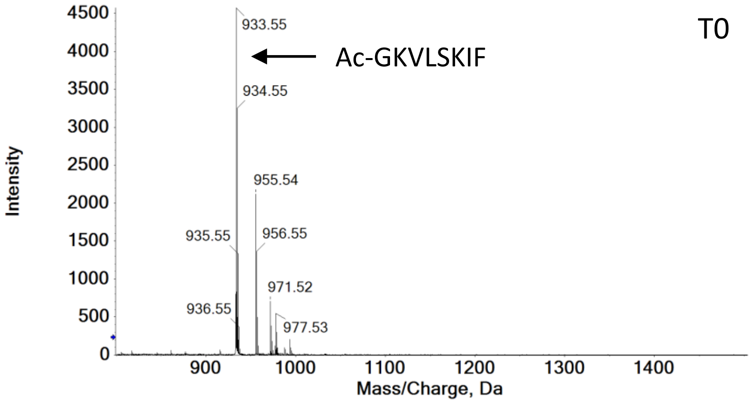


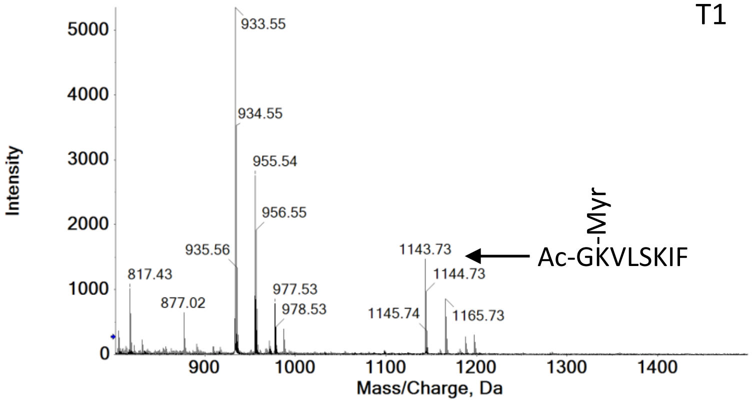


MSMS T1 ac-GK(Myr)VLSKIF (1143)

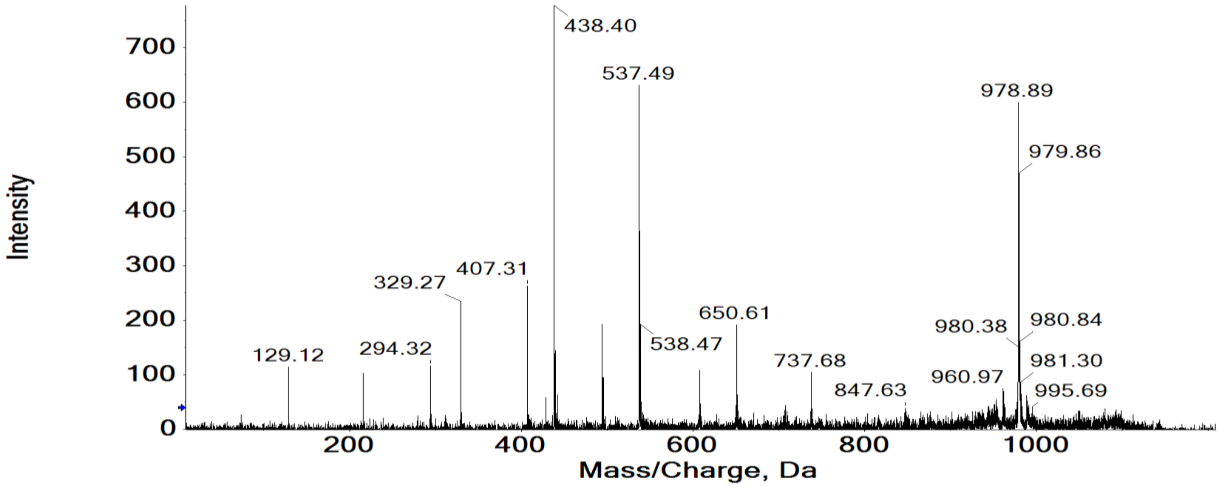


MSMS T1 ac-GK(Myr)VLSKIF (1143) (2)

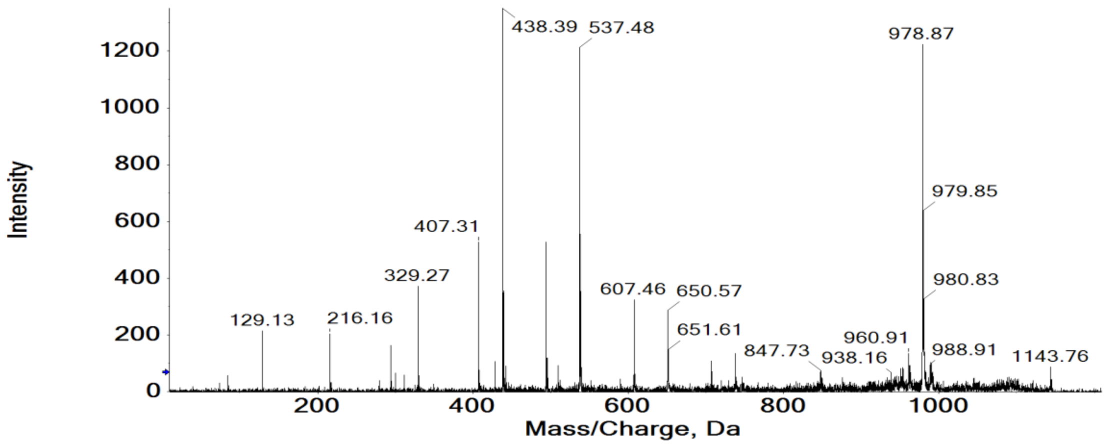

### AKQNSKLR

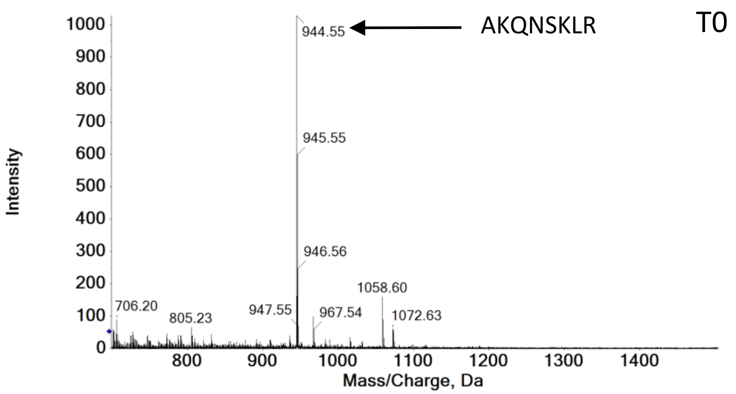


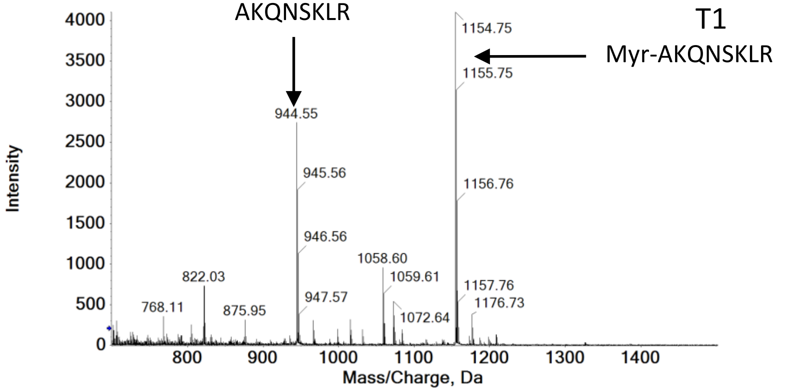


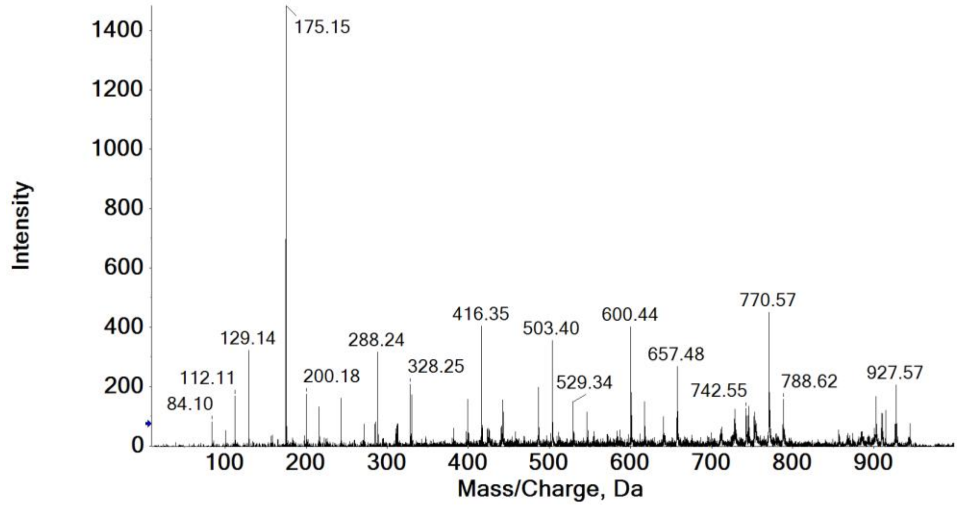


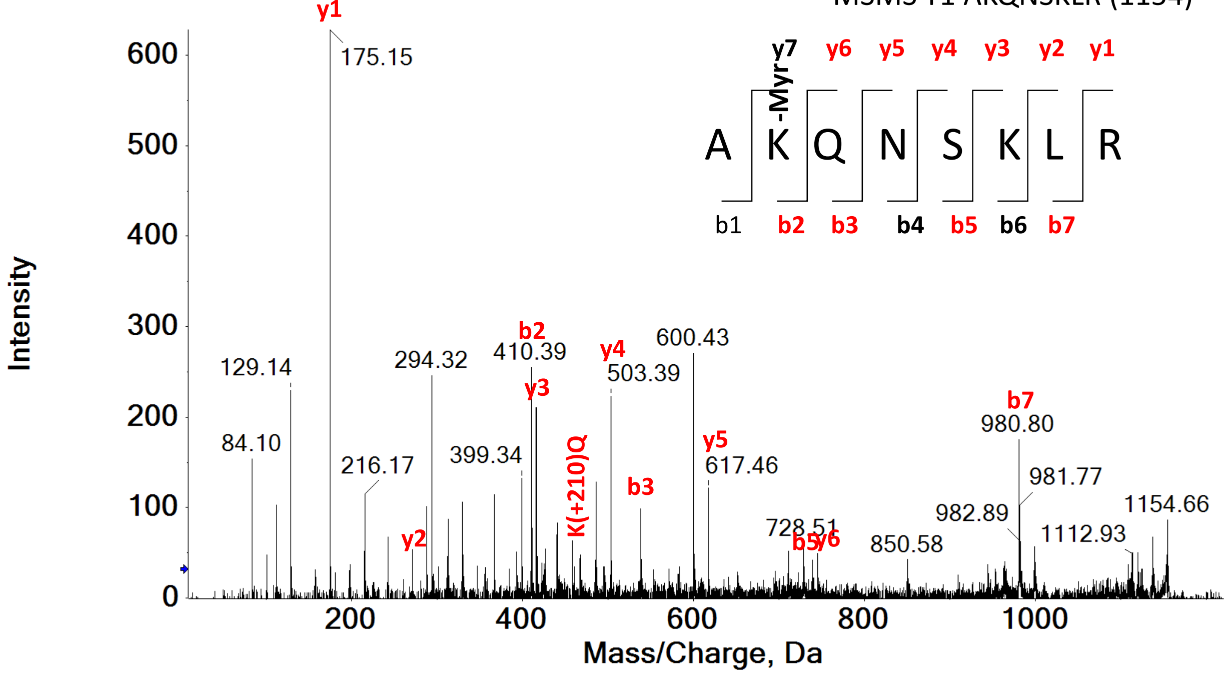


| K24 1154 | Mass/Charge (Da) | Intensity | |
|---|---|---|---|
| 535 | 175,0203 | 35,45098 | y1 |
| 1231 | 288,1639 | 31,52941 | y2 |
| 2914 | 410,3834 | 254,7451 | b2 |
| 3049 | 416,3236 | 210,8235 | y3 |
| 3905 | 467,378 | 43,60785 | K(+210)Q ? |
| 4432 | 503,3777 | 223,3726 | y4 |
| 4875 | 538,4333 | 98,98039 | b3 |
| 5859 | 600,4333 | 249,8824 | y5-NH3 |
| 6146 | 617,4308 | 121,4118 | y5 |
| 8017 | 739,5452 | 42,5098 | b5 |
| 8118 | 745,4847 | 49,09804 | y6 |
| 9226 | 850,5908 | 42,5098 | b6-NH3 |
| 10884 | 980,7767 | 175,5294 | b7 |
| 12877 | 1137,707 | 67,45098 | v8 |

### AKVLSKIF

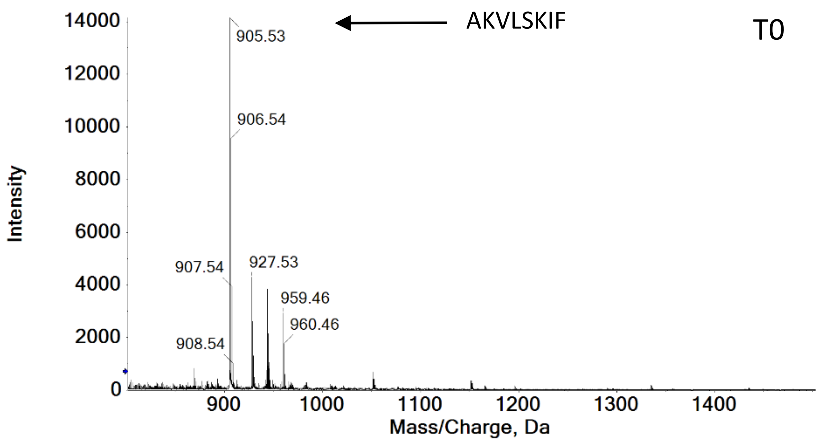


MSMS T0 AKVLSKIF (905)

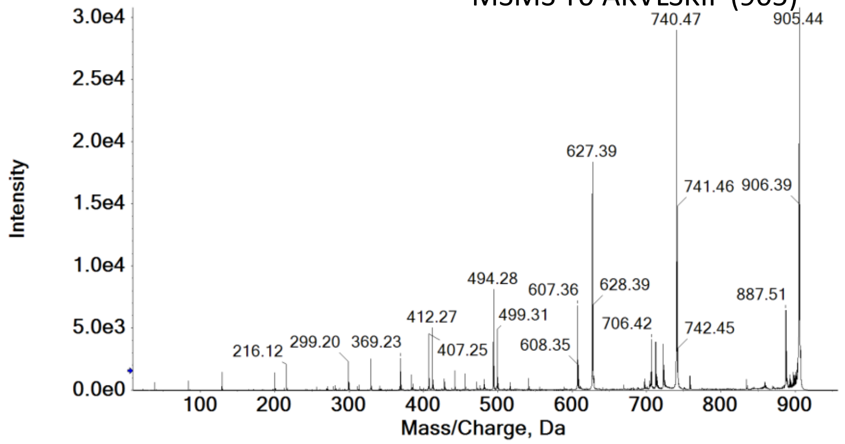


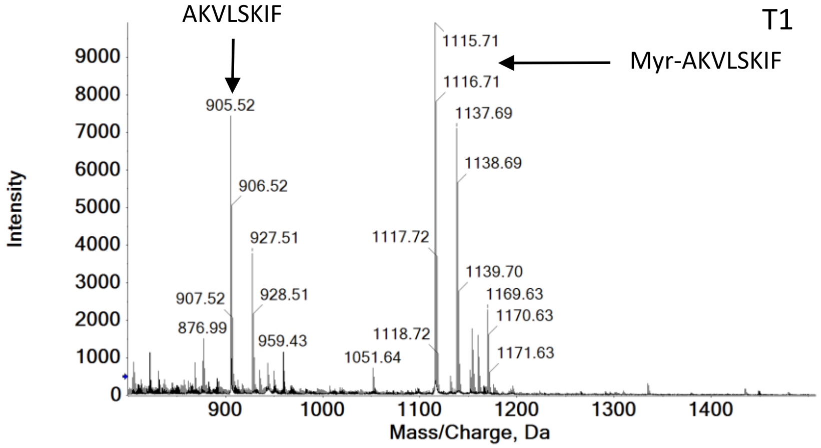


MSMS T1 Myr-AKVLSKIF (1115)

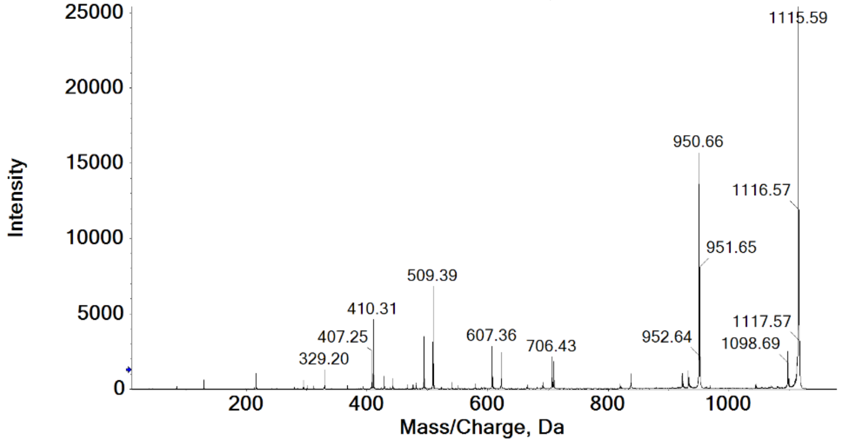

### AKSFSKPR

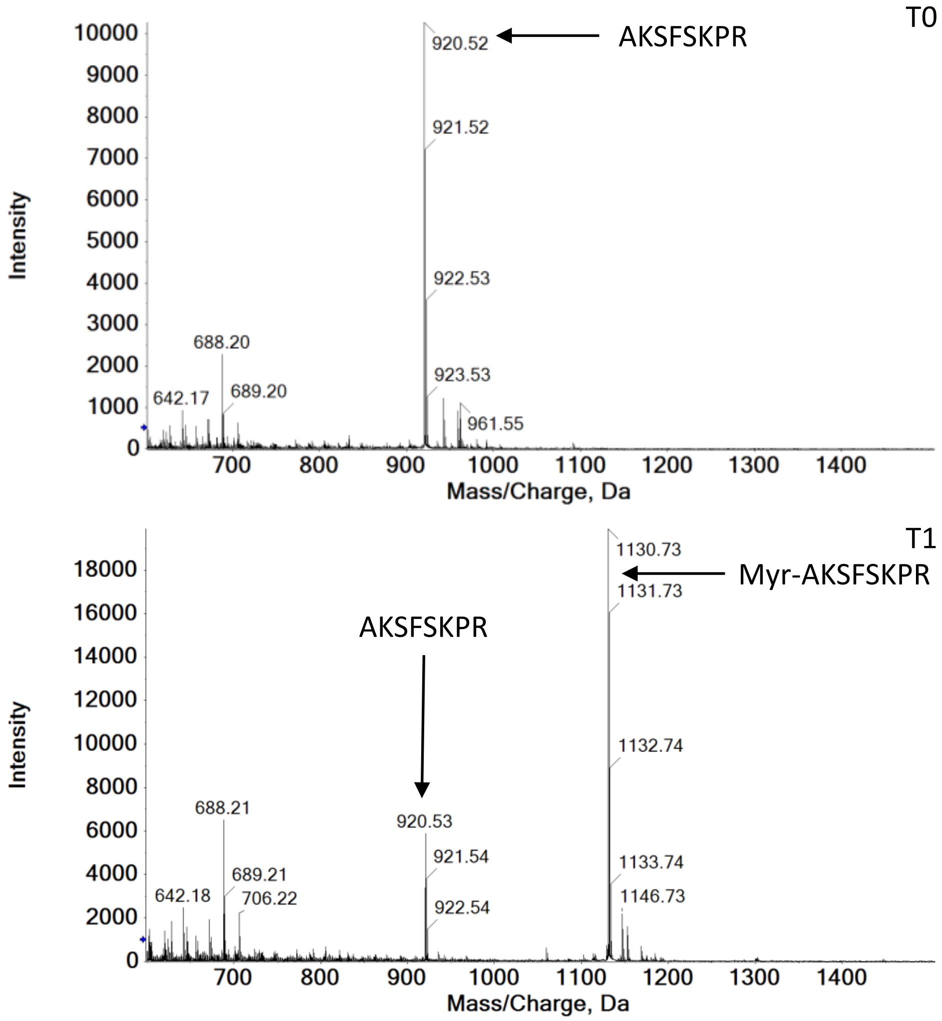


MSMS T1 Myr-AKSFSKPR (1130)

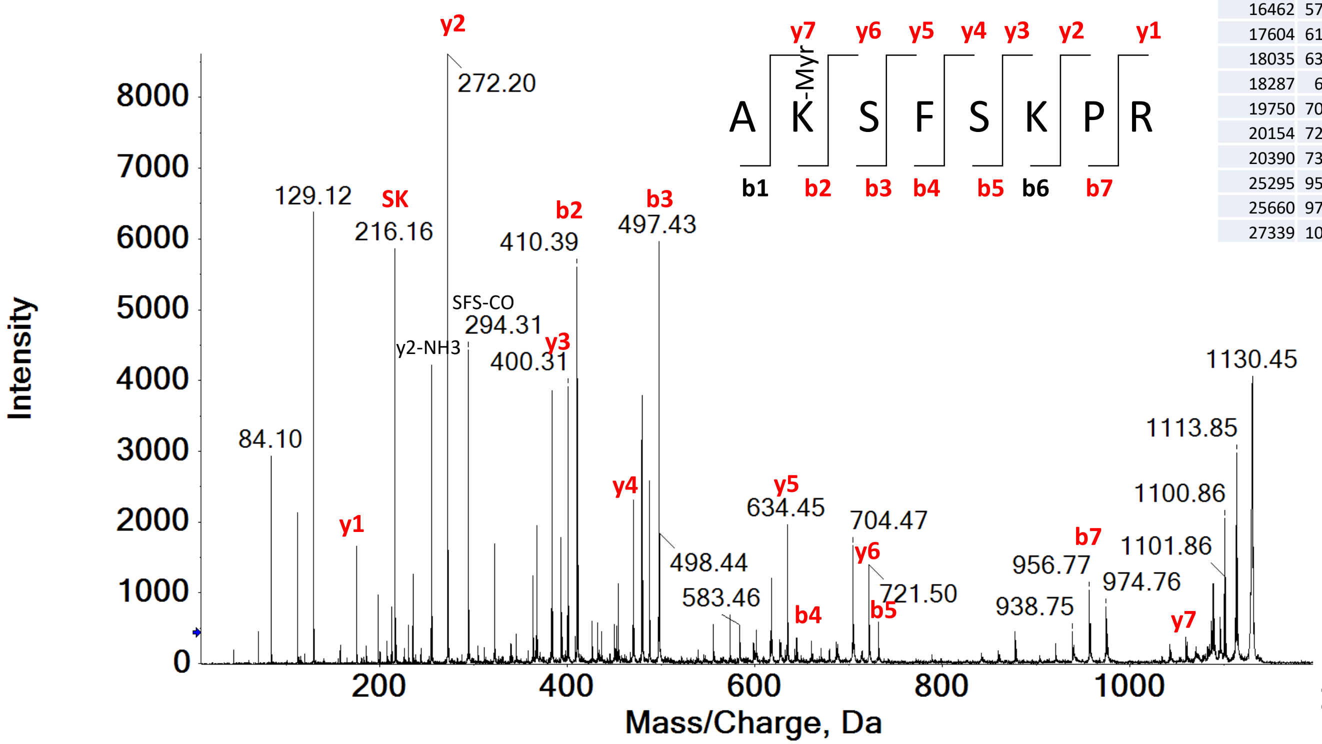


| | | | |
|---|---|---|---|
| 766 | 84,09495 | 2935,84 | K |
| 1481 | 112,1094 | 2126,90 | R |
| 2056 | 129,1225 | 6378,35 | K |
| 3482 | 175,1427 | 1653,96 | y1 |
| 4180 | 198,1471 | 973,80 | SK-H2O/ KP-CO |
| 4900 | 216,147 | 5869,18 | SK |
| 5668 | 235,1309 | 1267,45 | FS |
| 6392 | 255,174 | 4213,96 | y2-NH3 |
| 7039 | 272,2352 | 6608,47 | y2 |
| 7848 | 294,3016 | 4431,22 | SFS-CO |
| 8846 | 322,1745 | 1693,18 | SFS |
| 9640 | 345,2368 | 414,43 | FSK-H2O |
| 10247 | 363,2515 | 1236,71 | FSK |
| 10905 | 383,2669 | 3857,57 | y3-NH3 |
| 11231 | 393,3545 | 1791,69 | b2-NH3 |
| 11453 | 400,3001 | 3915,61 | y3 |
| 11709 | 408,3862 | 384,78 | K(+210)S-H2O |
| 11772 | 410,3888 | 5604,39 | b2 |
| 12270 | 426,3936 | 597,18 | K(+210)S |
| 12996 | 450,2837 | 557,65 | SFSK |
| 13590 | 470,3228 | 2309,80 | y4-NH3 |
| 13855 | 479,4061 | 3797,80 | $b_3$-$H_2O$ |
| 13884 | 480,4055 | 1171,45 | b3-NH3 |
| 14085 | 487,3613 | 2589,18 | y4 |
| 14373 | 497,4166 | 5962,98 | b3 |
| 16462 | 573,4848 | 685,96 | K(+210)SF |
| 17604 | 617,4413 | 1208,78 | y5-NH3 |
| 18035 | 634,4463 | 1964,86 | y5 |
| 18287 | 644,498 | 369,88 | b4 |
| 19750 | 704,4481 | 1667,14 | y6-NH3 |
| 20154 | 721,4829 | 1395,76 | y6 |
| 20390 | 731,5301 | 590,59 | b5 |
| 25295 | 956,7646 | 1045,80 | b7 |
| 25660 | 974,7401 | 803,45 | b7+H2O |
| 27339 | 1059,846 | 375,53 | y7 |

### AKPTSKDSGLK

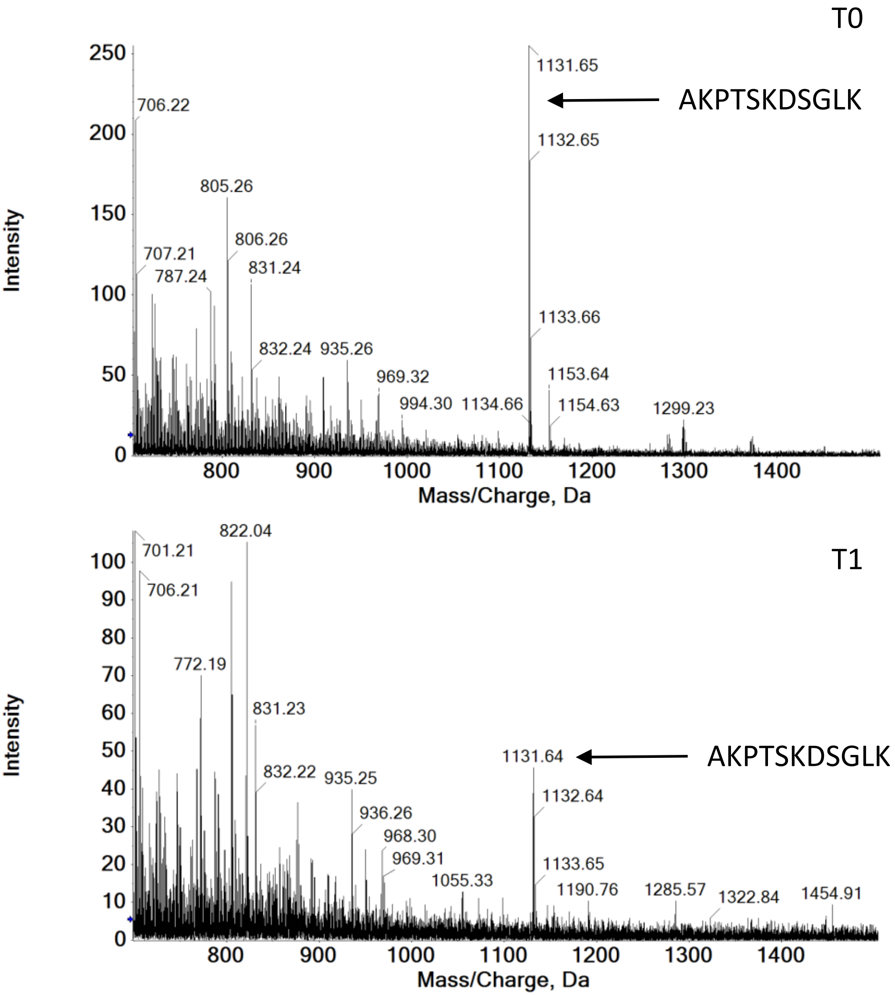

### A{ORN}SFSKPR

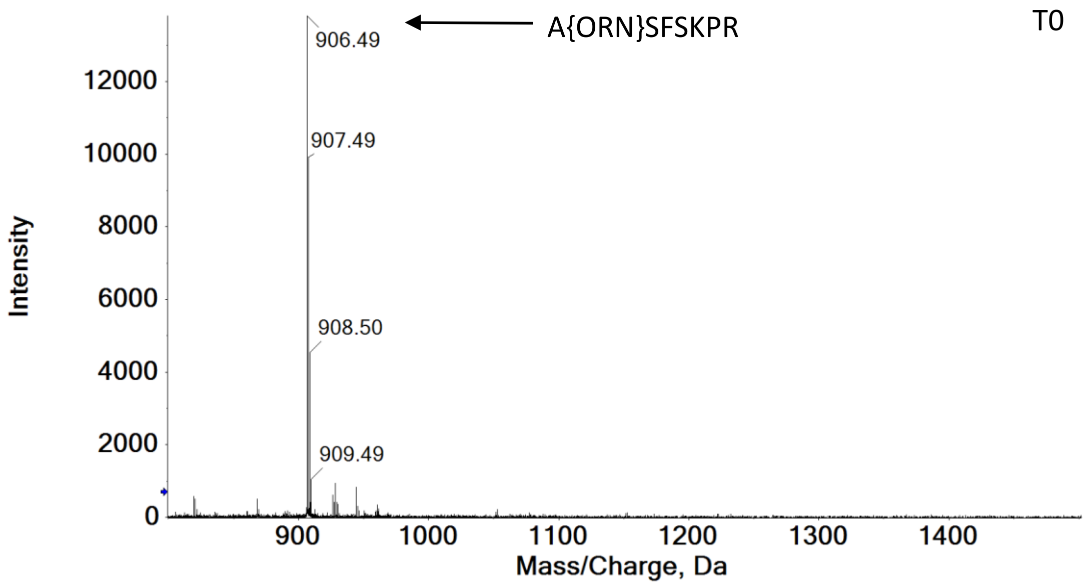


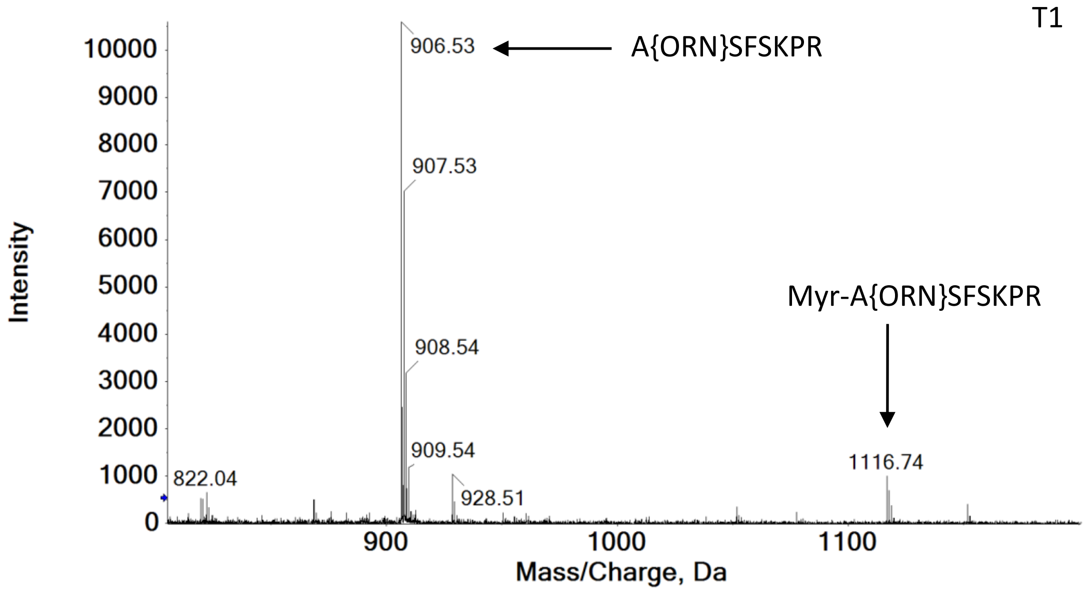


MSMS T1 Myr-A{ORN}SFSKPR (1116)

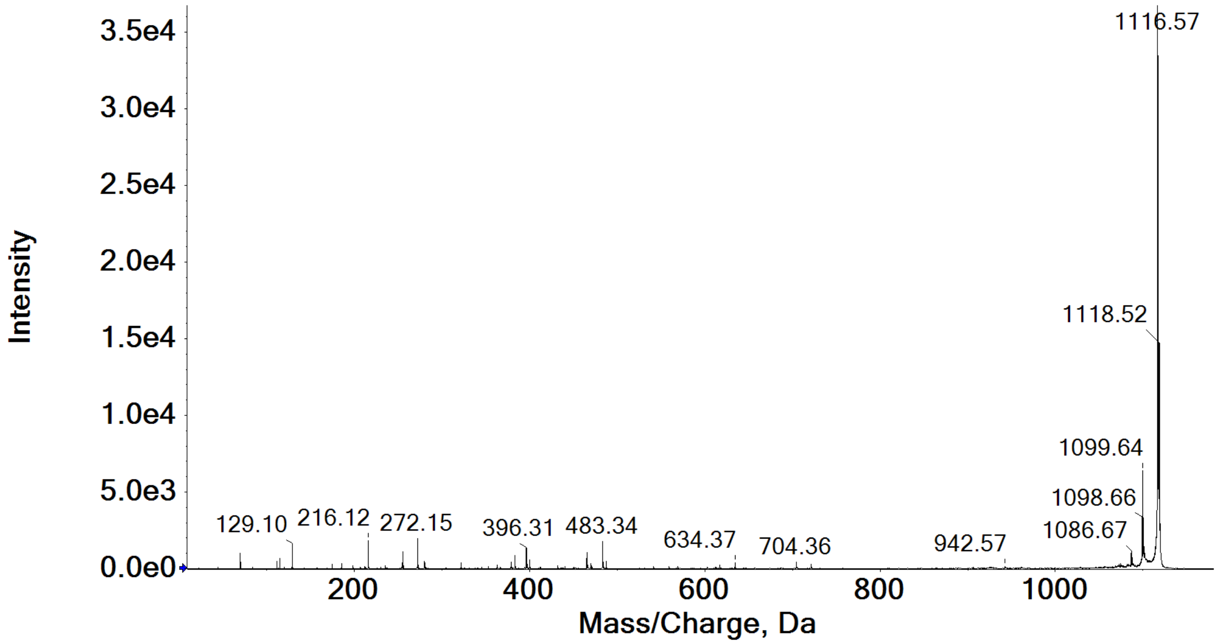

### A{D-ORN}SFSKPR

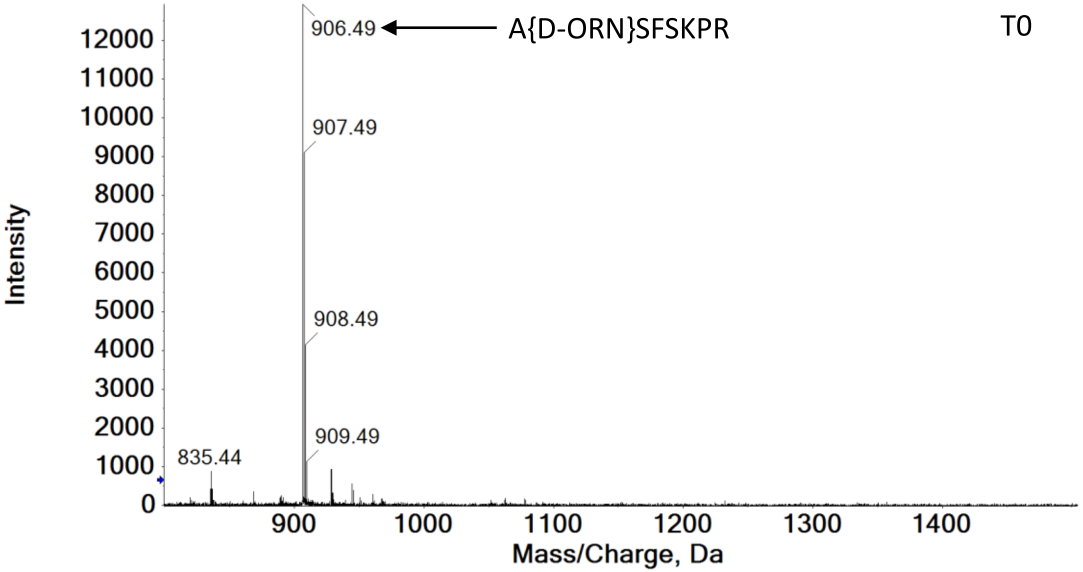


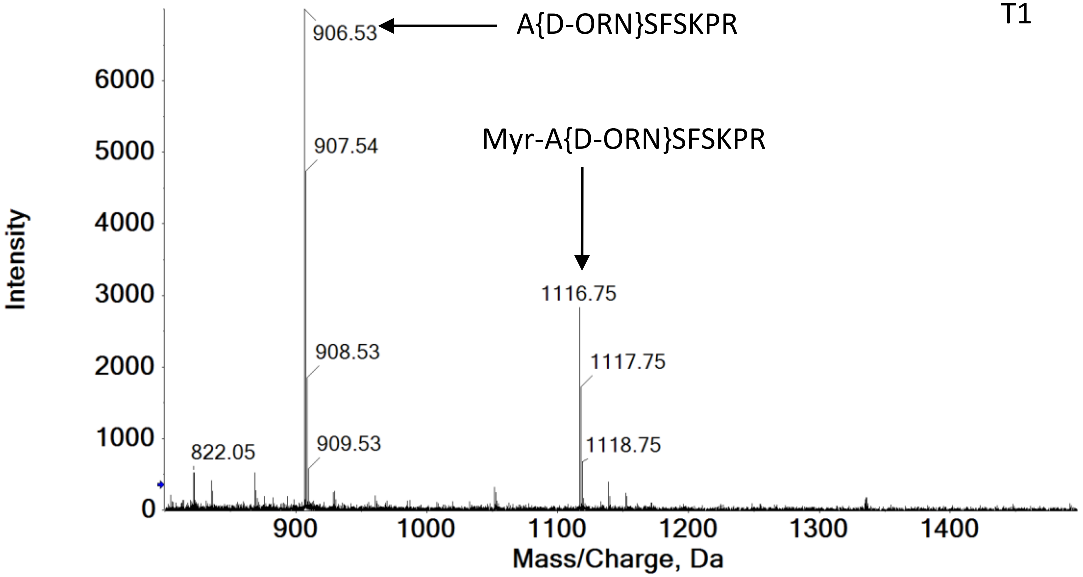


MSMS T1 Myr-A{D-ORN}SFSKPR (1116)

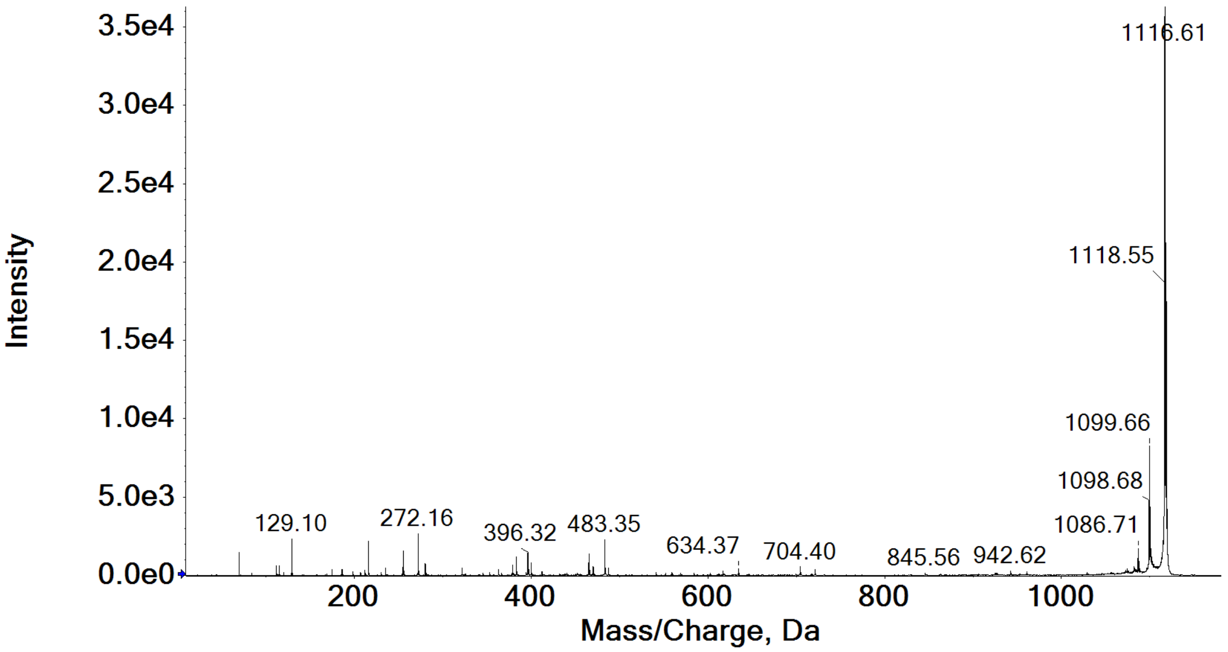

### SKSFSKPR

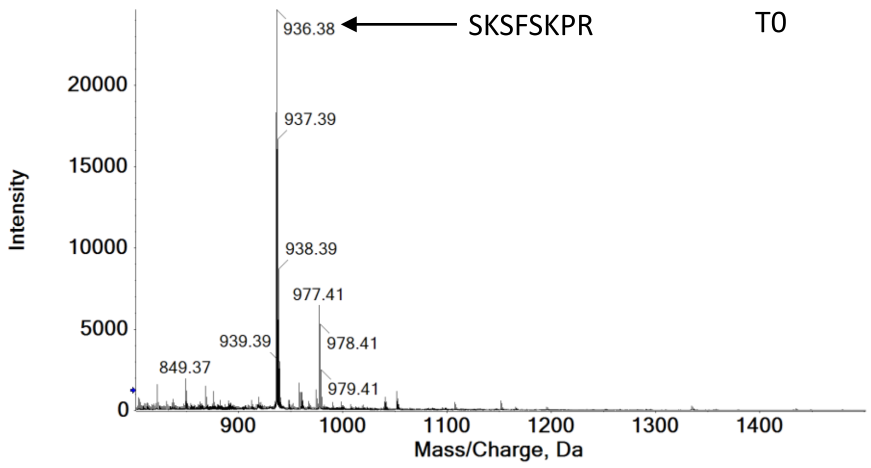


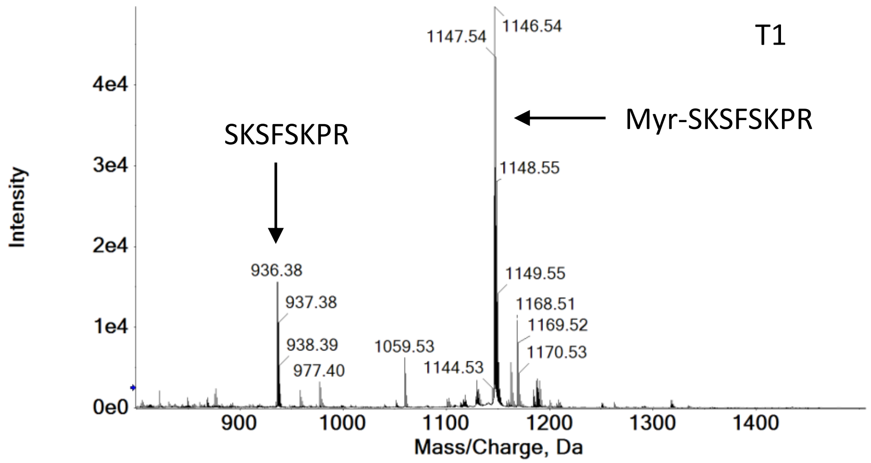


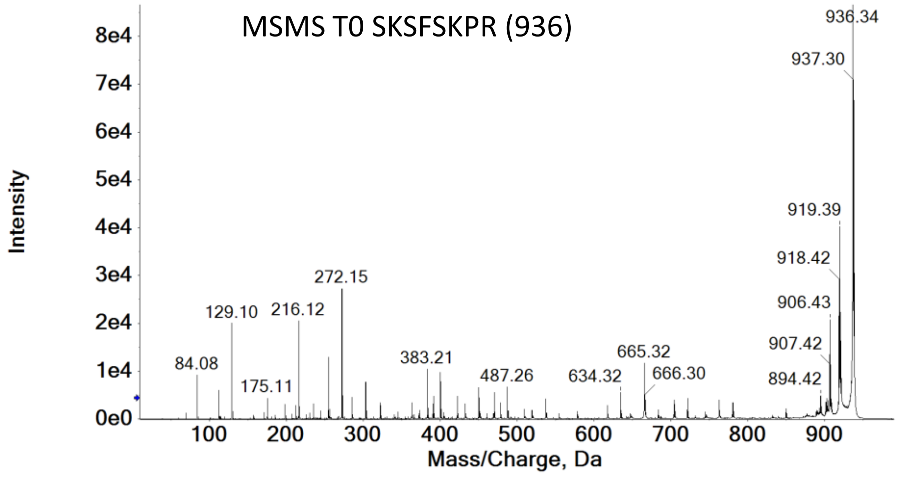


MSMS T1 SK(Myr)SFSKPR (1146)

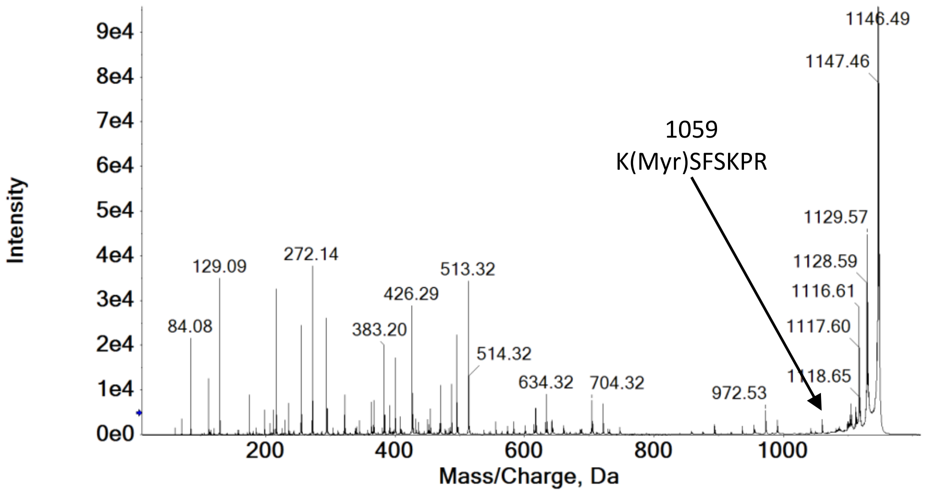

### MKSFSKPR

T0

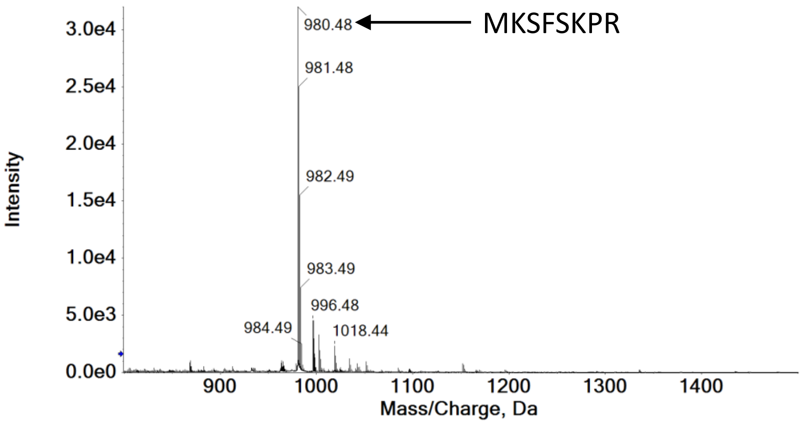


T1

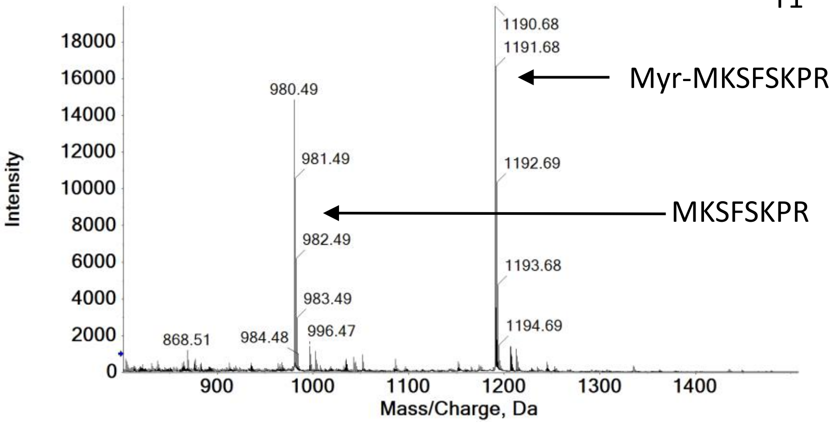


MSMS T0 MKSFSKPR (980)

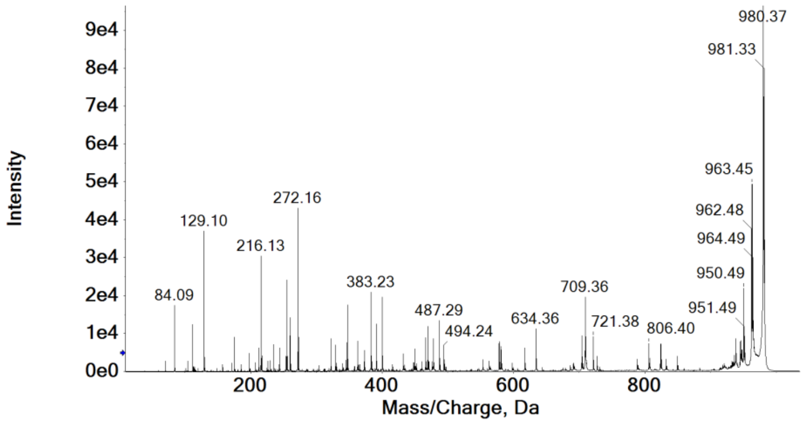


MSMS T1 MK(Myr)SFSKPR (1158)

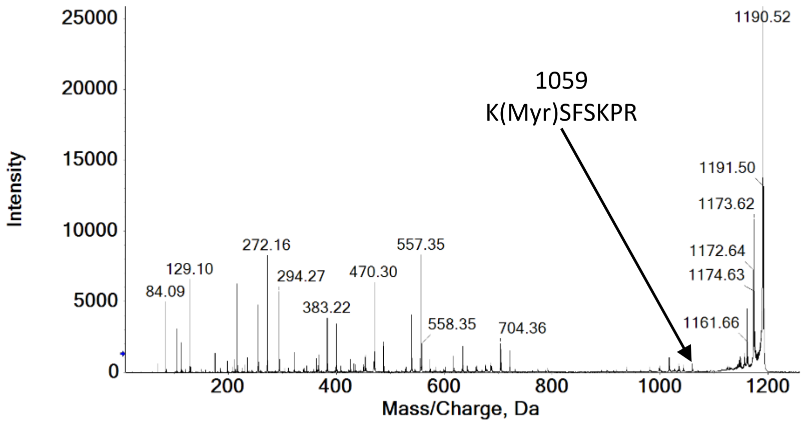

### PKSFSKPR

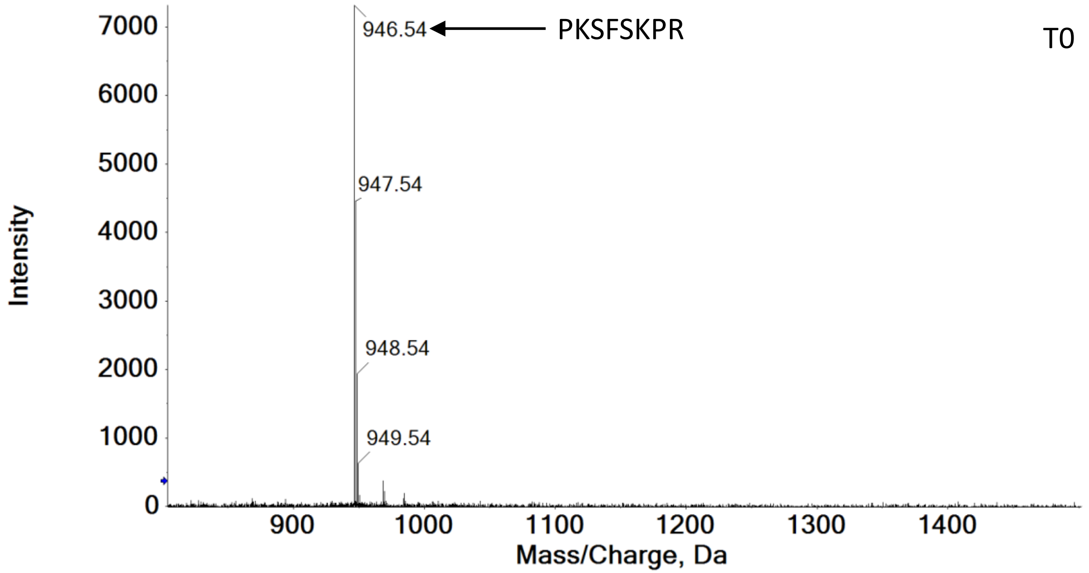


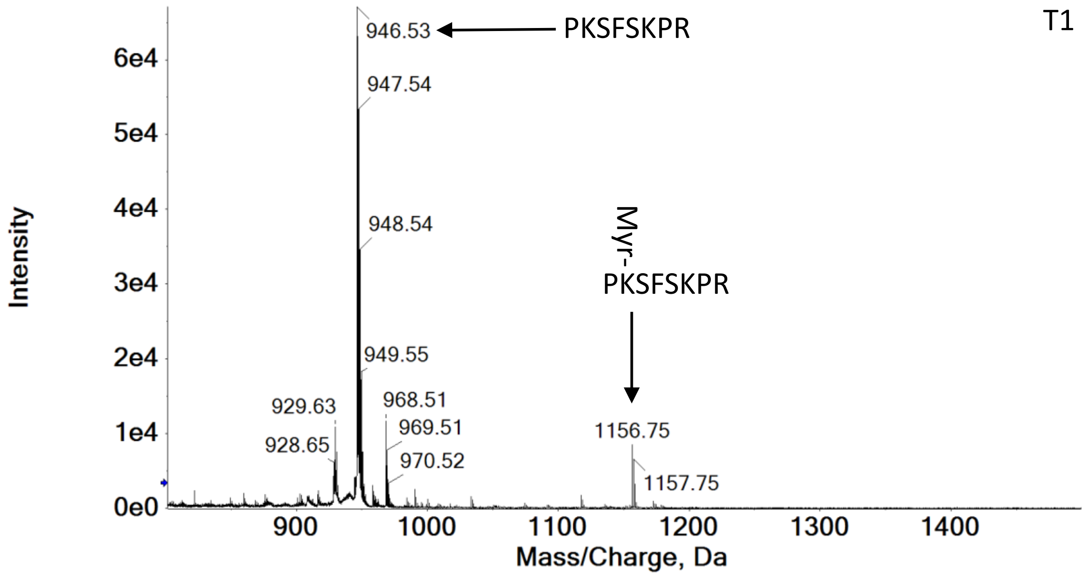


MSMS T1 PK(Myr)SFSKPR (1156)

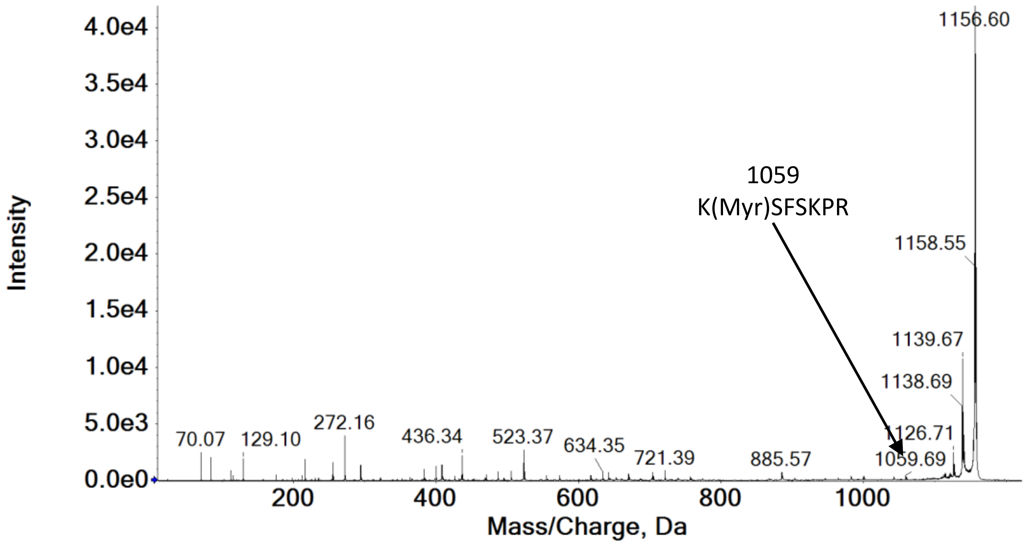

### G{D-ORN}SFSKPR

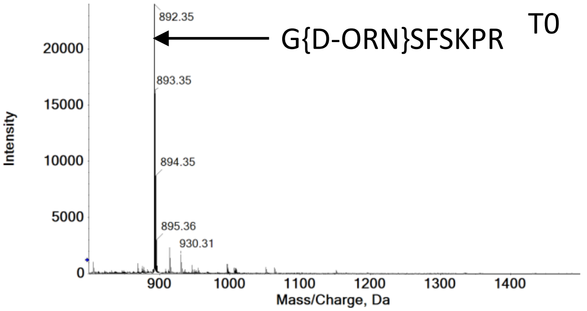


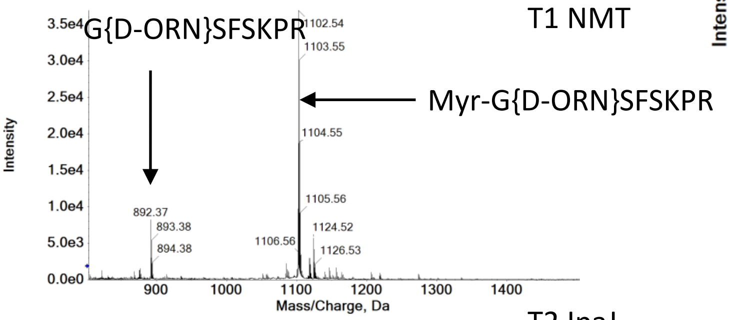


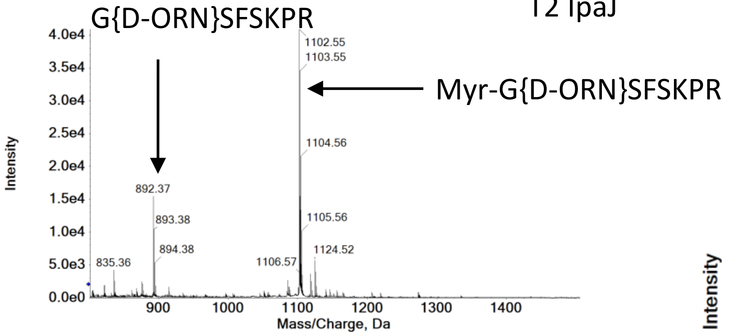


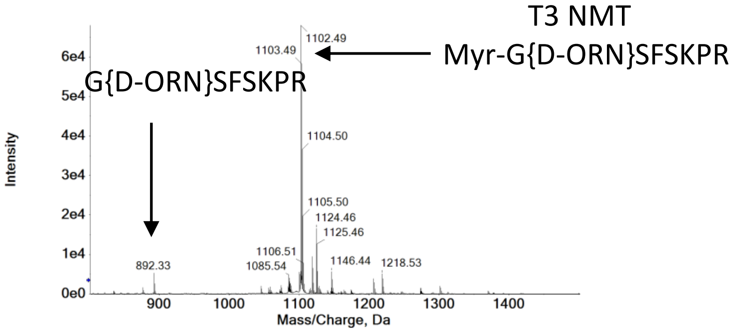


MSMS T1 Myr-G{D-ORN}SFSKPR (1102)

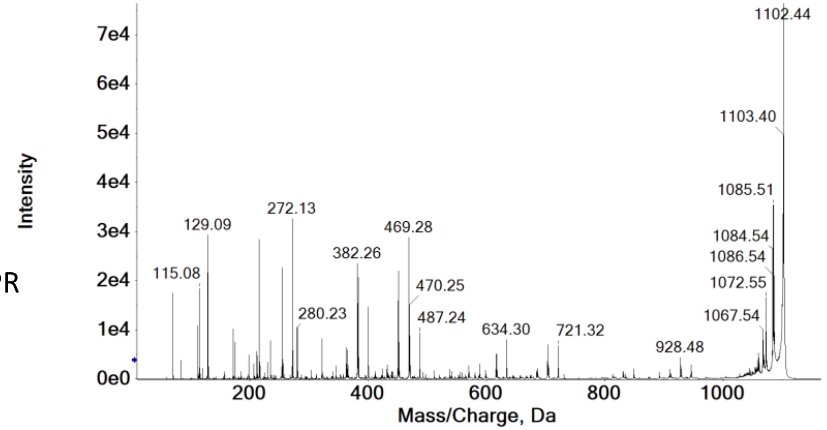


MSMS T0 G{D-ORN}SFSKPR (892)

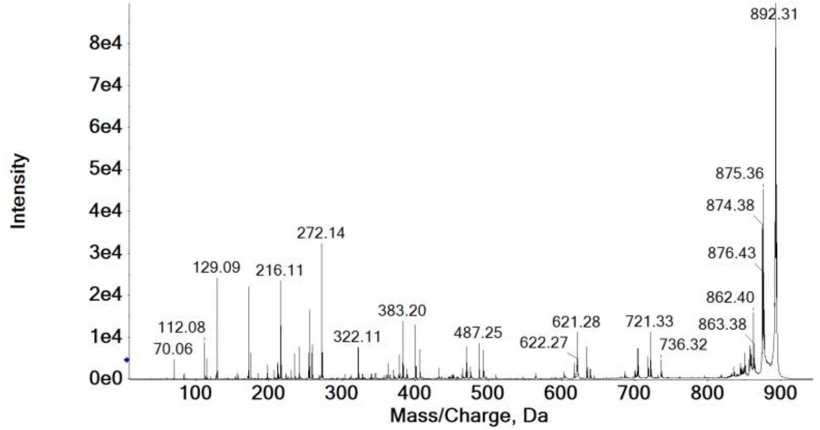


MSMS T2 Myr-G{D-ORN}SFSKPR (1102)

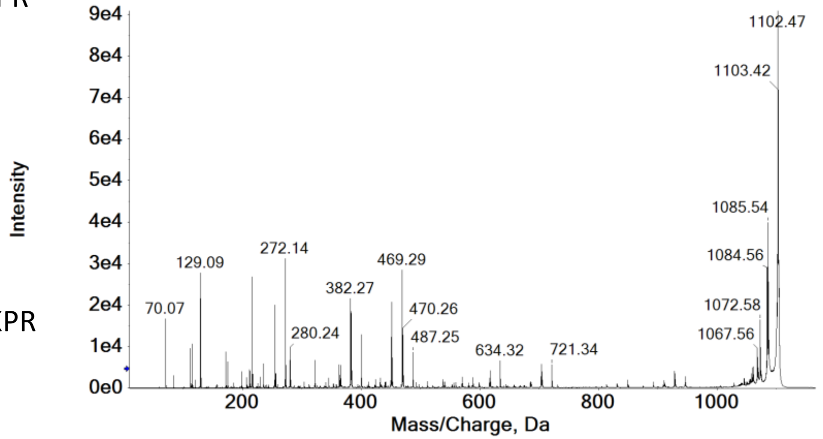


MSMS T3 Myr-G{D-ORN}SFSKPR (1102)

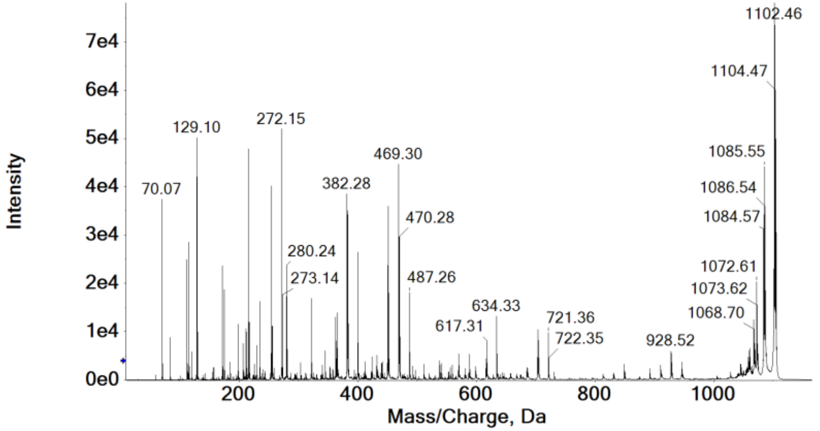

### Ac-KSFSKPR

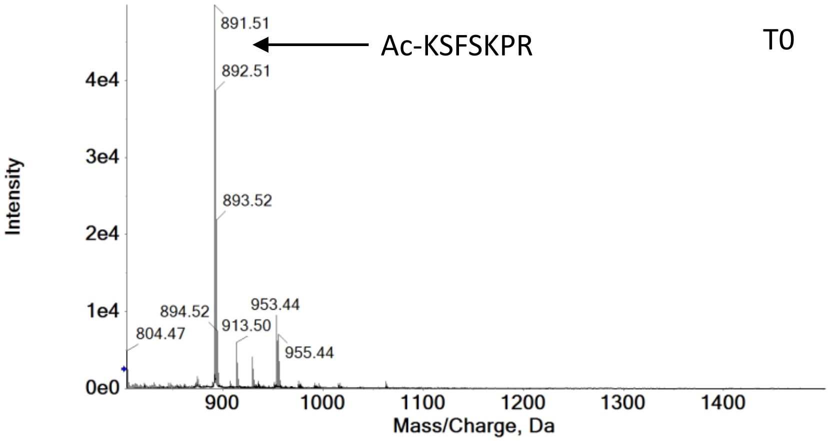


MSMS T0 Ac-KSFSKPR (891)

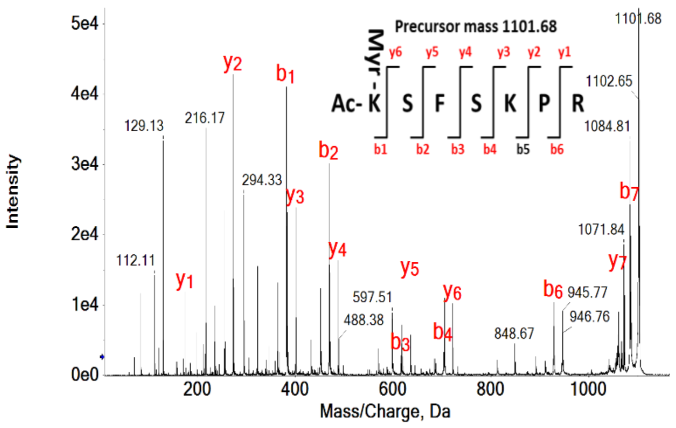


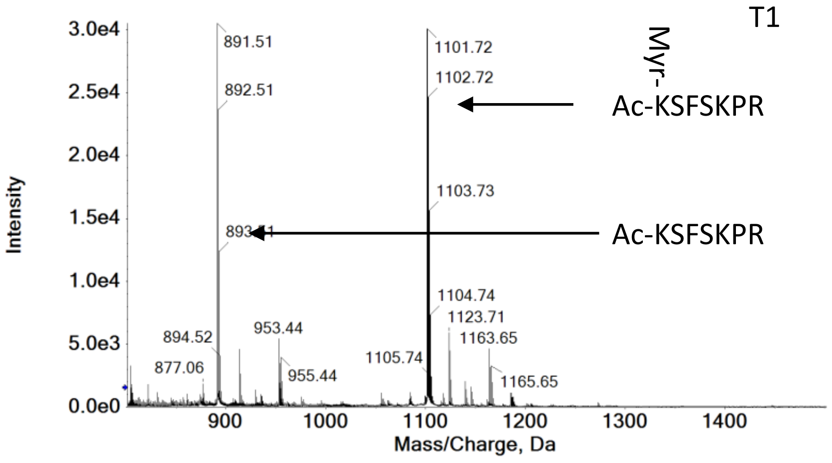


MSMS T1 Ac-K(Myr)SFSKPR (1101)

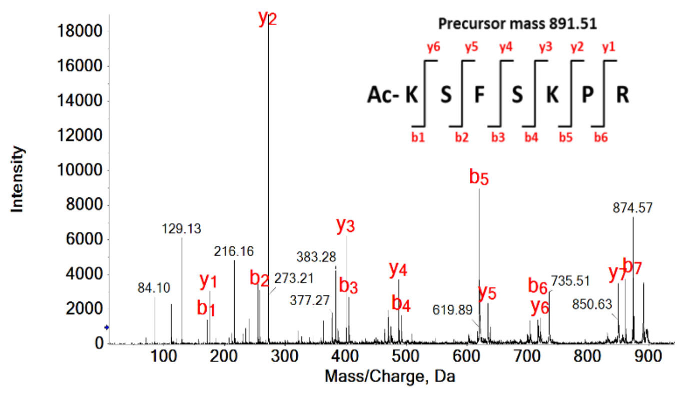

### Ac-{L-ORN}SFSKPR

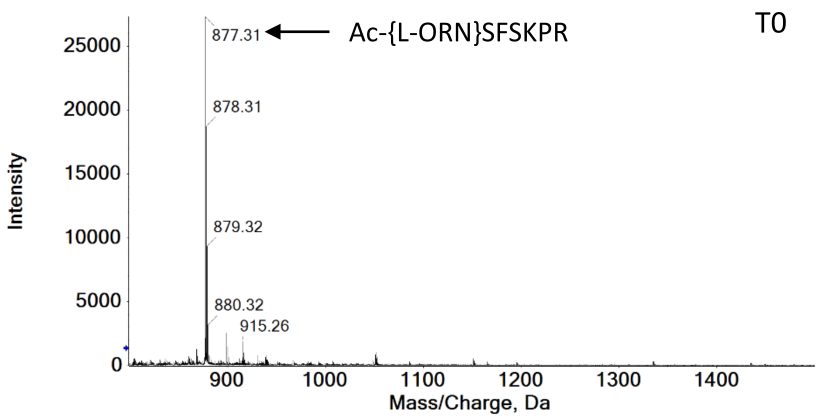


MSMS T0 Ac-KSFSKPR (891)

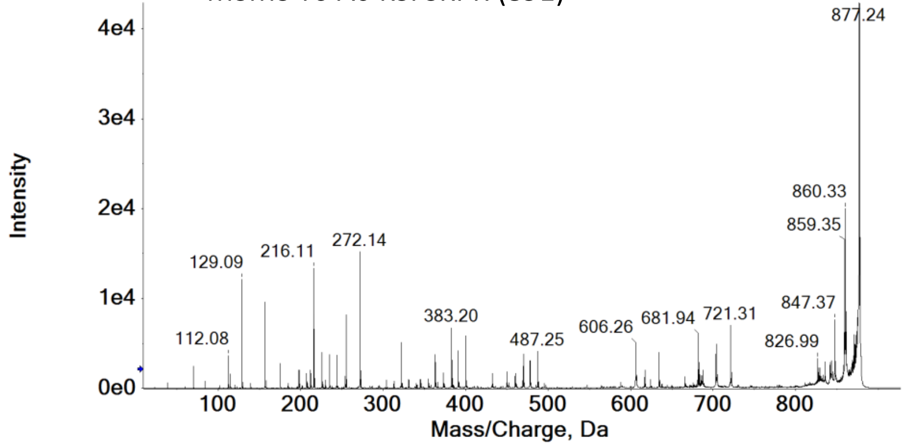


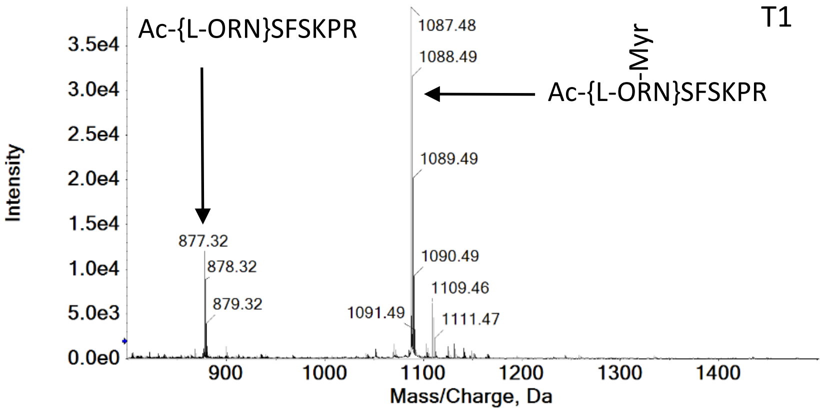


MSMS T1 Ac-K(Myr)SFSKPR (1101)

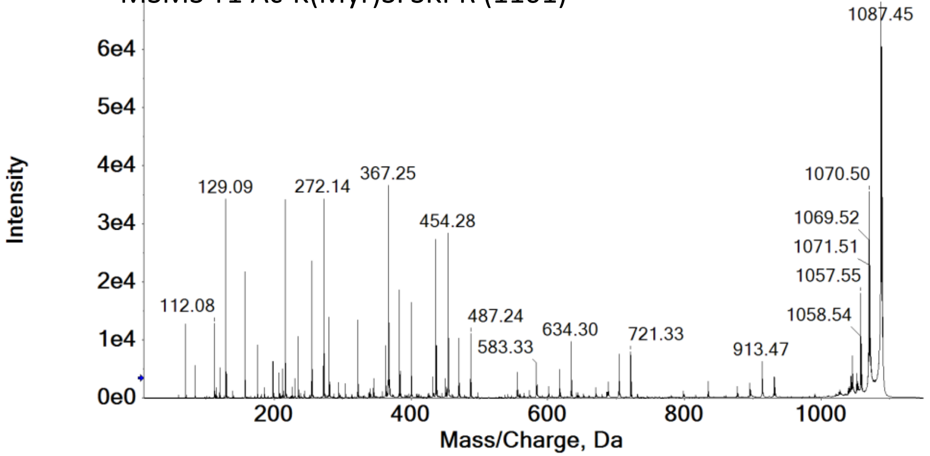

### Ac-{D-ORN}SFSKPR

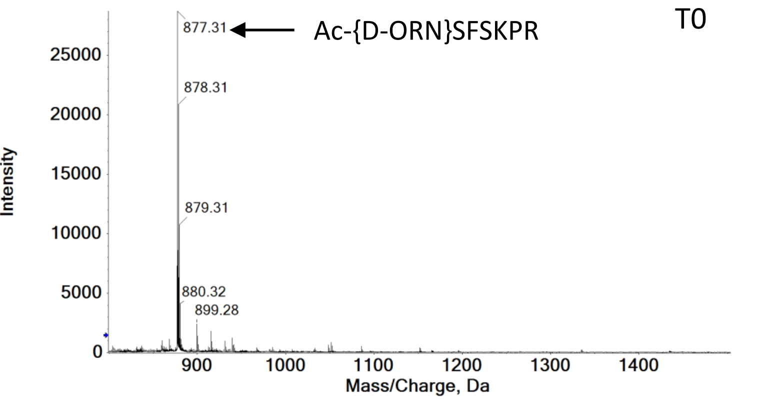


MSMS T0 Ac-{D-ORN}SFSKPR (877)

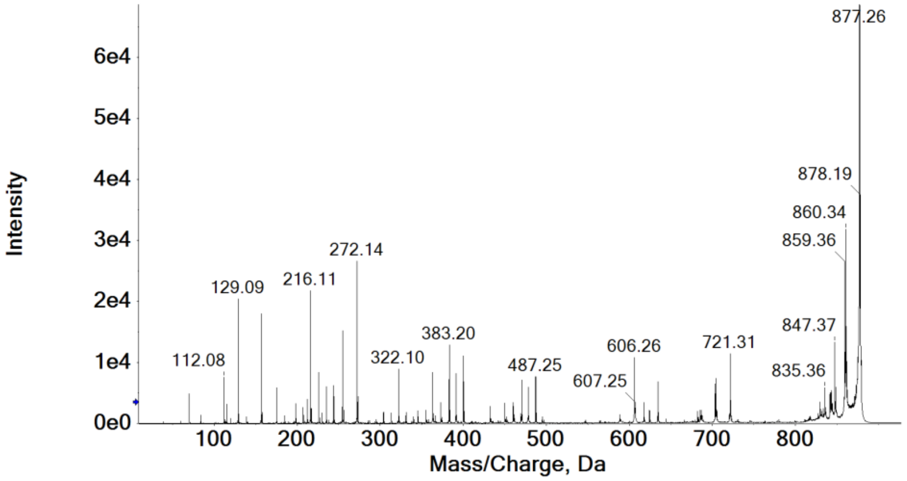


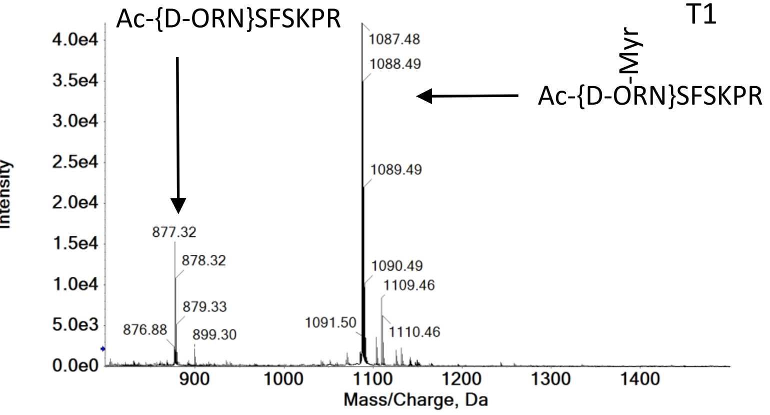


MSMS T1 Ac-{D-ORN}(Myr)SFSKPR (1087)

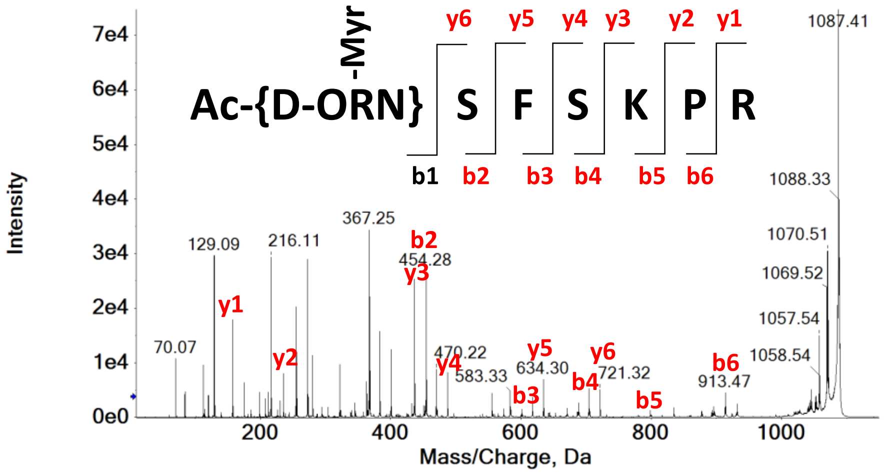


| N16 1087 | Mass/Charge (Da) | Intensity | ion |
|---|---|---|---|
| 9780 | 175,1064 | 6416,418 | y1 |
| 13956 | 272,2247 | 8455,216 | y2 |
| 18426 | 400,2117 | 12488,37 | y3 |
| 20091 | 454,2782 | 26812,13 | b2 |
| 21057 | 487,2445 | 8257,673 | y4 |
| 24166 | 601,3394 | 1466,144 | b3 |
| 25006 | 634,266 | 6512,837 | y5 |
| 26339 | 688,3574 | 2666,876 | b4 |
| 27125 | 721,3122 | 5124,183 | y6 |
| 29297 | 816,4788 | 361,3072 | b5 |
| 31381 | 913,469 | 4516,706 | b6 |

### Ac-{D-DAB}SFSKPR

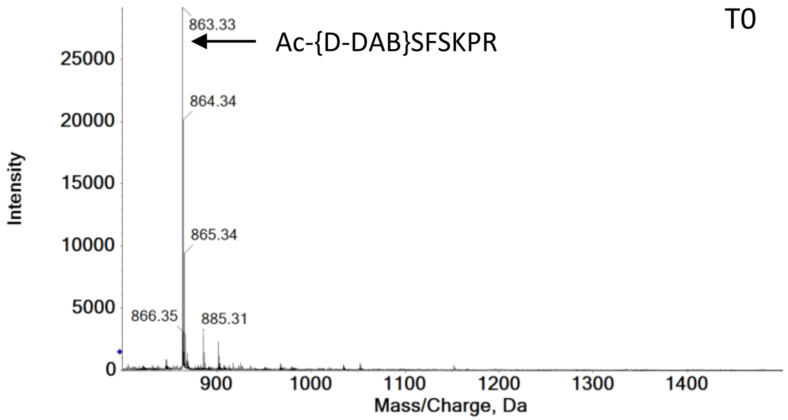


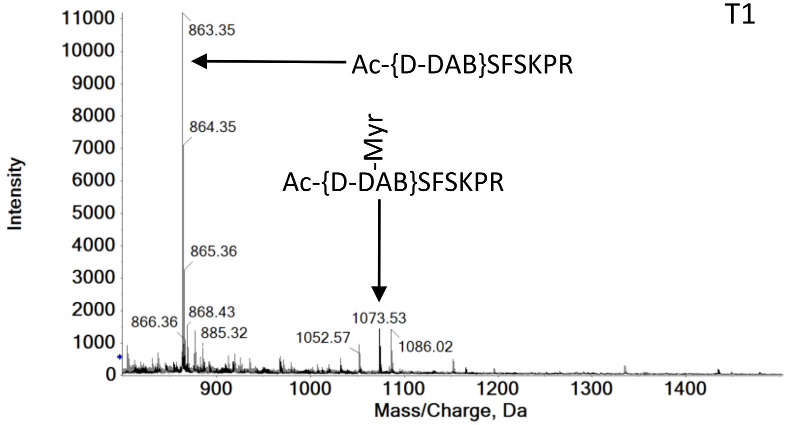


MSMS T0 Ac-{D-DAB}SFSKPR (863)

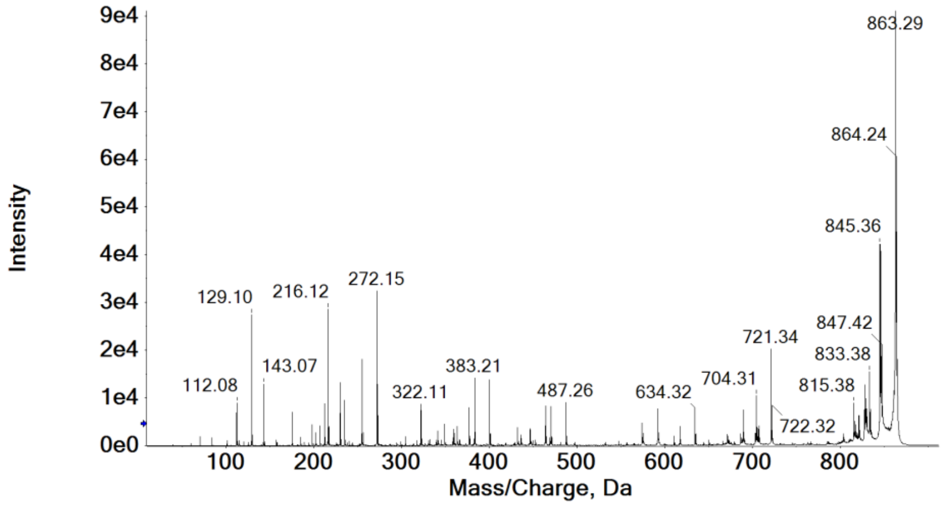


MSMS T1 Ac-{D-DAB}(Myr)SFSKPR (1073)

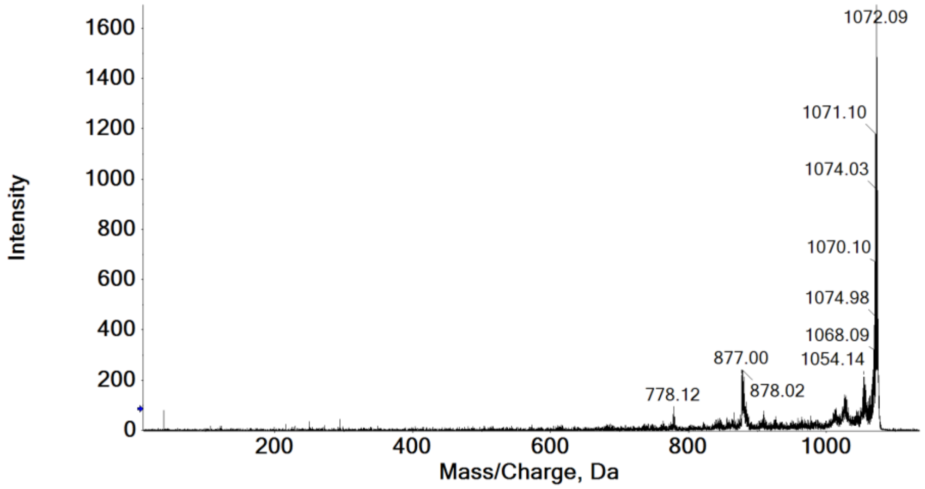

### Ac-{DAP}SFSKPR

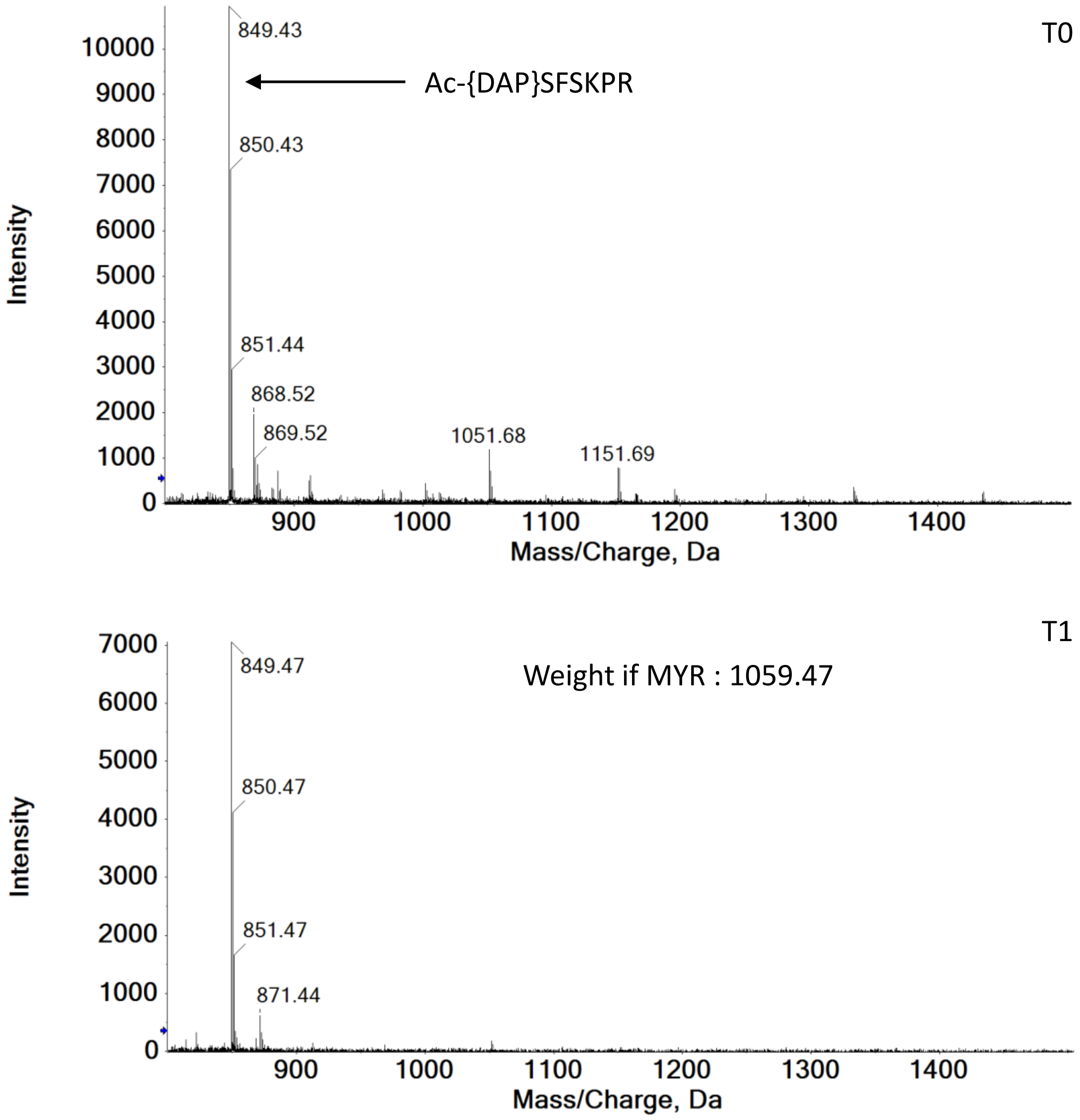

### KSFSKPR

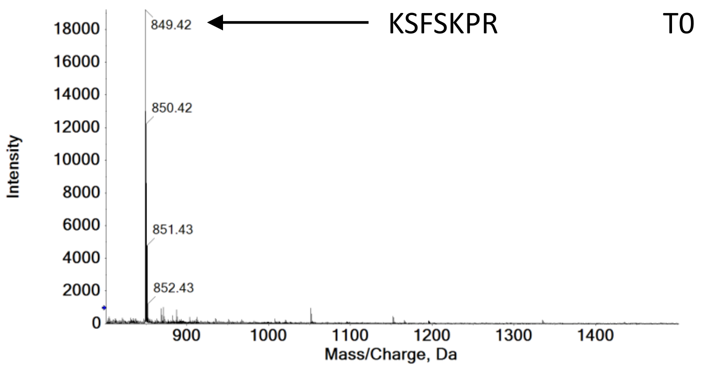


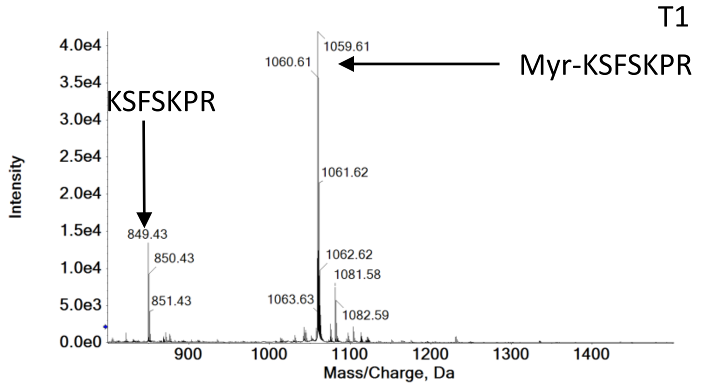


MSMS T0 KSFSKPR (849)

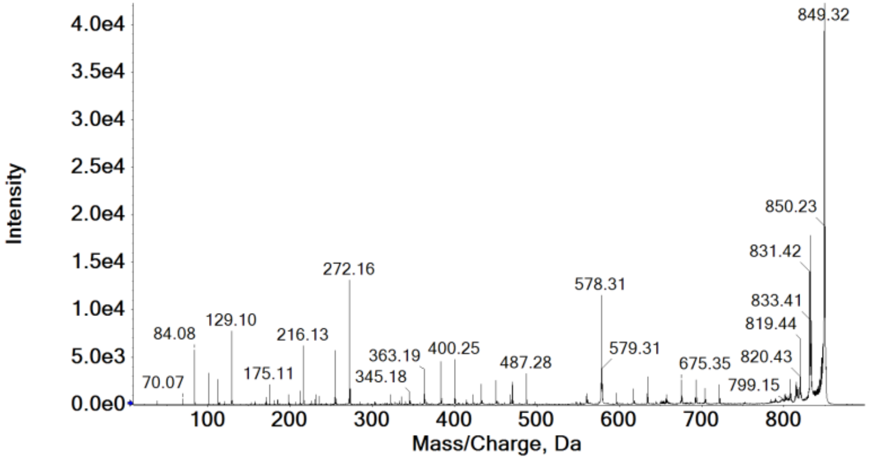


MSMS T1 Myr-KSFSKPR (1059)

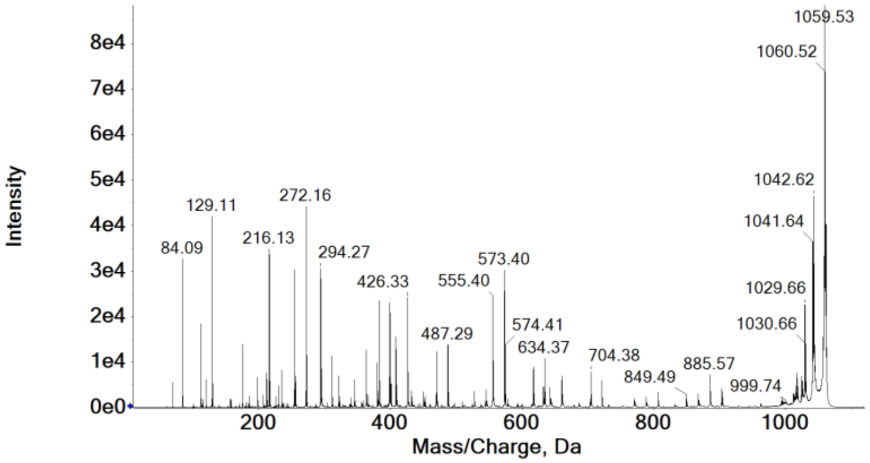

### {ORN}SFSKPR

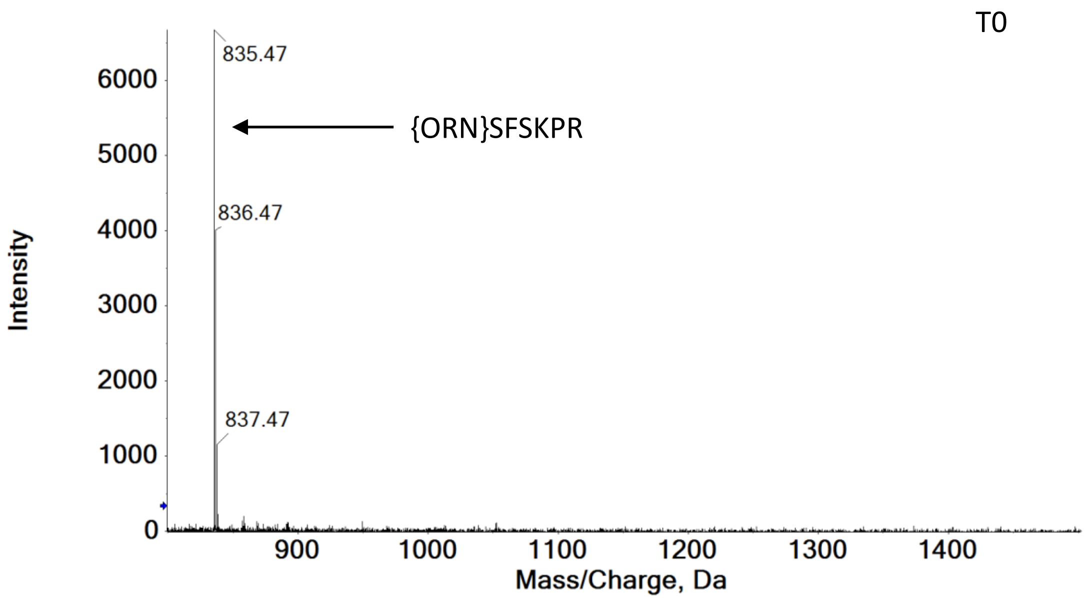
T0
835.47
{ORN}SFSKPR
836.47
837.47
Intensity
6000
5000
4000
3000
2000
1000
0
900
1000
1100
1200
1300
1400
Mass/Charge, Da

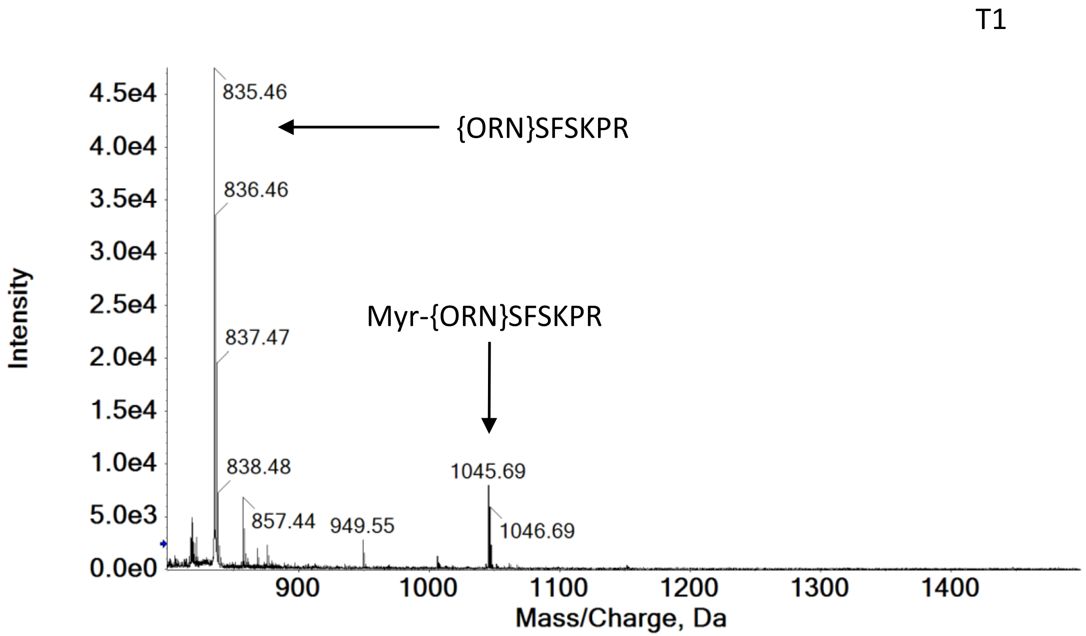
T1
835.46
{ORN}SFSKPR
836.46
837.47
Myr-{ORN}SFSKPR
838.48
857.44
949.55
1045.69
1046.69
Intensity
4.5e4
4.0e4
3.5e4
3.0e4
2.5e4
2.0e4
1.5e4
1.0e4
5.0e3
0.0e0
900
1000
1100
1200
1300
1400
Mass/Charge, Da

### {D-ORN}SFSKPR

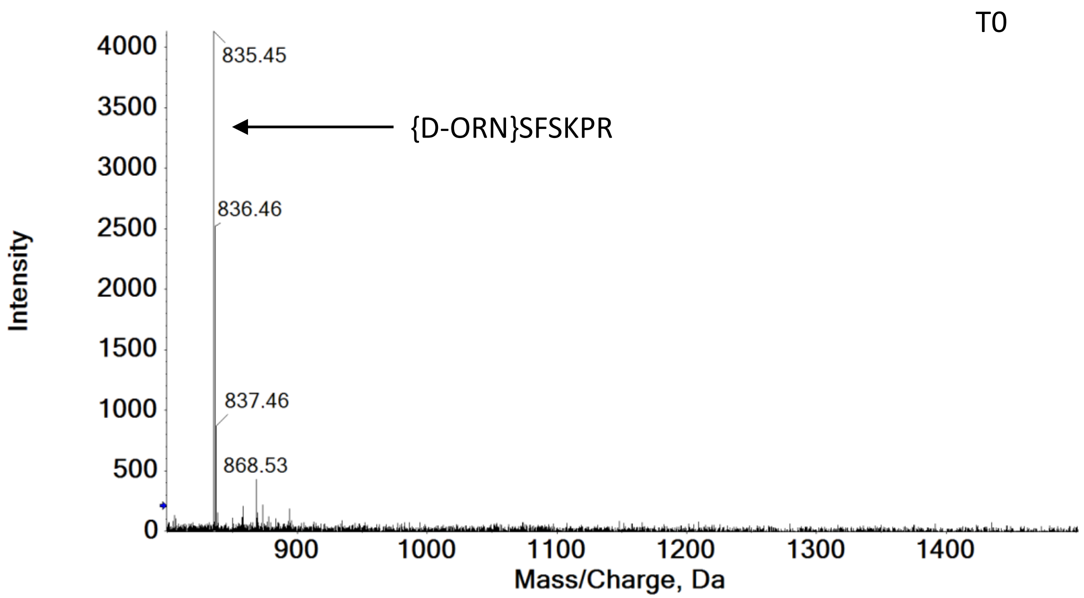

T0
Intensity
4000
3500
3000
2500
2000
1500
1000
500
0
835.45
{D-ORN}SFSKPR
836.46
837.46
868.53
900
1000
1100
1200
1300
1400
Mass/Charge, Da


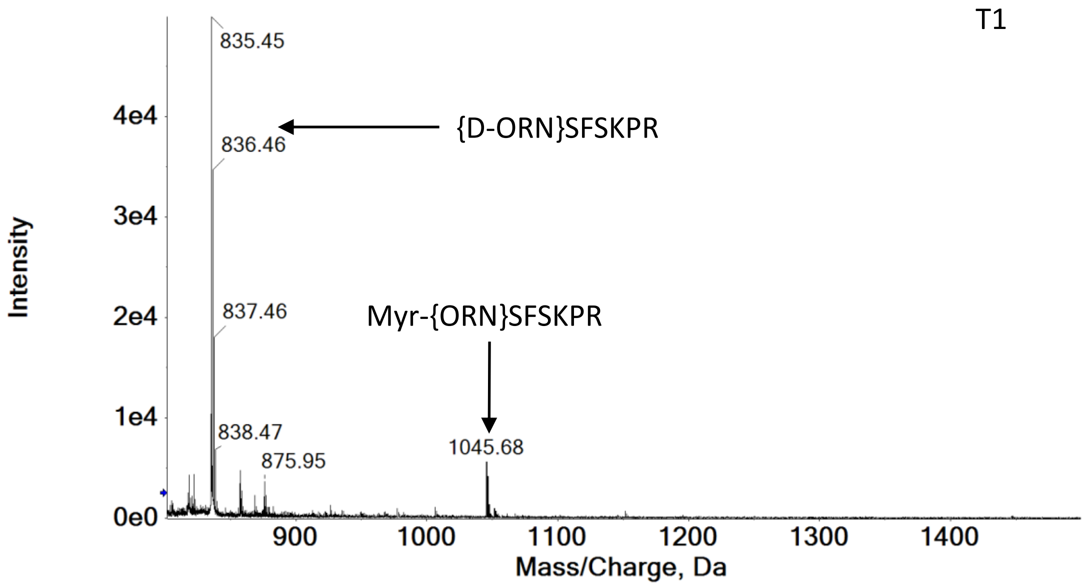

T1
Intensity
4e4
3e4
2e4
1e4
0e0
835.45
{D-ORN}SFSKPR
836.46
837.46
Myr-{ORN}SFSKPR
838.47
875.95
1045.68
900
1000
1100
1200
1300
1400
Mass/Charge, Da

### {DAB}SFSKPR

T0

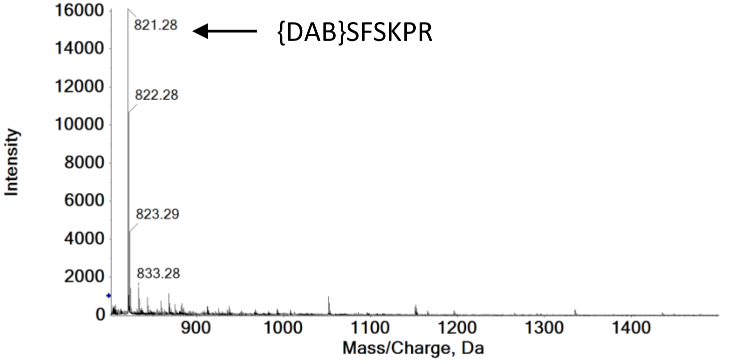


MSMS T0 {DAB}SFSKPR (821)

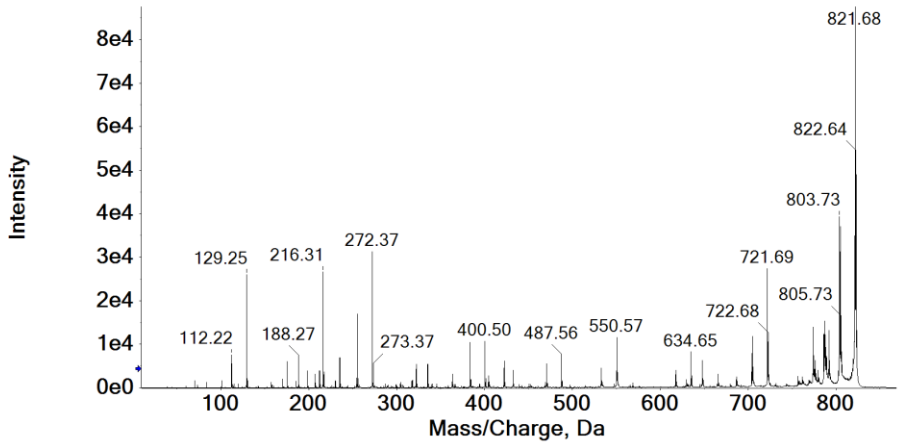


T1

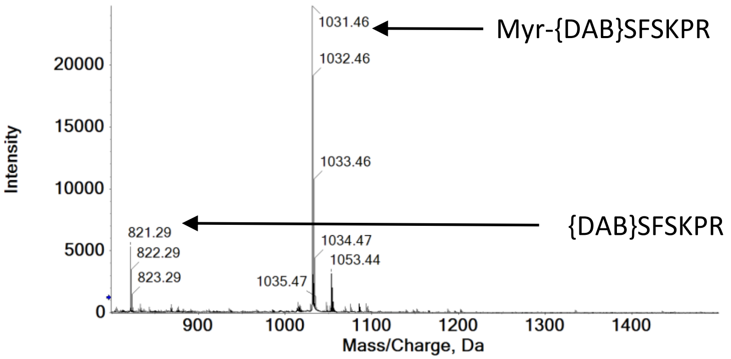


MSMS T1 Myr-{D-DAB}SFSKPR (1031)

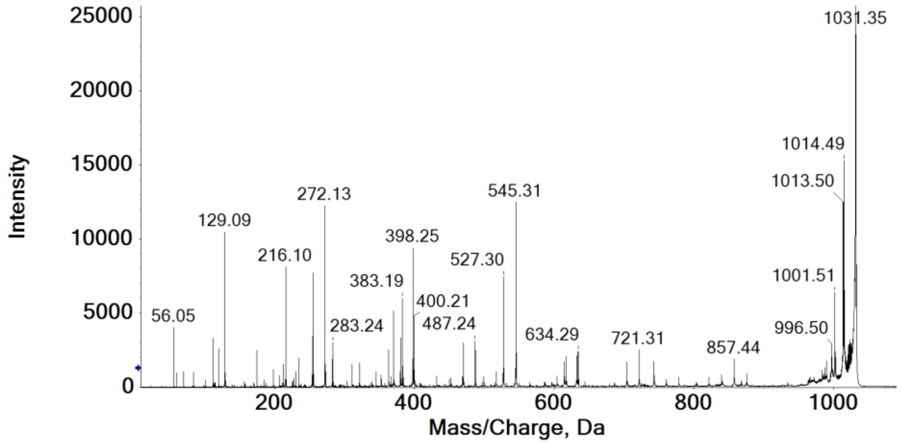

### KQNSKLRP

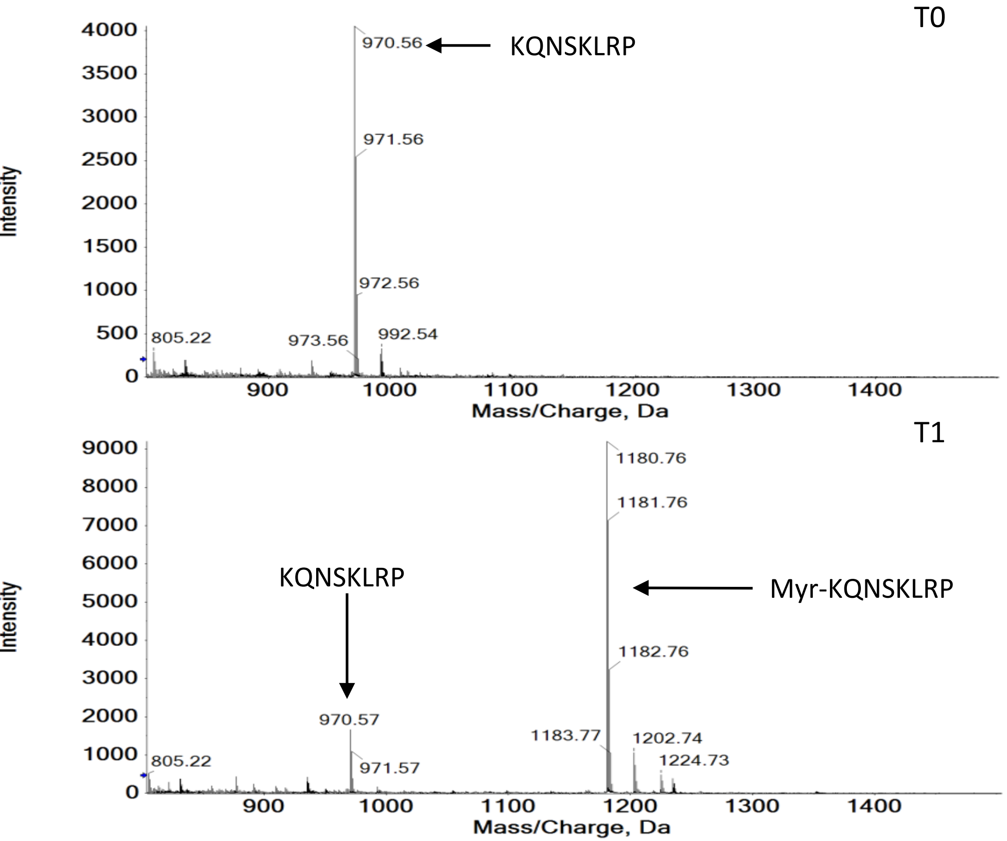
T0
KQNSKLRP
970.56
971.56
972.56
973.56
992.54
805.22
Intensity
Mass/Charge, Da
T1
KQNSKLRP
Myr-KQNSKLRP
1180.76
1181.76
1182.76
1183.77
1202.74
1224.73
970.57
971.57
805.22
Intensity
Mass/Charge, Da

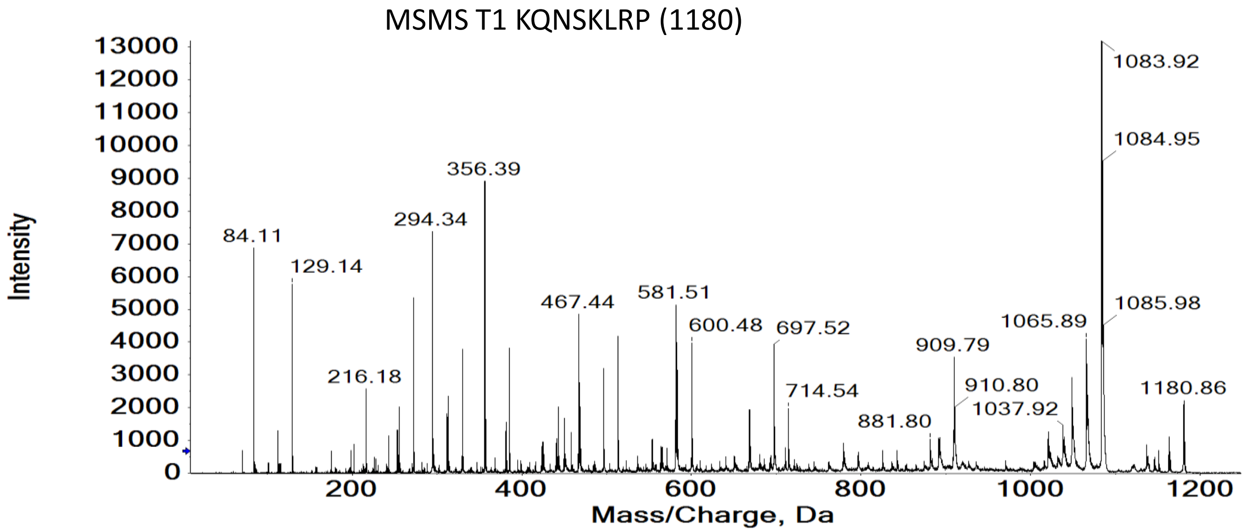
MSMS T1 KQNSKLRP (1180)
1083.92
1084.95
1085.98
1065.89
1037.92
1180.86
909.79
910.80
881.80
356.39
294.34
84.11
129.14
216.18
467.44
581.51
600.48
697.52
714.54
Intensity
Mass/Charge, Da

### Ac-KFSKPR

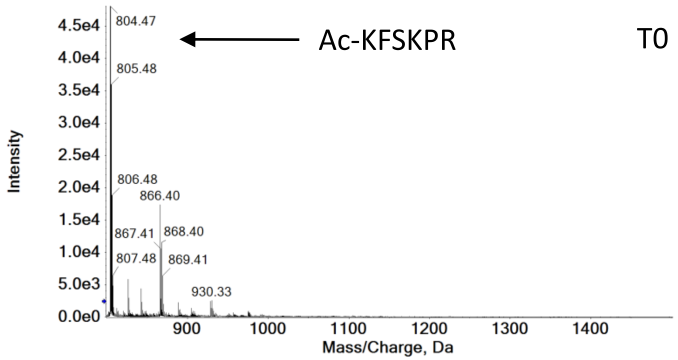

T0
Ac-KFSKPR
804.47
805.48
806.48
866.40
867.41
868.40
807.48
869.41
930.33
Intensity
Mass/Charge, Da


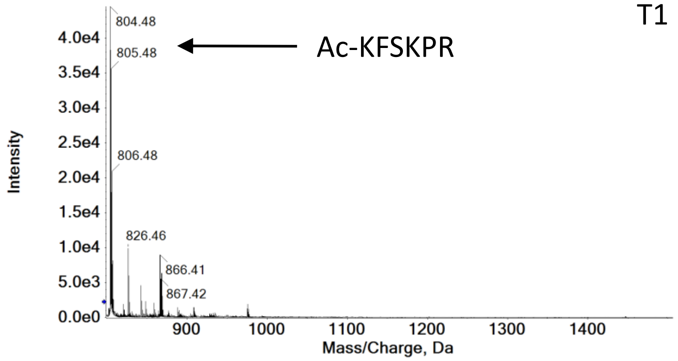

T1
Ac-KFSKPR
804.48
805.48
806.48
826.46
866.41
867.42
Intensity
Mass/Charge, Da

### KQNS{ORN}LRP

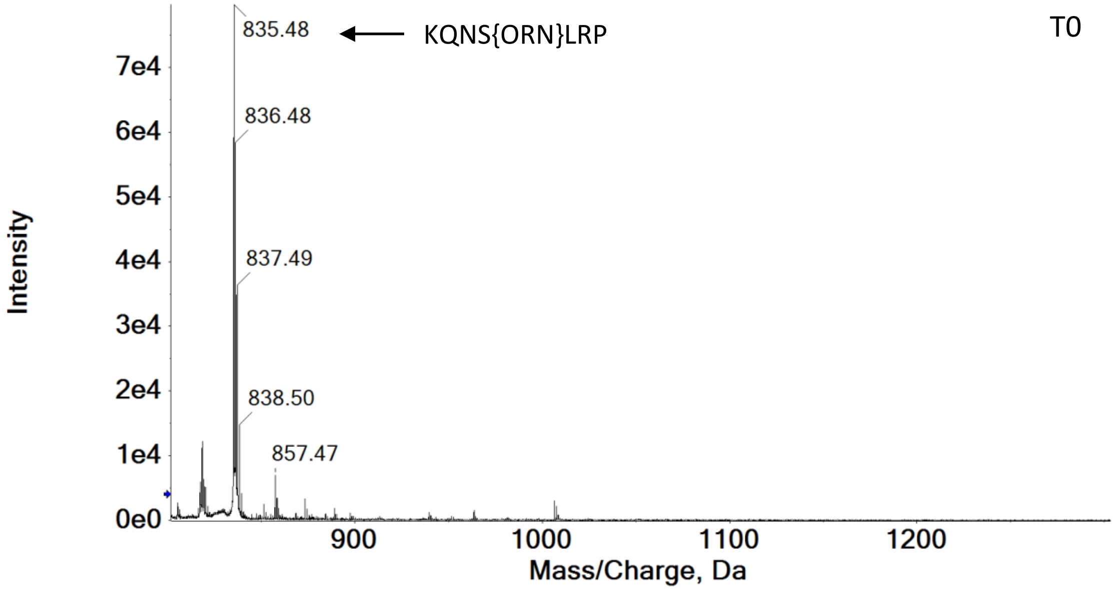
T0
835.48
KQNS{ORN}LRP
836.48
837.49
838.50
857.47
7e4
6e4
5e4
4e4
3e4
2e4
1e4
0e0
Intensity
900
1000
1100
1200
Mass/Charge, Da

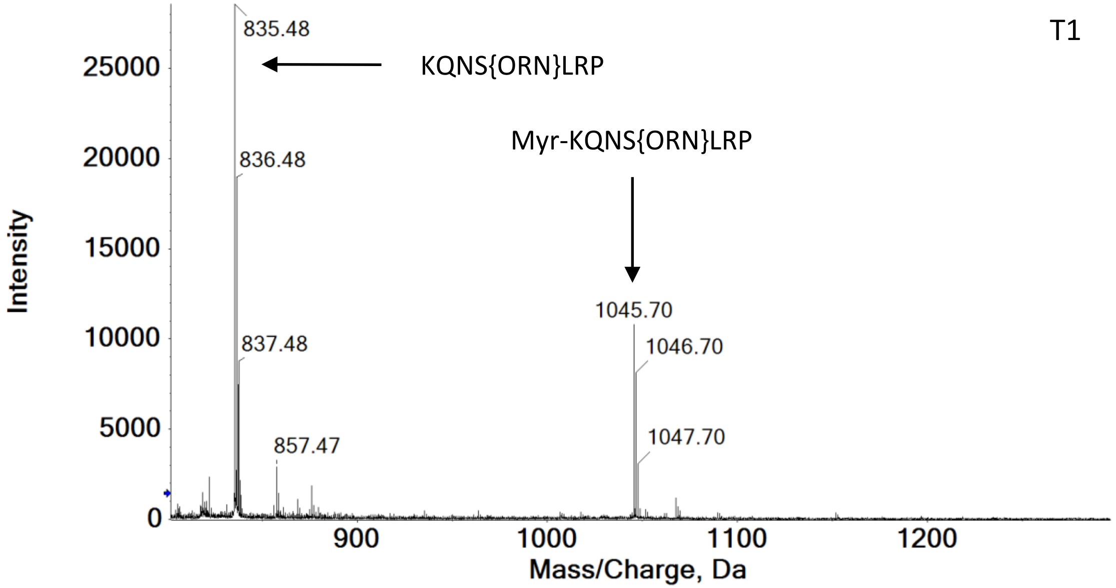
T1
835.48
KQNS{ORN}LRP
Myr-KQNS{ORN}LRP
836.48
837.48
857.47
1045.70
1046.70
1047.70
25000
20000
15000
10000
5000
0
Intensity
900
1000
1100
1200
Mass/Charge, Da

### KQNS{DAB}LRP

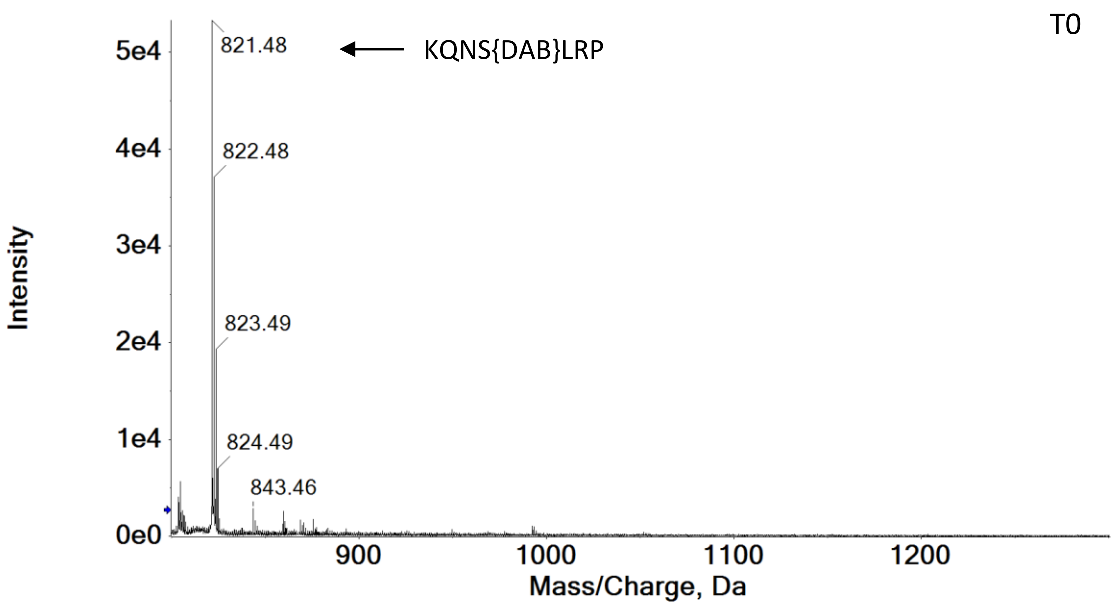

T0
821.48
KQNS{DAB}LRP
822.48
823.49
824.49
843.46
5e4
4e4
3e4
2e4
1e4
0e0
Intensity
900
1000
1100
1200
Mass/Charge, Da


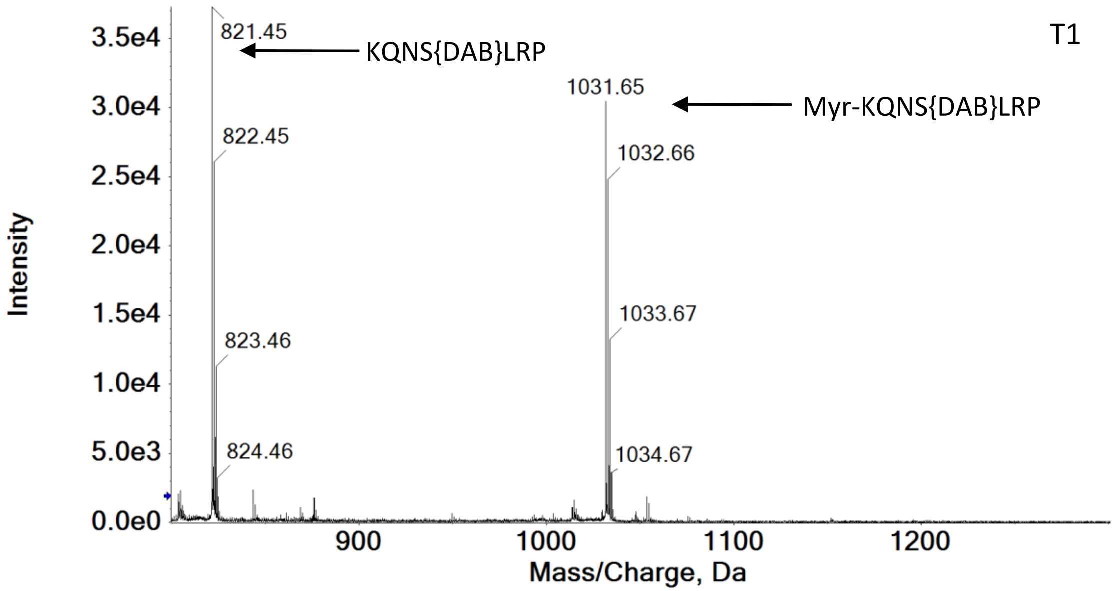

T1
821.45
KQNS{DAB}LRP
1031.65
Myr-KQNS{DAB}LRP
822.45
1032.66
1033.67
823.46
824.46
1034.67
3.5e4
3.0e4
2.5e4
2.0e4
1.5e4
1.0e4
5.0e3
0.0e0
Intensity
900
1000
1100
1200
Mass/Charge, Da

#### KQNS{DAP}LRP

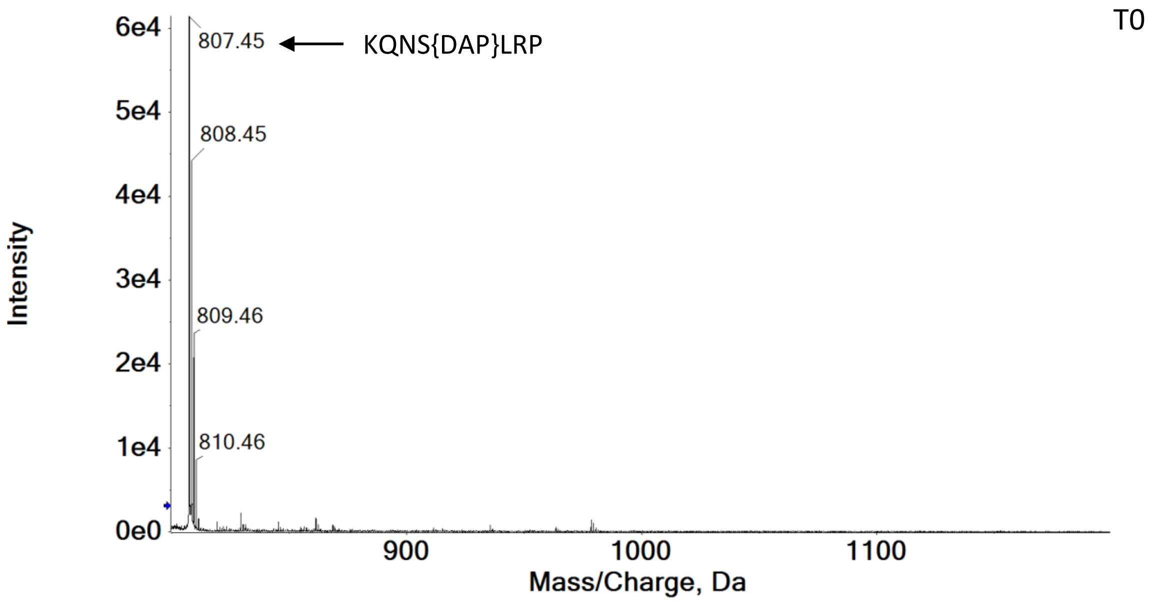
T0
Intensity
6e4
5e4
4e4
3e4
2e4
1e4
0e0
807.45
KQNS{DAP}LRP
808.45
809.46
810.46
900
1000
1100
Mass/Charge, Da

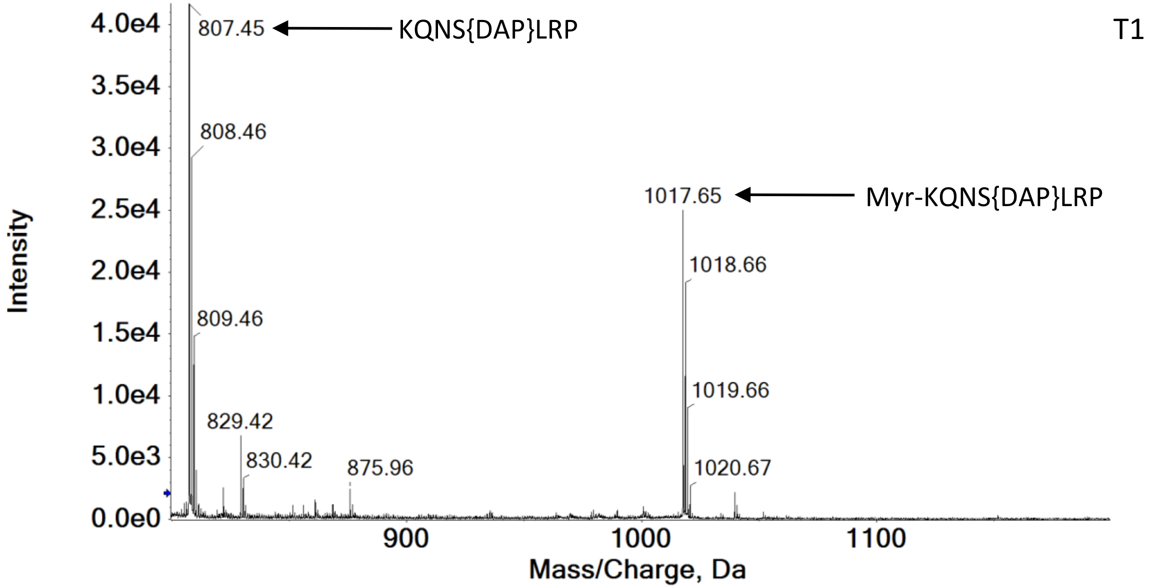
T1
Intensity
4.0e4
3.5e4
3.0e4
2.5e4
2.0e4
1.5e4
1.0e4
5.0e3
0.0e0
807.45
KQNS{DAP}LRP
808.46
809.46
829.42
830.42
875.96
1017.65
Myr-KQNS{DAP}LRP
1018.66
1019.66
1020.67
900
1000
1100
Mass/Charge, Da

### Ac-{D-ORN}SNSKLK

T0

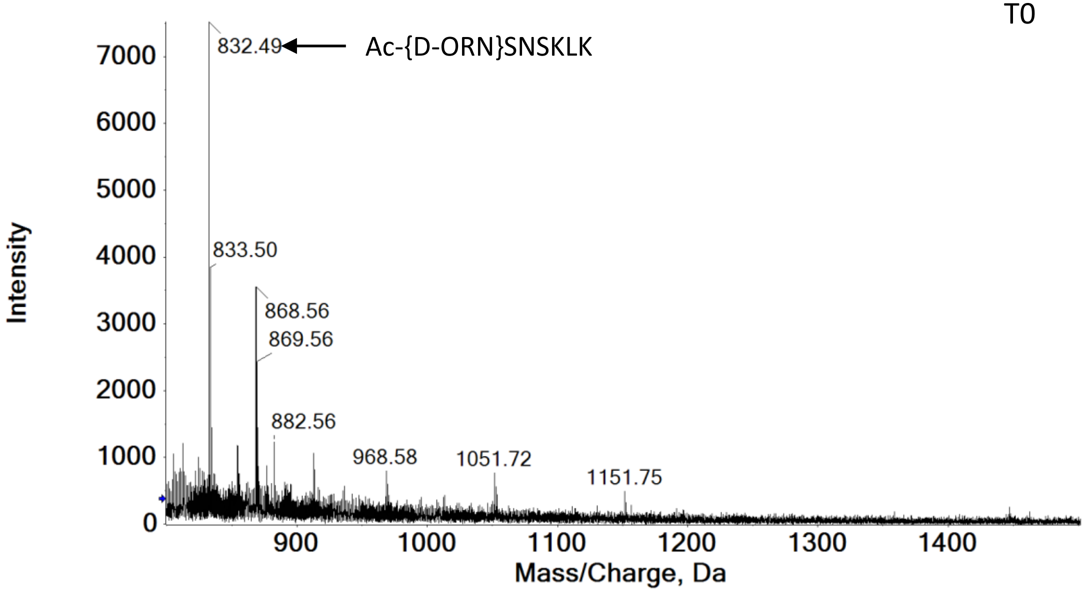


T1

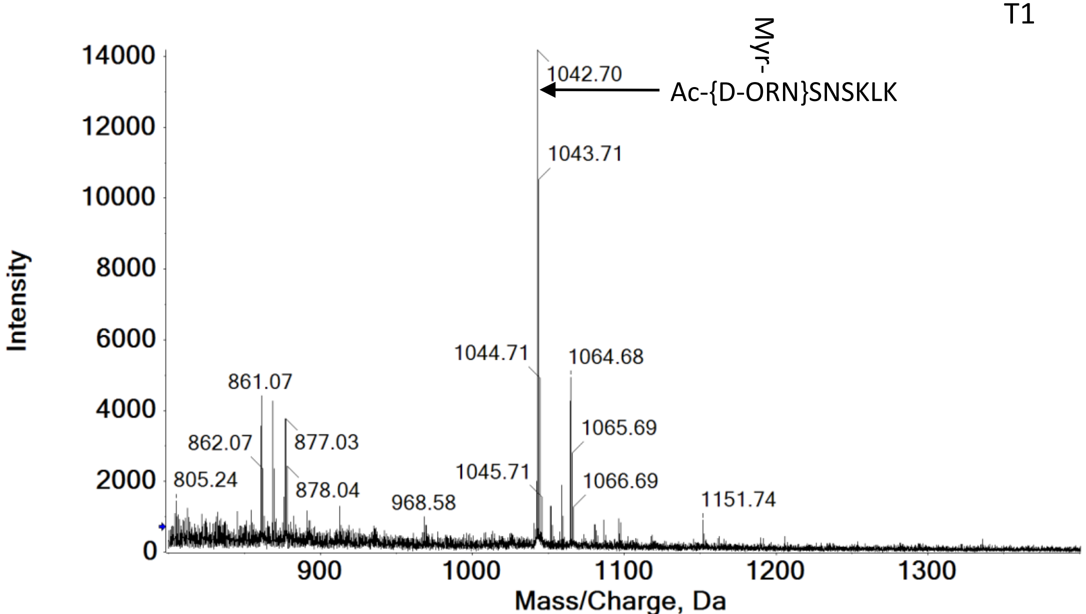

#### KSNAKLKP

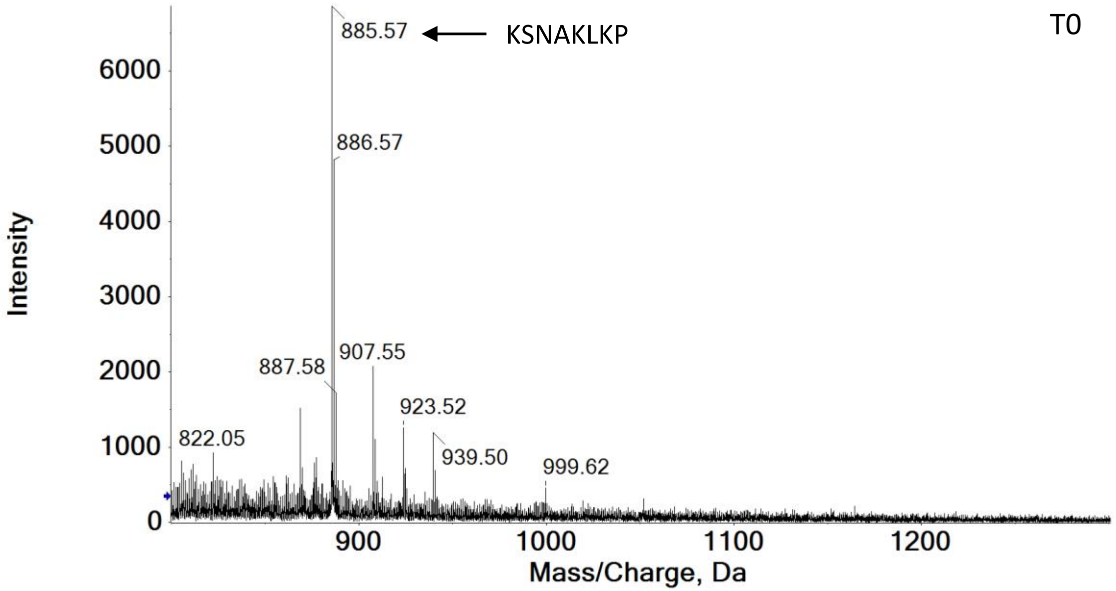
T0
885.57
KSNAKLKP
886.57
887.58
907.55
923.52
822.05
939.50
999.62
6000
5000
4000
3000
2000
1000
0
Intensity
900
1000
1100
1200
Mass/Charge, Da

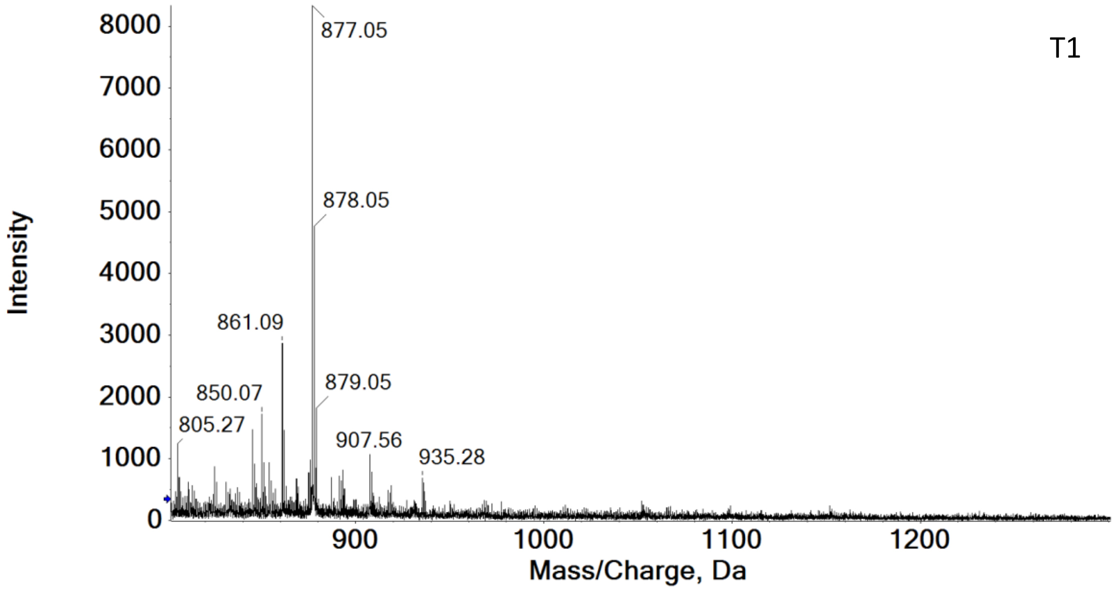
T1
877.05
878.05
861.09
850.07
879.05
805.27
907.56
935.28
8000
7000
6000
5000
4000
3000
2000
1000
0
Intensity
900
1000
1100
1200
Mass/Charge, Da

### GGKSFSKP

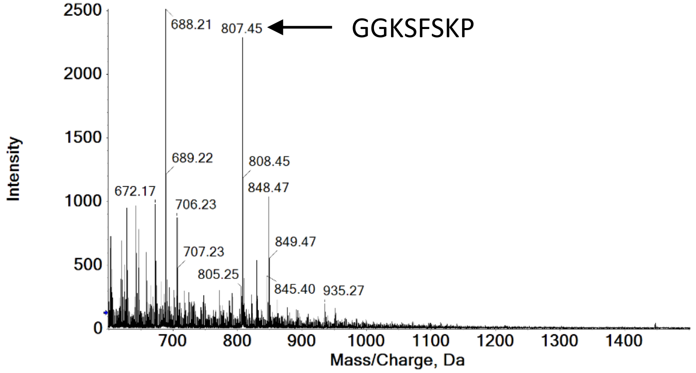


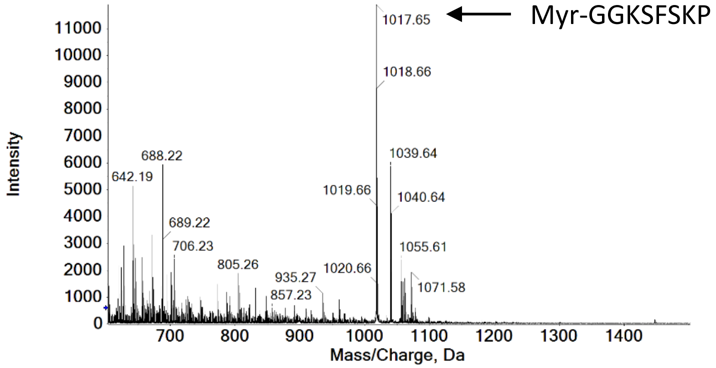


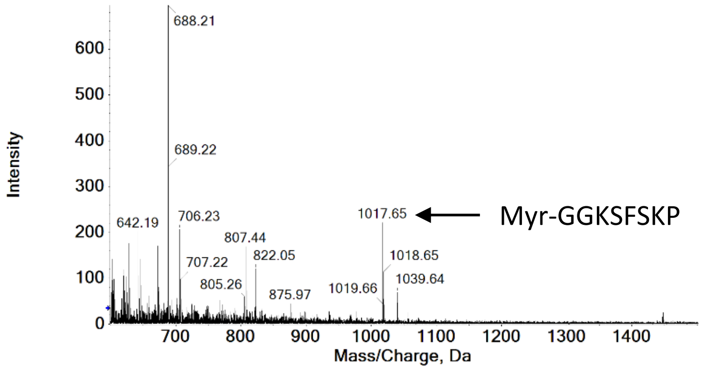


MSMS T0 GGKSFSKP (807)

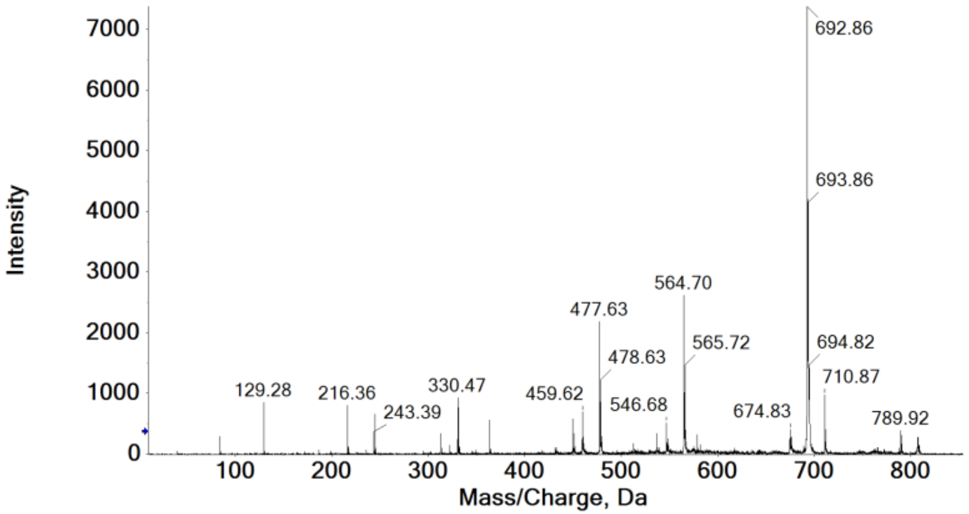


MSMS T1 Myr-GGKSFSKP (1017)

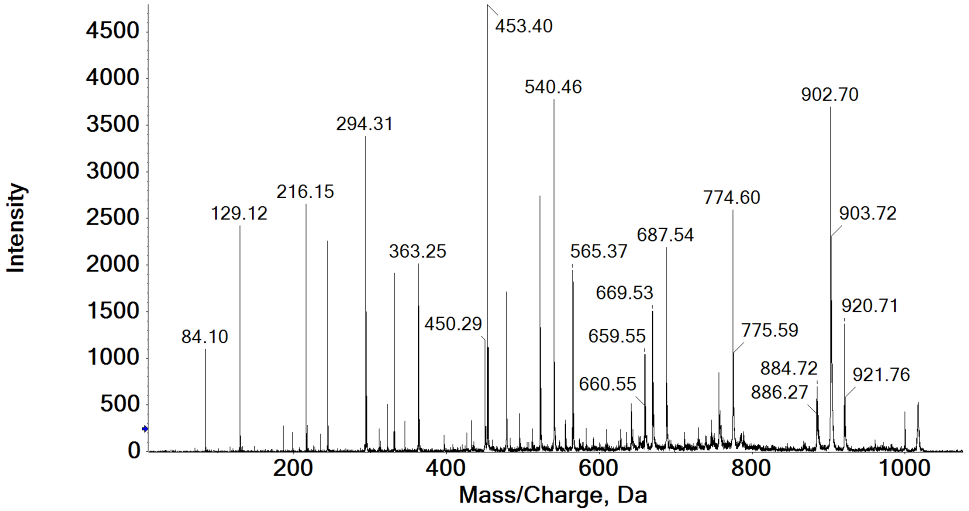

MSMS T1 Myr-GGKSFSKP (1017)

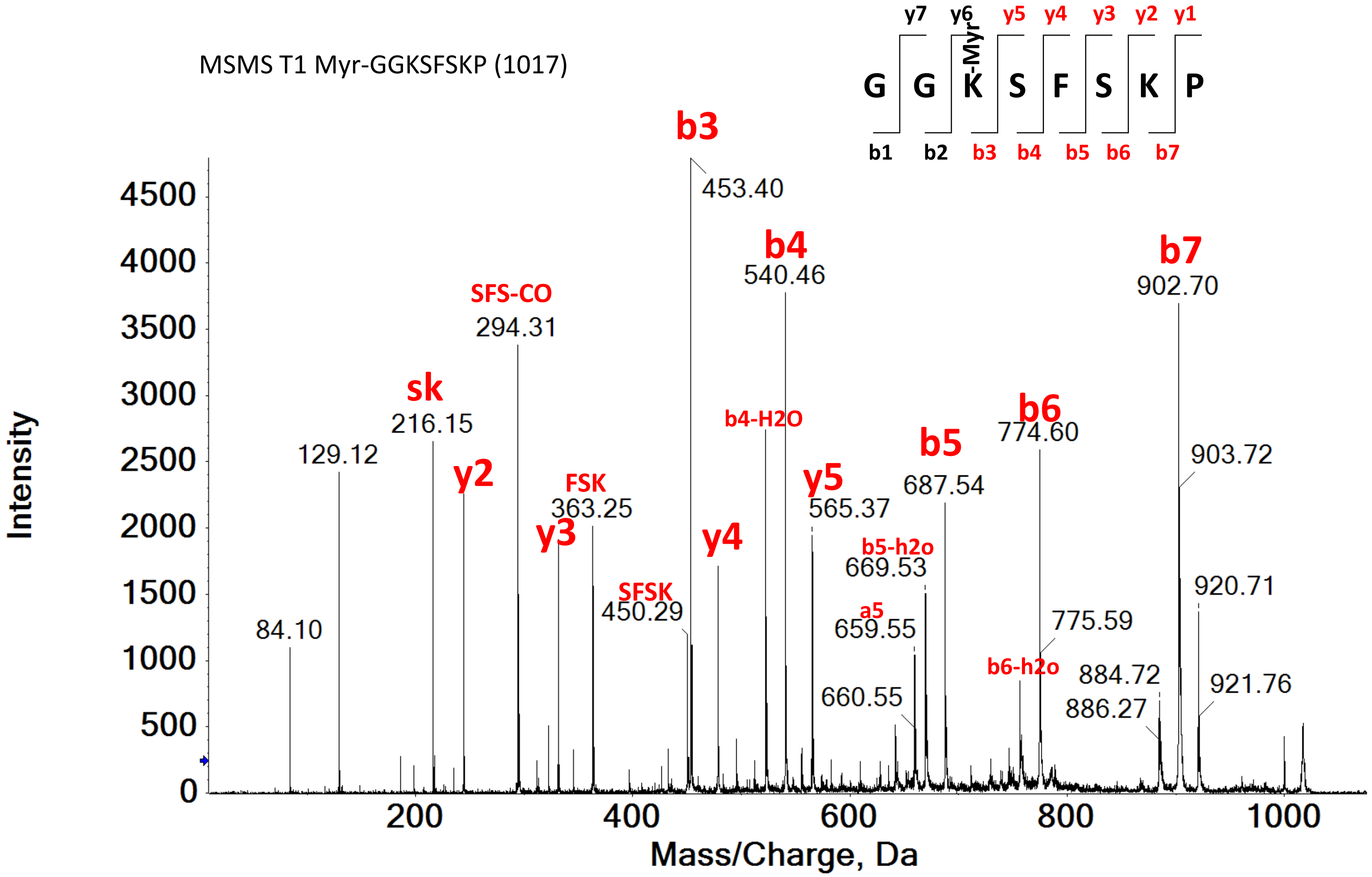

### AGKSFSKPR

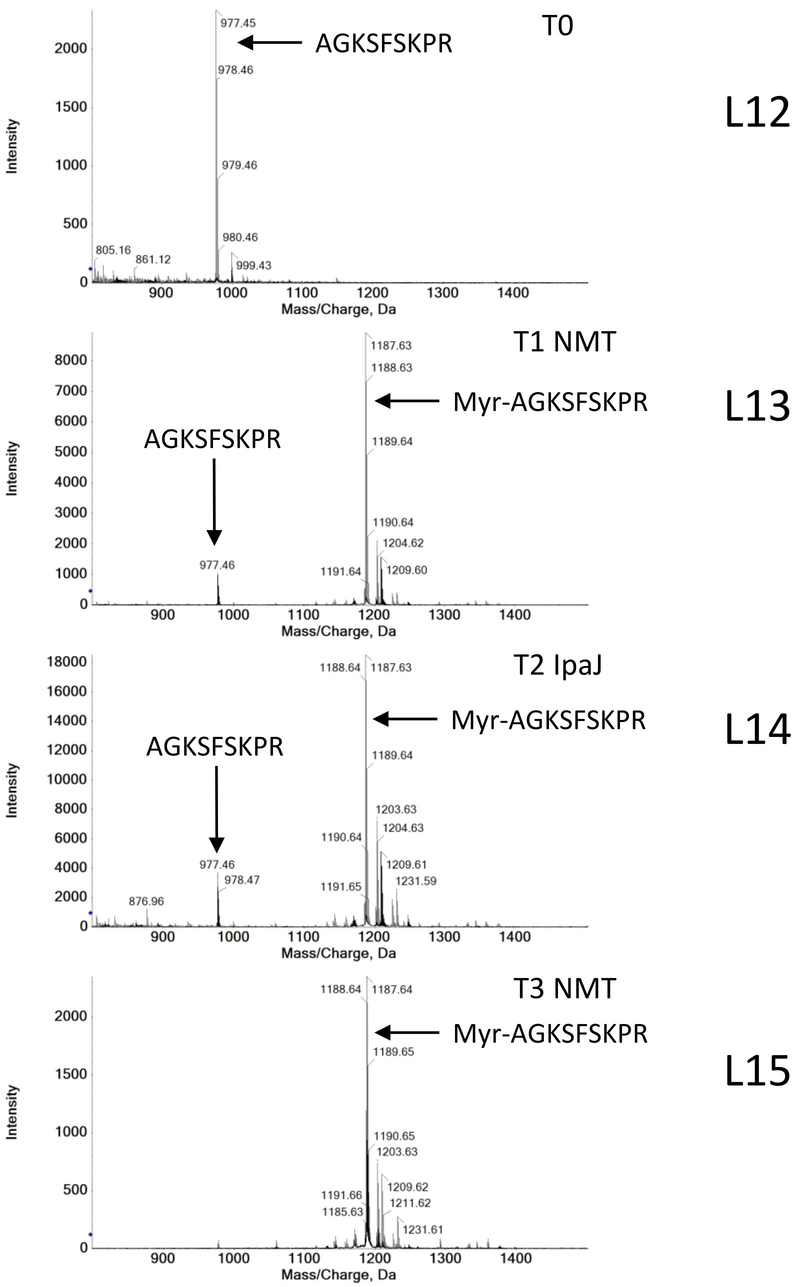
T0
AGKSFSKPR
977.45
978.46
979.46
980.46
999.43
805.16
861.12
L12
T1 NMT
Myr-AGKSFSKPR
AGKSFSKPR
1187.63
1188.63
1189.64
1190.64
1204.62
1209.60
1191.64
977.46
L13
T2 IpaJ
Myr-AGKSFSKPR
AGKSFSKPR
1188.64
1187.63
1189.64
1203.63
1204.63
1190.64
1209.61
1231.59
1191.65
977.46
978.47
876.96
L14
T3 NMT
Myr-AGKSFSKPR
1188.64
1187.64
1189.65
1190.65
1203.63
1209.62
1211.62
1191.66
1185.63
1231.61
L15
Intensity
Mass/Charge, Da

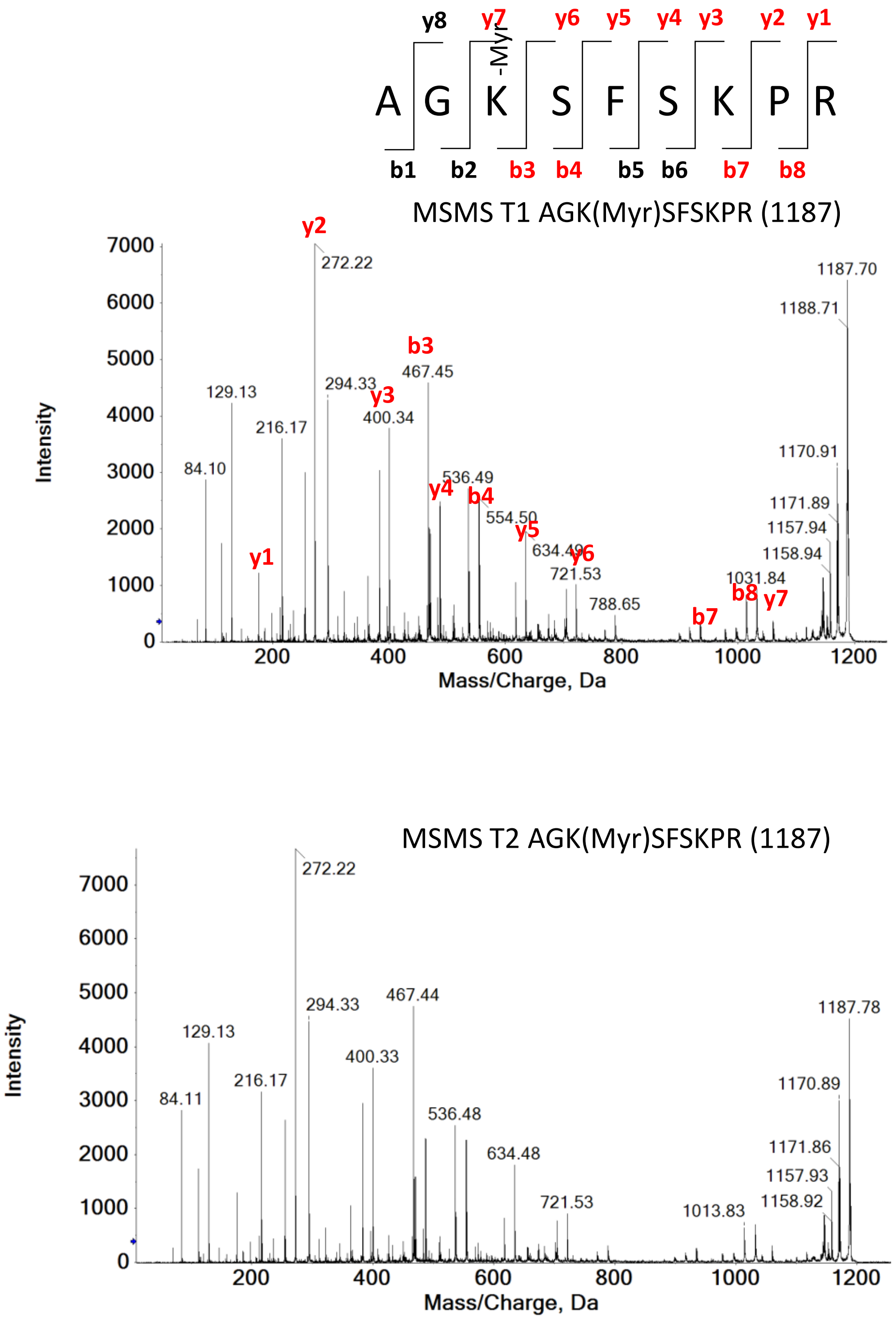
y8 y7 y6 y5 y4 y3 y2 y1
A G K S F S K P R
-Myr
b1 b2 b3 b4 b5 b6 b7 b8
MSMS T1 AGK(Myr)SFSKPR (1187)
272.22
1187.70
1188.71
294.33
467.45
129.13
400.34
216.17
84.10
1170.91
536.49
1171.89
554.50
1157.94
634.49
1158.94
721.53
1031.84
788.65
MSMS T2 AGK(Myr)SFSKPR (1187)
272.22
294.33
467.44
1187.78
129.13
400.33
216.17
84.11
1170.89
536.48
634.48
1171.86
1157.93
721.53
1013.83
1158.92
Intensity
Mass/Charge, Da

### Ac-GGKSFSKPR

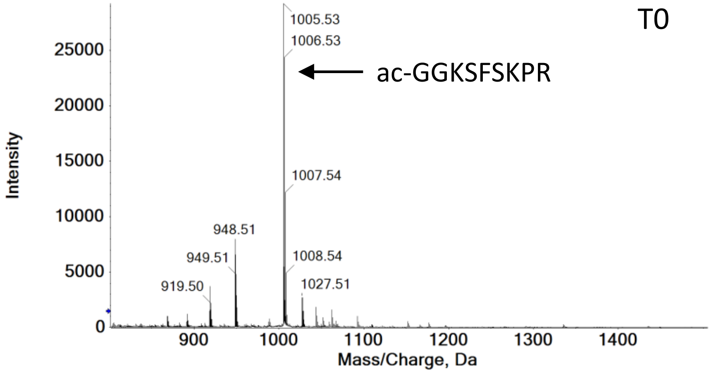


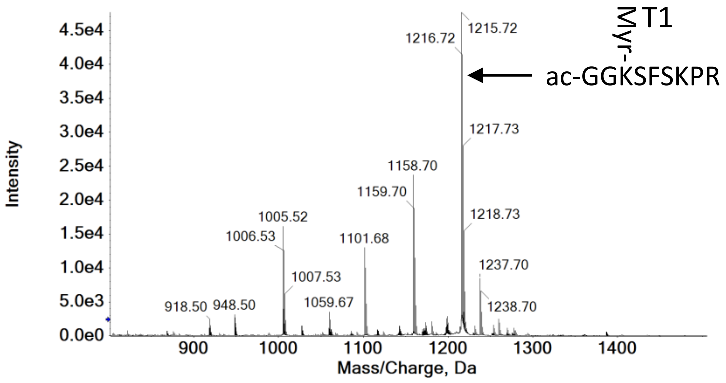


MSMS T0 ac-GGKSFSKPR (1005)

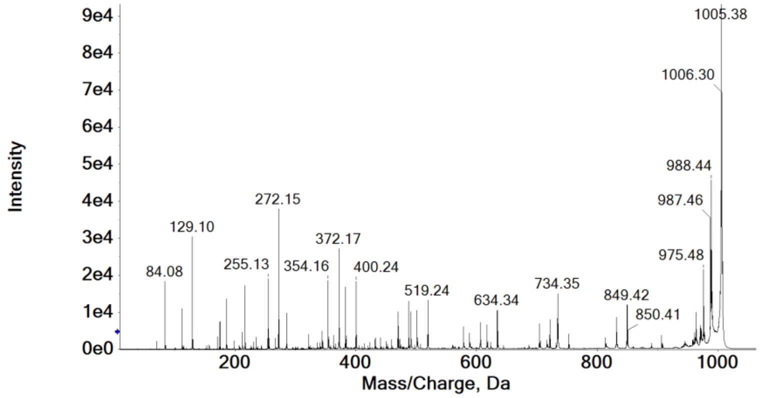


MSMS T1 ac-GGK(Myr)SFSKPR (1215)

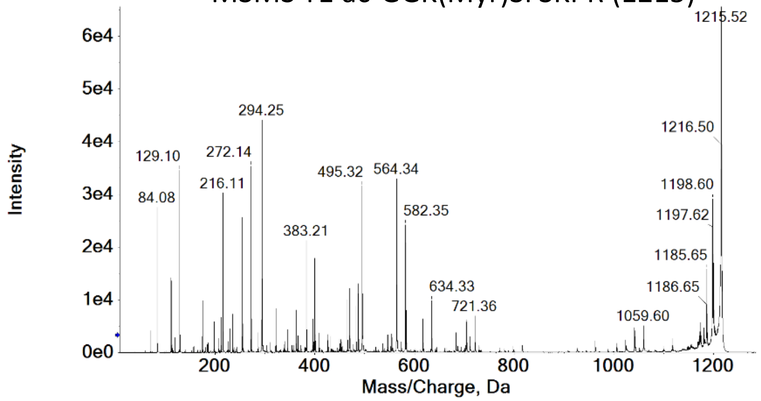

### Ac-GGGKSFSKPR

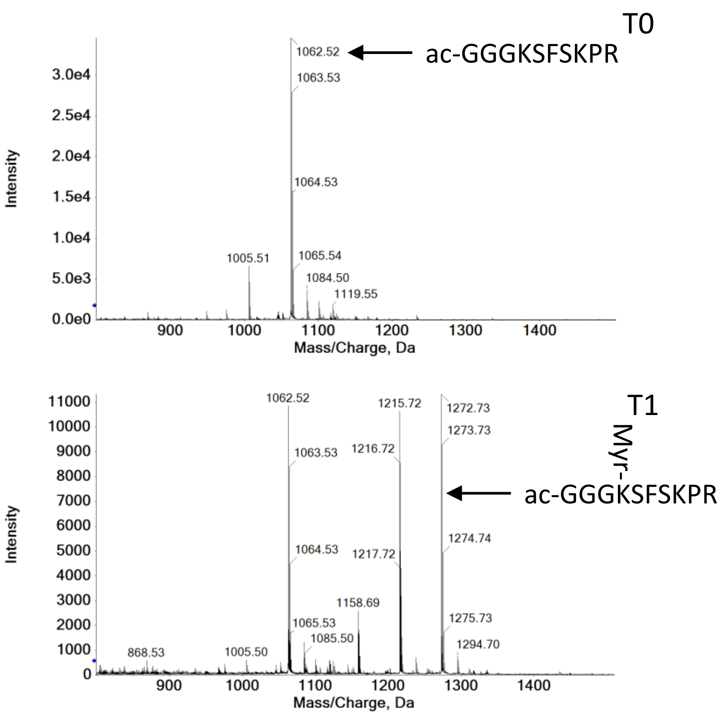


MSMS T0 ac-GGGKSFSKPR (1062)

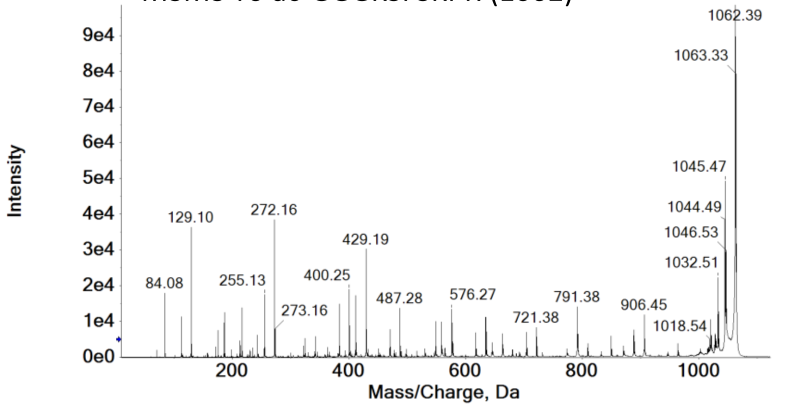


MSMS T1 ac-GGGK(Myr)SFSKPR (1272)

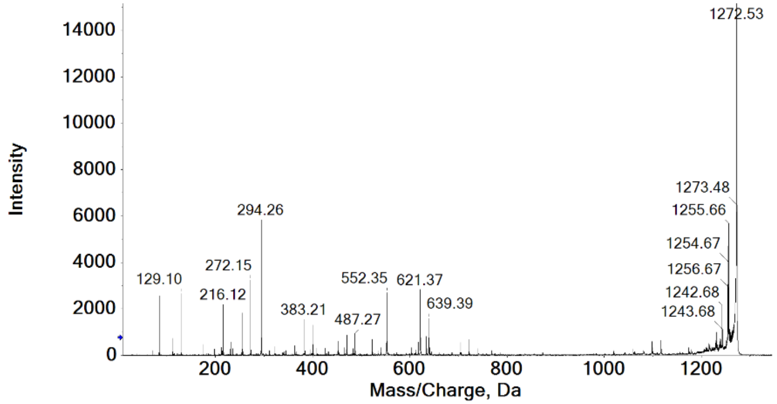

### Ac-GGGGKSFSKPR

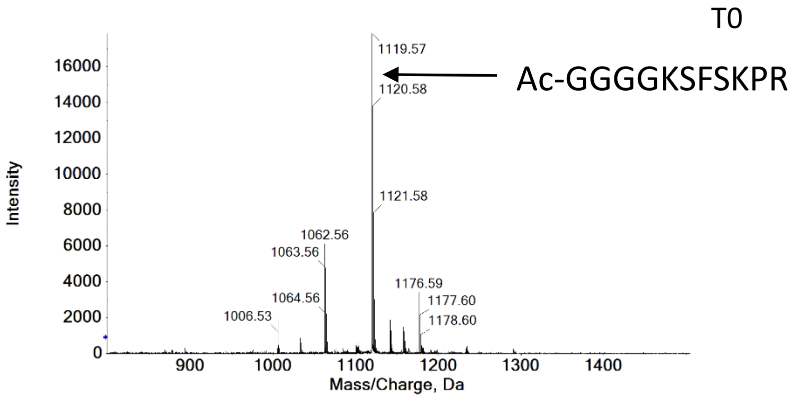


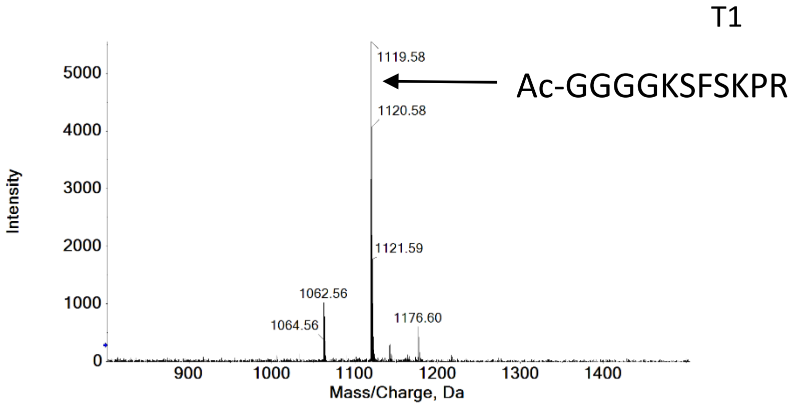

### Ac-GGGGGKSFSKPR

T0

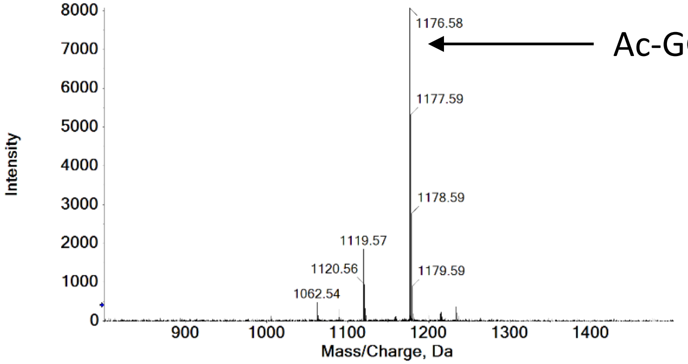


Ac-GGGGGKSFSKPR

T1

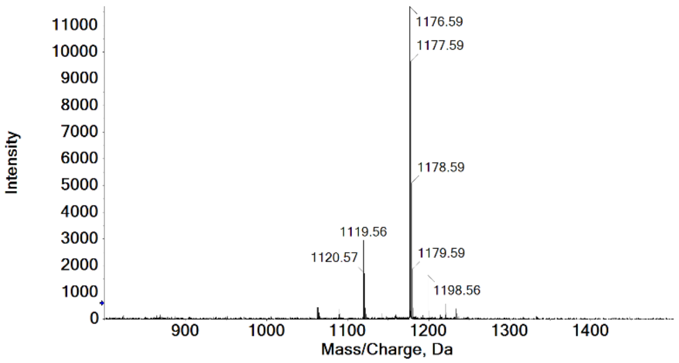
