## Supplementary Dataset 1d for "Proteome-wide probing of the dual NMT-dependent myristoylation tradeoff unveils potent, mechanism-based suicide inhibitors"

### Sup Dataset 1d

| Sequence | Cleavage by IpaJ | MS data in Supplementary Dataset 1e |
|---|---|---|
| Myr-GLSFGKLFSKLFAK | + | 2 |
| Myr-GSMFSGNR | + | 3 |
| Myr-GQGPSGGLNR | + | 4 |
| Myr-GNLHGIHR | + | 5 |
| Myr-GKVLSKIFGNK | + | 6 |
| Myr-GNCFSKPR | + | 7 |
| ac-GNCFSKPR | - | 8 |
| Myr-ATNGSKVA | - | 9 |
| | IpaJ cleavage after MYR | |
| GIGWSLHN | + | 10 |
| GNLHGIHR | + | 11 |
| GCFHSKAA | + | 12 |
| GHRHSKSK | + | 13 |
| GMVFGKIA | + | 14 |
| GGTTSTRR | + | 15 |
| GKVLSKIF | + | 16 |
| GFCFSKFG | + | 17 |
| GRKWSGPT | + | 18 |
| GTSVSKPV | + | 19 |

### Myr-GLSFGKLFSKLFAK

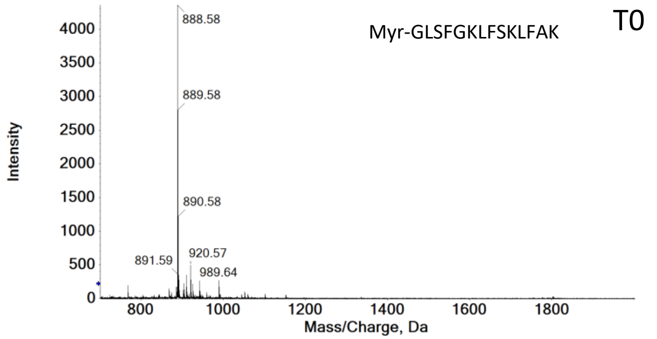


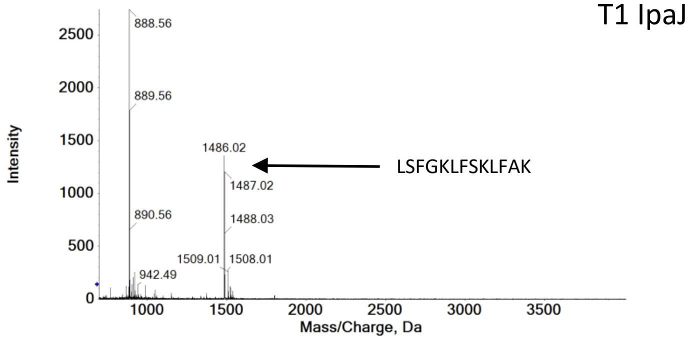

### Myr-GSMFSGNR

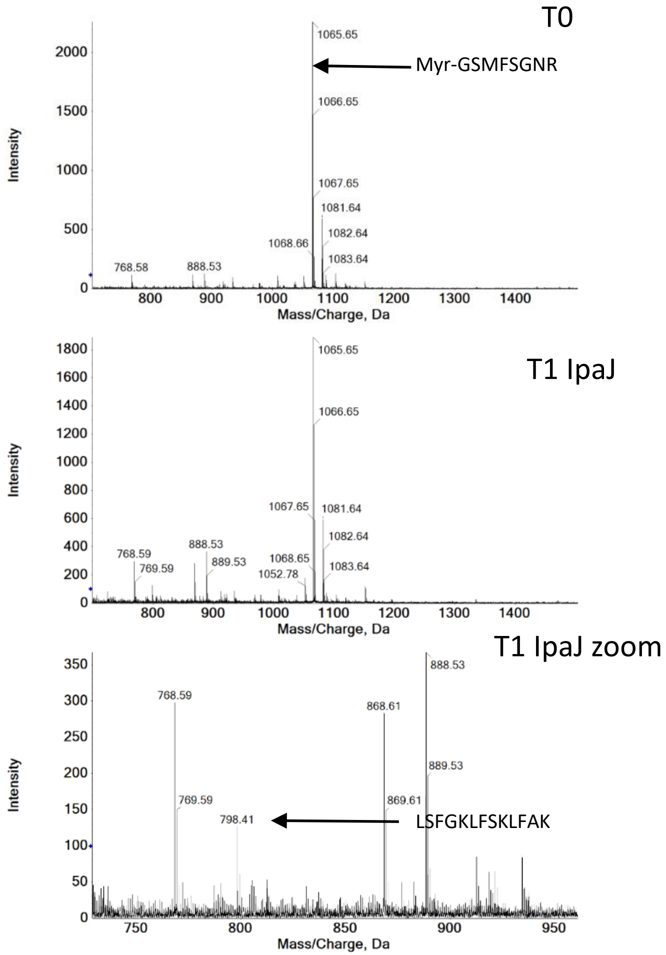

### Myr-GQGPSGGLNR

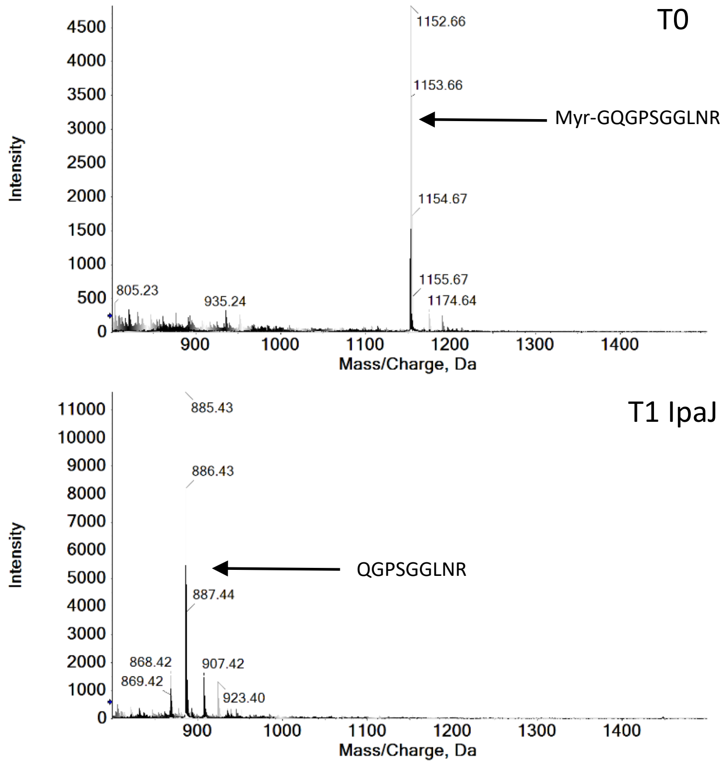

### Myr-GNLHGIHR

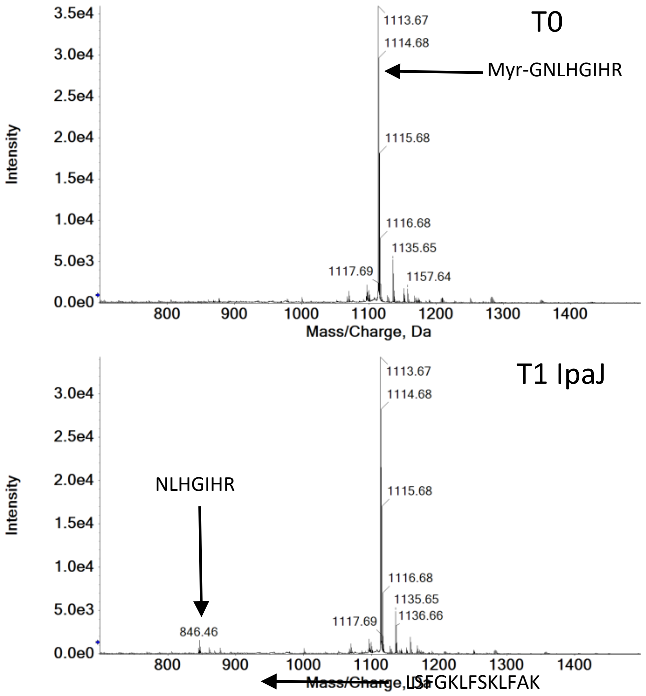

### Myr-GKVLSKIFGNK

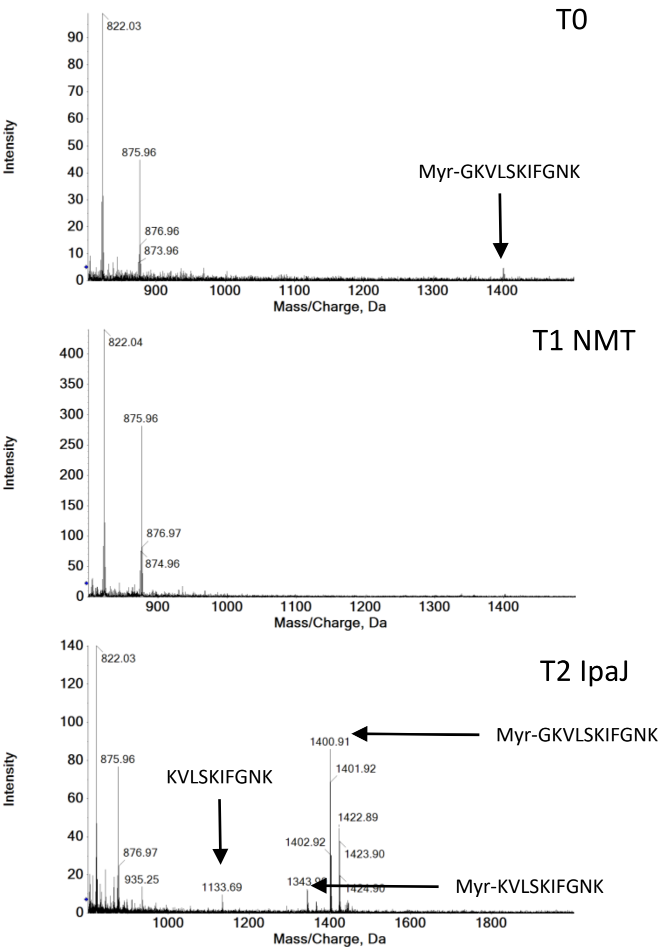

### Myr-GNCFSKPR

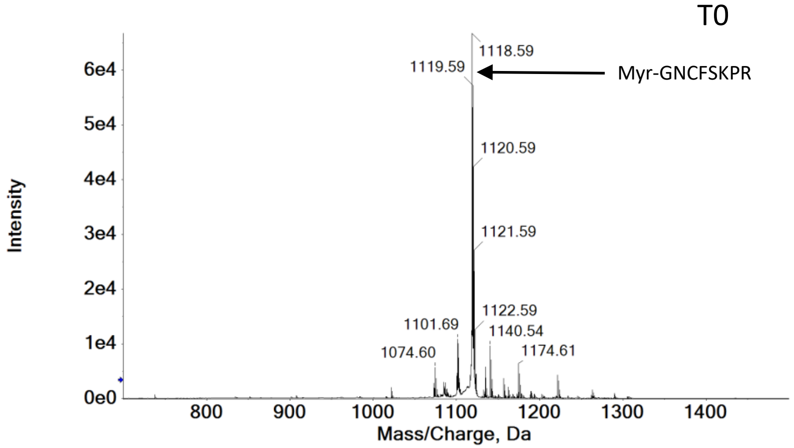


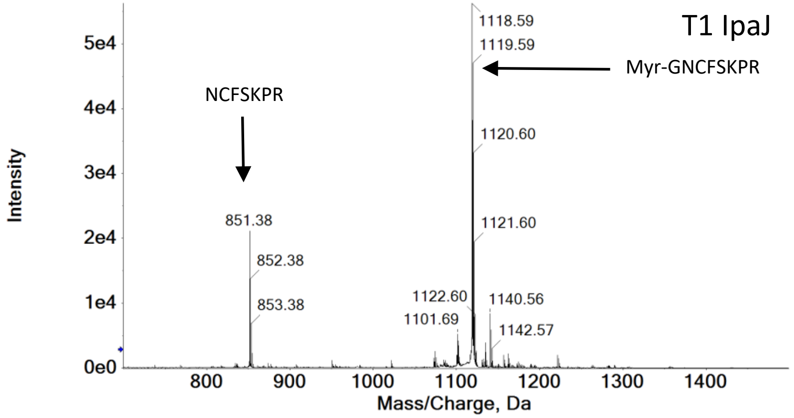

### Ac-GNCFSKPR

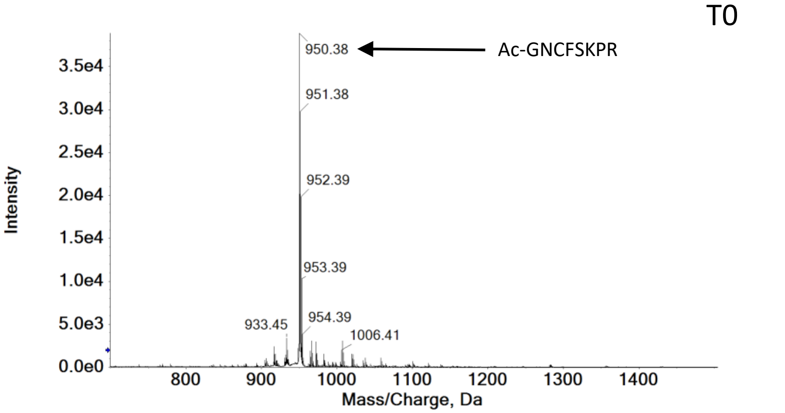
T0
950.38
Ac-GNCFSKPR
951.38
952.39
953.39
933.45
954.39
1006.41
Intensity
3.5e4
3.0e4
2.5e4
2.0e4
1.5e4
1.0e4
5.0e3
0.0e0
800
900
1000
1100
1200
1300
1400
Mass/Charge, Da

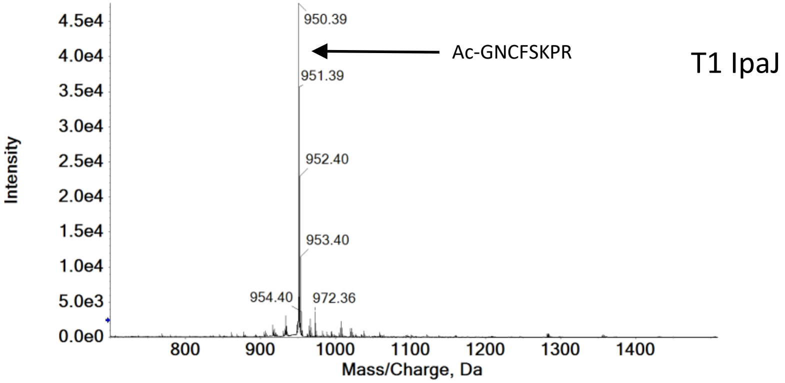
950.39
Ac-GNCFSKPR
T1 IpaJ
951.39
952.40
953.40
954.40
972.36
Intensity
4.5e4
4.0e4
3.5e4
3.0e4
2.5e4
2.0e4
1.5e4
1.0e4
5.0e3
0.0e0
800
900
1000
1100
1200
1300
1400
Mass/Charge, Da

### Myr-ATNGSKVA

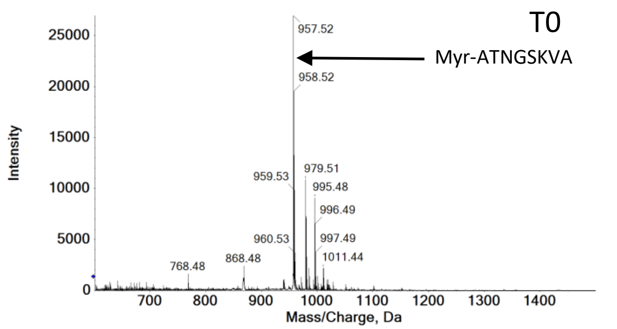


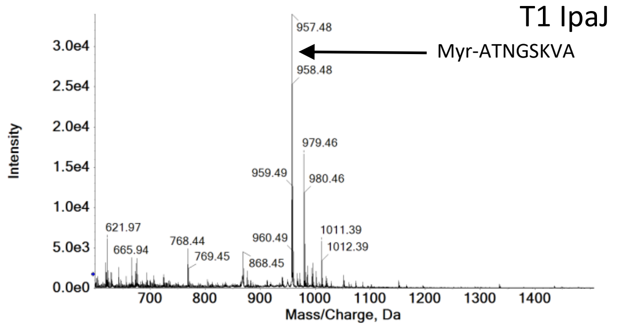


MSMS T0 Myr-ATNGSKVA (957)

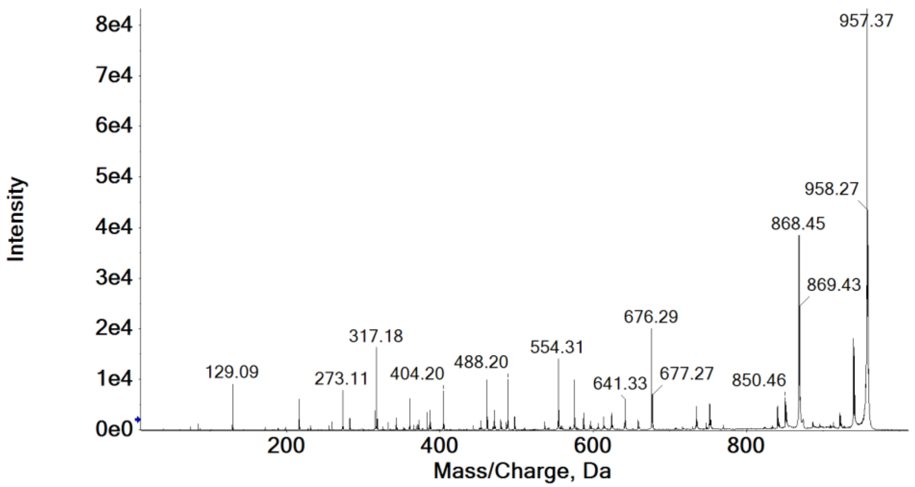

### GIGWSLHN

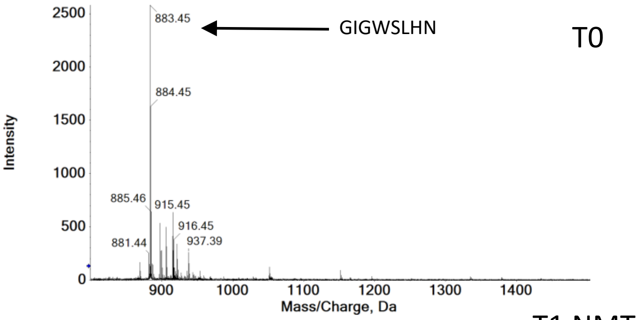
T0
GIGWSLHN
883.45
884.45
885.46
915.45
916.45
881.44
937.39
Intensity
Mass/Charge, Da

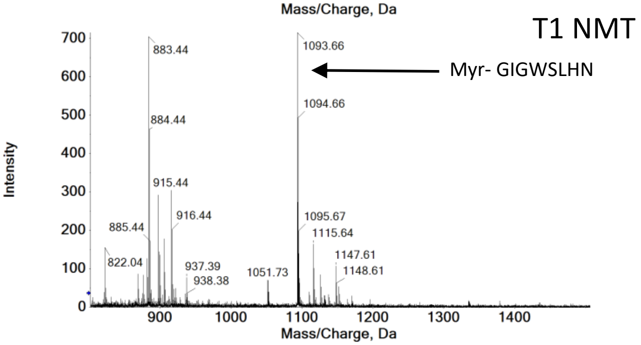
T1 NMT
Myr- GIGWSLHN
883.44
884.44
915.44
916.44
885.44
822.04
937.39
938.38
1051.73
1093.66
1094.66
1095.67
1115.64
1147.61
1148.61
Intensity
Mass/Charge, Da

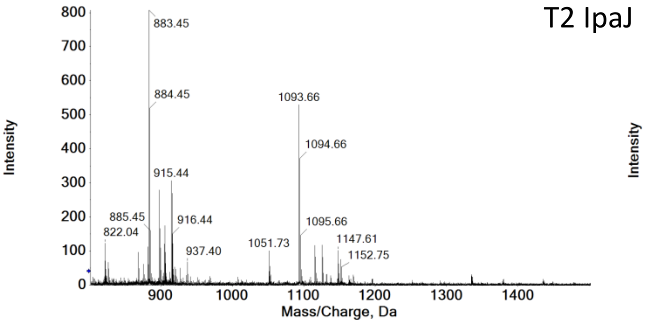
T2 IpaJ
883.45
884.45
915.44
885.45
822.04
916.44
937.40
1051.73
1093.66
1094.66
1095.66
1147.61
1152.75
Intensity
Mass/Charge, Da

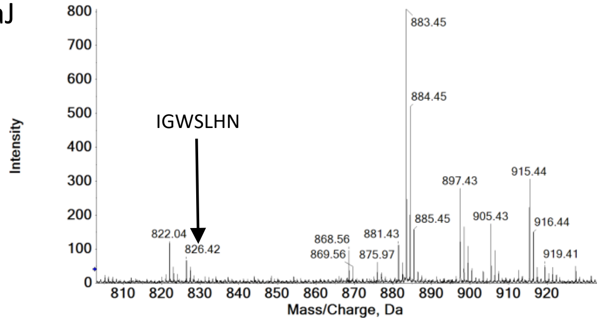
IGWSLHN
822.04
826.42
868.56
869.56
875.97
881.43
883.45
884.45
885.45
897.43
905.43
915.44
916.44
919.41
Intensity
Mass/Charge, Da

### GNLHGIHR

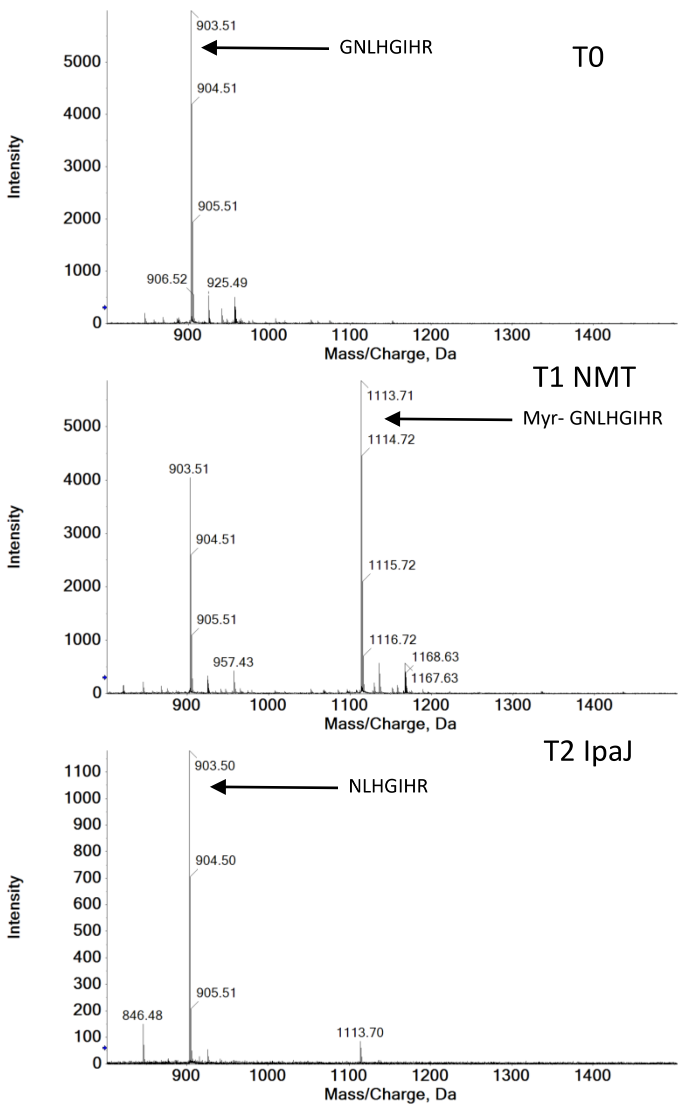

T0
GNLHGIHR
903.51
904.51
905.51
906.52
925.49
Intensity
Mass/Charge, Da
T1 NMT
Myr- GNLHGIHR
1113.71
1114.72
1115.72
1116.72
1168.63
1167.63
903.51
904.51
905.51
957.43
T2 IpaJ
NLHGIHR
903.50
904.50
905.51
846.48
1113.70

### GCFHSKAA

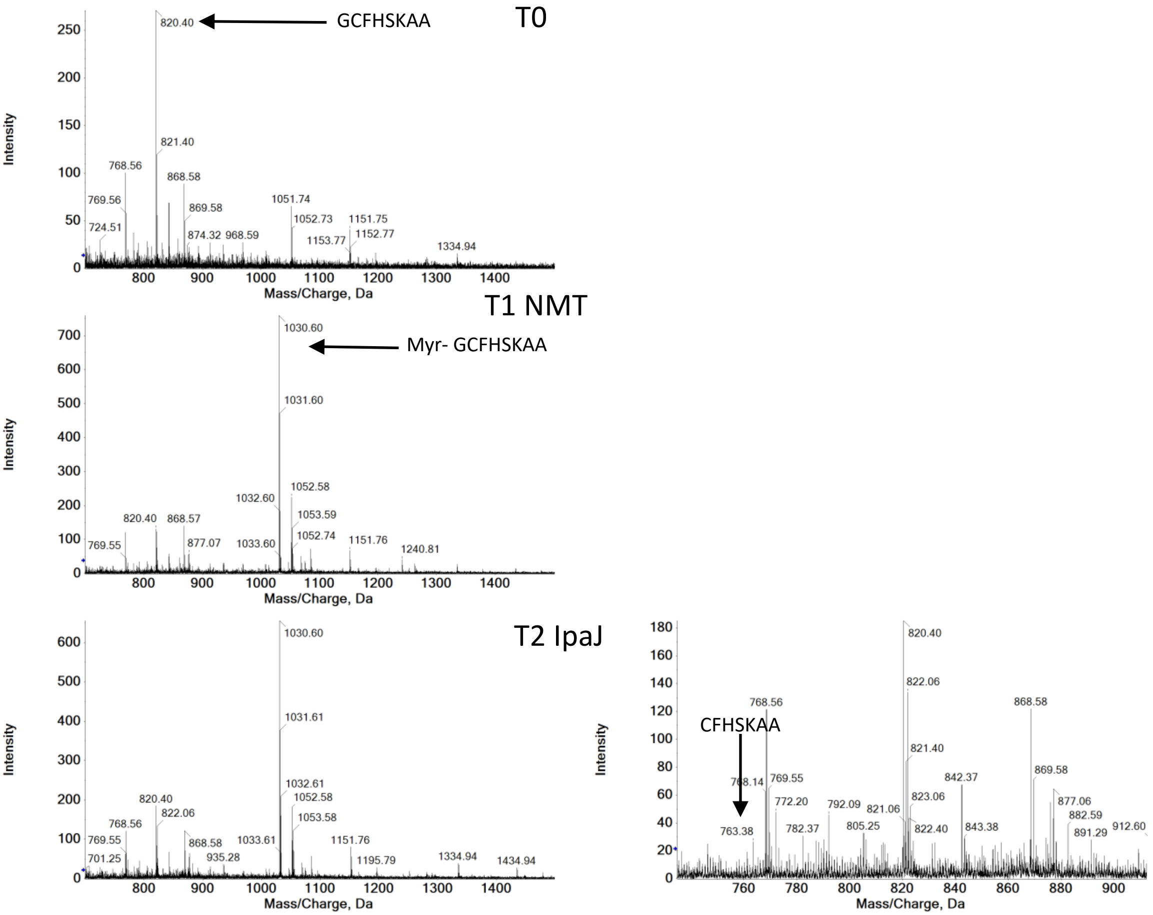
T0
GCFHSKAA
820.40
821.40
768.56
868.58
769.56
869.58
724.51
874.32
968.59
1051.74
1052.73
1153.77
1151.75
1152.77
1334.94
T1 NMT
Myr- GCFHSKAA
1030.60
1031.60
1032.60
1052.58
1053.59
1052.74
1033.60
820.40
868.57
769.55
877.07
1151.76
1240.81
T2 IpaJ
1030.60
1031.61
1032.61
1052.58
1053.58
1033.61
820.40
822.06
768.56
769.55
701.25
868.58
935.28
1151.76
1195.79
1334.94
1434.94
CFHSKAA
820.40
822.06
821.40
823.06
822.40
768.56
769.55
768.14
763.38
772.20
782.37
792.09
805.25
821.06
842.37
843.38
868.58
869.58
877.06
882.59
891.29
912.60
Intensity
Mass/Charge, Da

### GHRHSKSK

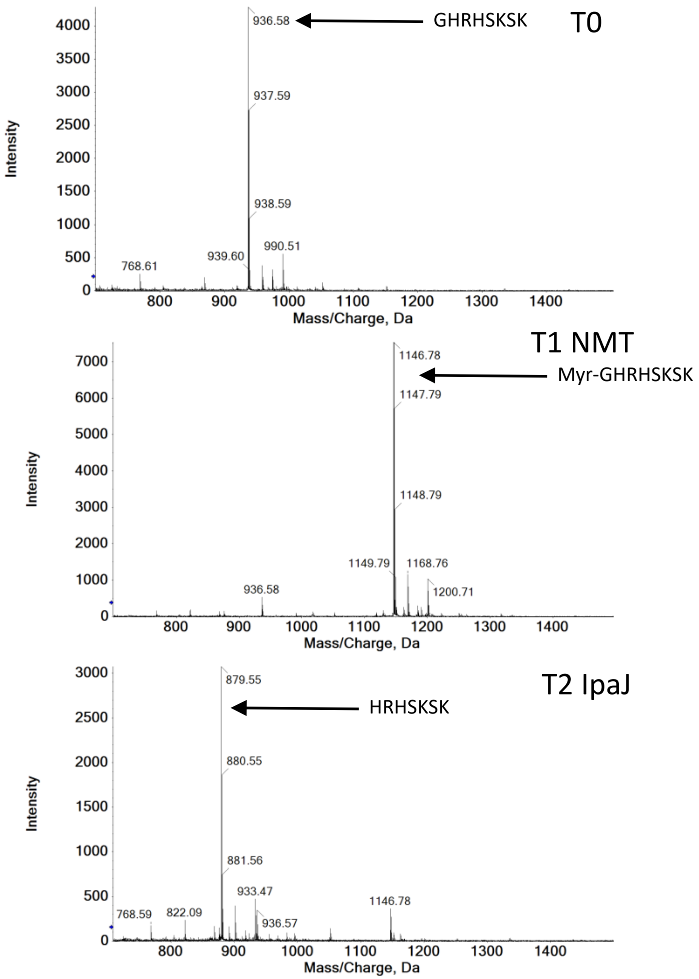

T0
GHRHSKSK
936.58
937.59
938.59
939.60
990.51
768.61
Intensity
Mass/Charge, Da
T1 NMT
Myr-GHRHSKSK
1146.78
1147.79
1148.79
1149.79
1168.76
1200.71
936.58
Intensity
Mass/Charge, Da
T2 IpaJ
HRHSKSK
879.55
880.55
881.56
933.47
936.57
1146.78
768.59
822.09
Intensity
Mass/Charge, Da

### GMVFGKIA

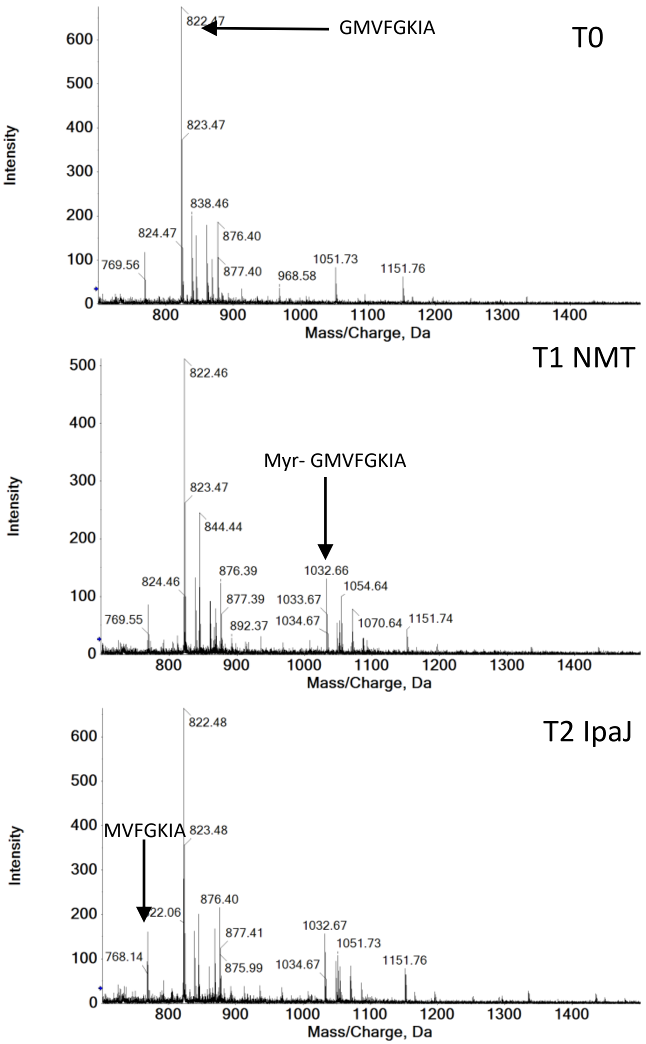

T0
GMVFGKIA
822.47
823.47
838.46
824.47
876.40
769.56
877.40
968.58
1051.73
1151.76
T1 NMT
822.46
Myr- GMVFGKIA
823.47
844.44
876.39
1032.66
824.46
1054.64
877.39
1033.67
769.55
892.37
1034.67
1070.64
1151.74
T2 IpaJ
822.48
MVFGKIA
823.48
876.40
1032.67
877.41
1051.73
768.14
1034.67
875.99
1151.76
Intensity
Mass/Charge, Da

### GGTTSTRR

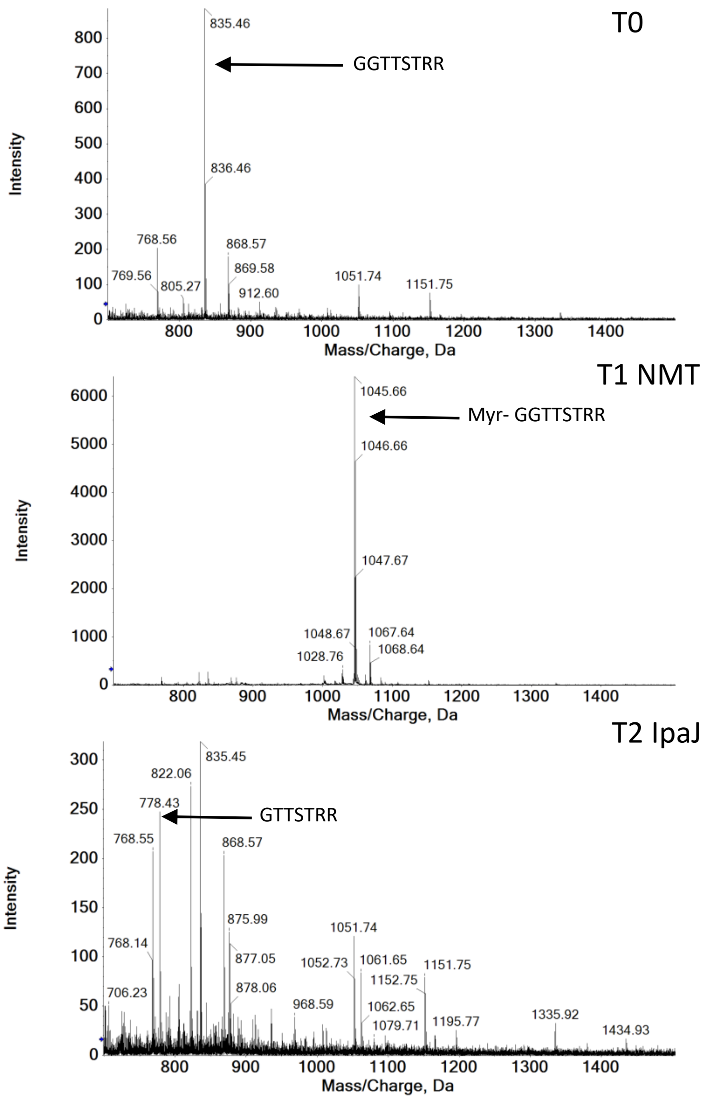
T0
GGTTSTRR
835.46
836.46
768.56
868.57
769.56
805.27
869.58
912.60
1051.74
1151.75
Intensity
Mass/Charge, Da
T1 NMT
Myr- GGTTSTRR
1045.66
1046.66
1047.67
1048.67
1067.64
1028.76
1068.64
T2 IpaJ
GTTSTRR
835.45
822.06
778.43
768.55
868.57
875.99
768.14
877.05
878.06
706.23
968.59
1051.74
1052.73
1061.65
1151.75
1152.75
1062.65
1079.71
1195.77
1335.92
1434.93

### GKVLSKIF

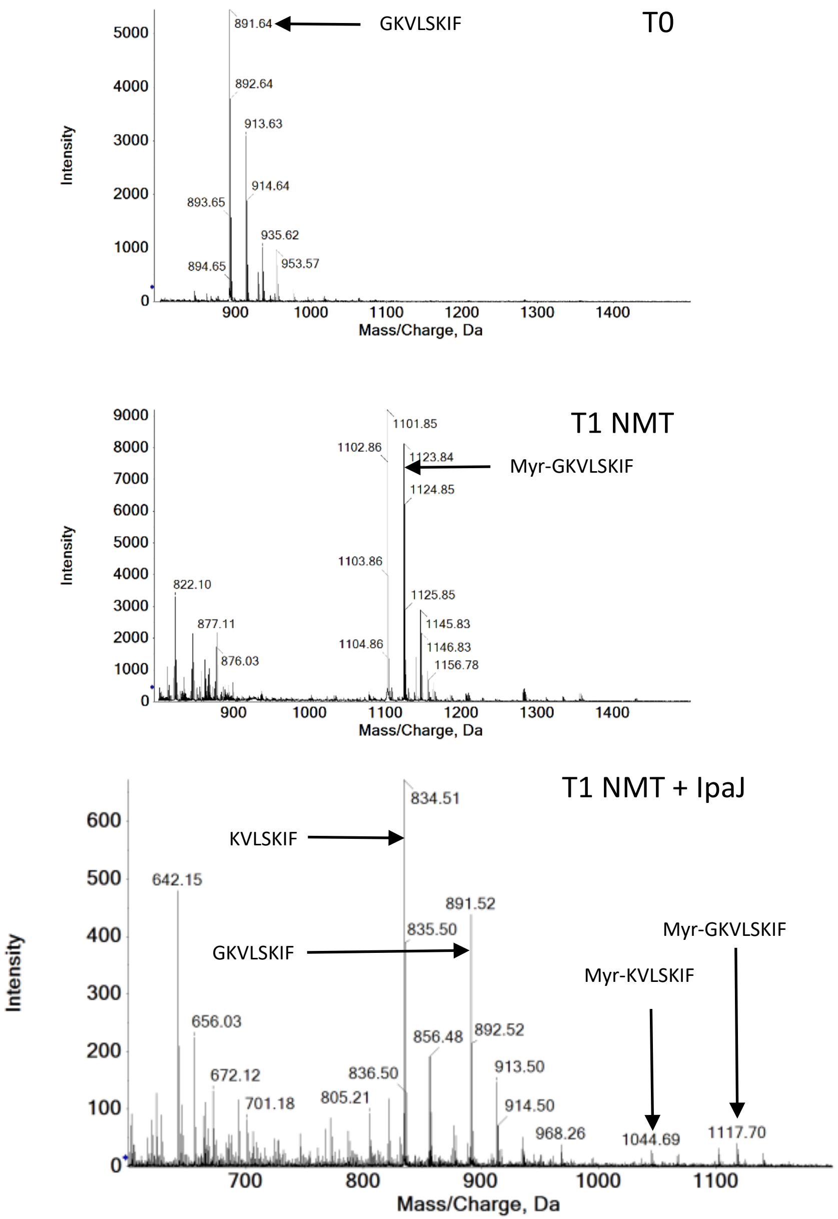
T0
GKVLSKIF
891.64
892.64
913.63
914.64
893.65
935.62
953.57
894.65
Intensity
Mass/Charge, Da
T1 NMT
Myr-GKVLSKIF
1101.85
1102.86
1123.84
1124.85
1103.86
1125.85
1145.83
1146.83
1156.78
1104.86
822.10
877.11
876.03
T1 NMT + IpaJ
KVLSKIF
GKVLSKIF
Myr-KVLSKIF
Myr-GKVLSKIF
834.51
642.15
891.52
835.50
856.48
892.52
656.03
913.50
672.12
836.50
914.50
701.18
805.21
968.26
1044.69
1117.70

### GFCFSKFG

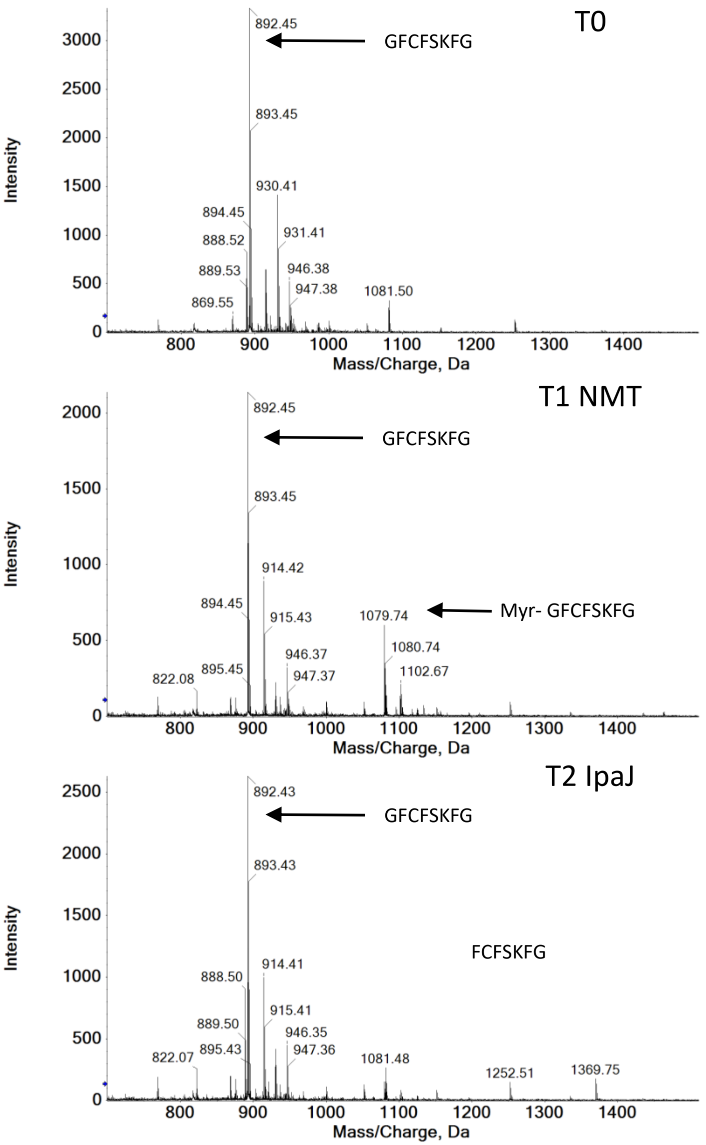
T0
GFCFSKFG
892.45
893.45
930.41
894.45
888.52
931.41
889.53
946.38
947.38
1081.50
869.55
Intensity
Mass/Charge, Da
T1 NMT
GFCFSKFG
892.45
893.45
914.42
894.45
915.43
Myr- GFCFSKFG
1079.74
1080.74
946.37
947.37
1102.67
822.08
895.45
T2 IpaJ
GFCFSKFG
892.43
893.43
FCFSKFG
914.41
888.50
915.41
889.50
946.35
895.43
947.36
822.07
1081.48
1252.51
1369.75

### GRKWSGPT

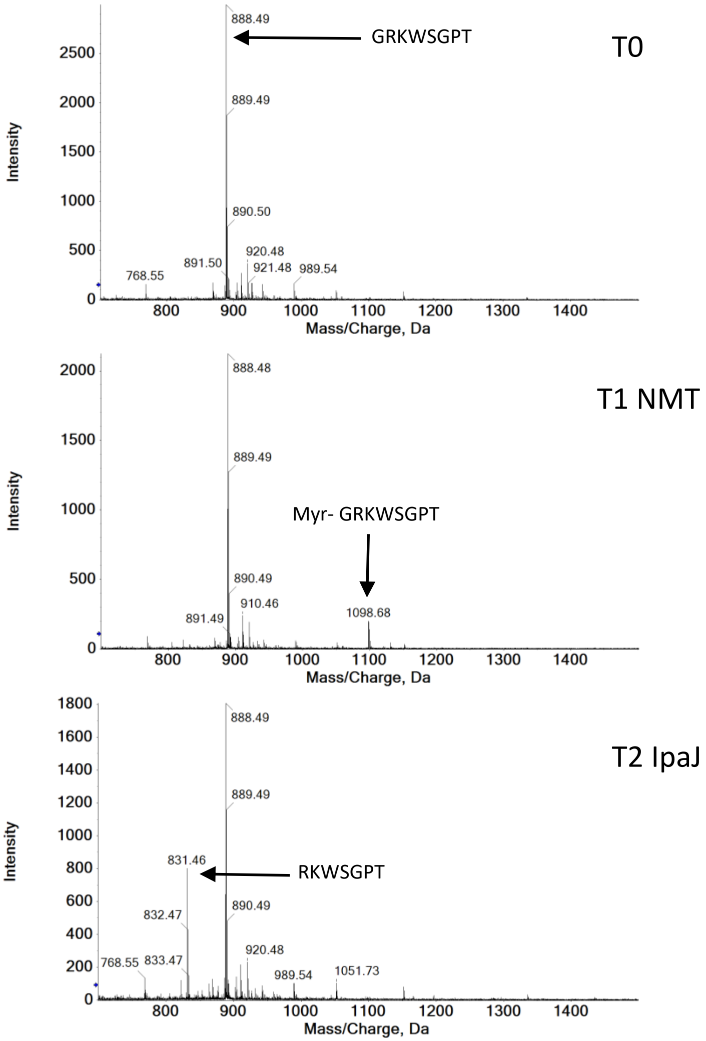

T0
GRKWSGPT
888.49
889.49
890.50
891.50
920.48
921.48
989.54
768.55
Intensity
Mass/Charge, Da
T1 NMT
888.48
889.49
890.49
891.49
910.46
Myr- GRKWSGPT
1098.68
T2 IpaJ
888.49
889.49
890.49
831.46
832.47
833.47
RKWSGPT
768.55
920.48
989.54
1051.73

### GTSVSKPV

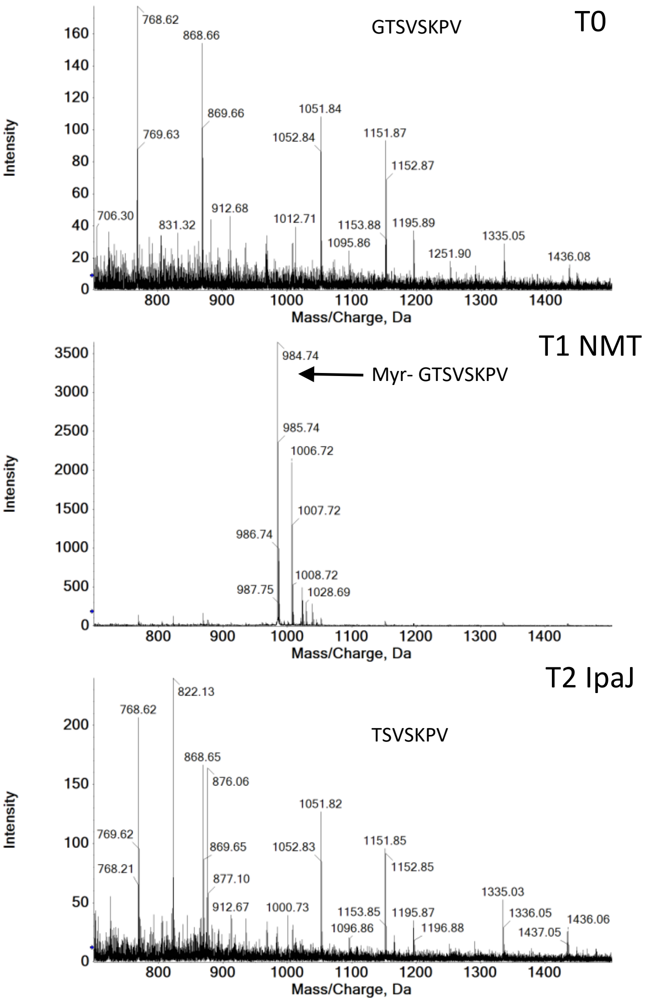

T0
GTSVSKPV
768.62
868.66
869.66
1051.84
769.63
1052.84
1151.87
1152.87
706.30
831.32
912.68
1012.71
1153.88
1095.86
1195.89
1251.90
1335.05
1436.08
Intensity
Mass/Charge, Da
T1 NMT
984.74
Myr- GTSVSKPV
985.74
1006.72
1007.72
986.74
1008.72
987.75
1028.69
T2 IpaJ
822.13
768.62
TSVSKPV
868.65
876.06
1051.82
769.62
869.65
1052.83
1151.85
768.21
1152.85
877.10
1335.03
912.67
1000.73
1153.85
1195.87
1336.05
1436.06
1096.86
1196.88
1437.05
