## Supplementary Dataset 1e for "Proteome-wide probing of the dual NMT-dependent myristoylation tradeoff unveils potent, mechanism-based suicide inhibitors"

### Sup Dataset 1e

| Peptide sequence | MS data in Supplementary |
|---|---|
| met-GNCFSKPR | 2 |
| GKSFSKPR | 3 |
| G{ORN}SFSKPR | 4 |
| GKVLSKIFGNK | 5/6/7/8 |
| GKVLSKIFGN | 5/6/7/8 |
| GKVLSKIF | 5/6/7/8 |
| Myr-GKVLSKIFGNK | 9 |
| G-K(ε-MYR)VLSKIFGNK | 10 |
| GKFSSKPR | 11 |
| GKQNSKLR | 12 |
| KQNSKLRP | 13 |
| GKQNSKLA | 14 |
| GKSNSKLK | 15 |
| GKTNSKLA | 16 |
| GKSASKQF | 17 |
| GAKQSGPA | 18 |
| GGKQSTAA | 19 |
| GGKLSKKK | 20 |
| GGKWSKLS | 21 |
| AKSFSKPR | 22 |
| GKSWSKGR | 23 |
| AKQNSKLR | 24 |
| GGKQSKAA | 25 |
| GGKSFSKP | 26 |
| GGKFSSKPR | 27/28 |
| AKPTSKDSGLK | 29 |
| AGKFSKPR | 30 |
| AGKSFSKPR | 31 |

### Met-GNCFSKPR

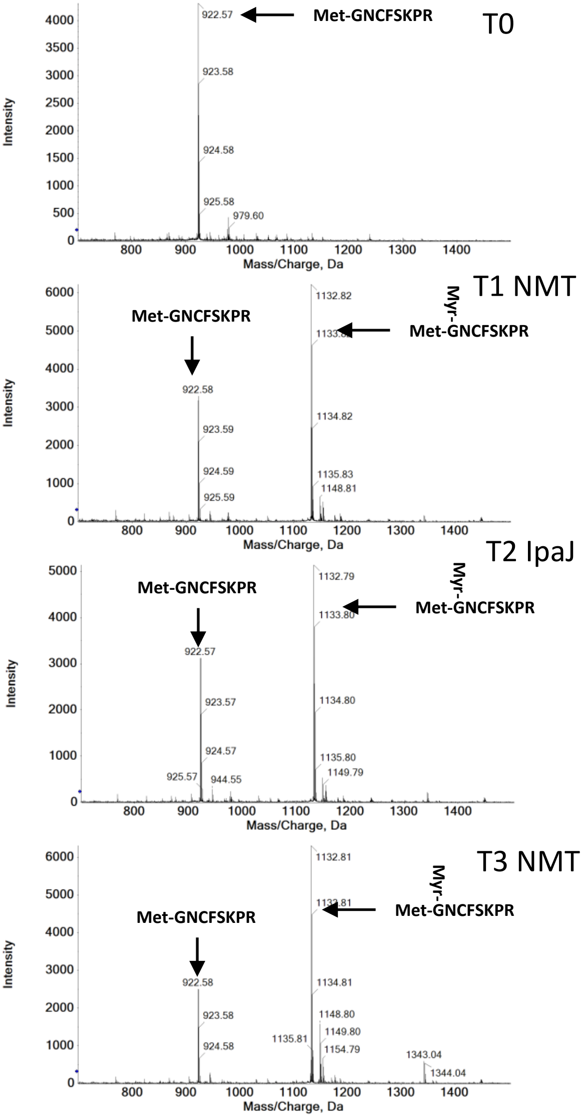


MSMS T1 Met-G(Myr)NCFSKPR (1132)

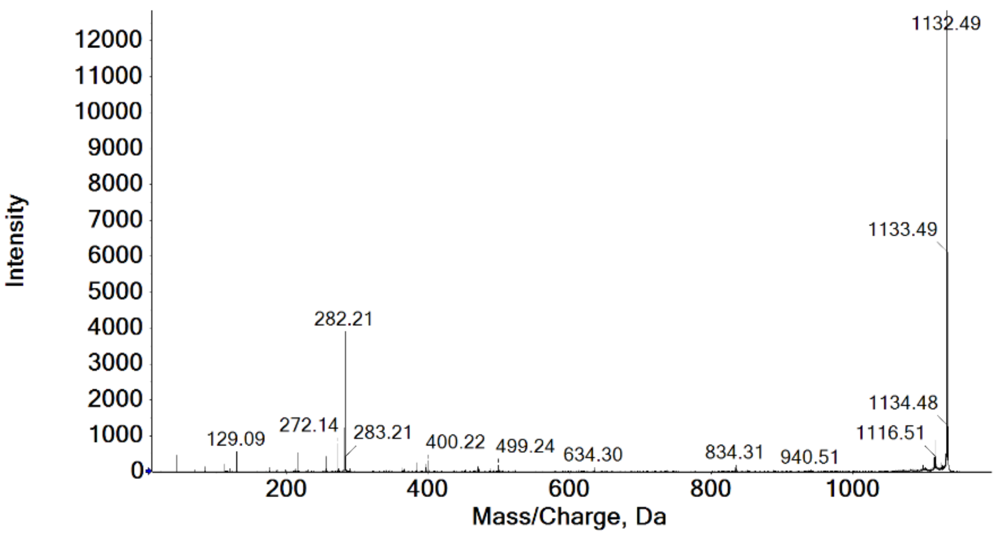

### GKSFSKPR

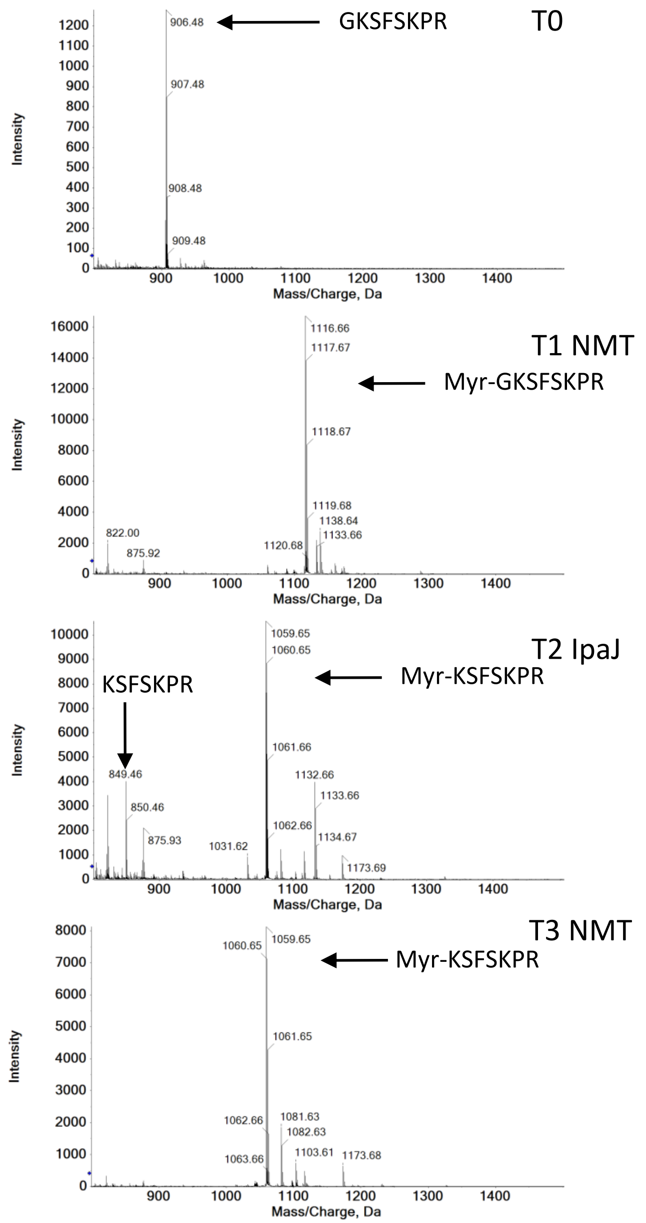

GKSFSKPR
T0
906.48
907.48
908.48
909.48
T1 NMT
Myr-GKSFSKPR
1116.66
1117.67
1118.67
1119.68
1138.64
1133.66
1120.68
822.00
875.92
T2 IpaJ
KSFSKPR
Myr-KSFSKPR
1059.65
1060.65
1061.66
1132.66
1133.66
849.46
850.46
875.93
1031.62
1062.66
1134.67
1173.69
T3 NMT
Myr-KSFSKPR
1060.65
1059.65
1061.65
1062.66
1081.63
1082.63
1063.66
1103.61
1173.68
Intensity
Mass/Charge, Da


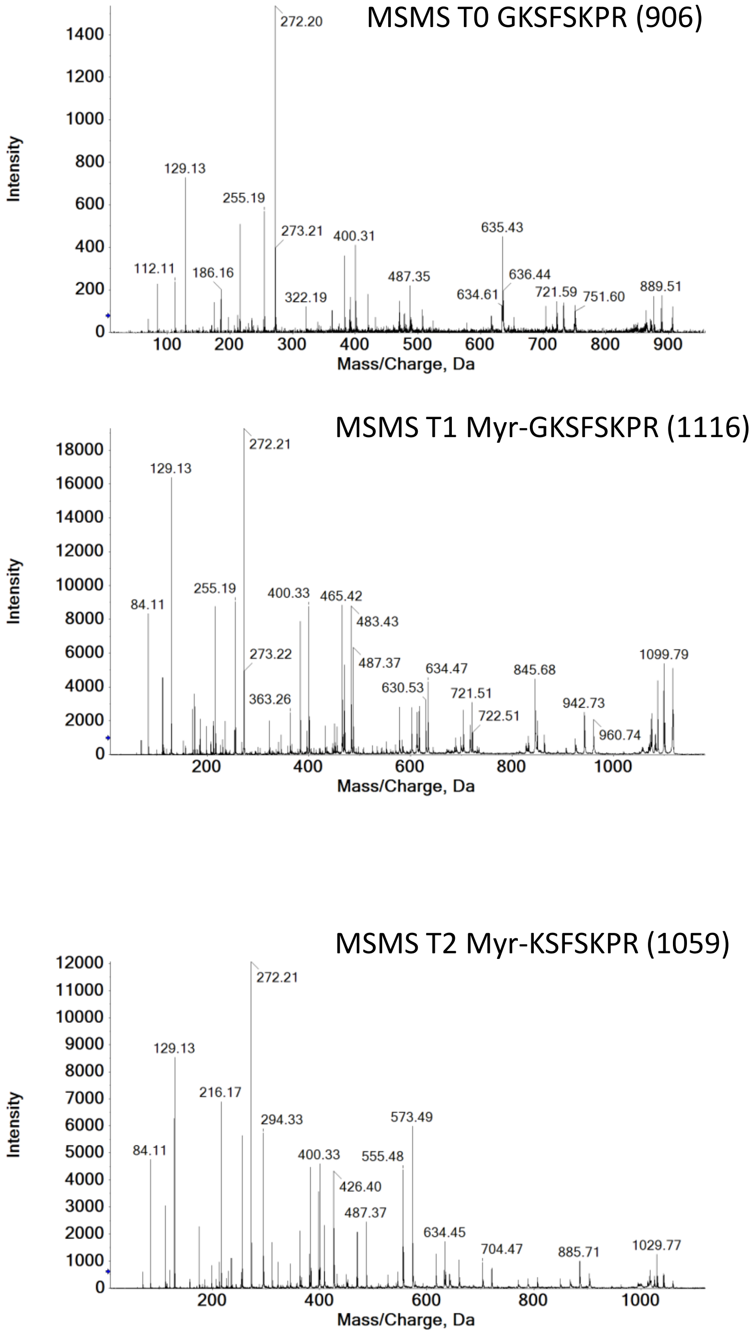

MSMS T0 GKSFSKPR (906)
272.20
129.13
255.19
273.21
400.31
635.43
112.11
186.16
487.35
636.44
322.19
634.61
721.59
751.60
889.51
MSMS T1 Myr-GKSFSKPR (1116)
272.21
129.13
255.19
400.33
465.42
483.43
84.11
273.22
487.37
634.47
845.68
1099.79
363.26
630.53
721.51
722.51
942.73
960.74
MSMS T2 Myr-KSFSKPR (1059)
272.21
129.13
216.17
294.33
573.49
84.11
400.33
555.48
426.40
487.37
634.45
704.47
885.71
1029.77
Intensity
Mass/Charge, Da

### G{ORN}SFSKPR

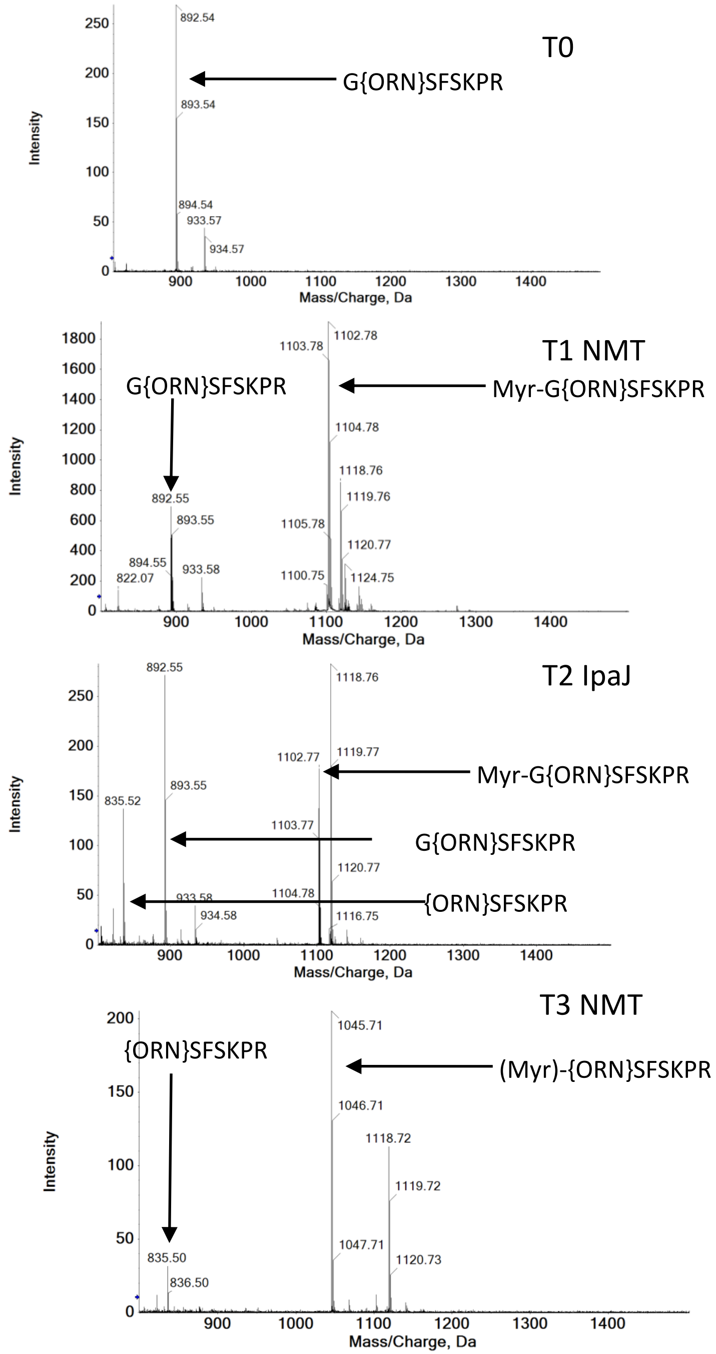


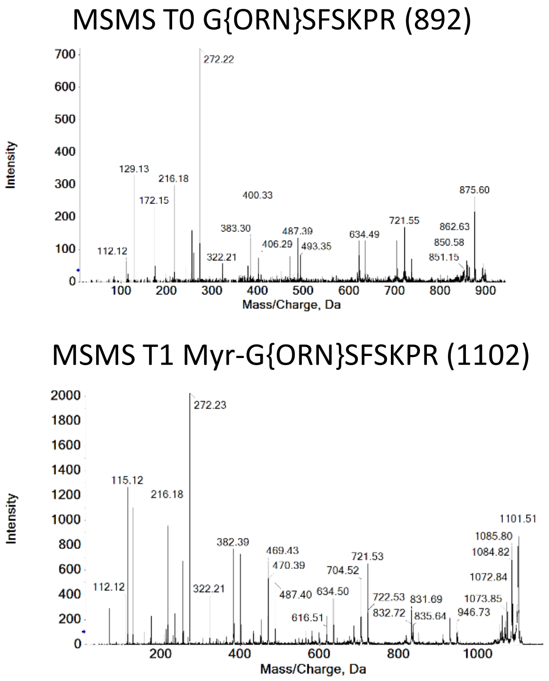


| | Mass/Charge (Da) | Intensity | ion |
|---|---|---|---|
| 875 | 175,1545 | 170,5098 | y1 |
| 1983 | 272,224 | 1113,882 | y2 |
| 4152 | 400,3378 | 282,5098 | y3 |
| 4407 | 412,3786 | 371,6078 | b2 |
| 5781 | 487,3956 | 134,5882 | y4 |
| 6965 | 559,4784 | 496 | b3 |
| 8430 | 634,4864 | 97,41177 | y5 |
| 8686 | 646,5124 | 76,39216 | b4 |
| 9878 | 721,5168 | 61,96078 | y6 |
| 11989 | 871,7861 | 56,62745 | b6 |

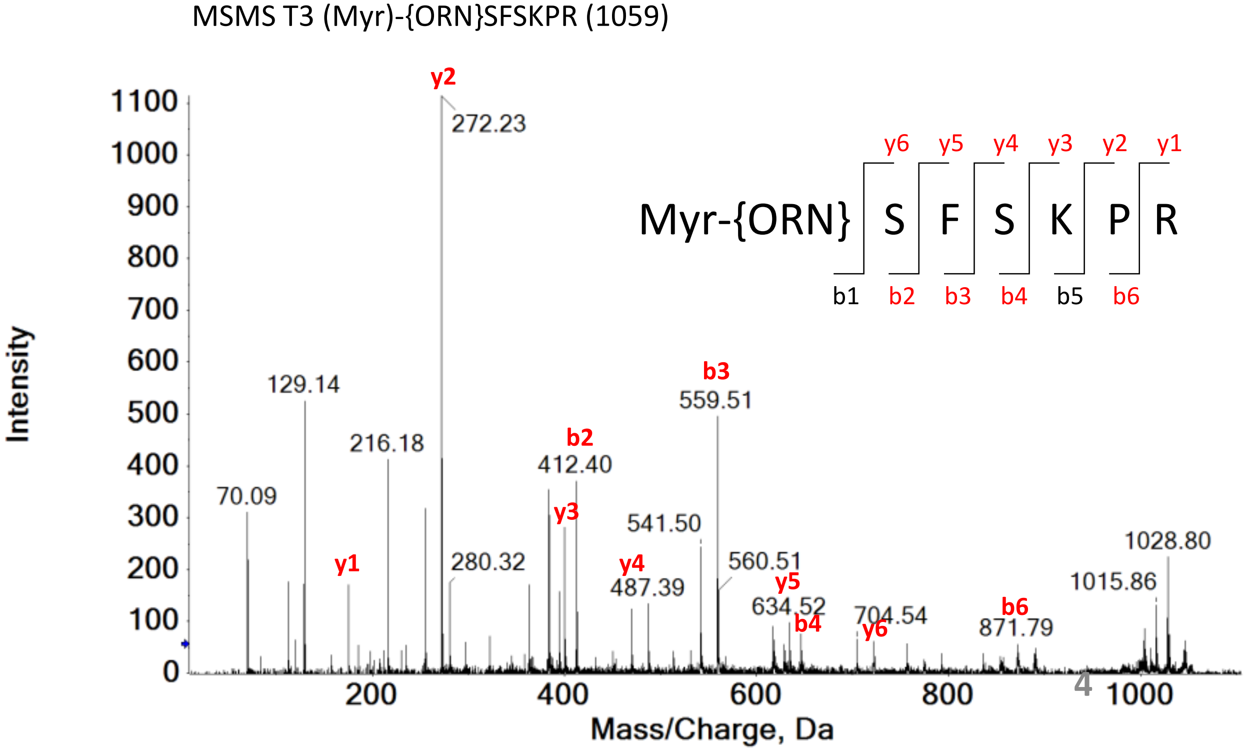

### ARF6 +/- Long NMT 1

Final point after NMT IpaJ NMT

#### KVLSKIF

T1 NMT + IpaJ + NMT

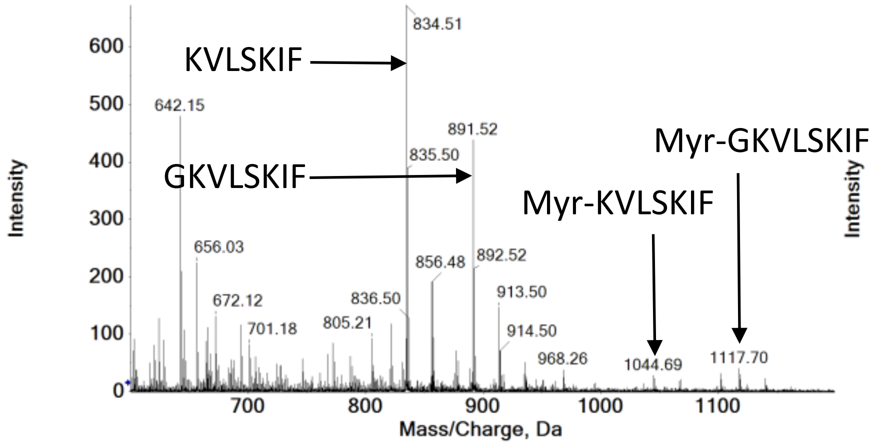


#### GKVLSKIFGNK

T1 NMT + IpaJ + NMT

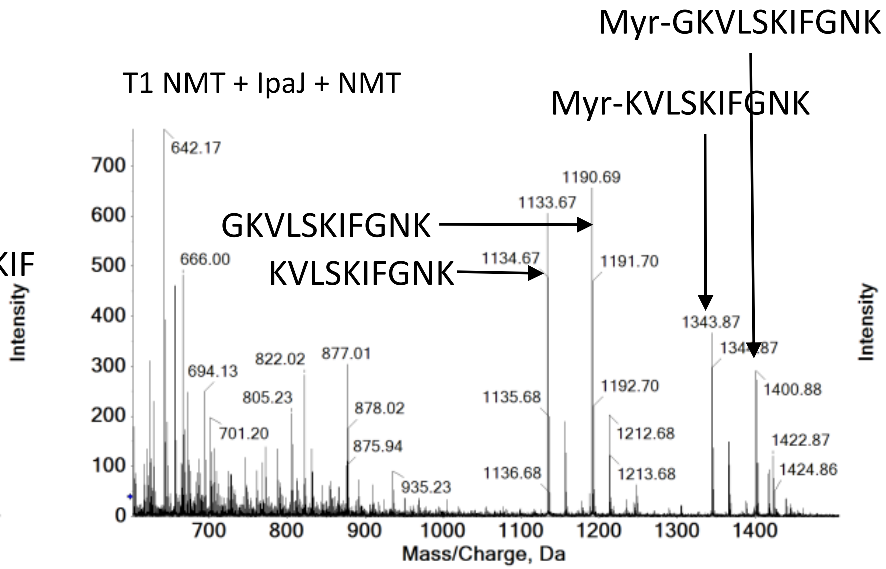


#### GKVLSKIFGN

T1 NMT + IpaJ + NMT

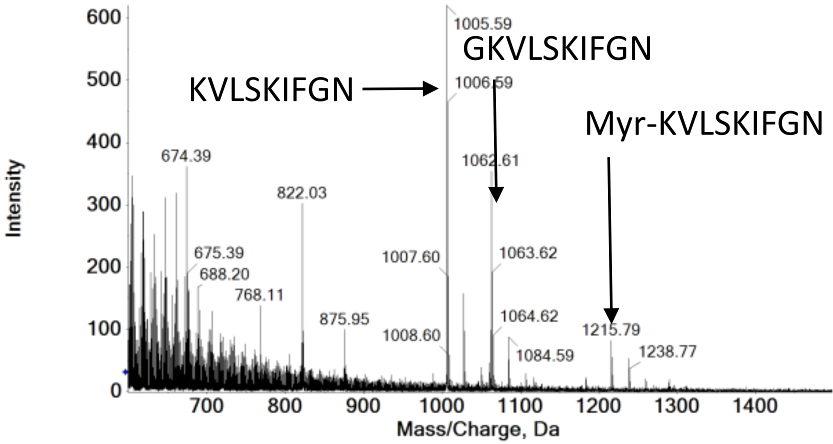

| E24_MSMS_1343.9000_2 | | | ion |
|---|---|---|---|
| pic | Mass/Charge (Da) | Intensity | ion |
| 2965 | 438,4012 | 270,4314 | b2 |
| 4415 | 551,5016 | 387,9216 | b3 |
| 5943 | 638,5012 | 113,7255 | b4 |
| 8104 | 766,6613 | 253,8039 | b5 |
| 10210 | 879,7369 | 1377,098 | b6 |
| 12772 | 1026,816 | 591,8431 | b7 |
| 13697 | 1083,844 | 281,4118 | b8 |
| 15474 | 1197,918 | 197,8039 | b9 |
| 3370 | 465,2638 | 324,8628 | y4 |
| 4874 | 578,3805 | 445,6471 | y5 |
| 7099 | 706,5034 | 506,9804 | y6 |
| 8686 | 793,5415 | 932,7059 | y7 |
| 10753 | 906,6525 | 703,3726 | y8 |
| 12389 | 1005,731 | 115,451 | y9 |

MSMS T2 Myr-KVLSKIFGNK (1343)

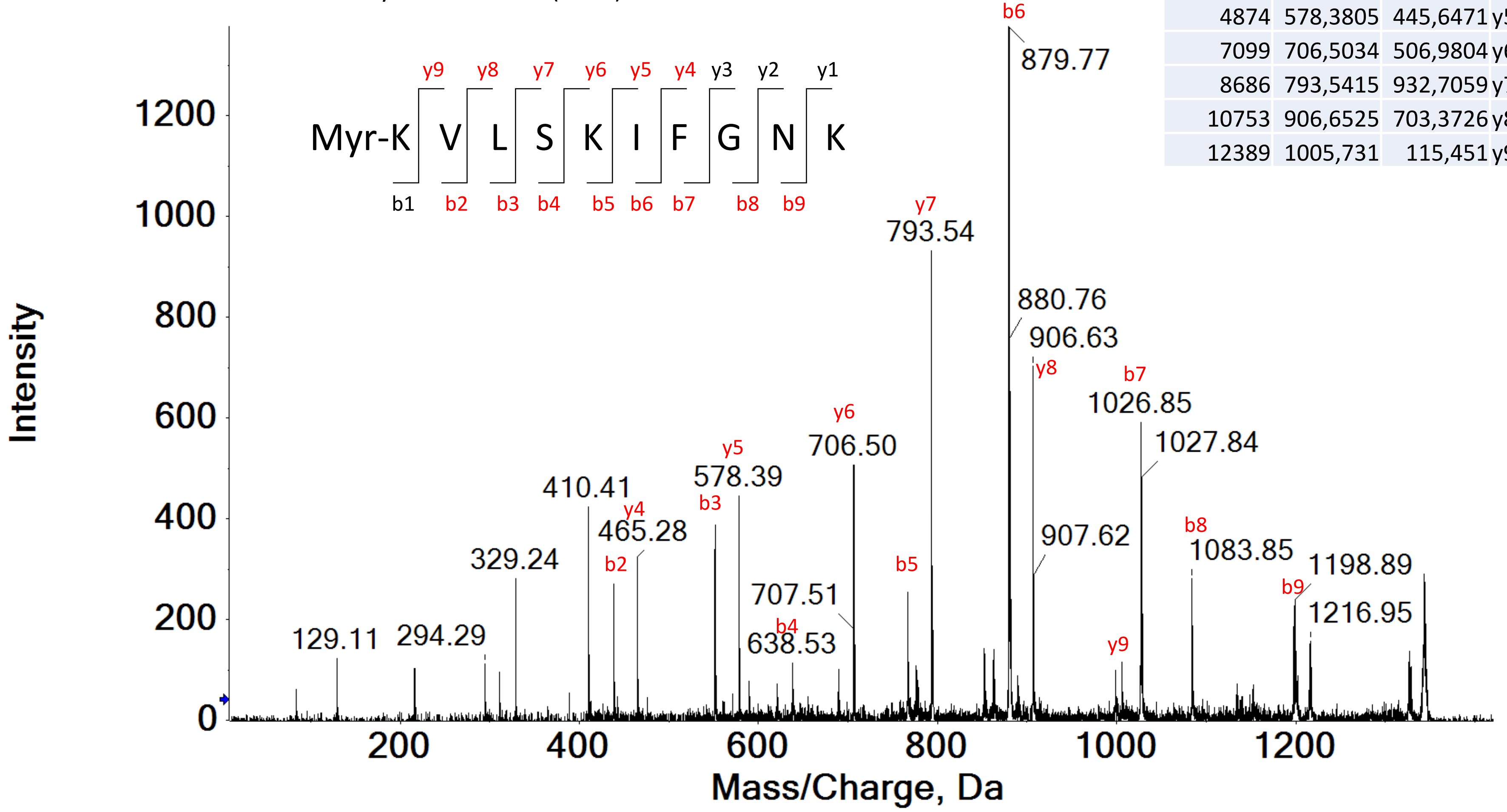

### ARF6 +/- Long NMT 2

Final point after NMT IpaJ NMT

KVLSKIF

T1 NMT + IpaJ + NMT

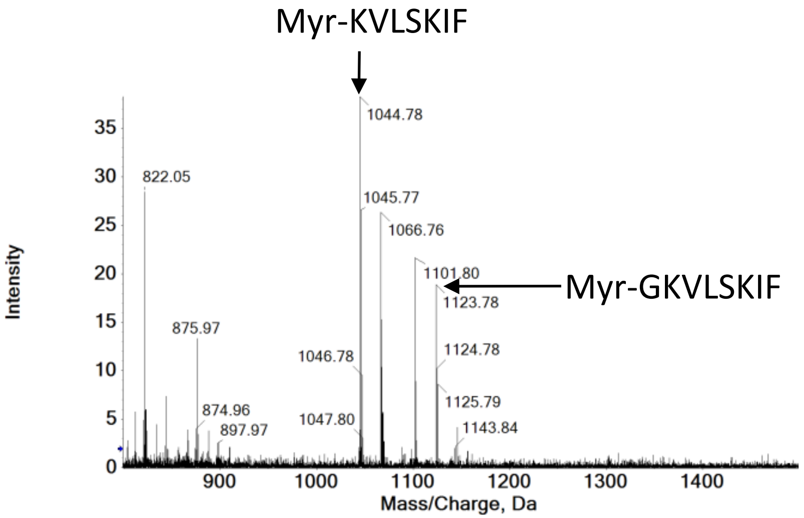


GKVLSKIFGNK

T1 NMT + IpaJ + NMT

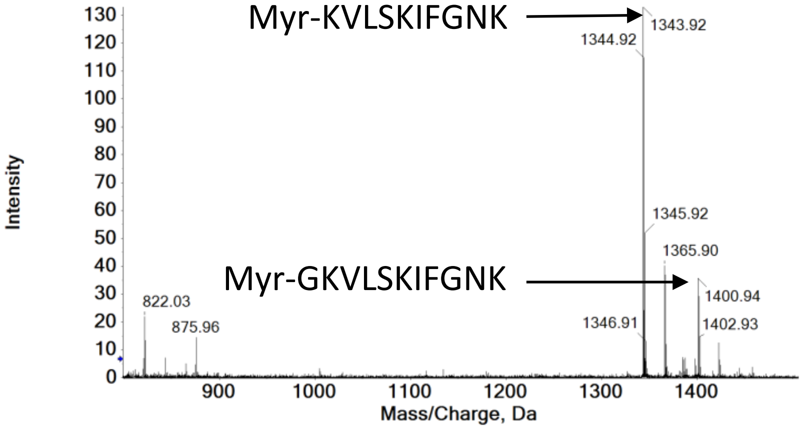


GKVLSKIFGN

T1 NMT + IpaJ + NMT

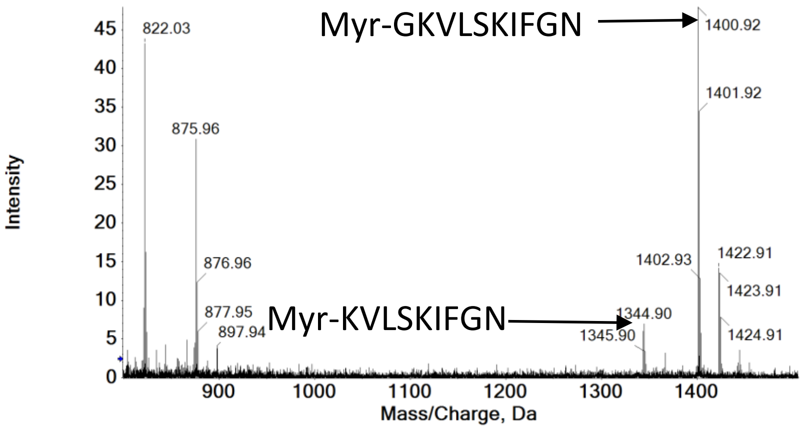

### T1 (1h NMT2 30°C)
### ARF6 short

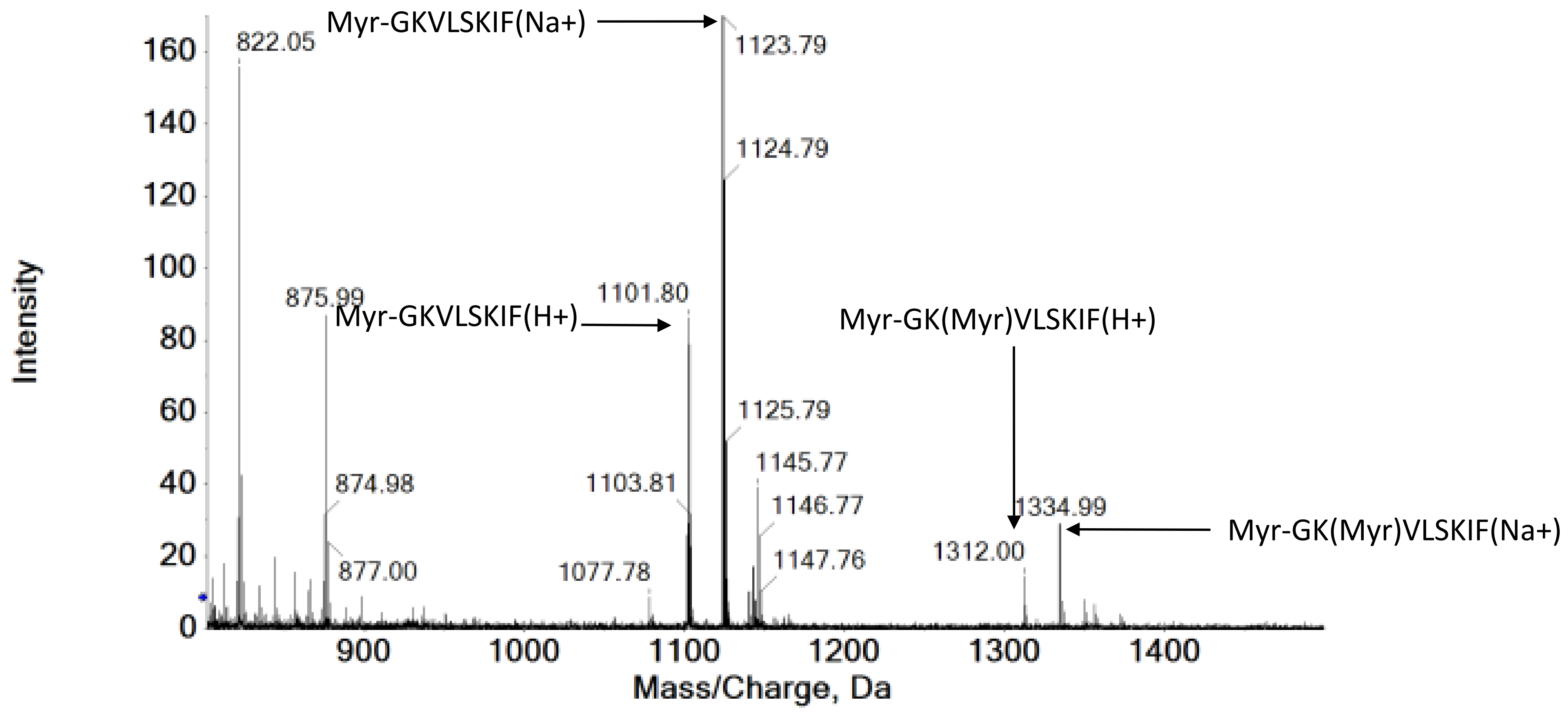

### Myr-GKVLSKIFGNK

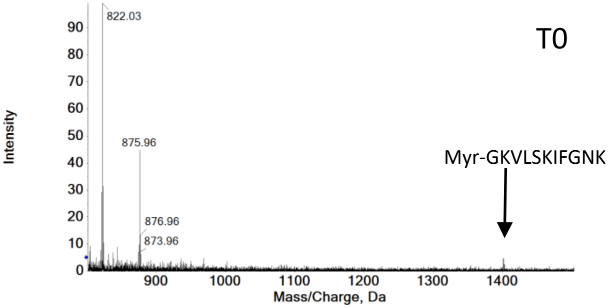


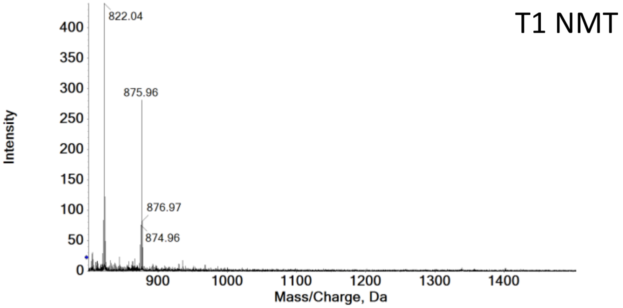


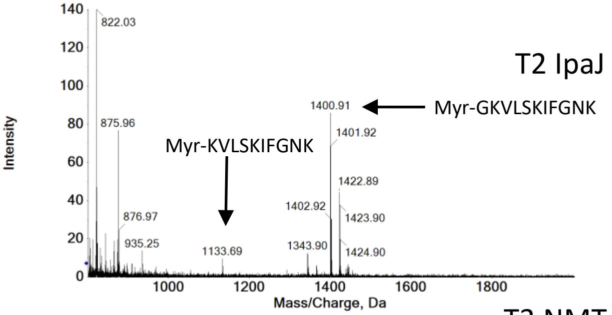


T3 NMT

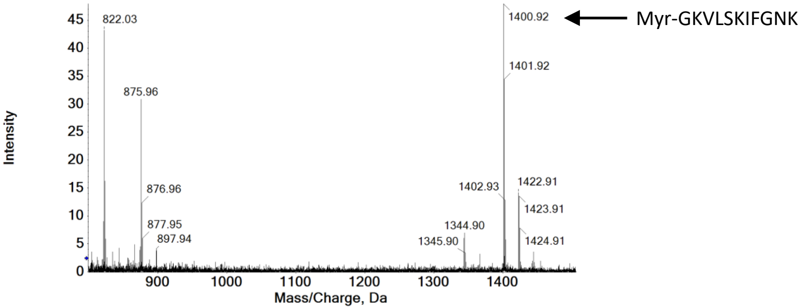


MSMS T2 Myr-GKVLSKIFGNK (1400)

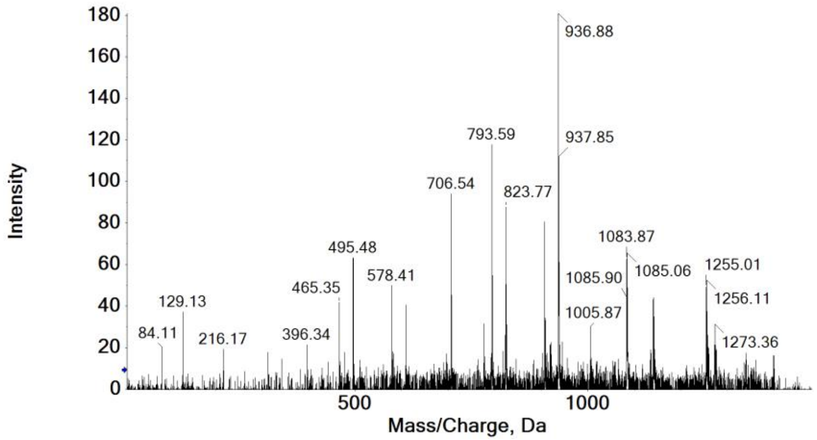

### G-K(ε-MYR)VLSKIFGNK

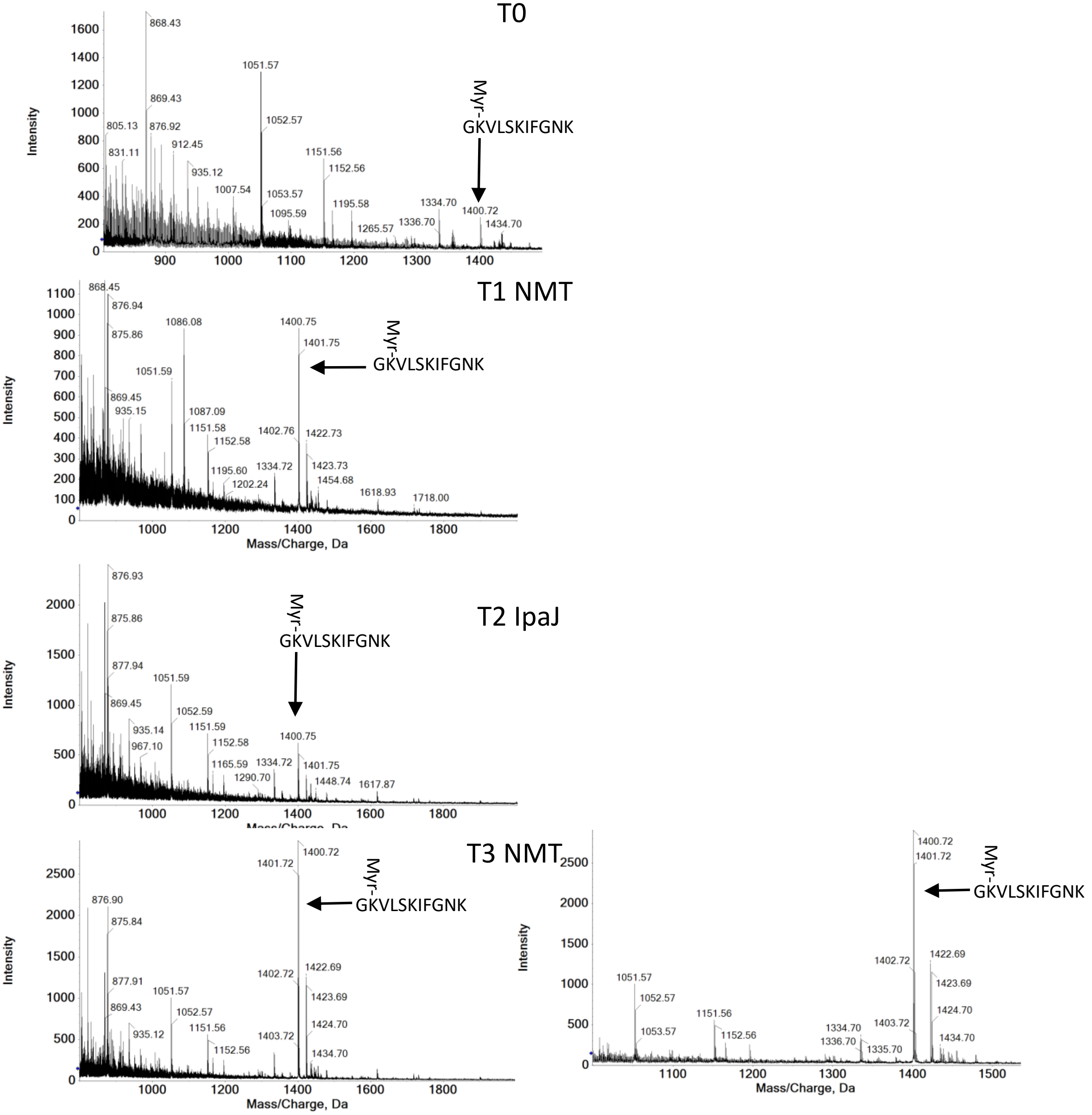


MSMS T1 Myr-GKVLSKIFGNK (1400)

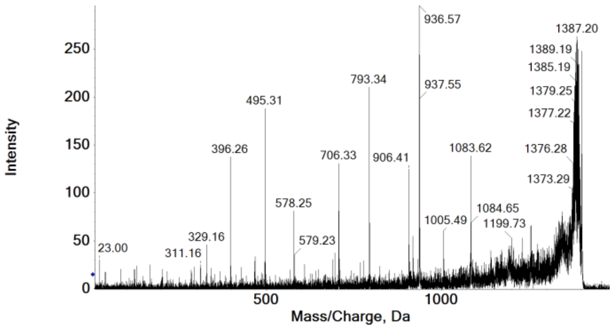


MSMS T3 Myr-GKVLSKIFGNK (1400)

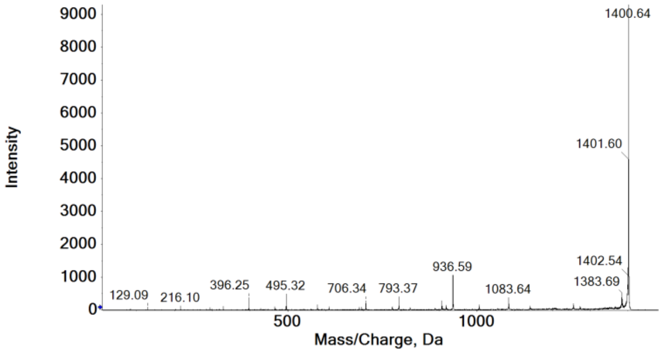

### GKFSSKPR

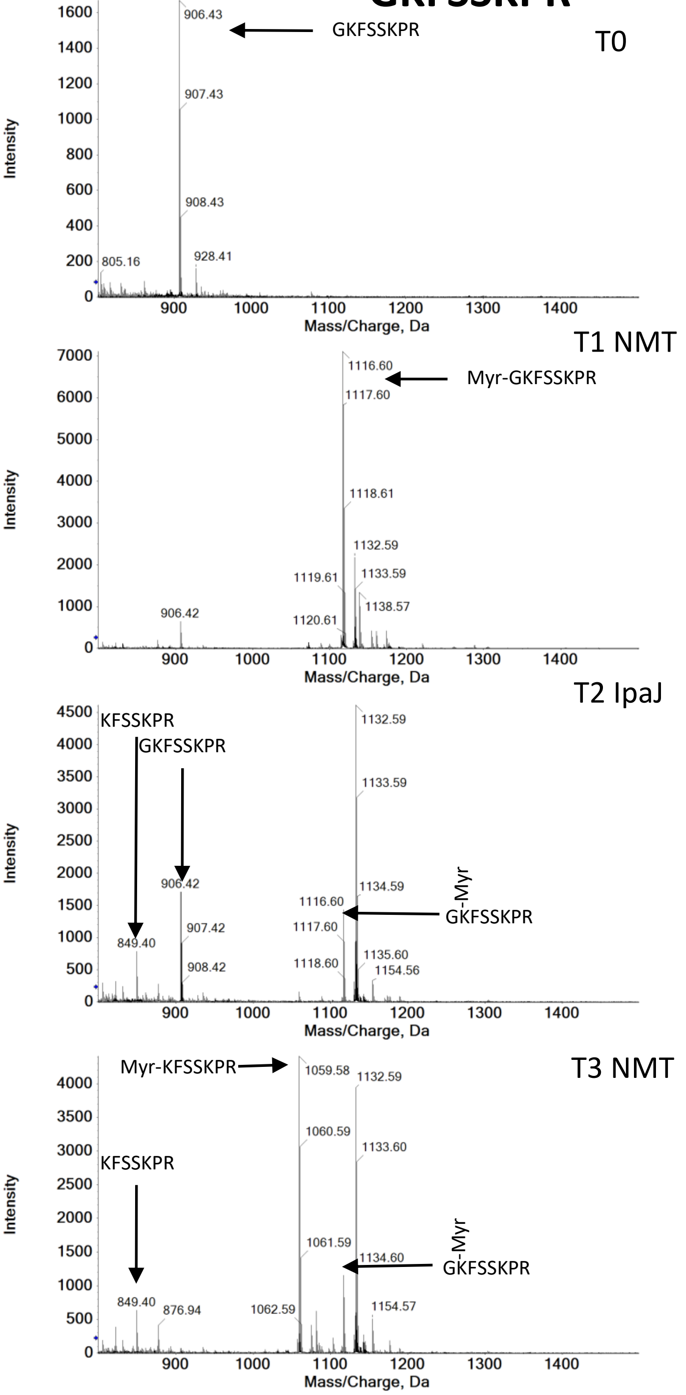
T0
GKFSSKPR
906.43
907.43
908.43
928.41
805.16
T1 NMT
Myr-GKFSSKPR
1116.60
1117.60
1118.61
1119.61
1120.61
1132.59
1133.59
1138.57
906.42
T2 IpaJ
KFSSKPR
GKFSSKPR
-Myr GKFSSKPR
849.40
906.42
907.42
908.42
1116.60
1117.60
1118.60
1132.59
1133.59
1134.59
1135.60
1154.56
T3 NMT
Myr-KFSSKPR
KFSSKPR
-Myr GKFSSKPR
1059.58
1060.59
1061.59
1062.59
1132.59
1133.60
1134.60
1154.57
849.40
876.94
Intensity
Mass/Charge, Da

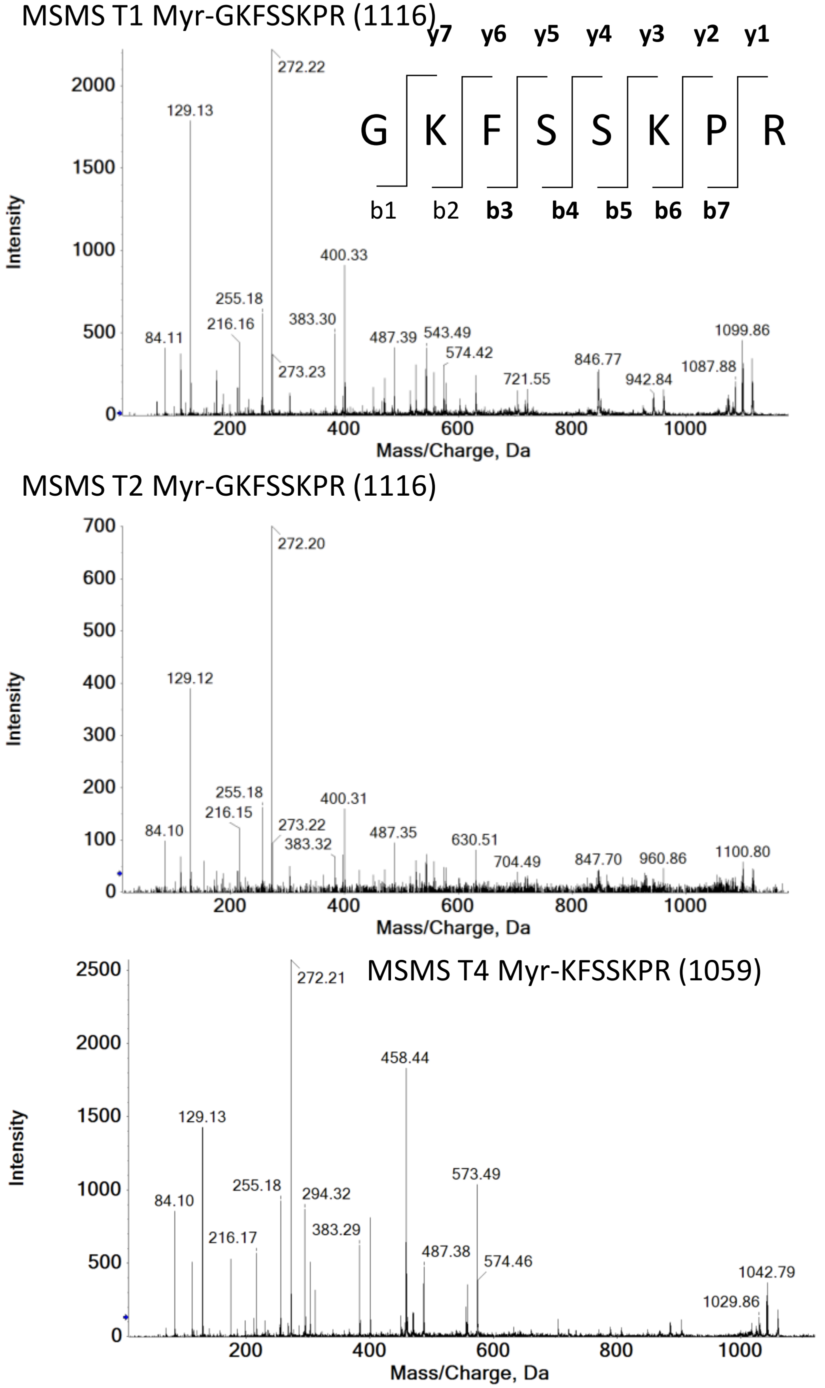
MSMS T1 Myr-GKFSSKPR (1116)
y7 y6 y5 y4 y3 y2 y1
G K F S S K P R
b1 b2 b3 b4 b5 b6 b7
272.22
129.13
400.33
255.18
216.16
383.30
84.11
487.39
543.49
574.42
273.23
721.55
846.77
942.84
1087.88
1099.86
MSMS T2 Myr-GKFSSKPR (1116)
272.20
129.12
255.18
216.15
400.31
273.22
383.32
84.10
487.35
630.51
704.49
847.70
960.86
1100.80
MSMS T4 Myr-KFSSKPR (1059)
272.21
458.44
129.13
255.18
294.32
84.10
216.17
383.29
573.49
487.38
574.46
1042.79
1029.86
Intensity
Mass/Charge, Da

### GKQNSKLR

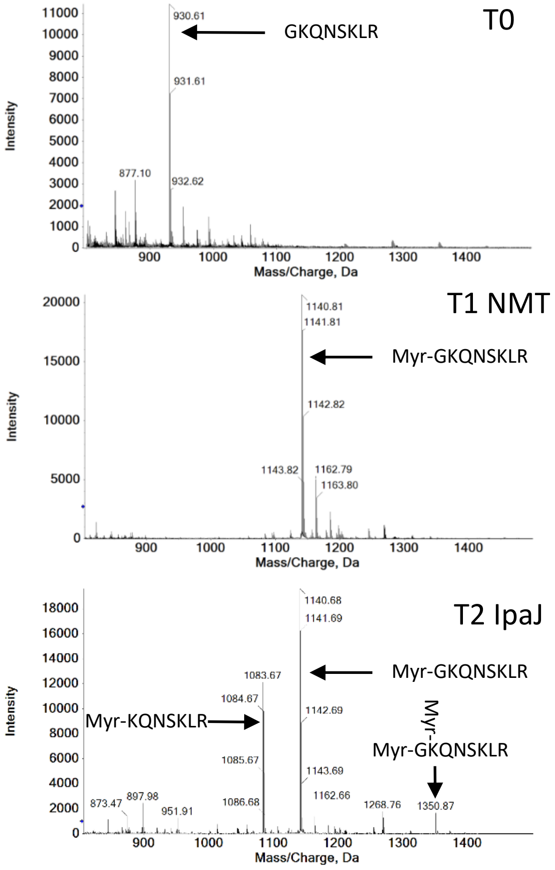


MSMS T0 GKQNSKLR (930)

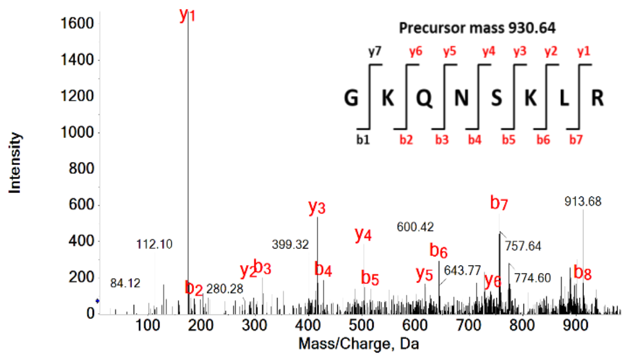


MSMS T1 Myr-GKQNSKLR (1140)

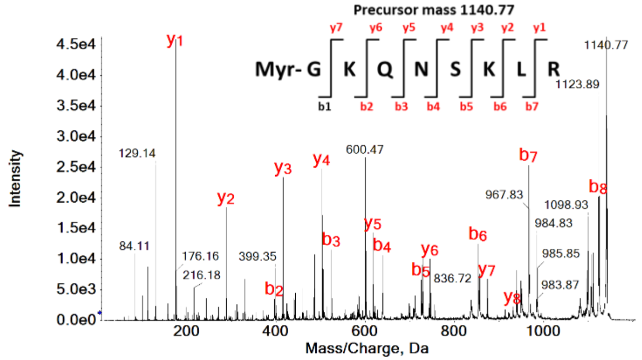


MSMS T2 Myr-KQNSKLR (1083)

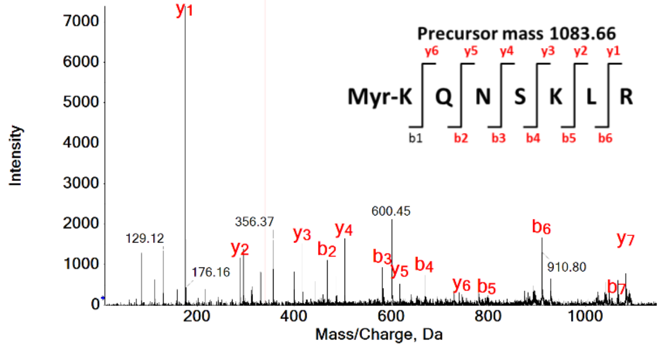


MSMS T2 Myr-GK(Myr)QNSKLR (1350)

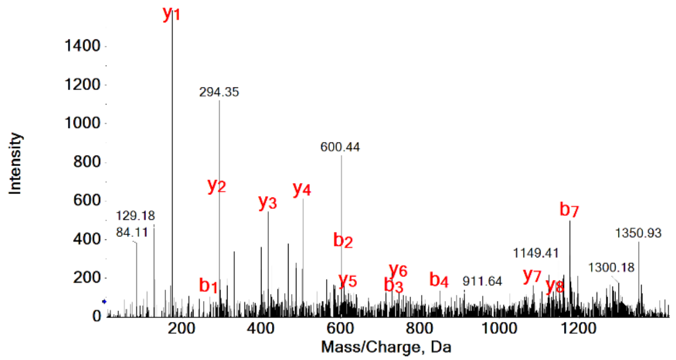


MSMS T2 Myr-GK(Myr)QNSKLR (1350)

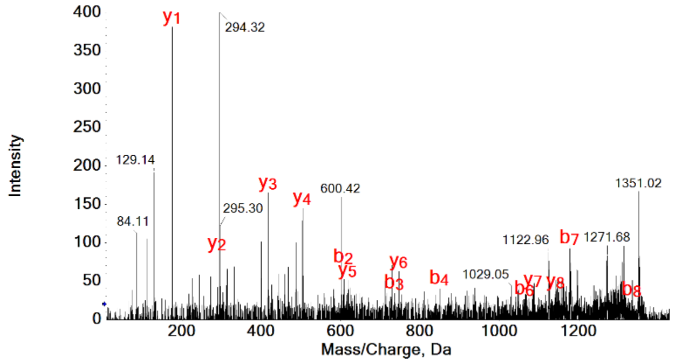


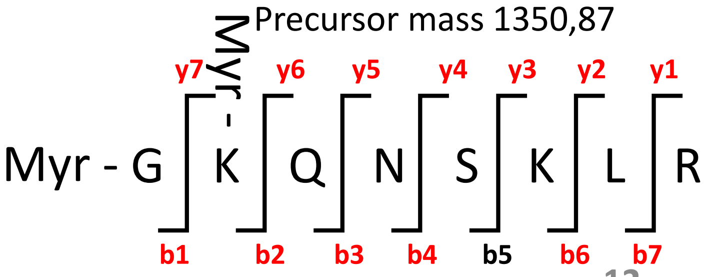

### KQNSKLRP

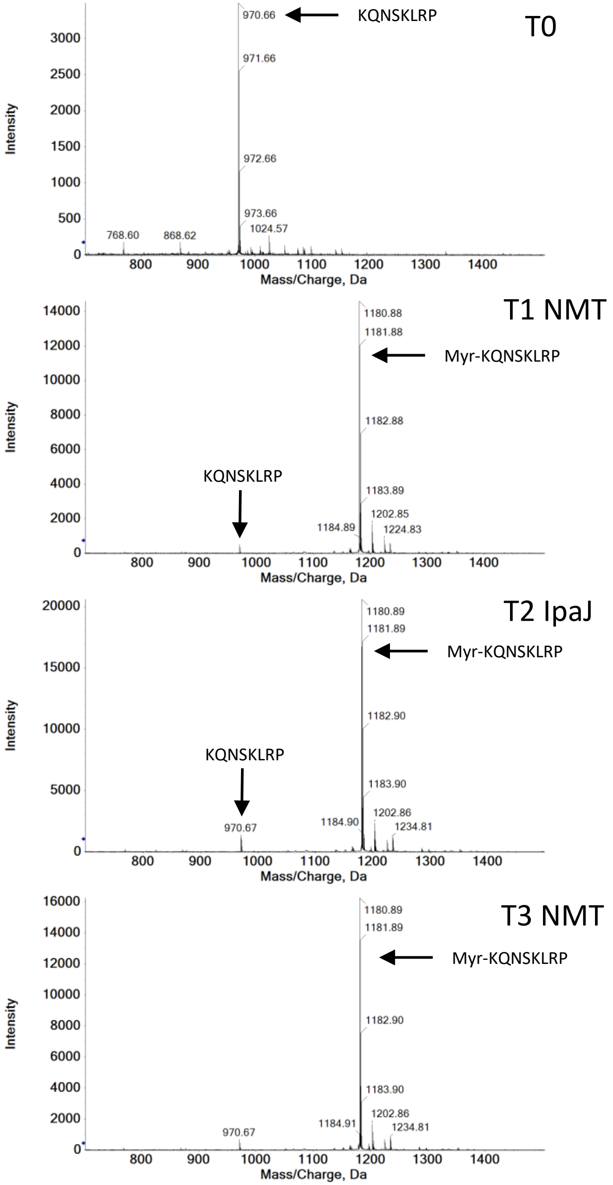


MSMS T2 Myr-KQNSKLR (1180)

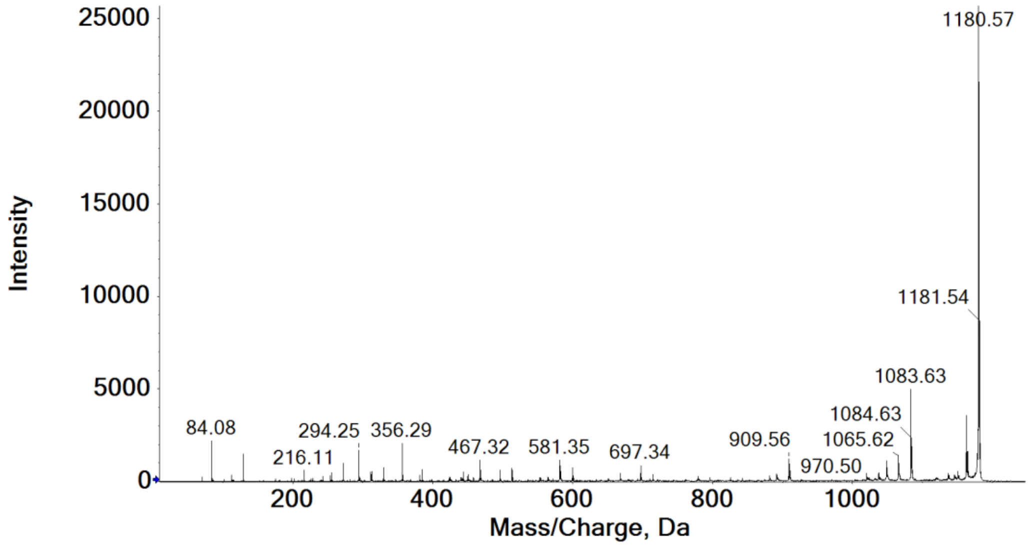

### GKQNSKLA

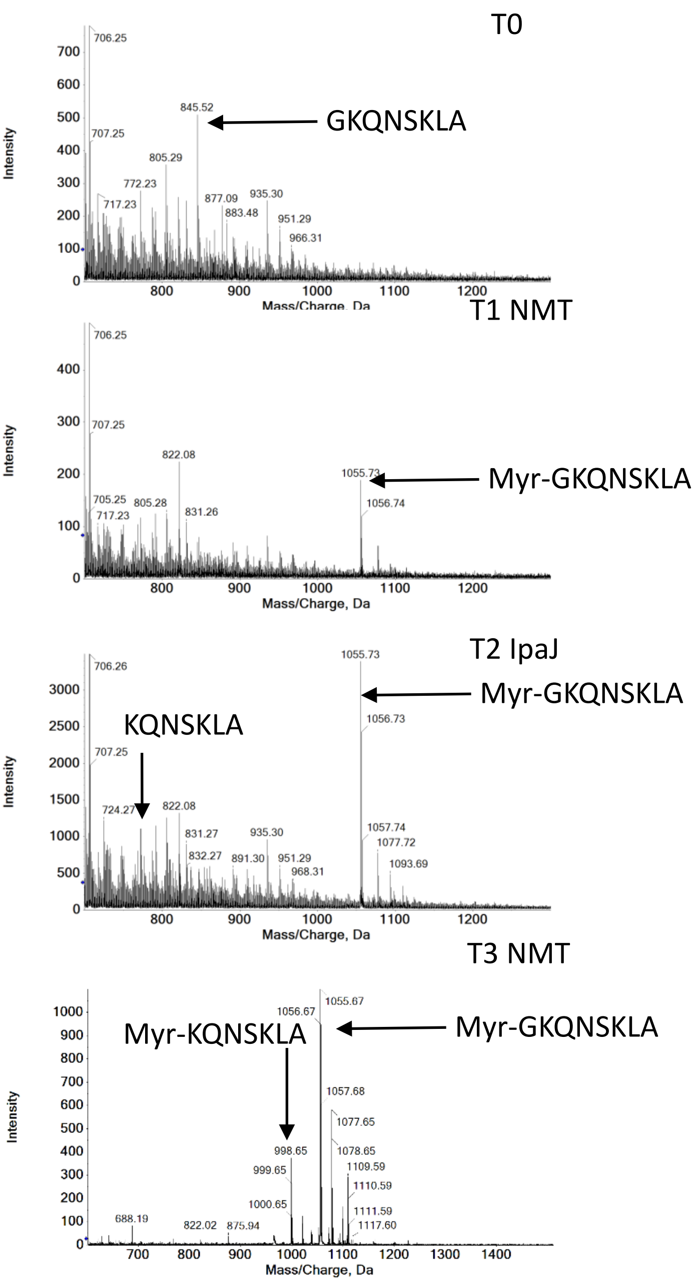


| F10_MSMS_998.6400_3 | | | ion |
|---|---|---|---|
| pic | Mass/Charge (Da) | Intensity | ion |
| 2155 | 331,2692 | 744,6275 | y3 |
| 3523 | 418,3104 | 564,8628 | y4 |
| 4794 | 467,4386 | 1193,098 | b2 |
| 6248 | 532,3957 | 2316,863 | y5 |
| 7446 | 581,477 | 1642,667 | b3 |
| 9310 | 660,4426 | 707,451 | y6 |
| 9508 | 668,5073 | 1420,863 | b4 |
| 12015 | 779,6317 | 800,9412 | b5 |
| 14810 | 909,7243 | 5384,157 | b6 |

MSMS T2 Myr-KQNSKLA (998)

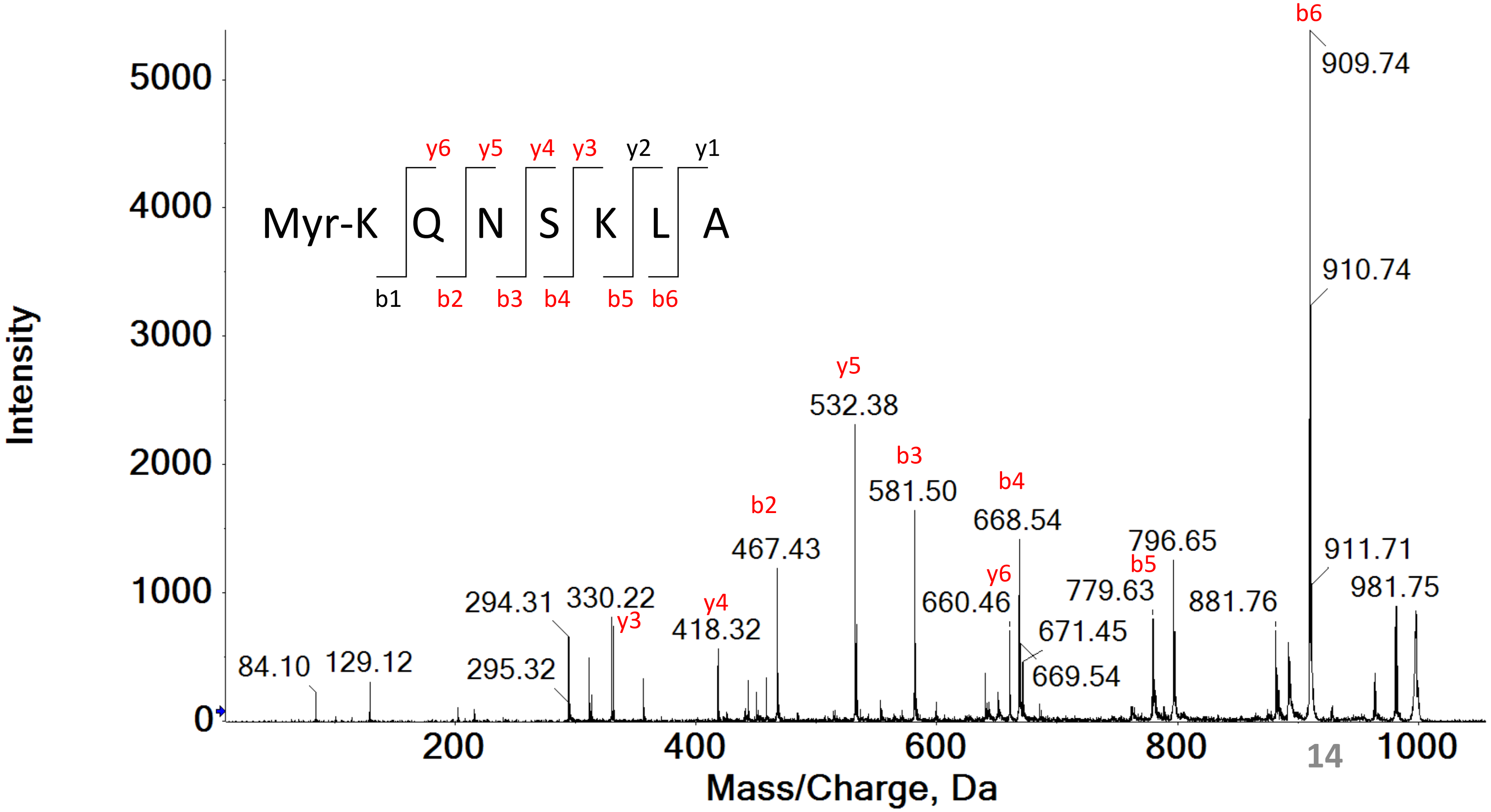

### GKSNSKLK

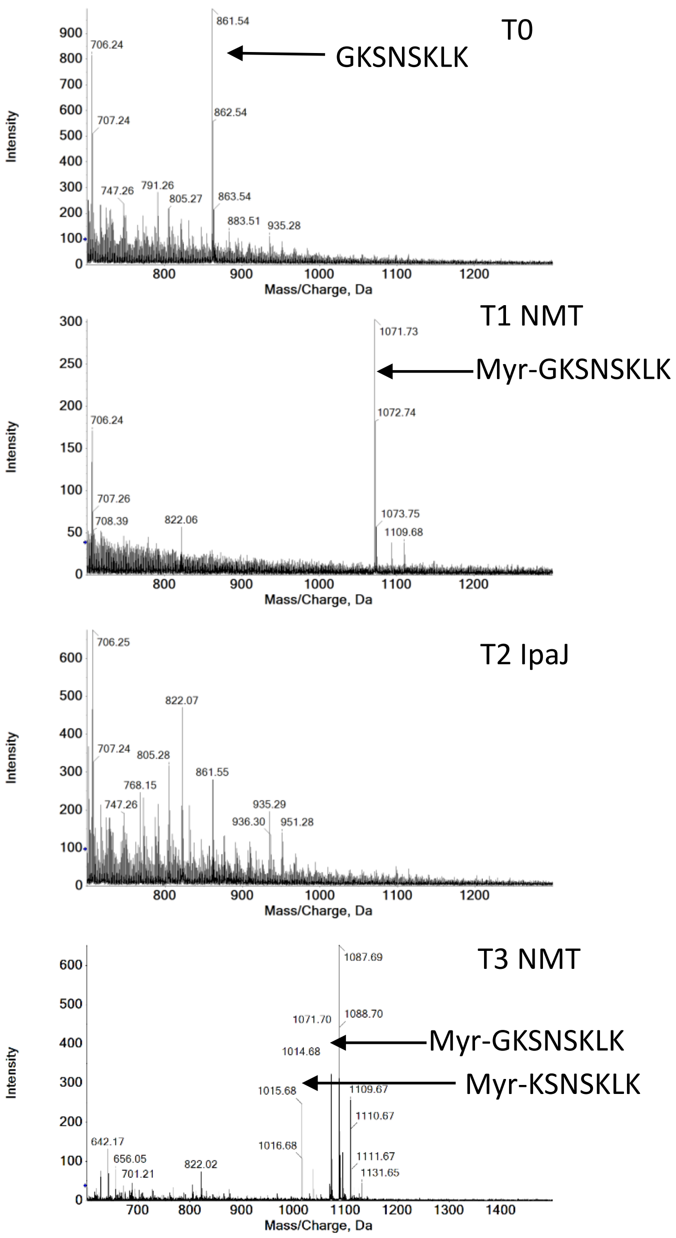


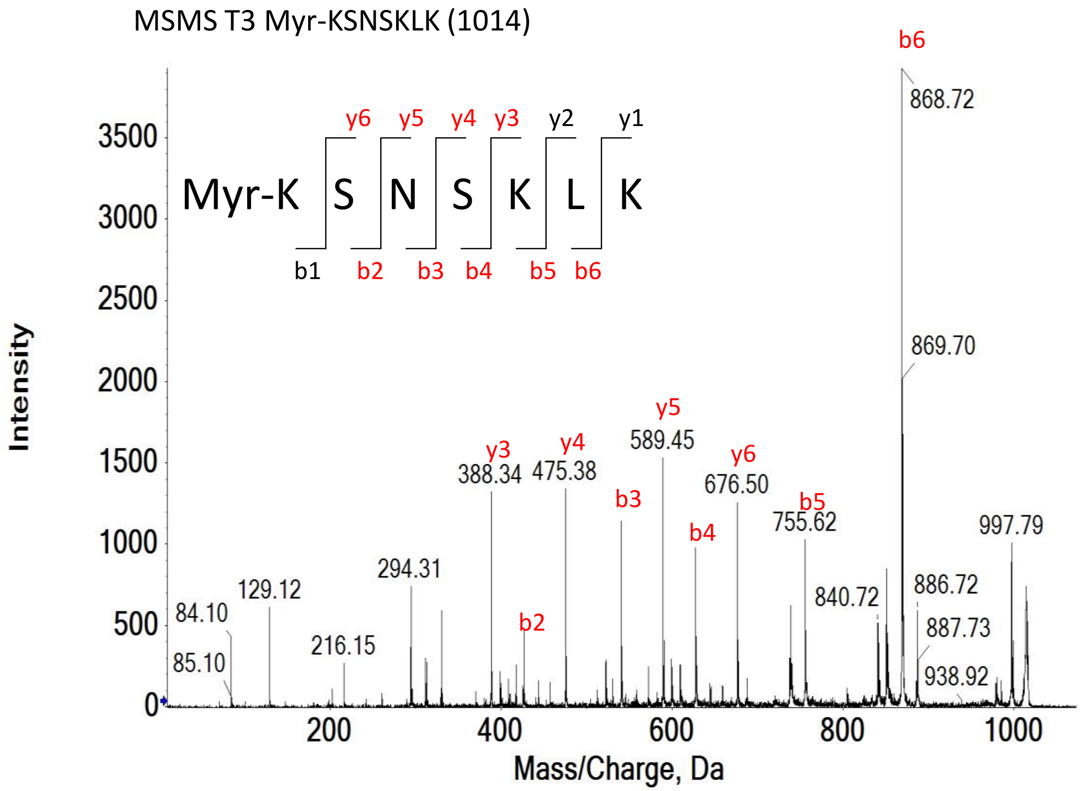


| F9_MSMS_1014.6000_2 | | | ion |
|---|---|---|---|
| pic | Mass/Charge (Da) | Intensity | ion |
| 4036 | 388,3268 | 1320,784 | y3 |
| 5032 | 426,3984 | 484,2353 | b2 |
| 6101 | 475,3693 | 1344,157 | y4 |
| 7622 | 540,4544 | 1142,118 | b3 |
| 8831 | 589,4373 | 1535,059 | y5 |
| 9779 | 627,5093 | 978,5098 | b4 |
| 10957 | 676,5035 | 1258,196 | y6 |
| 12771 | 755,6361 | 1029,804 | b5 |
| 15221 | 868,7002 | 3926,588 | b6 |

### GKTNSKLA

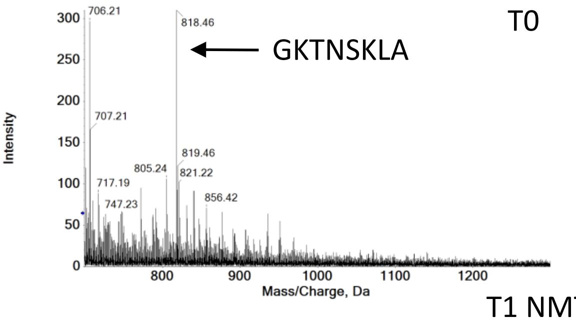


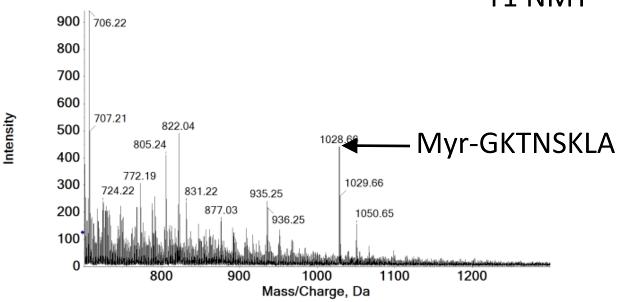


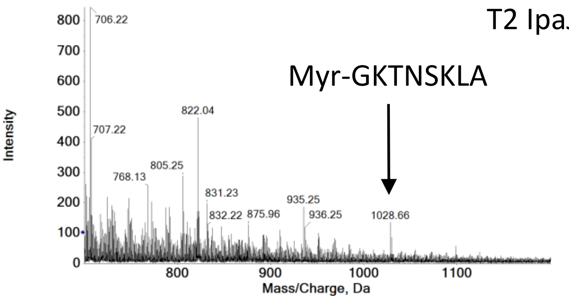


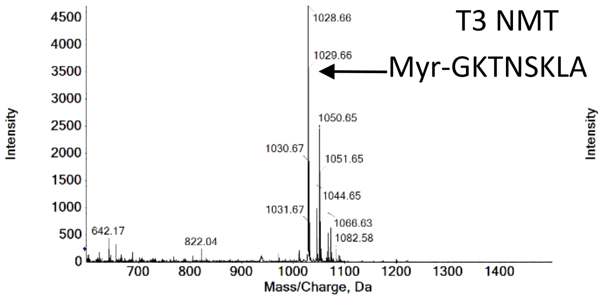


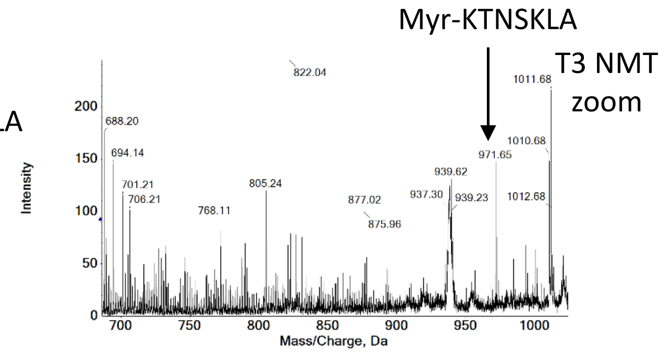


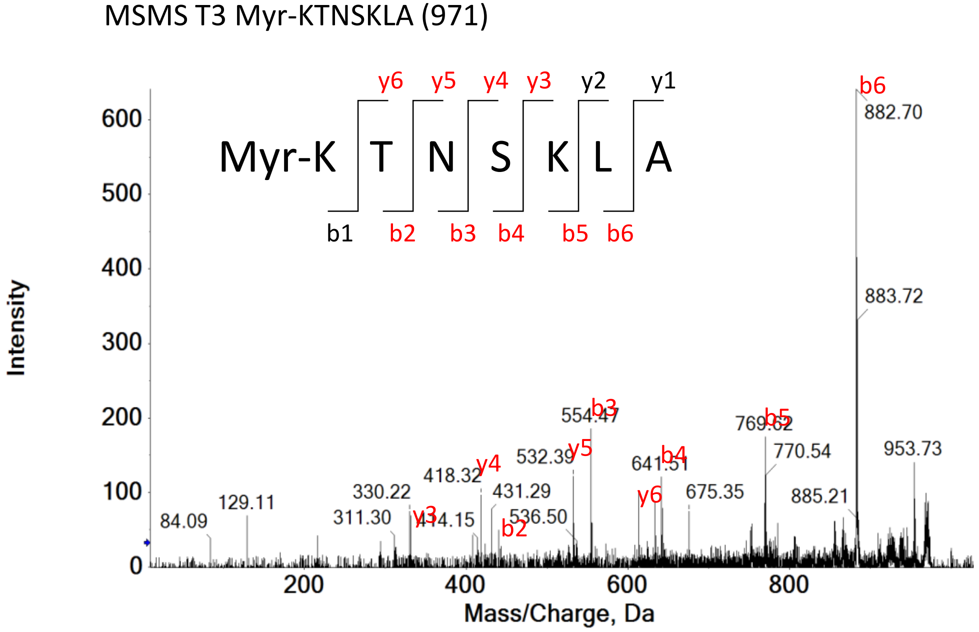


| F7_MSMS_971.6000_2 | | | ion |
|---|---|---|---|
| pic | Mass/Charge (Da) | Intensity | ion |
| 1835 | 331,2539 | 69,4902 | y3 |
| 2576 | 418,3141 | 97,09805 | y4 |
| 2865 | 440,4206 | 49,56863 | b2 |
| 3905 | 532,3945 | 121,8824 | y5 |
| 4239 | 554,4536 | 185,5686 | b3 |
| 5292 | 633,4332 | 87,84314 | y6 |
| 5428 | 641,4906 | 121,4118 | b4 |
| 7520 | 769,2246 | 62,27451 | b5 |
| 9452 | 882,458 | 73,56863 | b6 |

### GKSASKQF

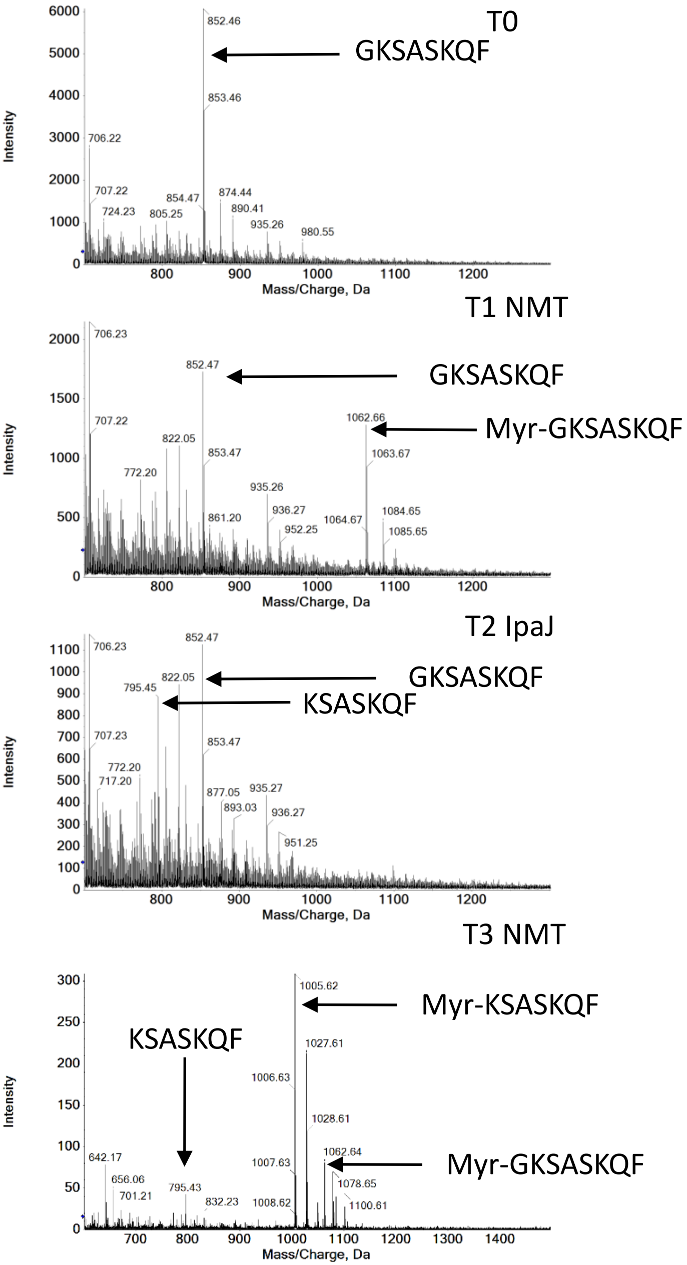


MSMS T3 Myr-KSASKQF (1005)

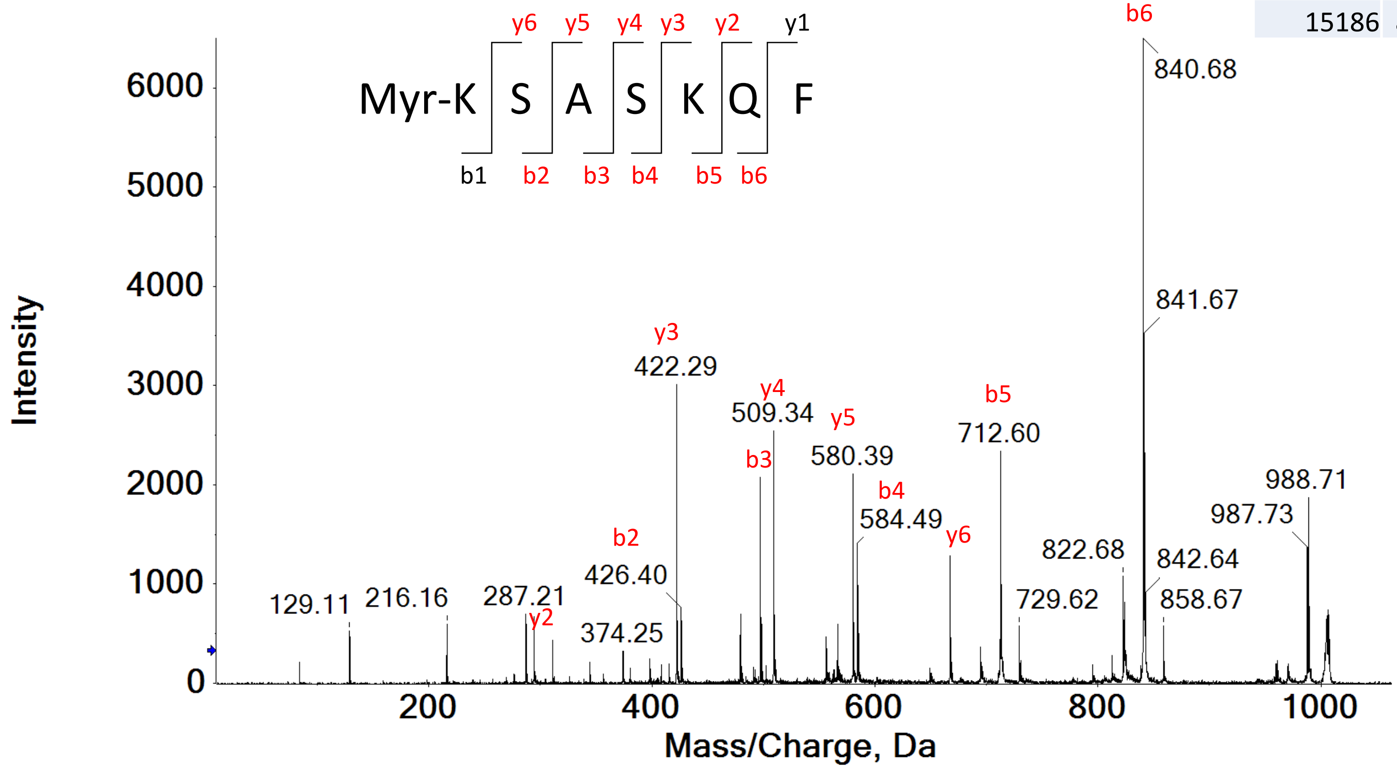


| F8_MSMS_1005.6000_3 | | | ion |
|---|---|---|---|
| pic | Mass/Charge (Da) | Intensity | ion |
| 2238 | 294,1691 | 387,451 | y2 |
| 5108 | 422,2837 | 3010,667 | y3 |
| 5235 | 426,3989 | 758,902 | b2 |
| 6967 | 497,4295 | 2081,255 | b3 |
| 7300 | 509,3415 | 2547,294 | y4 |
| 9172 | 580,4089 | 2050,039 | y5 |
| 9279 | 584,4832 | 1414,902 | b4 |
| 11315 | 667,4761 | 958,5883 | y6 |
| 12367 | 712,6038 | 2342,745 | b5 |
| 15186 | 840,6764 | 6497,726 | b6 |

### GAKQSGPA

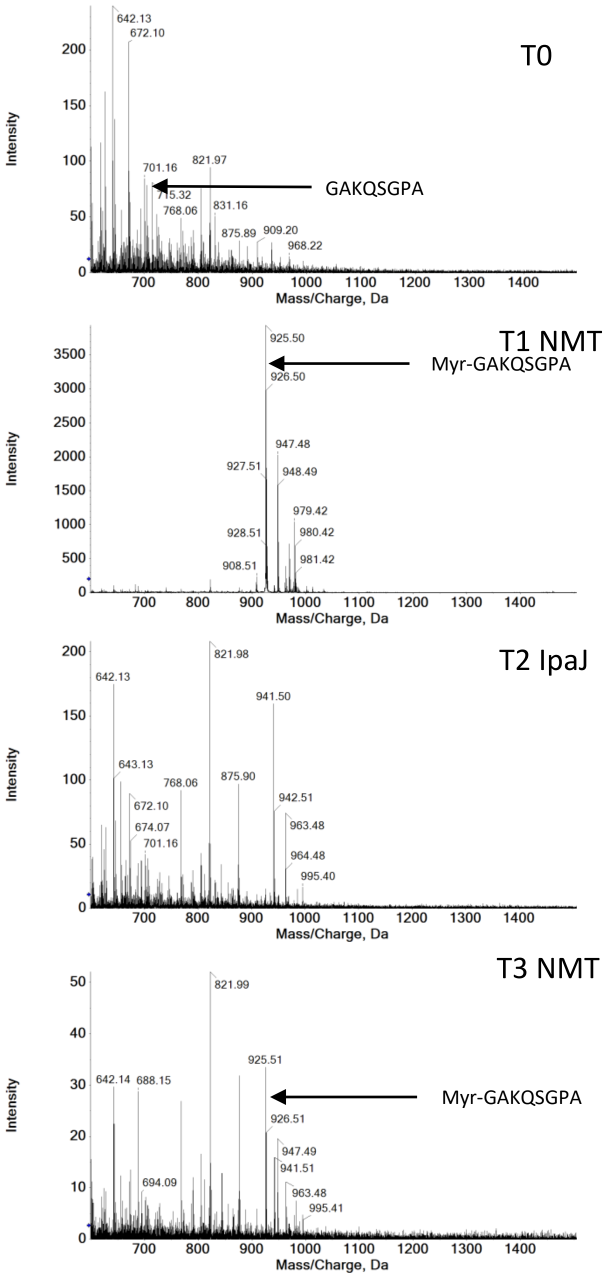
T0
GAKQSGPA
642.13
672.10
701.16
821.97
715.32
768.06
831.16
875.89
909.20
968.22
T1 NMT
Myr-GAKQSGPA
925.50
926.50
947.48
927.51
948.49
979.42
928.51
980.42
981.42
908.51
T2 IpaJ
821.98
642.13
941.50
643.13
768.06
875.90
672.10
942.51
963.48
674.07
701.16
964.48
995.40
T3 NMT
Myr-GAKQSGPA
821.99
925.51
642.14
688.15
926.51
947.49
941.51
694.09
963.48
995.41
Intensity
Mass/Charge, Da

### GGKQSTAA

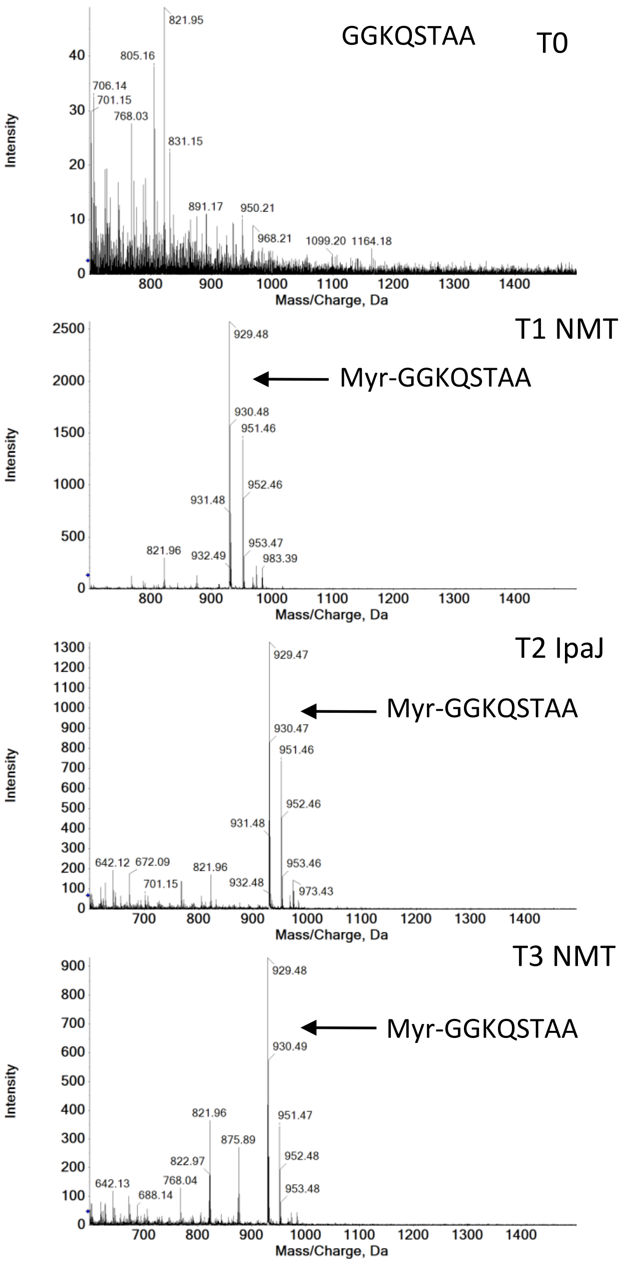
GGKQSTAA
T0
821.95
805.16
706.14
701.15
768.03
831.15
891.17
950.21
968.21
1099.20
1164.18
Intensity
Mass/Charge, Da
T1 NMT
929.48
Myr-GGKQSTAA
930.48
951.46
952.46
931.48
821.96
932.49
953.47
983.39
T2 IpaJ
929.47
Myr-GGKQSTAA
930.47
951.46
952.46
931.48
642.12
672.09
701.15
821.96
932.48
953.46
973.43
T3 NMT
929.48
Myr-GGKQSTAA
930.49
821.96
951.47
875.89
822.97
952.48
768.04
642.13
688.14
953.48

### GGKLSKKK

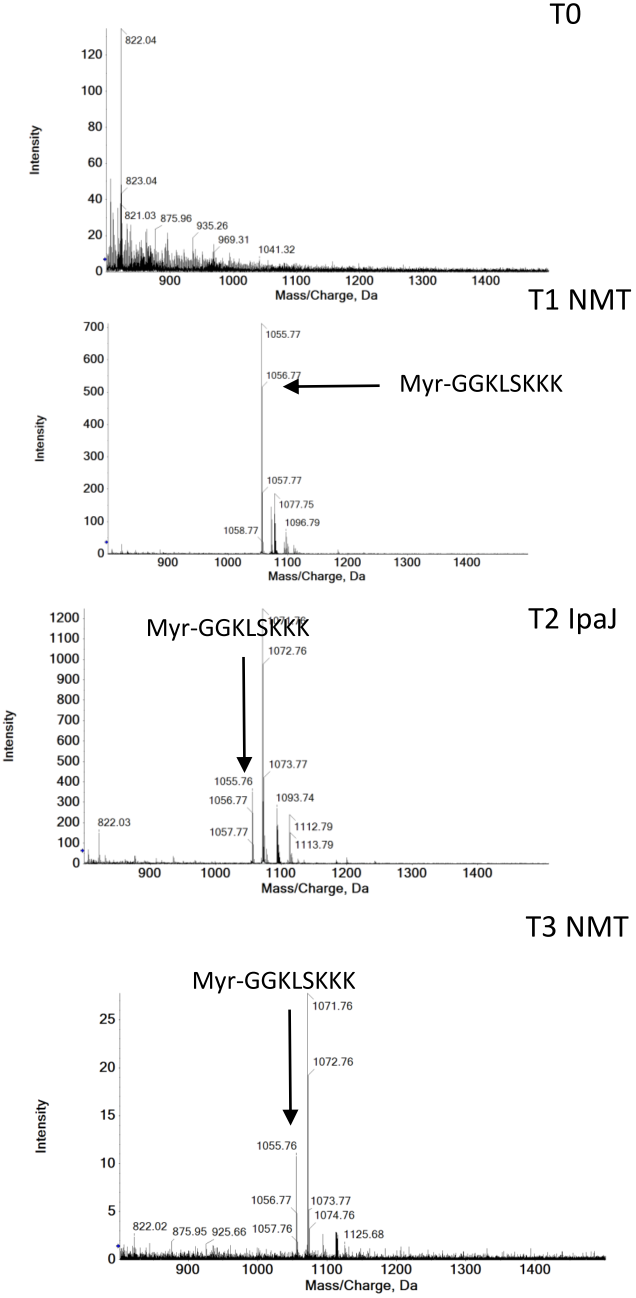


| | | | |
|---|---|---|---|
| y2 | 2642 | 275,2397 | 400,9412 |
| y3 | 5363 | 403,3621 | 1237,961 |
| b3 | 6526 | 453,4303 | 375,8431 |
| y4 | 7591 | 490,8696 | 179,6078 |
| b4 | 9598 | 566,5328 | 1952,314 |
| y5 | 10561 | 603,5288 | 1129,098 |
| b5 | 11824 | 653,5715 | 1144,941 |
| y6 | 13686 | 731,6654 | 131,2941 |
| b6 | 14812 | 781,6852 | 615,8431 |
| y7 | 14969 | 788,6453 | 359,6863 |
| b7 | 17555 | 909,8929 | 1265,882 |

MSMS T1 Myr-GGKLSKKK (1055)

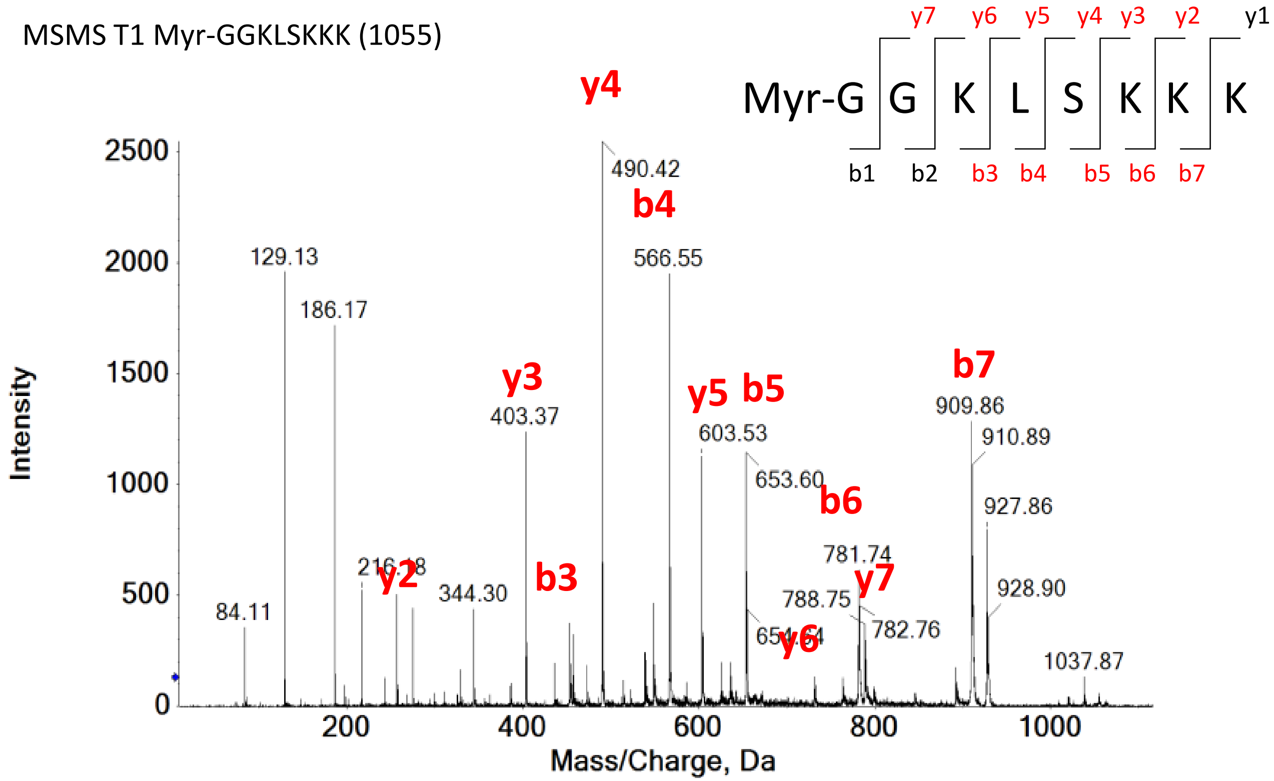

### GGKWSKLS

T0

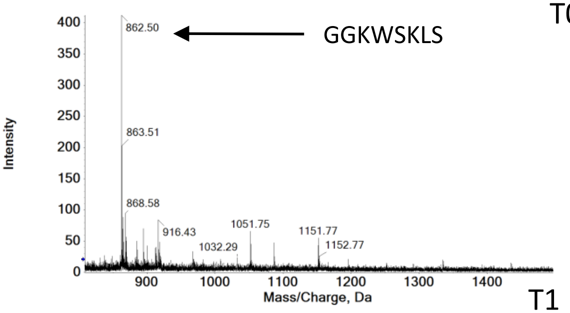


T1 NMT

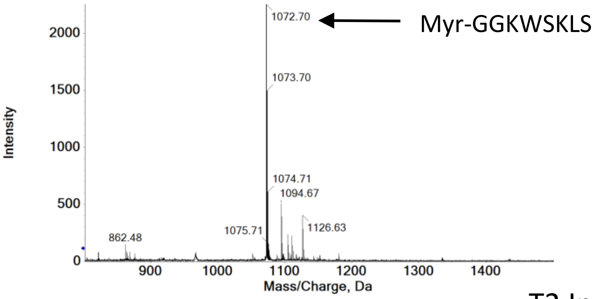


T2 IpaJ

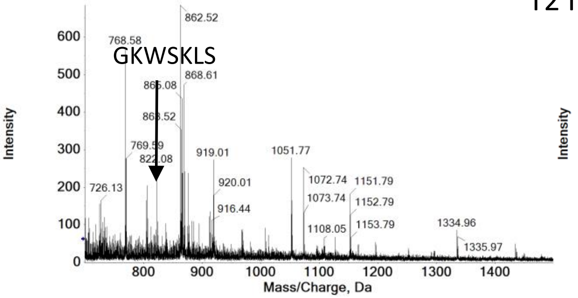


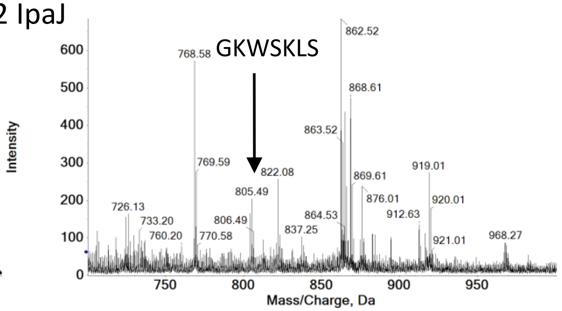


T3 NMT

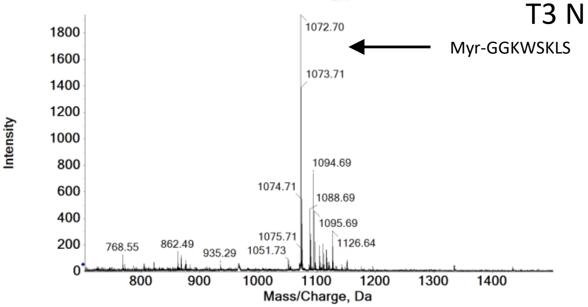


MSMS T1 Myr-GGKWSKLS (1072) (1)

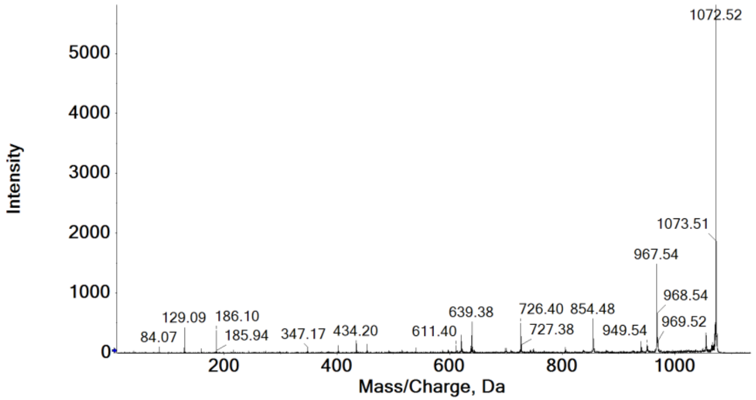


MSMS T1 Myr-GGKWSKLS (1072) (2)

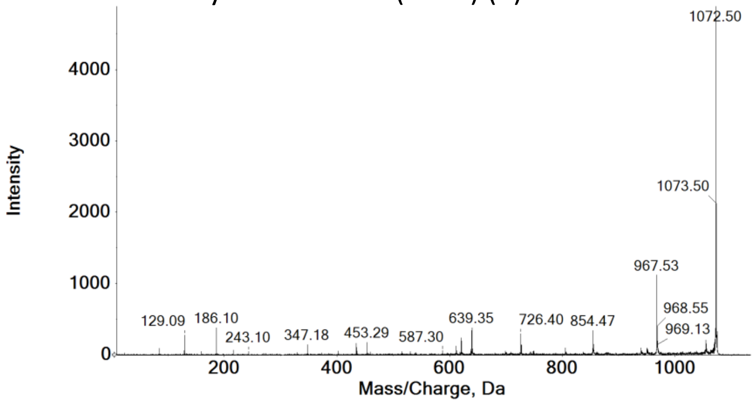

### AKSFSKPR

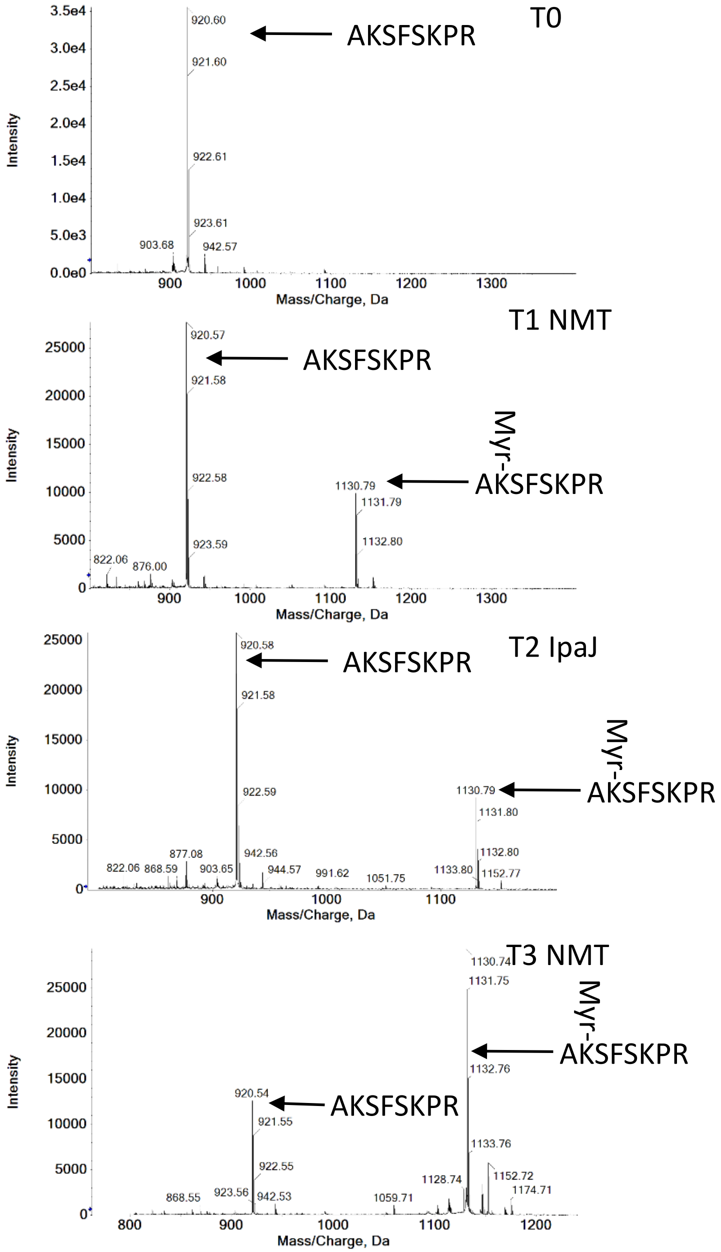


MSMS T1 AK(Myr)SFSKPR (1130)

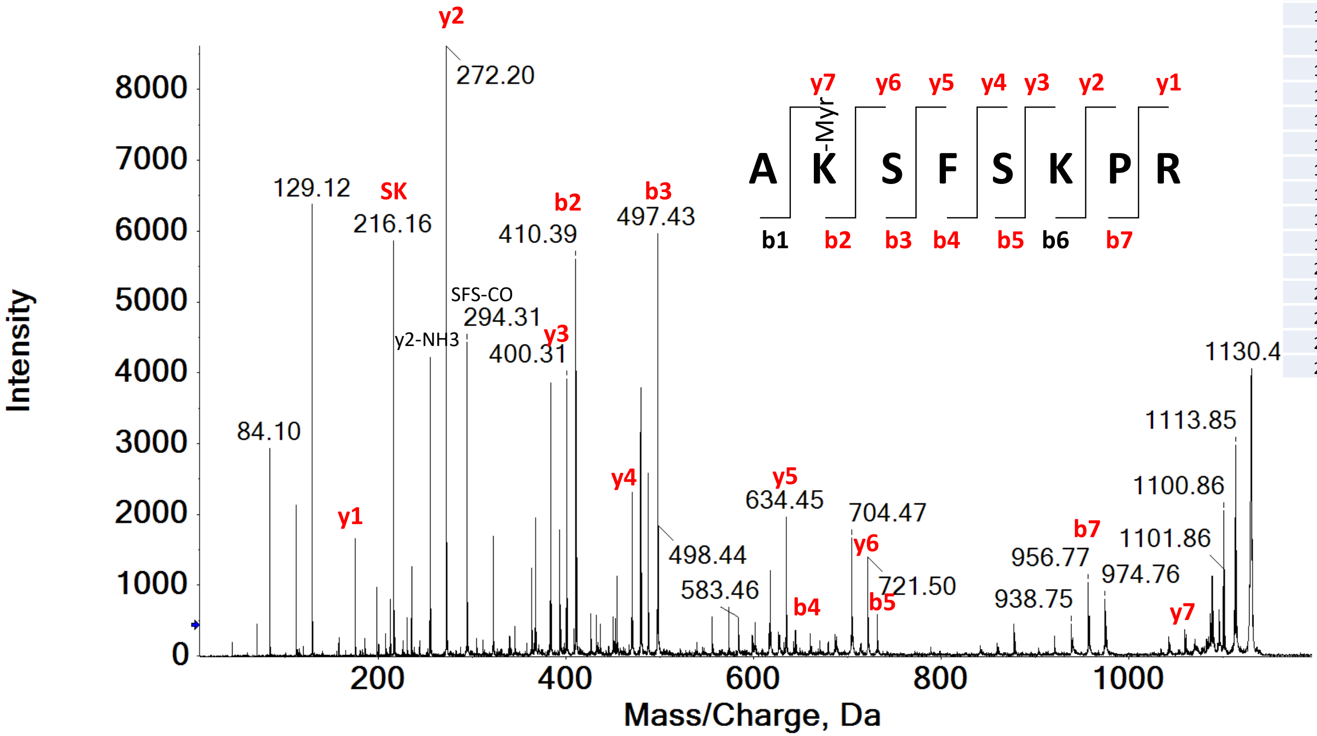


| | | | |
|---|---|---|---|
| 766 | 84,09495 | 2935,84 | K |
| 1481 | 112,1094 | 2126,90 | R |
| 2056 | 129,1225 | 6378,35 | K |
| 3482 | 175,1427 | 1653,96 | y1 |
| 4180 | 198,1471 | 973,80 | SK-H2O/ KP-CO |
| 4900 | 216,147 | 5869,18 | SK |
| 5668 | 235,1309 | 1267,45 | FS |
| 6392 | 255,174 | 4213,96 | y2-NH3 |
| 7039 | 272,2352 | 6608,47 | y2 |
| 7848 | 294,3016 | 4431,22 | SFS-CO |
| 8846 | 322,1745 | 1693,18 | SFS |
| 9640 | 345,2368 | 414,43 | FSK-H2O |
| 10247 | 363,2515 | 1236,71 | FSK |
| 10905 | 383,2669 | 3857,57 | y3-NH3 |
| 11231 | 393,3545 | 1791,69 | b2-NH3 |
| 11453 | 400,3001 | 3915,61 | y3 |
| 11709 | 408,3862 | 384,78 | K(+210)S-H2O |
| 11772 | 410,3888 | 5604,39 | b2 |
| 12270 | 426,3936 | 597,18 | K(+210)S |
| 12996 | 450,2837 | 557,65 | SFSK |
| 13590 | 470,3228 | 2309,80 | y4-NH3 |
| 13855 | 479,4061 | 3797,80 | $b_3$-$H_2O$ |
| 13884 | 480,4055 | 1171,45 | b3-NH3 |
| 14085 | 487,3613 | 2589,18 | y4 |
| 14373 | 497,4166 | 5962,98 | b3 |
| 16462 | 573,4848 | 685,96 | K(+210)SF |
| 17604 | 617,4413 | 1208,78 | y5-NH3 |
| 18035 | 634,4463 | 1964,86 | y5 |
| 18287 | 644,498 | 369,88 | b4 |
| 19750 | 704,4481 | 1667,14 | y6-NH3 |
| 20154 | 721,4829 | 1395,76 | y6 |
| 20390 | 731,5301 | 590,59 | b5 |
| 25295 | 956,7646 | 1045,80 | b7 |
| 25660 | 974,7401 | 803,45 | b7+H2O |
| 27339 | 1059,846 | 375,53 | y7 |

### GKSWSKGR

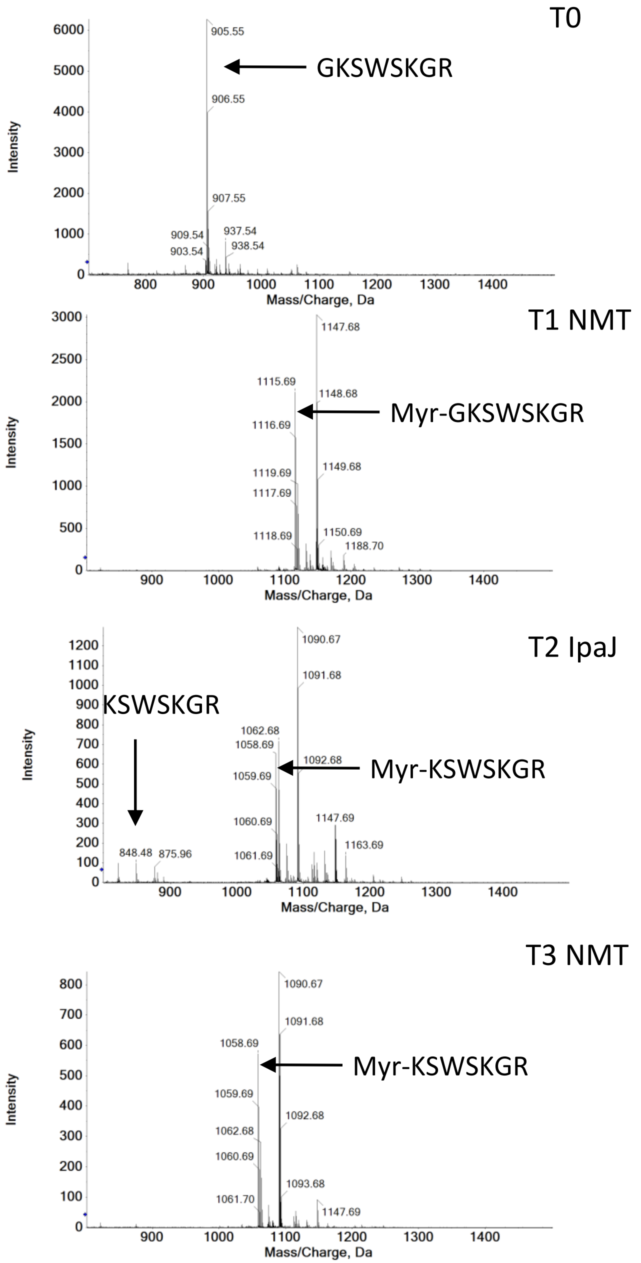


MSMS T2 Myr-KSWSKGR (1158)

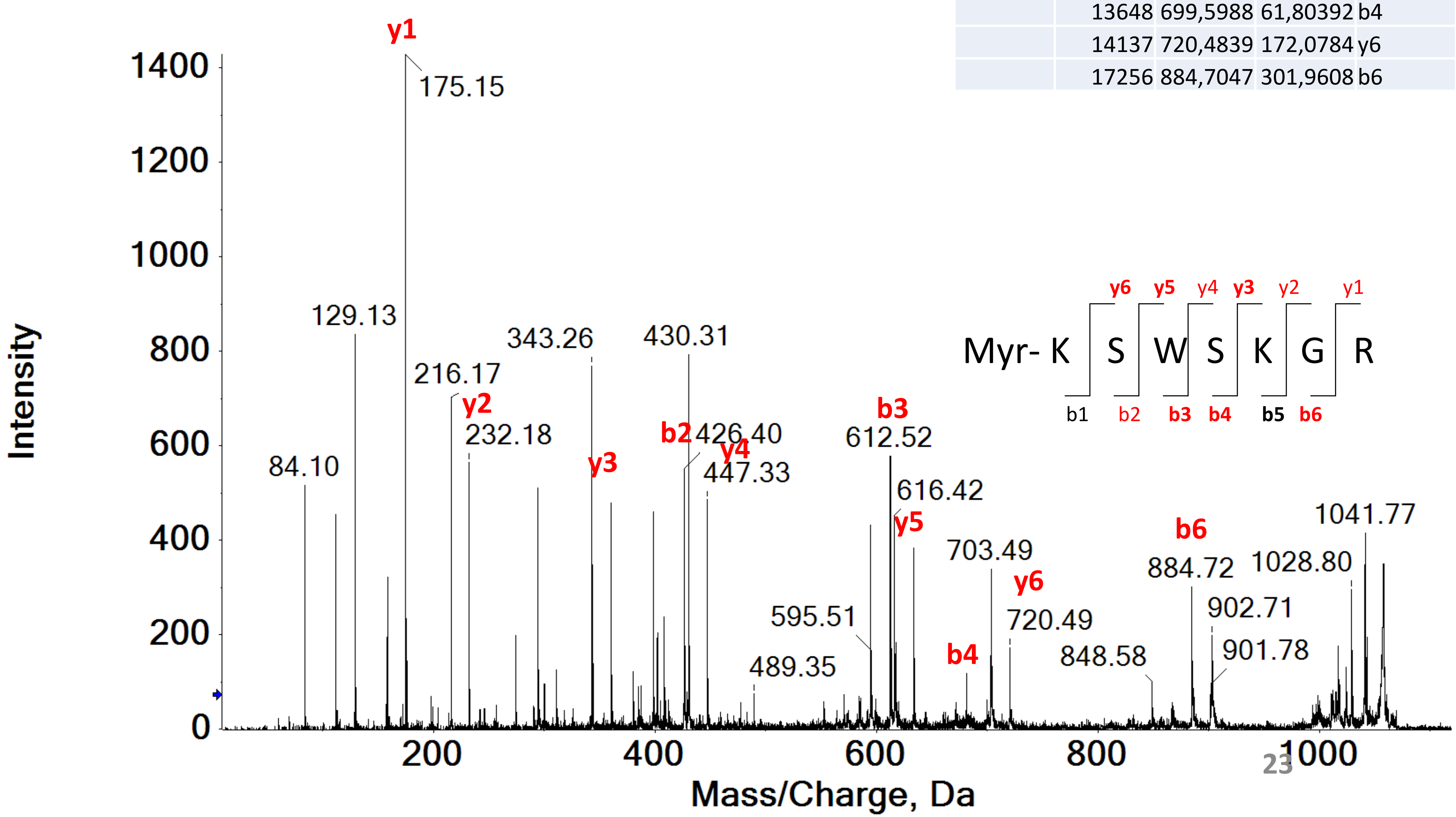


| F20 1058.68 | | Mass/Charge (Da) | Intensity | |
|---|---|---|---|---|
| | 1520 | 175,1429 | 1428,392 | y1 |
| | 2460 | 232,1888 | 496,1569 | y2 |
| | 5185 | 360,3024 | 479,3726 | y3 |
| | 7074 | 426,405 | 552,1569 | b2 |
| | 7655 | 447,3432 | 485,4902 | y4 |
| | 11575 | 612,537 | 578,0392 | b3 |
| | 12102 | 633,4509 | 383,6863 | y5 |
| | 13648 | 699,5988 | 61,80392 | b4 |
| | 14137 | 720,4839 | 172,0784 | y6 |
| | 17256 | 884,7047 | 301,9608 | b6 |

### AKQNSKLR

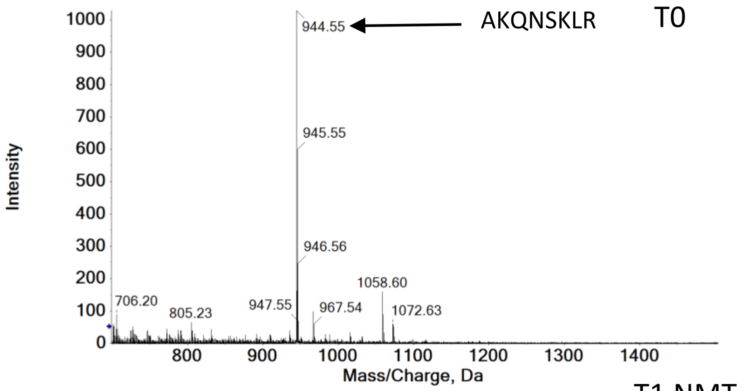


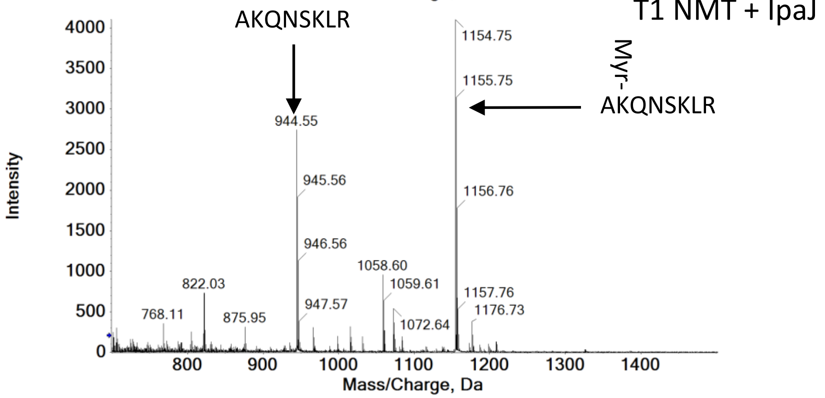


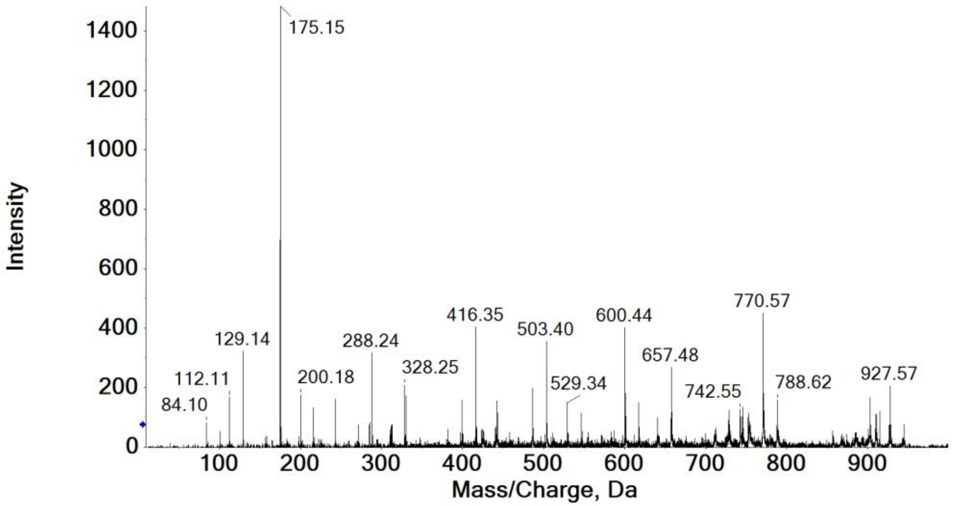


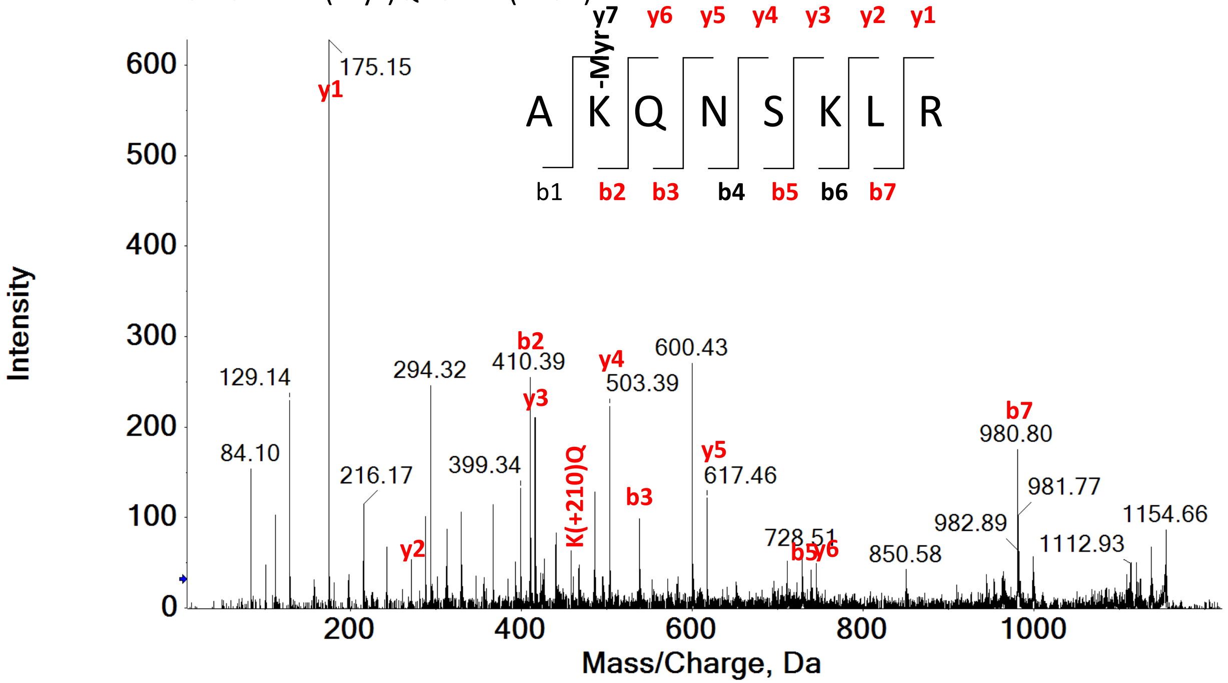


| K24 1154 | Mass/Charge (Da) | Intensity | |
|---|---|---|---|
| 535 | 175,0203 | 35,45098 | y1 |
| 1231 | 288,1639 | 31,52941 | y2 |
| 2914 | 410,3834 | 254,7451 | b2 |
| 3049 | 416,3236 | 210,8235 | y3 |
| 3905 | 467,378 | 43,60785 | K(+210)Q ? |
| 4432 | 503,3777 | 223,3726 | y4 |
| 4875 | 538,4333 | 98,98039 | b3 |
| 5859 | 600,4333 | 249,8824 | y5-NH3 |
| 6146 | 617,4308 | 121,4118 | y5 |
| 8017 | 739,5452 | 42,5098 | b5 |
| 8118 | 745,4847 | 49,09804 | y6 |
| 9226 | 850,5908 | 42,5098 | b6-NH3 |
| 10884 | 980,7767 | 175,5294 | b7 |
| 12877 | 1137,767 | 67,45098 | v8 |

### GGKQSKAA

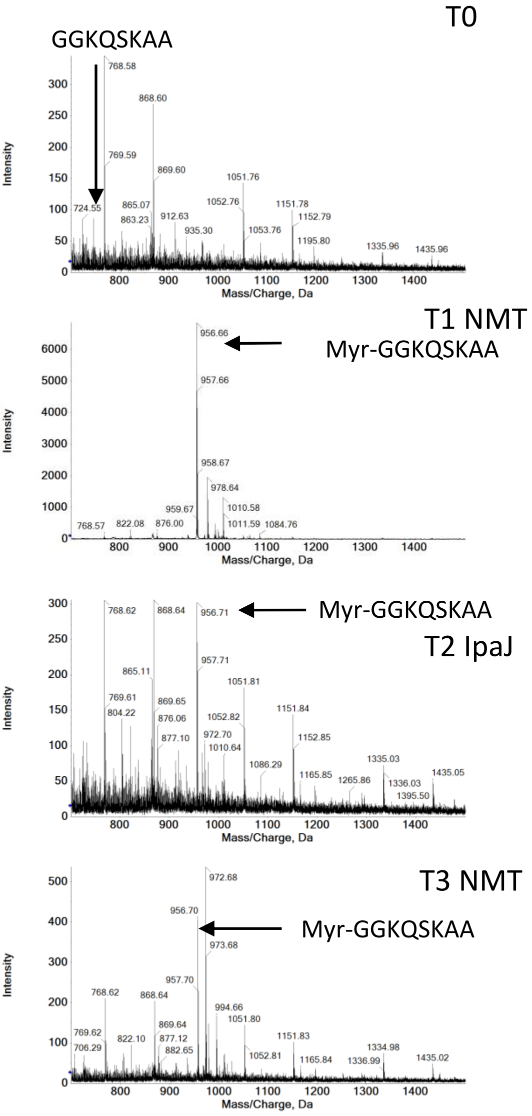


MSMS T1 Myr-GGKQSKAA (956)

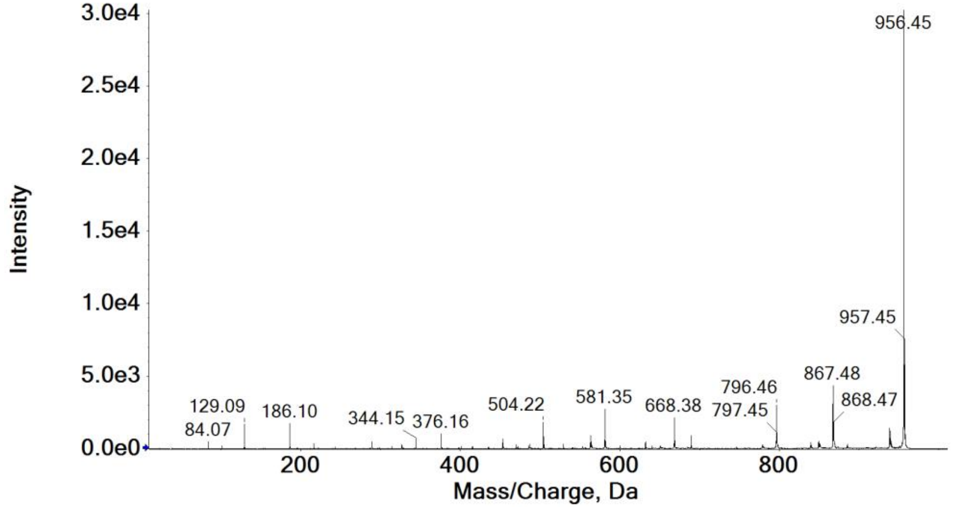

### GGKSFSKP

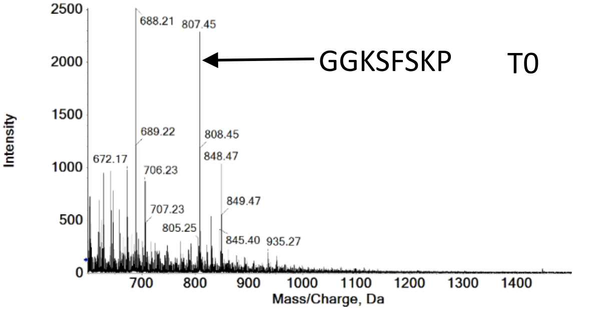
GGKSFSKP
T0
688.21
807.45
689.22
808.45
848.47
672.17
706.23
849.47
707.23
805.25
845.40
935.27
Intensity
Mass/Charge, Da

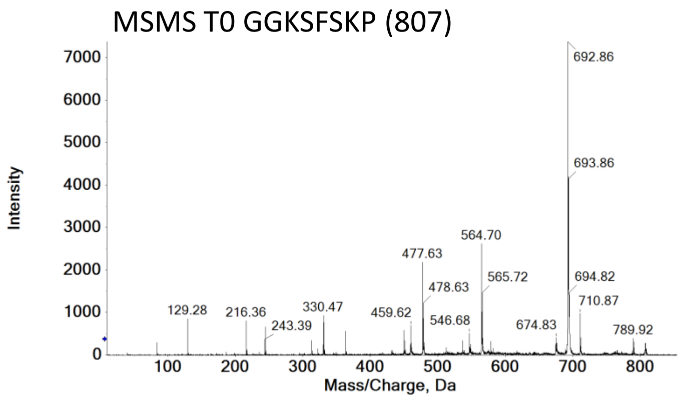
MSMS T0 GGKSFSKP (807)
692.86
693.86
564.70
477.63
478.63
565.72
694.82
710.87
129.28
216.36
330.47
243.39
459.62
546.68
674.83
789.92
Intensity
Mass/Charge, Da

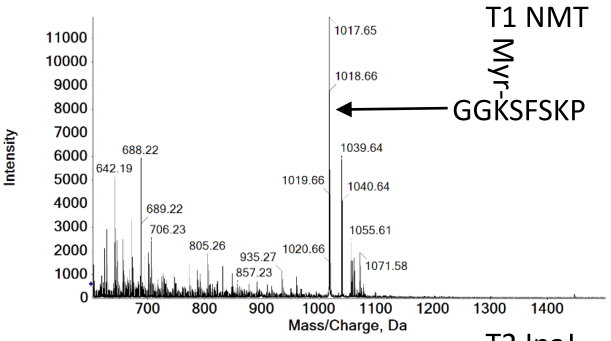
T1 NMT
Myr-
GGKSFSKP
1017.65
1018.66
1039.64
1040.64
1019.66
688.22
642.19
689.22
706.23
805.26
935.27
857.23
1020.66
1055.61
1071.58
Intensity
Mass/Charge, Da

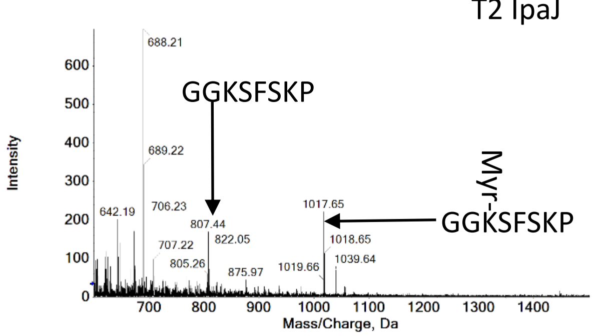
T2 IpaJ
GGKSFSKP
Myr-
GGKSFSKP
688.21
689.22
706.23
642.19
807.44
707.22
822.05
805.26
875.97
1017.65
1018.65
1039.64
1019.66
Intensity
Mass/Charge, Da

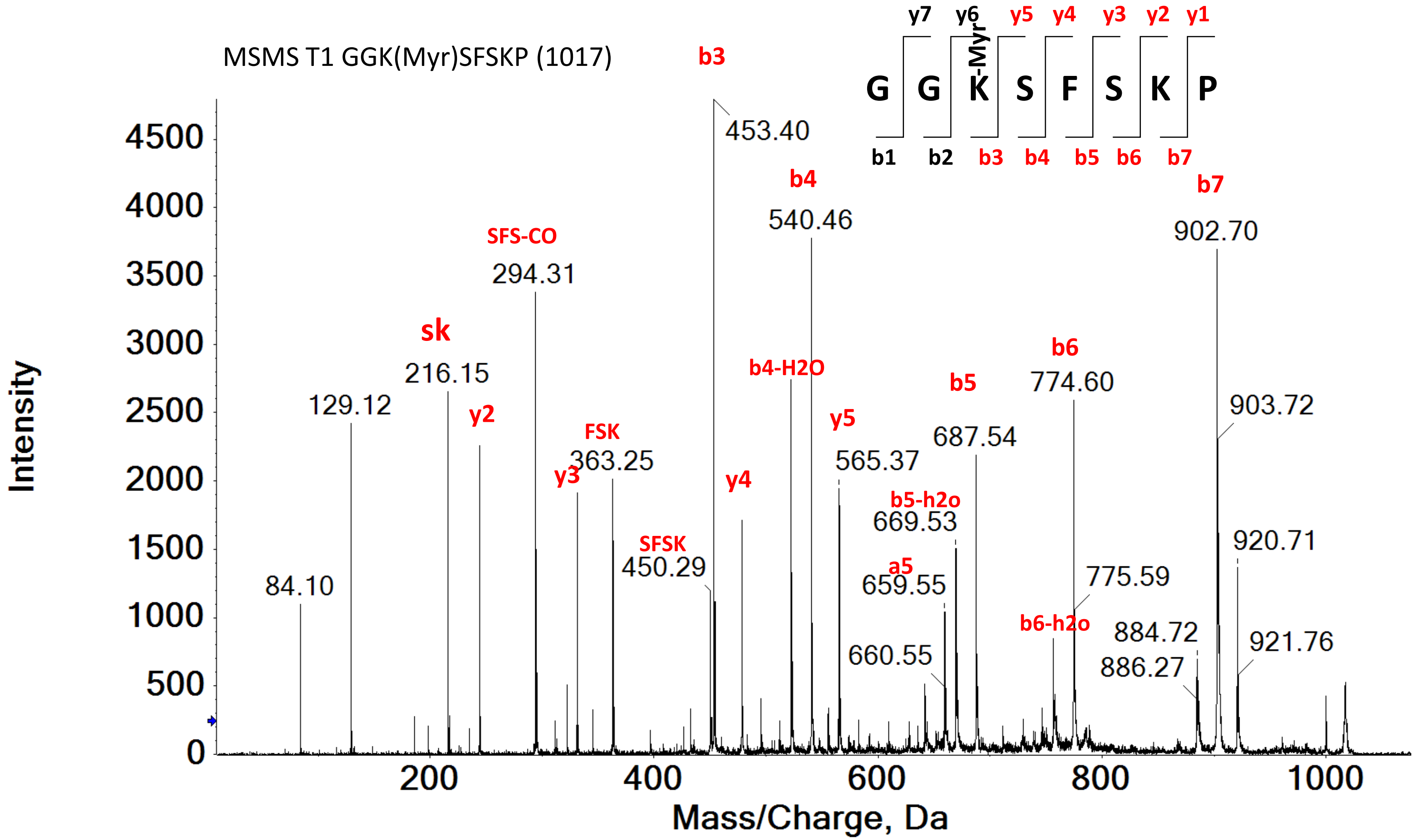
MSMS T1 GGK(Myr)SFSKP (1017)
y7 y6 y5 y4 y3 y2 y1
G G K S F S K P
-Myr
b1 b2 b3 b4 b5 b6 b7
b3
453.40
b4
540.46
b7
902.70
SFS-CO
294.31
sk
216.15
129.12
y2
b4-H2O
b6
774.60
b5
903.72
687.54
y5
565.37
FSK
363.25
y3
y4
b5-h2o
669.53
SFSK
450.29
a5
659.55
920.71
775.59
84.10
b6-h2o
884.72
886.27
921.76
660.55
Intensity
Mass/Charge, Da

### GGKFSSKPR

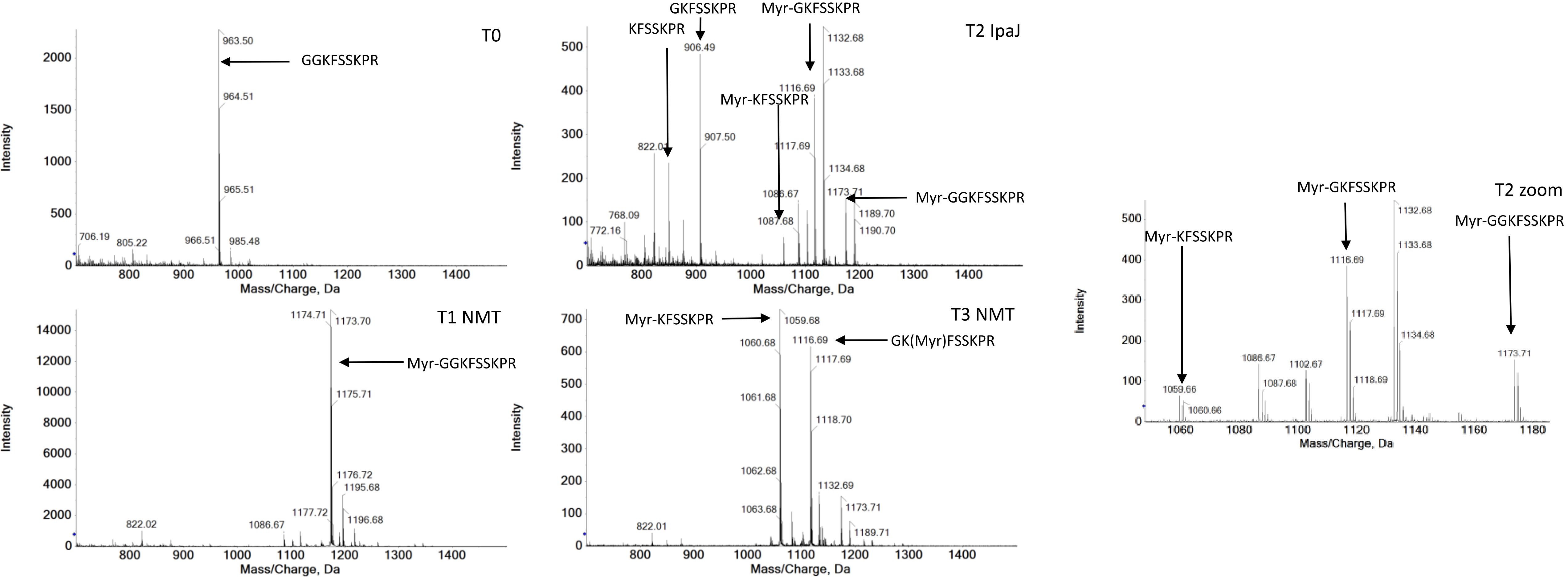
T0
GGKFSSKPR
963.50
964.51
965.51
966.51
985.48
706.19
805.22
Intensity
Mass/Charge, Da
T2 IpaJ
GKFSSKPR
Myr-GKFSSKPR
KFSSKPR
Myr-KFSSKPR
Myr-GGKFSSKPR
906.49
907.50
822.01
768.09
772.16
1086.67
1087.68
1116.69
1117.69
1132.68
1133.68
1134.68
1173.71
1189.70
1190.70
T1 NMT
Myr-GGKFSSKPR
1174.71
1173.70
1175.71
1176.72
1195.68
1196.68
1177.72
1086.67
822.02
T3 NMT
Myr-KFSSKPR
GK(Myr)FSSKPR
1059.68
1060.68
1061.68
1062.68
1063.68
1116.69
1117.69
1118.70
1132.69
1173.71
1189.71
822.01
T2 zoom
Myr-GKFSSKPR
Myr-KFSSKPR
Myr-GGKFSSKPR
1059.66
1060.66
1086.67
1087.68
1102.67
1116.69
1117.69
1118.69
1132.68
1133.68
1134.68
1173.71

### GGKFSSKPR

MSMS T0 GGKFSSKPR (963)

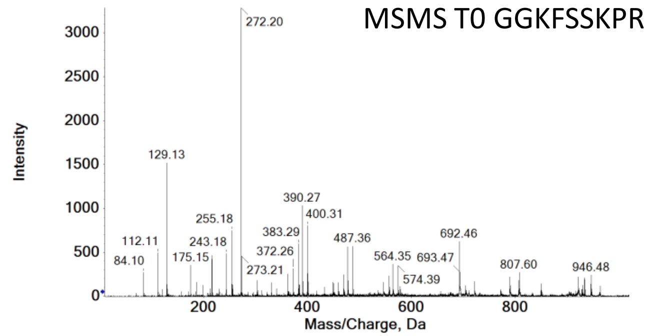


MSMS T3 GK(Myr)FSSKPR (1116)

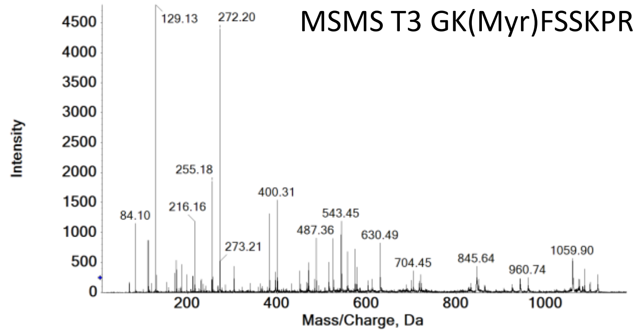


MSMS T3 Myr-KFSSKPR (1059)

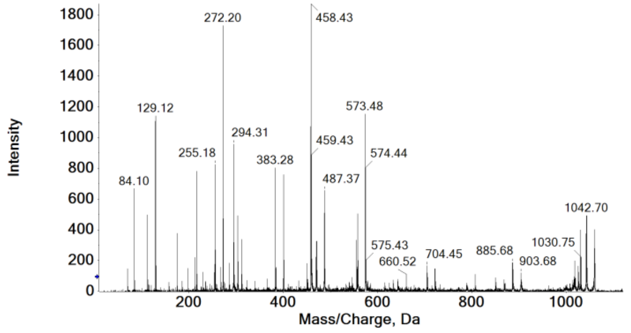


MSMS T0 Myr-GGKFSSKPR (1173)

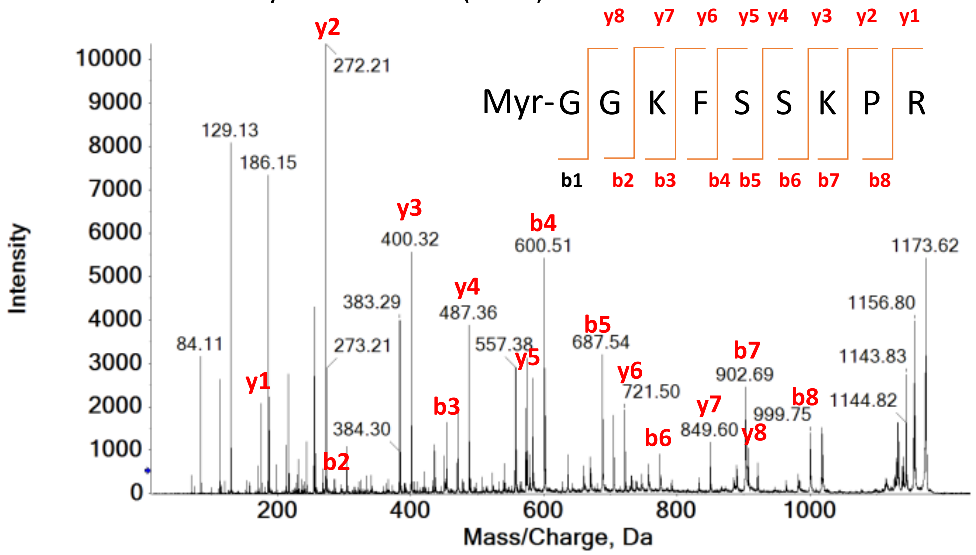


| K20 1173 | Mass/Charge (Da) | Intensity | |
|---|---|---|---|
| 4372 | 175,1503 | 2072 | y1 |
| 8355 | 272,1954 | 10360,16 | y2 |
| 10293 | 325,2772 | 320 | b2 |
| 12791 | 400,3083 | 5570,51 | y3 |
| 14427 | 453,4173 | 1645,961 | b3 |
| 15423 | 487,3767 | 3883,765 | y4 |
| 17825 | 574,4145 | 3119,059 | y5 |
| 18508 | 600,4936 | 5425,726 | b4 |
| 20684 | 687,5204 | 3206,745 | b5 |
| 21495 | 721,4933 | 1923,922 | y6 |
| 22724 | 774,5721 | 912,7844 | b6 |
| 24390 | 849,6013 | 1189,02 | y7 |
| 25523 | 902,6548 | 2460,706 | b7 |
| 25606 | 906,606 | 1046,588 | y8 |
| 27511 | 999,7244 | 1402,039 | b8 |

MSMS T3 GK(Myr)FSSKPR (1116)

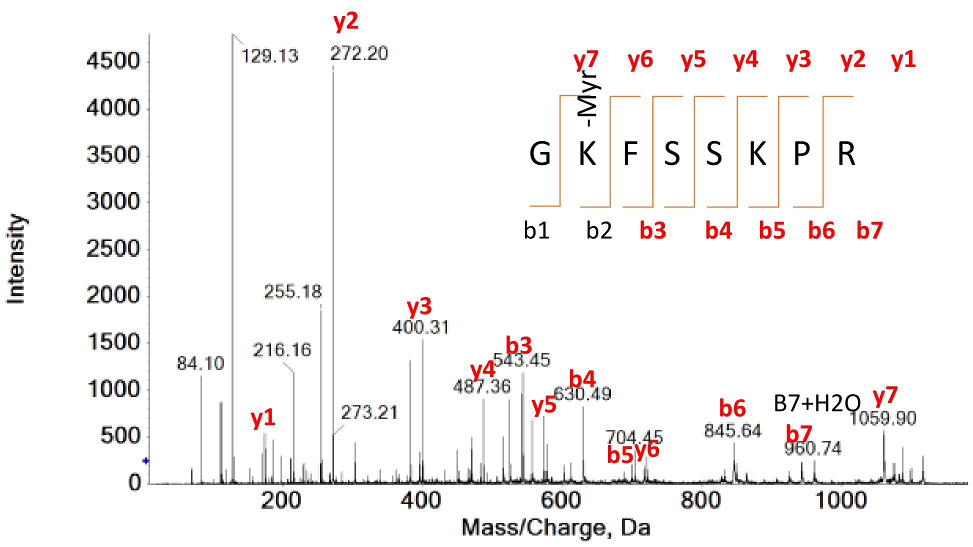


| K22 1116 | Mass/Charge (Da) | Intensity | |
|---|---|---|---|
| 2008 | 175,1541 | 538,35 | y1 |
| 2891 | 216,1627 | 1182,90 | SK |
| 3999 | 255,1691 | 1852,24 | y2-NH3 |
| 4553 | 272,2057 | 4394,67 | y2 |
| 5462 | 303,1855 | 437,33 | SSK |
| 7699 | 383,2735 | 1310,43 | SSKP-NH3 |
| 8218 | 400,3085 | 1541,02 | y3 |
| 9657 | 450,2645 | 359,53 | FSSK |
| 10244 | 470,3054 | 500,24 | y4-NH3 |
| 10729 | 487,3455 | 905,41 | y4 |
| 11494 | 515,4305 | 506,04 | a3 |
| 12266 | 543,4355 | 1191,69 | b3 |
| 13097 | 574,384 | 724,55 | y5 |
| 14532 | 630,4798 | 822,75 | b4 |
| 16634 | 717,5125 | 190,59 | b5 |
| 16727 | 721,4527 | 298,67 | y6 |
| 19445 | 845,6232 | 432,31 | b6 |
| 21453 | 942,7505 | 231,69 | b7 |
| 21812 | 960,7414 | 246,90 | b7+H2O |
| 23625 | 1059,864 | 569,10 | y7 |

### AKPTSKDSGLK

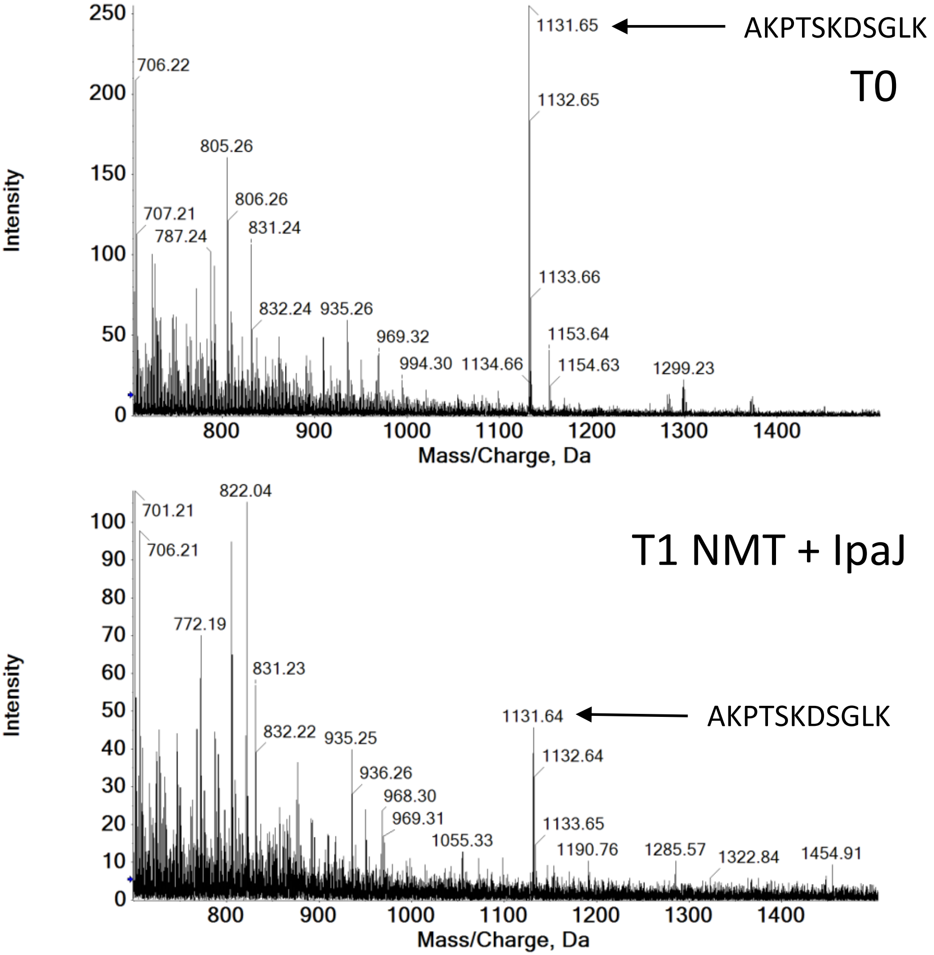

### AGKFSKPR

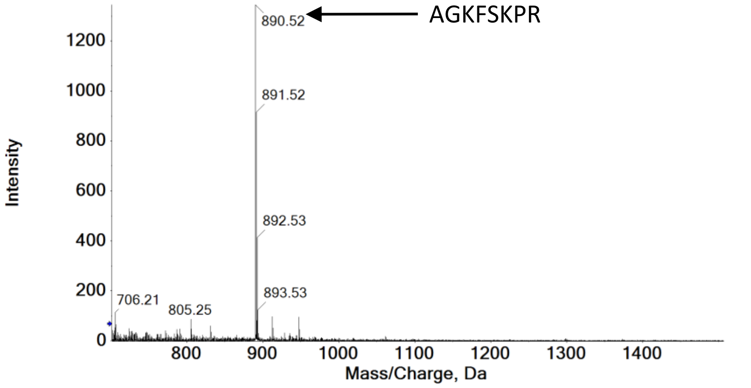


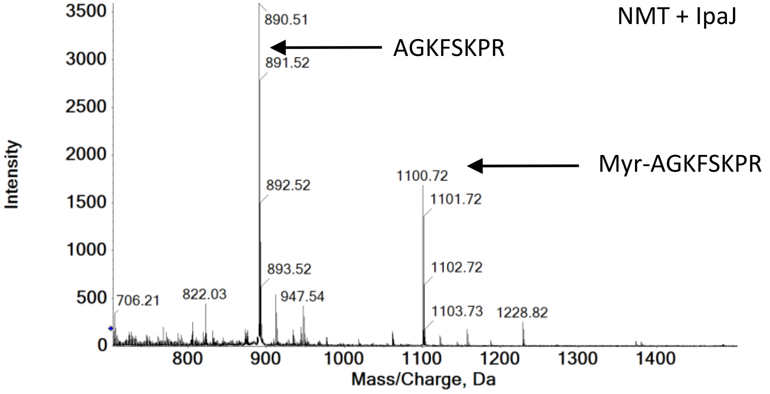


MSMS T0 AGKFSKPR (890)

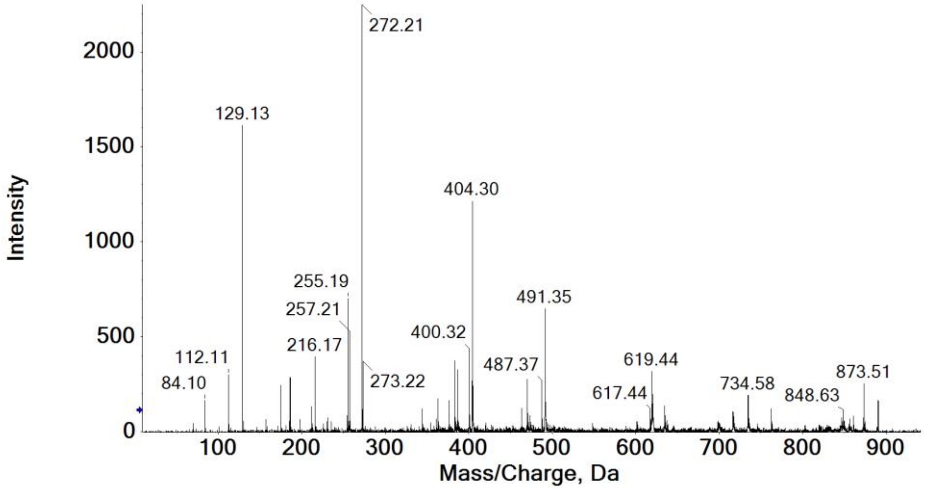


MSMS T1 Myr-AGKFSKPR (1100)

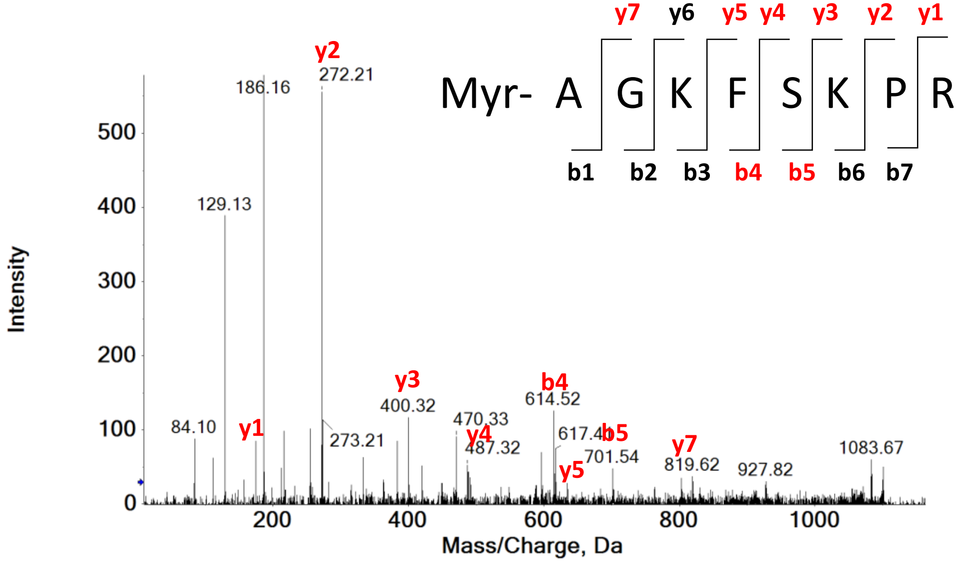


| L4 1100 | Mass/Charge (Da) | Intensity | |
|---|---|---|---|
| 542 | 129,138 | 388,8628 | K |
| 801 | 175,1467 | 84,86275 | y1 |
| 894 | 186,1513 | 578,353 | GK |
| 1149 | 216,181 | 98,66667 | SK |
| 1590 | 272,2057 | 555,9216 | y2 |
| 2024 | 333,2545 | 63,21569 | GKF |
| 2715 | 400,3234 | 116,8627 | y3 |
| 2896 | 420,3111 | 51,45098 | GKFS |
| 3333 | 470,3304 | 90,82353 | y4-NH3 |
| 3503 | 487,3036 | 52,07843 | y4 |
| 4591 | 614,5147 | 126,1177 | b4 |
| 4836 | 634,4205 | 28,07843 | y5 |
| 5421 | 701,5098 | 47,68628 | b5 |
| 6483 | 819,5577 | 37,17647 | y7 |

### AGKSFSKPR

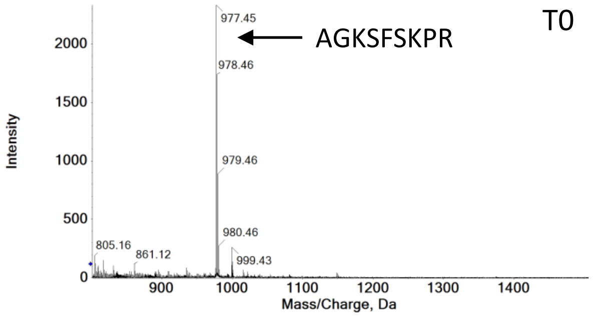


L12

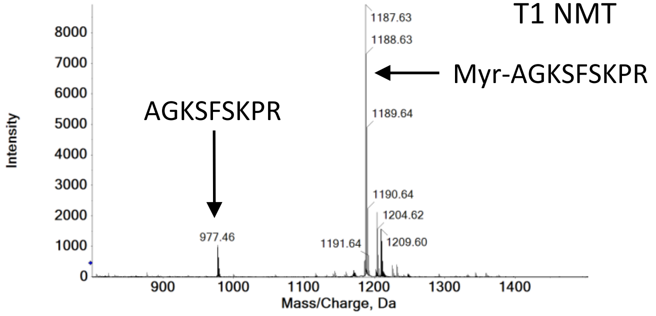


L13

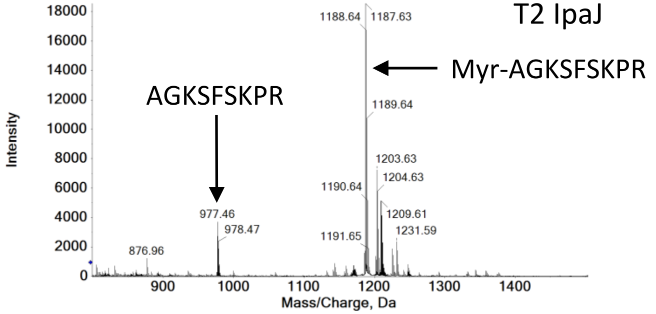


L14

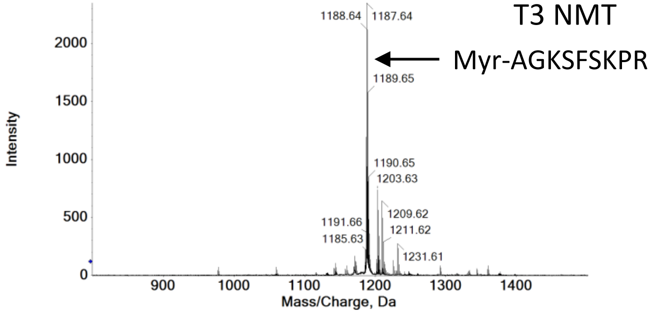


L15

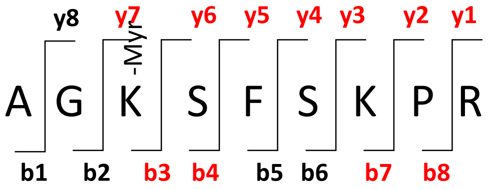


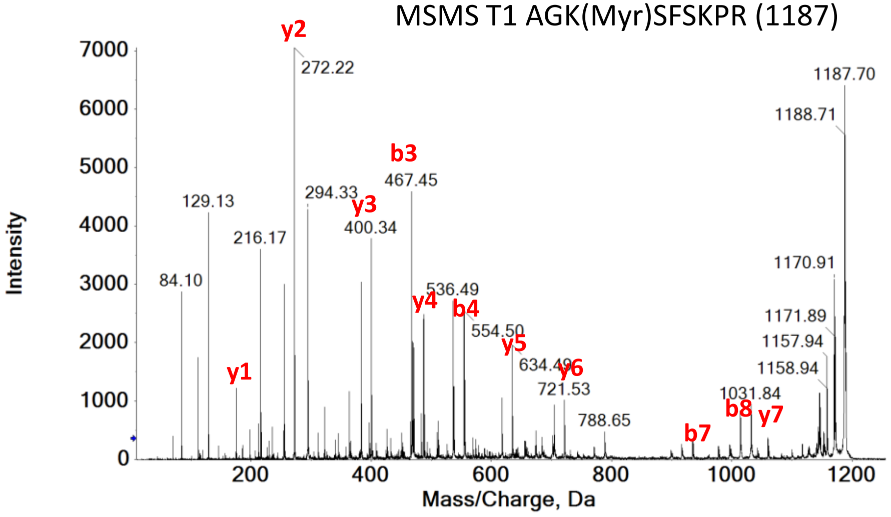


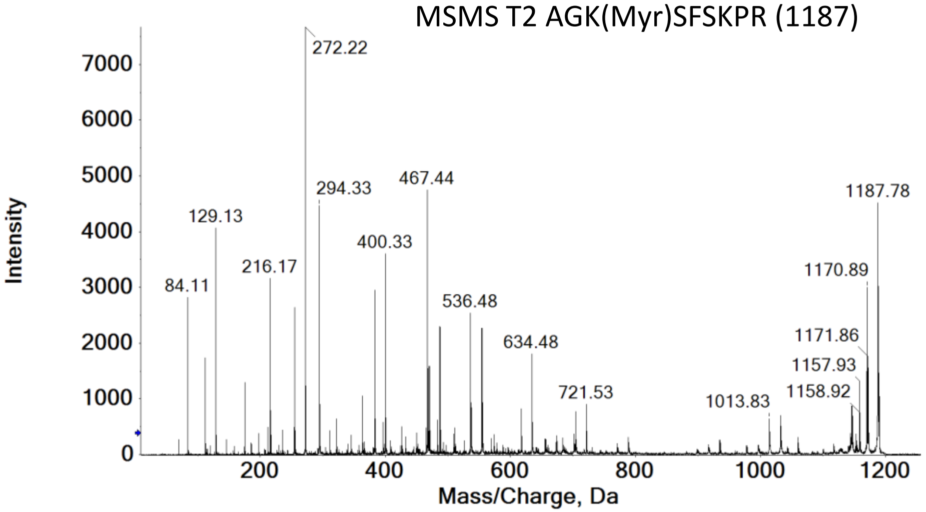
